# Protocell formation on micrometeorites

**DOI:** 10.1101/2025.03.31.646313

**Authors:** Aldo Jesorka, Esteban Pedrueza Villalmanzo, Ezgi Ciftcioglu, Piotr Jedrasik, Jon Larsen, Irep Gözen

## Abstract

We report on the formation of membranous protocells by self-assembly of lipids on micrometeorites, the extraterrestrial particles that have been continuously reaching the surface of the Earth ever since its formation. Synergistic interactions of lipid compartments with pristine extraterrestrial surfaces are entirely unexplored, but constitute a possible scenario for early evolution of primitive cells by a surface energy-driven transformation mechanism. Lipids utilize the surface energy of the particles to adhere to them and autonomously transform into spherical compartments, typically through formation of lipid nanotubes. Natural sand particles of similar composition and shape were simultaneously investigated for reference, showing that certain lipid compositions prefer micrometeorite surfaces. The elemental composition of the particles, their surface texture and cleanness altogether may be contributing to the differences observed in lipid behavior. Lipid nanotubes on- and extending out of- the micrometeorites were observed to carry lipid particles and connect to other objects in the surrounding environment.

## Introduction

Micrometeorites are extraterrestrial particles, 10-2000 μm in size, originating from asteroids and comets. They are a subset of cosmic dust particles, i.e., interstellar dust. 40±20 kilotons shower our planet every year^1^, of which according to van Ginneken et al.^2^, only 10% survive the atmospheric entry. There are many different types of micrometeorites of varying composition and structure. Specimens were recovered from direct atmospheric sampling^3^, surface ice and snow of the Antarctic^4^, glacial sediments, deep-sea sediments, deserts, and most recently from the rooftops in urban environments^2, 5^.

Various types of organic matter were detected in micrometeorites, such as polycyclic aromatic hydrocarbons^6^, nitrogen-rich organic matter ^7-8^ and even amino acids^9^, although their ability to remain intact is debated^10^. It has been suggested that on the early Earth, micrometeorites could have been a significant source of organic material and could have functioned as microscopic chemical reactors for the synthesis of prebiotic molecules^8, 11^. This is important in the context of abiogenesis, i.e., the origin of life, at which the assembly of prebiotic molecules is hypothesized to have led to the formation of protocells, precursors of the first biological cells. The possible synergistic combination of such natural, self-supplied chemical microreactors with dynamic membrane compartments in direct contact with their surface has never been considered as initiator of a viable development pathway at the origin of life.

Experimental work focusing on protocells at the origin of life employs physical soft matter model structures^12-13^. One common physical protocell model, among coacervates, is the giant unilamellar lipid vesicles (GUVs)^12^. GUVs are lipid compartments suspended in water enveloping an aqueous volume with a continuous spherical lipid bilayer, similar to the membranes surrounding contemporary biological cells. We have previously performed extensive work on protocell transformations on *terrestrial* solid rock and mineral surfaces and a Martian meteorite specimen, and formulated the hypothesis that surface energy contributed to the transformation of lipid agglomerates to primitive cells under straightforward, early Earth-compatible assumptions. Other groups also reported earlier that terrestrial mineral nano and microparticles induced the formation of lipid vesicles from fatty acid micelles^14-16^.

The intrinsic energy of surfaces is large enough to induce the self-assembly of lipid compartments into versatile morphologies, e.g. lipid nanotube-compartment networks^17-20^ and foam-like structures^21-23^ reminiscent of microbial colonies. These non-trivial protocell morphologies possess certain advantages compared to the GUVs freely suspended in an aqueous environment. For example, they can directly transport molecules, e.g. RNA or DNA via nano-tunnels between them^24^, or withstand osmotic pressures and remain intact while in the same conditions isolated GUVs immediately disintegrate^21^.

Here, we investigated the interaction of micrometeorite surfaces with archaeal and bacterial lipids as well as reference lipid mixtures of the exact composition we utilized in earlier studies; specifically the protocell growth on micrometeorite specimens collected from rooftops in urban areas. Three micrometeorites: porphyritic (PO), scoriaceous (ScMM) and barred olivine (BO) types, furthermore regular untreated, as well as oxygen plasma-exposed sand particles for reference were subjected to several different lipid types. Some of lipid compositions were found to favor protocell assembly on micrometeorites compared to other surfaces, where some consistently showed weak interactions with the micrometeorites. Lipid spreading, rupturing and formation of the nanotube networks have been observed on the micrometeorites, suggesting formation pathways similar to the ones earlier found to occur on terrestrial surfaces^17-19^. The nanotubes on- or extending out of- the micrometeorites were observed to carry lipid agglomerates and connect to other objects and surfaces in the surrounding solution.

If there were special life-supporting environments on the early Earth, which we assume existed^25^, micrometeorites would have landed on these locations. This is important because micrometeorites a) are uncontaminated pristine surfaces, b) have unique surface texture, c) often contain catalytically relevant metals, d) may contain organic molecules, e) are mobile. Furthermore, since cosmic dust also reaches other rocky planets^26-27^ and exoplanets^28^, the characterization of primitive compartments on micrometeorites provides clues not only about the origin of life on Earth, but also about the possibility of life on other planetary bodies with similar environments.

## Results and Discussion

The schematic drawing in **Figure 1** gives an overview of our experimental design. We used three micrometeorites: PO, ScMM and BO, collected from roof tops^2, 29^. The details of the micrometeorite recovery from roof top dust are described by Larsen et al.^5, 29^. Briefly, the material collected from roof tops in the Oslo area (Norway, 2021) was subject to magnetic separation, cleaning and size separation using sieves. Micrometeorites were manually further separated from industrial particles of similar size by means of microscopy. For the experiments, we placed the micrometeorite specimens under a stereomicroscope into glass bottom dishes containing an aqueous buffer (cf. Materials&Methods for details). We then added a lipid suspension and left the sample overnight. The subsequent imaging was performed with an inverted fluorescence microscope.

**Figure 1.**
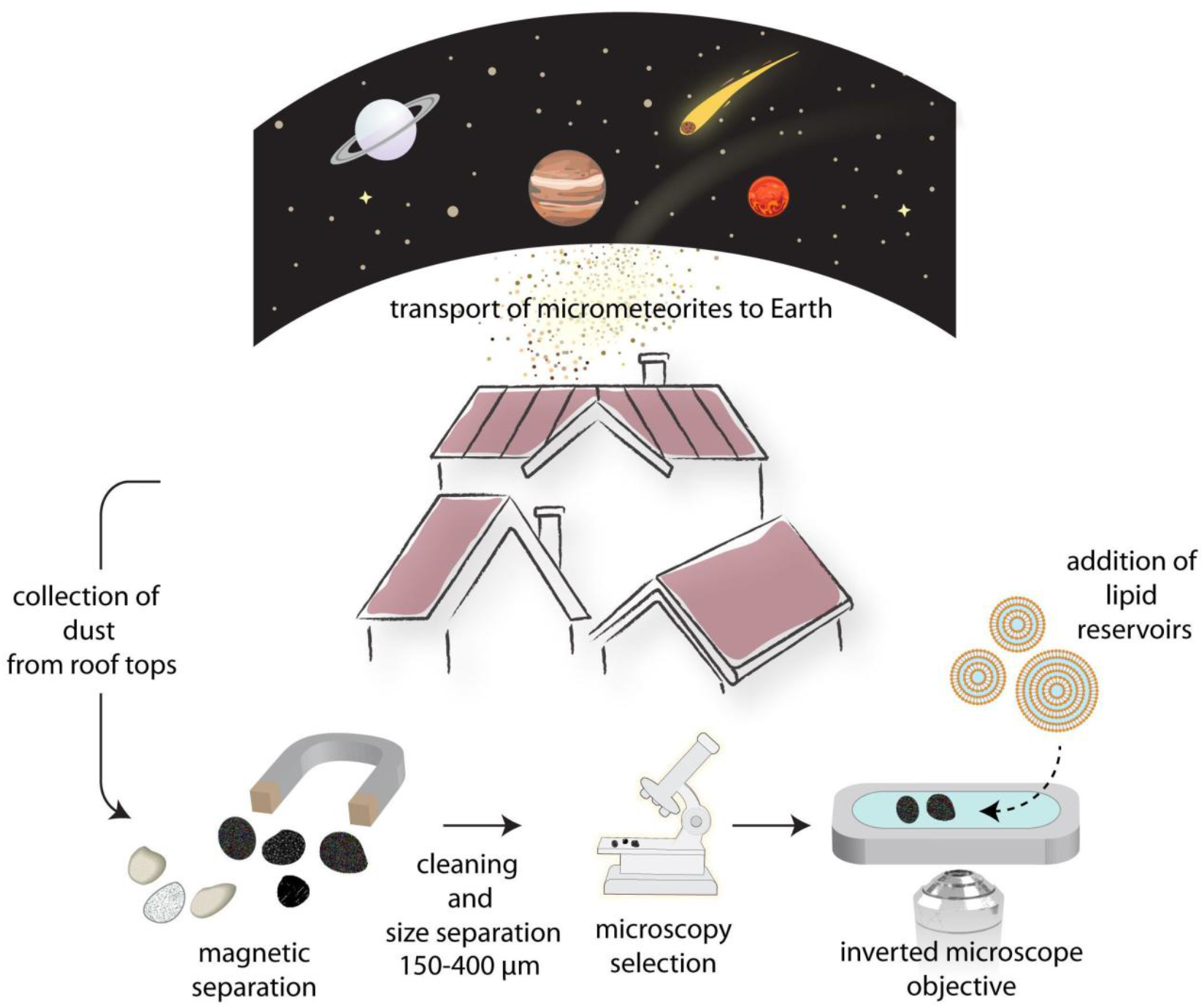
Schematic description of the experimental design, including the initial step of micrometeorite collection from urban environments. Debris from roof tops containing micrometeorites is collected, and all magnetically responsive particles are separated with the help of a permanent magnet. These magnetic particles can contain micrometeorites, but other natural and man-made particulates dominate the roof top dust. Sieves are used to eliminate large particles in the millimeter range. The remaining small particles are examined under a stereo microscope and particles carrying the unique characteristics of micrometeorites are collected. The micrometeorites are then transferred into an open-top sample chamber containing aqueous medium, followed by the addition of lipid reservoirs/agglomerates. The sample becomes ready for fluorescence microscopy observation after allowing the interaction of lipids with micrometeorite surface overnight.

Every lipid suspension was brought in contact with three different sets of particles. The first set consists of the micrometeorites. Before every experiment, the micrometeorites were cleaned with solvents and exposed to oxygen plasma. We also used two different sets of terrestrial reference particles besides the micrometeorites. Both sets consist of particles retrieved from a sand sample collected from a beach in Vrångö Island near Göteborg, Sweden. The terrestrial reference particles were chosen to match the characteristics of the micrometeorites as much as possible. We selected magnetic, i.e., iron-rich particles of similar size and shape. Magnetic particles within the sand sample, amounting to ∼10% of the total population, were separated with a neodymium magnet. Suitable particles were identified from this subset, and the remaining material was discarded.

Cosmic dust experiences plasma in space^30^ and meteorites produce a dense plasma layer around them upon atmospheric entry^31-32^. In order to simulate the physical conditions of meteorites arriving Earth, we exposed one set of magnetic sand particles to oxygen plasma, termed the ‘model meteorites’. Similar to the procedure applied to the micrometeorites, these magnetic particles were rinsed with solvents and oxygen plasma-treated before the experiments. We note that some micrometeorites experience low entry angle and/or grazing entry which leads to lower temperature, possibly without plasma formation (cold capture)^10^.

Besides real and model micrometeorites, several additional natural magnetic sand particles without any plasma treatment were used as a second set of controls. These sand particles were directly brought in contact with lipid suspensions. While the micrometeorites and the model micrometeorites were used repeatedly only altering the lipid compositions, a different batch of sand particles were used for each lipid suspension, as the sand particles contaminated with one type of lipid could not be used without cleaning for another experiment involving a different lipid suspension.

We used three different lipid compositions: diether lipids characteristic of archaeal membranes^33-34^, lipids derived from *E*.*coli* bacteria, and a reference mixture containing soy bean plant lipids and *E*.*coli* lipids for direct comparison to previous work where this mixture was studied on solid surfaces. The lipid compositions were selected due to their representation of different domains of life. The archaeal lipids contain ether bonds attaching isoprenoids to glycerol 1-phosphate, where bacteria and eukaryotes have ester bonds linking fatty acids to glycerol 3-phosphate^35^. All of the lipid compositions were fluorescently labeled for imaging.

Scanning electron microscope (SEM) images and compositional characterization data of the micrometeorites we used in our experiments are presented in **Figure 2. Fig. 2A-C** show the SEM images of the whole particles and **Fig. 2D-F** show close-ups of the particles corresponding to panels (A-C). **Fig. 2G-I** show the compositional analyses of the micrometeorites in (A-F) performed by energy dispersive X-ray spectroscopy (EDX). The table below each spectrum shows the elements present in each micrometeorite as obtained from the spectroscopy data. Mg, Si and Fe comprise the majority (89-90%) of the elemental make-up of the particles.

**Figure 2.**
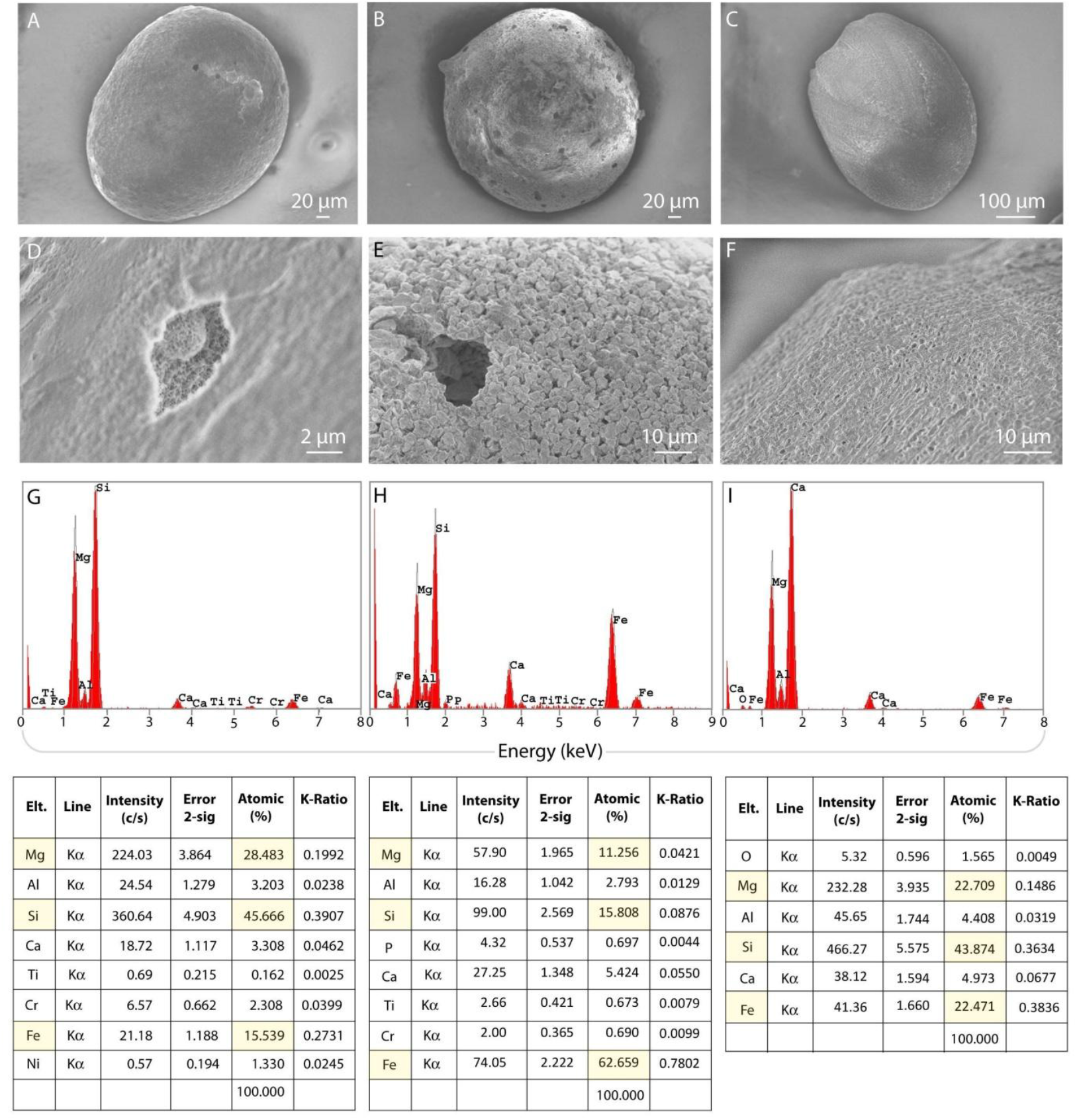
Scanning electron microscopy (SEM) and energy dispersive X-ray spectroscopy (EDX) characterization of the micrometeorites. (A-C) SEM images of the micrometeorites. (D-F) close up SEM images corresponding to (A-C). A: Porphyritic (PO), B: scoriaceous (ScMM), C: barred olivine (BO) (G-I) SEM-EDX spectra of the meteorites (intensity in counts versus energy in keV). Element assignments for each peak were provided by the instrument. Quantification of the elemental composition is presented below each spectrum. Elements which constitute more than 10% of the total are highlighted in the tables. Oxygen was not determined.

SEM images and the compositional analyses of all other particles used in our experiments were also obtained; they are presented in the Supporting Information (SI). When compared to all other particles, the shape, especially the surface textures of the micrometeorites are unique (**Fig. 2D-F**). This is especially apparent in **Fig. 2E** which shows the typical texture of a scoriaceous micrometeorite. Unlike terrestrial particles, micrometeorites go through a formation process (atmospheric entry - frictional heating - deceleration/solidification/re-crystallization) with entry speeds greater than 11 km/s. The micrometeorites are classified^36^ according to the different peak temperatures they experience during atmospheric entry and flash heating (BO <1800 °C, PO <1600 °C, and ScMM <1350 °C). These are surface temperatures, all three types may contain relict (unmelted) grains inside. As a result, the textures commonly found on fresh micrometeorites are fundamentally different from particles formed on Earth. Furthermore, due to low peak temperature during formation, sc micrometeorites have a magnetite (Fe_2_O_3_/Fe_3_O_4_) surface layer.

The purpose of the experiments was to investigate the possible interactions between the lipid material and the particle surfaces, identify areas of lipid attachment, film formation, shape transformations and compartment formation. An obvious obstacle is the large particle size and the lack of optical transparency. Imaging of lipid structures requires high magnification, which does not allow for snapshots of the entire particle. Therefore, surface regions were imaged at different focal planes. Transparency is a comparatively minor problem, since the fluorescently labeled lipids provide the necessary contrast.

**Figure 3** shows self-assembled protocells on sand particles (A-C), model micrometeorites (D-F) and micrometeorites (G-I). Although not in the same quantities, all surfaces were able to generate lipid compartments. Some surfaces were densely populated, while others had only very few attachments. Below each panel, a plot is shown which corresponds to the fluorescence intensity along the arrows in the respective micrograph above the plot. These M-shaped fluorescence intensity graphs are characteristic to GUVs^37-38^. Two exceptions are the graph in panel (A) where lipid agglomerates inside the vesicle cause the internal fluorescence to exceed the membrane fluorescence at the vesicle periphery, and panel (E), where on one side the vesicle is adjacent to another vesicle, such that the fluorescence intensity reflects the presence of two bilayer membranes.

**Figure 3.**
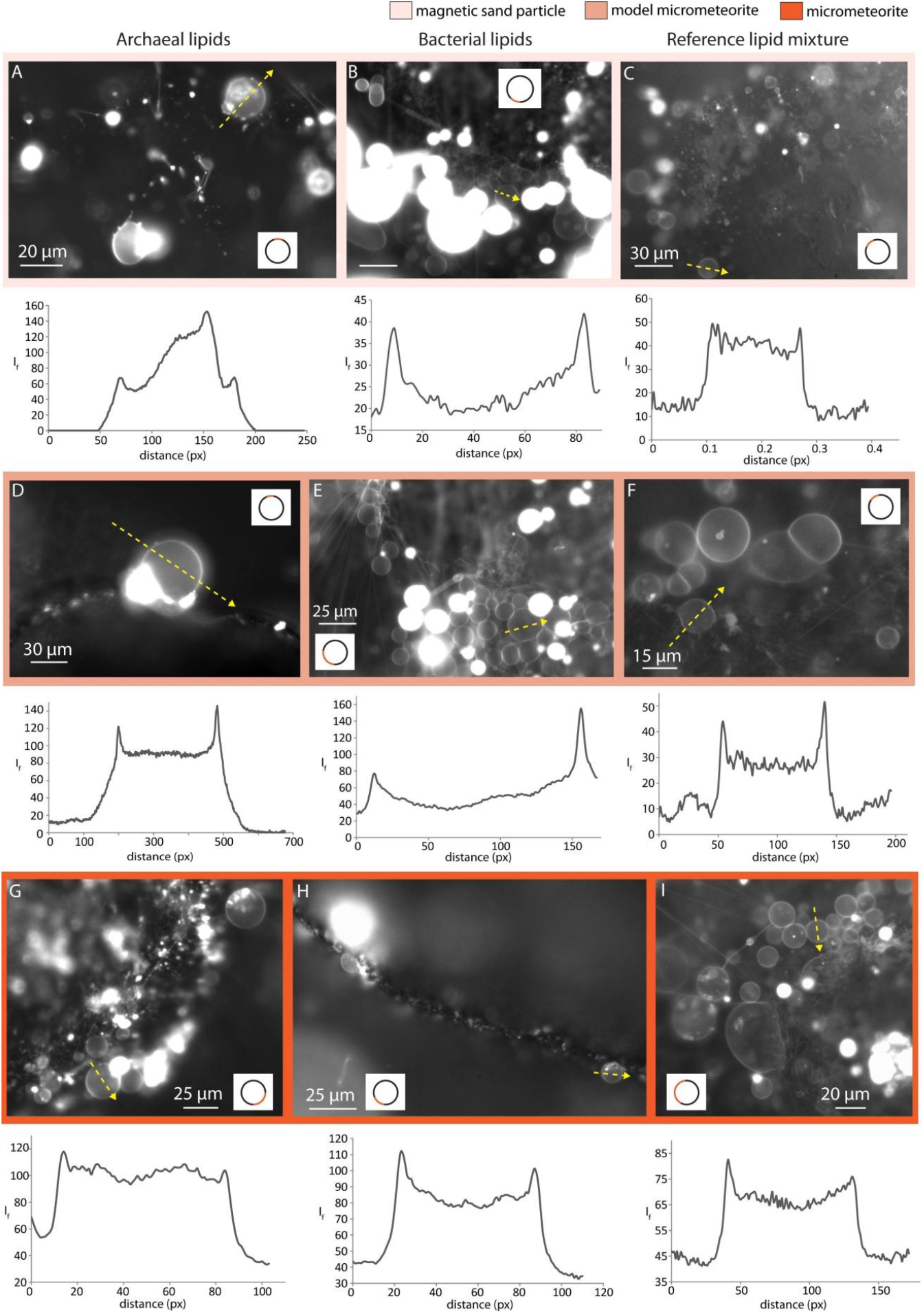
Fluorescence micrographs of protocells assembling on (A-C) sand particles, (D-F) model micrometeorites, (G-I) micrometeorites. The columns correspond to protocells composed of different lipid species: left: archaeal lipids, middle: *E*.*coli* lipids, right: reference lipid mixture. Below each micrograph, a fluorescence intensity plot corresponding to the arrow (yellow dashed line) across a lipid compartment in the microscopy image above is presented (units: gray value from 0 to 255 of 8-bit image, versus distance in pixels). The plots adopt M-shaped fluorescence intensity profiles, which are typical for GUVs. Deviations in panels A and E are explained in the text. In each image, an inset marks the approximate position of the protocells on the particle with an orange line on a circle.

The images suggest that the protocells form on micrometeorites by the same mechanism as established for protocells on planar terrestrial mineral and rock surfaces^17-19^, which is schematically described for reference in **Figure 4A-D**. A multilamellar lipid vesicle (MLV) spreads as a double lipid bilayer, i.e., a flat GUV, on the planar solid surface. The distal bilayer (upper lipid bilayer with respect to the surface) ruptures due to tension emerging from continuous expansion on the surface (**Fig. 4B**), and transforms into a network of lipid nanotubes (**Fig. 4C**). Some sections of the lipid nanotubes swell over time, forming a lipid nanotube-vesicle network (**Fig. 4D**). Note that lipid bilayers in aqueous environments do not have open edges; the section view is provided for better understanding of the illustrated membrane structures.

**Figure 4.**
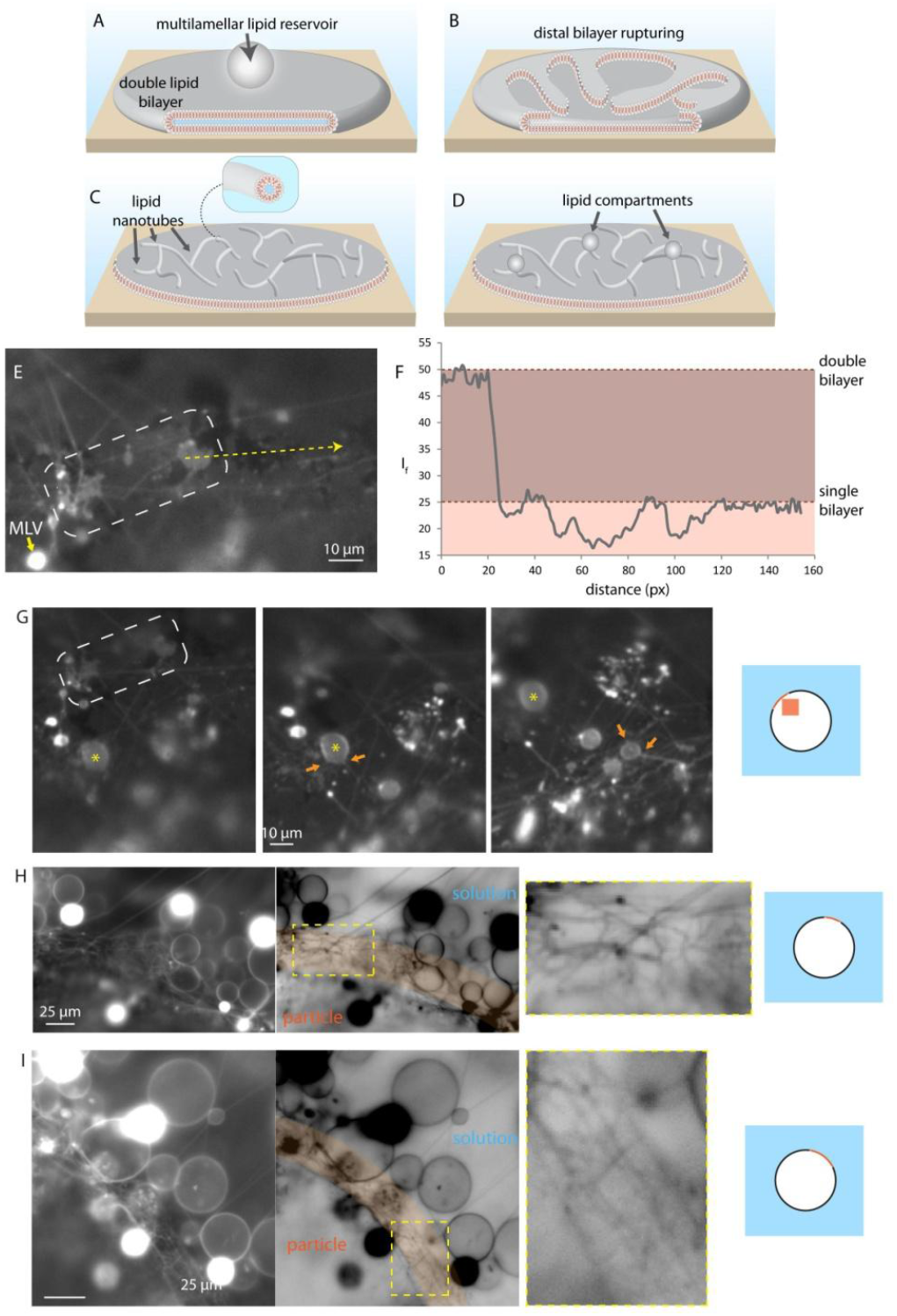
Protocell formation mechanism. (A-D) Formation of nanotube-lipid compartment networks on flat surfaces in aqueous environments. (A) Multilamellar lipid reservoirs, i.e., multilamellar vesicles (MLV) spread on high energy surfaces as a double lipid bilayer in the form of a flat giant unilamellar vesicle. (B) The distal lipid membrane (upper with respect to the surface) ruptures, (C) forming a lipid nanotube network. The inset to C shows the cross section of a lipid nanotube in the network. (D) Lipid compartments emerge from the lipid nanotubes. (E-I) Fluorescence microscopy images of protocells growing on model micrometeorites (plasma-treated sand particles) and micrometeorites. (E) Fluorescence micrograph of an archaeal lipid patch on a micrometeorite. The short yellow arrow points to the multilamellar reservoir. (F) Fluorescence intensity graph along the arrow in (E) (gray value from 0 to 255 of 8-bit image, distance in unit of pixels). The recorded 1:2 intensity ratio indicates single:double lipid bilayer membranes on the surface. (G) shows the vicinity of the region in (E) at three slightly different focal planes, where several lipid compartments grow out of the lipid nanotubes. The framed regions in (E) and (G) are identical. The same lipid compartment in different focal planes is marked with a yellow asterisk in (G). The nanotubes directly connected to the lipid compartments are shown with orange arrows. (H-I) fluorescence micrographs of bacteria-derived (H) and reference (I) lipids on a model micrometeorite along with inverted images for improved contrast. Magnified regions densely populated with lipid nanotubes are shown in yellow frames. The regions in orange color underscore the parts of the curved particle surface which are in focus. Blue squares attached to the images show schematically the approximate position of the protocells on the corresponding particle.

Since the micrometeorites are spherical and there is no single focal plane, it is challenging to capture this whole dynamic process on the particles step-by-step as we did earlier on planar surfaces^19^. However, we recorded regions of intact (double) and ruptured (single) lipid membranes on micrometeorite surface regions (**Fig. 4E-F**) as well as very dense surface-bound nanotube networks and compartments connected to them (**Fig 4G-I**).

The surface coverage density of the membranous protocells and the lipid nanotubes, i.e., their quantity per unit surface area, on all types of particle surfaces is presented in **Figure 5**. For the 9 different lipid– particle combinations, the formation of protocells of a given lipid composition was imaged three times on three particles of each particle type: i) sand particles, ii) model micrometeorites, iii) micrometeorites. For each particle, the three images were acquired from three different surface regions. The quantification results from each image, amounting to a total of 3×27 different data points, are represented as individual bars in the graphs in **Fig.5**.

**Figure 5.**
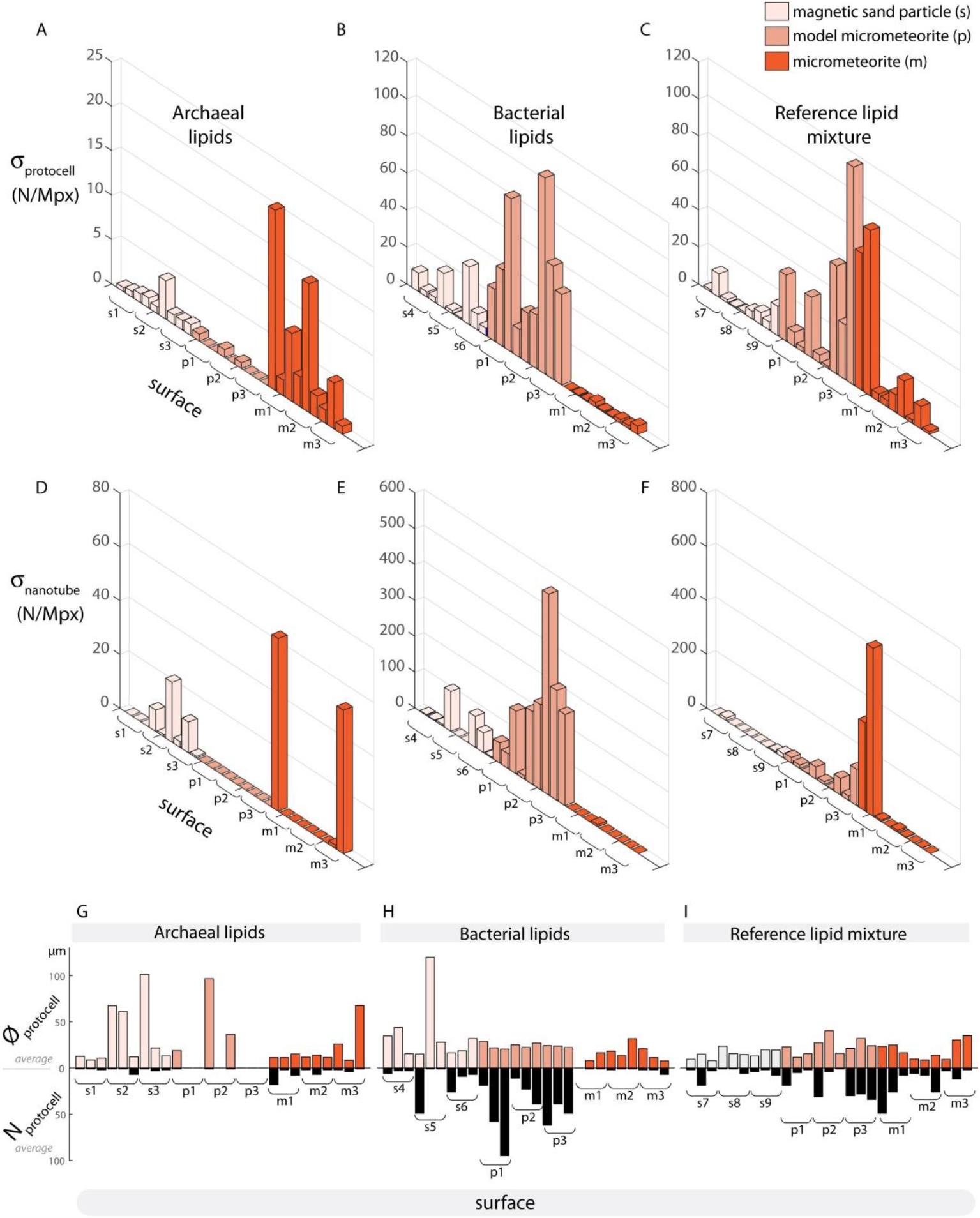
Analyses of protocells on particles. (A-F) Bar graphs showing protocell surface coverage (density: number of protocells per unit area, N/megapixels) on 27 different surface regions: 9 for each surface type, triplets of images for each particle type: s (sand), p (model micrometeorite particle), m (micrometeorite). (G-I) Top: bars representing the average diameter of protocells on each surface, bottom: bars representing the average number of protocells corresponding to that surface. Panels (A,D,G) represent protocells composed of archaeal lipids, panels (B,E, H) of bacterial (*E*.*coli*) lipids, and panels (C, F, I) of reference lipids containing plant- and bacteria-derived lipids.

One surprising result is that the archaeal lipids formed protocells preferably on the micrometeorites, whereas on sand particles and model micrometeorites their surface coverage density was much reduced (**Fig. 5A**). Lipids derived from the *E*.*coli* membranes on the other hand, appear to have avoided the micrometeorite surfaces and form protocells more densely on sand particles and model micrometeorites (**Fig. 5B**). No strong preference is discernible regarding the protocell formation from the reference lipid mixture, but the model- and actual micrometeorites are more densely populated (**Fig. 5C**). The three lipid types used are structurally different. In particular, the archaeal lipids with ether rather than ester links between hydrocarbon tails and polar heads, are attributed with greater robustness and temperature stability. They exist in the liquid crystalline phase over a wide temperature range^39^. Our data suggest that there are other functional features of archaeal lipids that have so far not been taken into consideration, specifically their surface wetting properties. **Fig. 5A** shows that the coverage density of archaeal protocells on all particle surfaces is less than the coverage density of protocells made from other lipids. The number of archaeal membrane lipid nanotubes is also smaller (**Fig. 5D**). The diameter of the formed compartments however is of comparable size. The archaeal lipids are no worse candidates to form cell-sized lipid containers than other lipid types, but they seem to prefer particular surface features in order to undergo membrane transformations.

The images that were the source of the bar graphs in **Fig. 5A-C** were also used to determine the number of nanotubes per unit surface area (**Fig. 5D-F**), as well as the average size of the vesicles and the length of the nanotubes (**Fig. 5G-I**).

There is no strong association between the surface coverage densities of nanotubes and protocells. In general, especially for the bacterial lipids and partly the reference lipid mixture, where the nanotube density is high, the vesicle density appears to be also high, and vice versa. The archaeal lipids on micrometeorites deviate from this trend, i.e., there are many surfaces on which the protocell density is high (**Fig. 5A**) but there are no visible nanotubes on those surfaces (**Fig.5D**). An example is m2 in **Fig. 5A** and **Fig. 5D**. We previously reported wetting experiments where areas with very few or no lipid nanotubes were situated in vesicle-dense areas^19^. Since the nanotubes transform into lipid compartments, compartments occasionally form locally in high density under complete consumption of all lipid nanotubes in that region. The opposite situation can be seen in m3, where there are over 40 nanotubes per unit surface area (**Fig. 5D**) corresponding to very few lipid compartments (**Fig. 5A**). In this case, most nanotubes are originating from the particle, but they are extending outwards rather than being part of a network fully adhered to the particle (cf. SI for corresponding image). If lipid compartments are formed on such nanoutubes, they could be anywhere on the tube but are not necessarily adhered to the particle (cf. **Fig. 6A** and discussion below).

**Figure 6.**
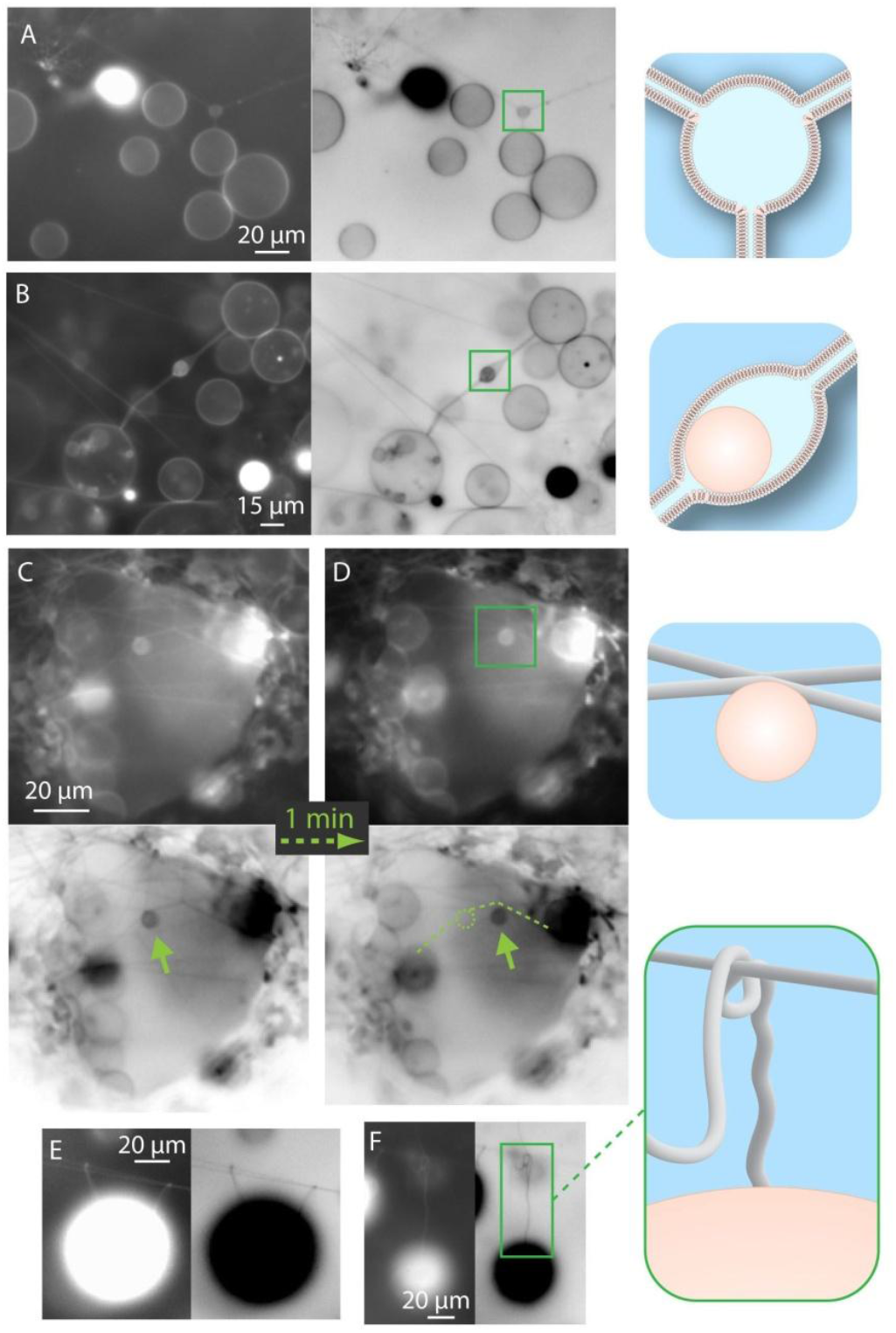
Different nanotube-vesicle configurations observed on particle surfaces. (A) Protocell-nanotube network where nanotubes are directly connected to a protocell with a continuous lipid bilayer and internal volume. (B) A lipid agglomerate/vesicle is carried inside an inter-vesicular nanotube. (C-D) A lipid agglomerate/vesicle is sliding along the surface of the nanotube network inside a cavity on the micrometeorite surface. (E-F) Multilamellar lipid reservoirs are entangled with/caught on nanotubes. Inverted versions of each fluorescence micrograph are depicted for better contrast. The schematic drawings to the right highlight the structures observed in the fluorescence micrographs A, B, D and F from top to bottom, respectively.

Lipid nanotubes were associated with particle-bound protocells in various ways, examples are shown in **Figure 6** (cf. supporting information for the full images corresponding to the panels). The most common form is the formation of bulges on the nanotubes. In this arrangement, protocells where the lipid membrane of the compartments and the lipid nanotube is continuous, the aqueous interior volumes of the protocell and the interior space of the lipid nanotubes are merged, i.e., directly connected (**Fig. 6A**). In other examples the compartments are located inside the nanotubes connecting different lipid compartments (**Fig. 6B**). Nanotubular connections between biological cells exist throughout all domains of life^40^ and translocate small molecules, vesicles, and organelles^41^. Small vesicles also attach to the membrane of the nanotubes and move along them (**Fig. 6C-D**). A previously reported similar observation is the attachment of *E*.*coli* bacteria surfing along the membrane of lipid nanotubes connecting two lipid compartments^42^. Briefly, driven by the surface tension gradient across the lipid membrane (Marangoni effect), bacteria attached to the lipid bilayer of a nanotube connecting two lipid vesicles were able to move from one attached vesicle to another.

MLVs in the surrounding aqueous environment get entangled with nearby nanotubes extending out of protocells on the particles (**Fig. 6E-F**). This can be a possible pathway to migration of fresh lipid material into membranes connected to the micrometeorites. In the present work we did not explore whether or how this occurs, but in a recent study we have already shown that fusion of MLVs of different lipid contents is enhanced on solid surfaces, and causes fusion and mixing of membrane lipids, leading to compositional diversity^20^.

## Conclusion

We experimentally gathered evidence that micrometeorites promote the formation of protocells from lipid reservoirs in a size range close to modern biological cells. Depending on the lipid type, we observed differences in the tendency to grow primitive compartments. Ether lipids representative of archaeal membranes prefer micrometeorites over reference particles; bacterial lipids exhibit the opposite behavior. A lipid mixture including plant-derived lipids confirms observations in earlier work; it is almost ideally suited to wet high energy surfaces and shows no preference for a specific particle type. It is a stark contrast to the purely prokaryotic lipid types which exhibit these notable preferences.

The imaging data suggest that the earlier proposed nanotube-mediated protocell formation mechanism also applies to micrometeorites. Protocell formation depending on differences in surface topography/structure and composition should be further investigated with a larger selection of micrometeorites of different types. In this work, we only considered three different classes of micrometeorites. Roughness, porosity and local differences in composition could be influential factors. The differences found between directly used and plasma treated reference particles (model micrometeorites) gives evidence that pristine surfaces outperform weathered and contaminated surfaces as substrates for protocell development. In this particular aspect they are reasonable physical models for micrometeorites, but in terms of composition and surface texture they are, according to the SEM data, not equivalent.

Micrometerorites have been continuously showering the planet since its formation and have been effectively distributed over the entire surface. On the early Earth, micrometeorites must have consistently reached regions of elevated prebiotic activity, such as warm ponds and other environmental niches, where molecular organic species assumed to have been involved in the origin of life were generated and accumulated. In the light of our earlier formulated hypothesis that solid surfaces might have been instrumental in the formation and development of protocells from simple biosurfactants, micrometeorites constituted a constant source of fresh and diversely structured high energy surfaces. This essentially means that activating surfaces were supplied to the locations where relevant molecules were accumulated. It is a clear advantage over the other scenario where molecular species would have had to amass on fixed locations where proper mineral surfaces were exposed. In addition, due to their small size these extraterrestrial microparticles are highly mobile in a dynamic exterior, and could have contributed to the exchange of prebiotic materials between various local microenvironments. Micrometeorites with their diverse composition, which not only comprises elements such as calcium, magnesium and iron, but typically also nickel, chromium, iridium and other less abundant elements in significant quantities, could have been relevant for catalytic processes that may have influenced the generation of chemical species which eventually enabled the transition from the non-living to the living world. Given the evidence from analytical studies that micrometeorites contain different organic species, it is not unreasonable to contemplate that the interaction of micrometeorites with organic matter under favorable conditions could have had a driving role in abiogenesis on the Earth and other planets of similar environmental make-up, which perhaps grants them a special place in the emerging field of astrobiology.

## Supporting information

Fig5 source images

## Acknowledgements

We are grateful for the support by the MC2 cleanroom staff, and acknowledge the European Union Erasmus program for supporting the training of E.C. at Chalmers University of Technology.

