## Supporting Information for "Protocell formation on micrometeorites"

### 1. Materials and Methods

*Preparation of lipids*: Three different lipid compositions were used to prepare the lipid suspensions: (1) 99 wt% 4ME 16:0 Diether PC (Sigma Aldrich) - 1 wt% 1,2-dipalmitoyl-sn-glycero-3-phosphoethanolamine-N-(lissamine rhodamine B sulfonyl) (16:0 Liss Rhod PE, Avanti Polar Lipids Inc., USA), (2) 99 wt% *E.coli* polar lipid extract (Avanti Polar Lipids Inc., USA) - 1wt% ATTO 655-1,2- dioleoyl-sn-glycero-3-phosphoethanolamine (ATTO 655-DOPE, ATTO-TEC GmbH, Germany), (3) 50 wt% Soybean polar lipid extract (Avanti Polar Lipids Inc., USA) - 49 wt% E. coli polar lipid extract - 1wt% ATTO 655-DOPE. To prepare each lipid suspension, the dehydration/rehydration method described by Karlsson et al.[1] was applied as follows: All lipids in the amounts specified above were dissolved in chloroform to a concentration of 10 mg/ml. 300 µl of this solution was transferred to a 10 ml round bottom flask, and by means of a rotary evaporator, the chloroform was removed at 20 kPa over a period of 6 hours. 3 ml of phosphate buffer/saline (PBS) containing 5 mM Trizma Base, 30 mM $K_3PO_4$, 30 mM $KH_2PO_4$, 3 mM $MgSO_4*7H_2O$, 0.5 mM $Na_2EDTA$ (pH=7.4 adjusted with KOH) and 30 µl glycerol, was added for rehydration to the dry lipid film in the flask. The flask was kept at +4 °C overnight, and then ultrasonicated for 5-10 seconds at room temperature to form the lipid suspension. For the experiments, 3 µl of this lipid suspension was desiccated for 20 minutes and rehydrated with 0.5 ml of HEPES buffer for 5 min. (10 mM HEPES and 100 mM NaCl, pH= 7.8, adjusted with NaOH). 4mM $CaCl_2$ was added to the HEPES buffer for all experiments except for the *E. coli* lipids, which aggregated under these conditions.

*Preparation of micrometeorites and reference particles*: The micrometeorites were collected according to the methods described by Larsen[2-3]. Before each experiment, the model micrometeorites and micrometeorites were treated with Tetrahydrofuran (Sigma Aldrich) , Triton-X (Sigma Aldrich), water, and were subsequently exposed to oxygen plasma at 250 mbar, 100 W and 10 sccm $O_2$ for 20 minutes (Diener Atto, Diener Electronic GmbH, Germany).

*Scanning electron microscopy and energy with dispersive X-ray spectroscopy*: Particles were SEM-imaged and analyzed with a Zeiss SUPRA SEM at MC2/Chalmers University of Technology (Göteborg, Sweden), equipped with a high sensitivity SDD detector with 133eV resolution for energy dispersive X-ray spectroscopy (EDX) unit (IXRF Systems, USA). Compositional analysis was performed with the IXRF-supplied software. Before transfer into the microscope, the particles were adhered to standard sticky carbon pads.

*Preparation of the sample chamber for optical microscopy*: Three circular frames made from polydimethylsiloxane (PDMS, Sigma Aldrich) were placed on a glass bottom dish (WillCo Wells B.V. Amsterdam, NL). Each particle type (mirometeorites and reference particles) was placed in one chamber under a stereomicroscope and and half-filled with HEPES buffer, using an automatic pipette. Rehydrated lipid solution was added to the chambers and with the lid of the petridish closed, and left overnight before imaging.

*Imaging:* An inverted Leica DM-IRB2 research microscope was used for fluorescence imaging with a 63x (oil) 1.40 NA objective. The fluorophores in the lipid suspensions were excited with diode laser sources; Rhodamine 561 nm: 240mW, Cobolt 06-MLD (Hübner Photonics, Germany), Atto 655: 150 mW, 635nm, (M-Series Dragon Lasers Ltd., UK). The emitted light was collected and processed using a digital camera (Prosilica GX, Allied Vision Technologies, Germany).

*Image analyses*: ImageJ (NIH) was used for the analysis of the fluorescence micrographs: fluorescence intensity profiles, lipid compartment and nanotube counts and size/length determination. In images containing multiple compartments and nanotubes, the mean average of these values were taken for that image. For calculation of the particle cross section area, the freehand selection tool was used to mark the contour of the section, and the area inside the contour was automatically calculated with the ImageJ *analyze* function. For counting of compartments and nanotubes, any relevant structure physically connected to the particle was considered as particle-bound. For example, if a vesicle was not directly connected to the particle, but connected to a vesicle on the particle, it was also considered as a particle-bound compartment. Bar-graphs and schematic drawings were produced with Matlab and Adobe Illustrator.

### 2. Extended Fig. 6

Complete images corresponding to the panels in Fig. 6 are shown below. The panels are marked in yellow frames in the image and the contour of the particles are shown by orange dashed lines.

Full image corresponding to Fig. 6A:

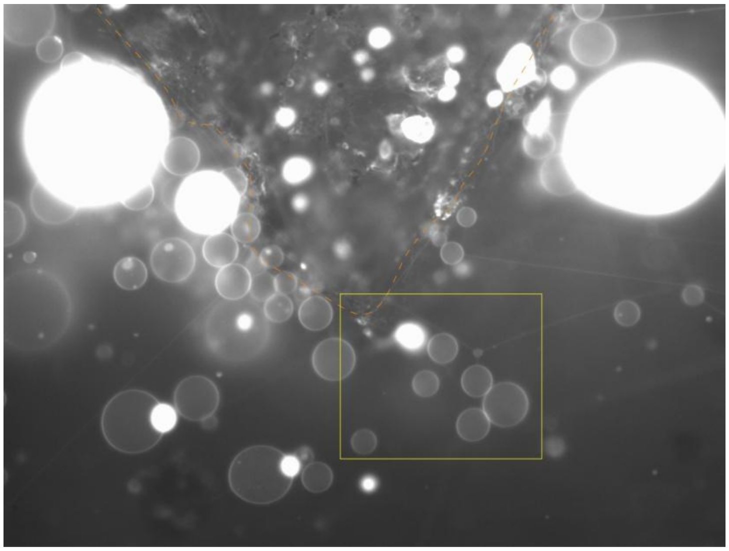

Full image corresponding to Fig. 6B and 6C:

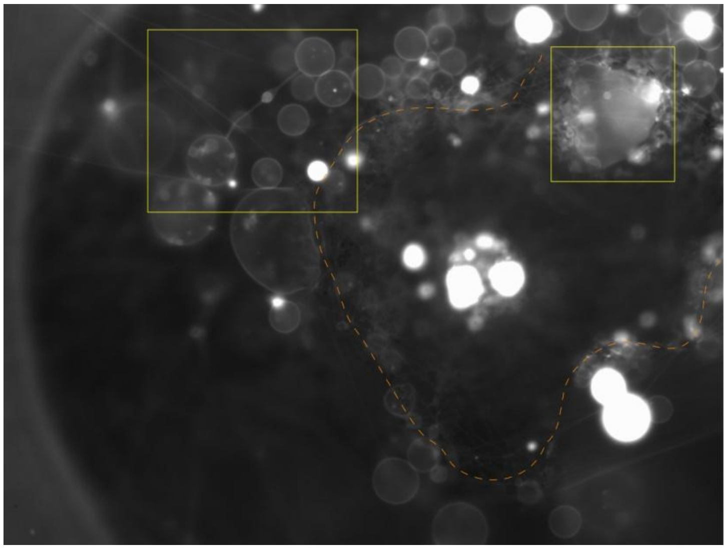

Full image corresponding to Fig. 6D:

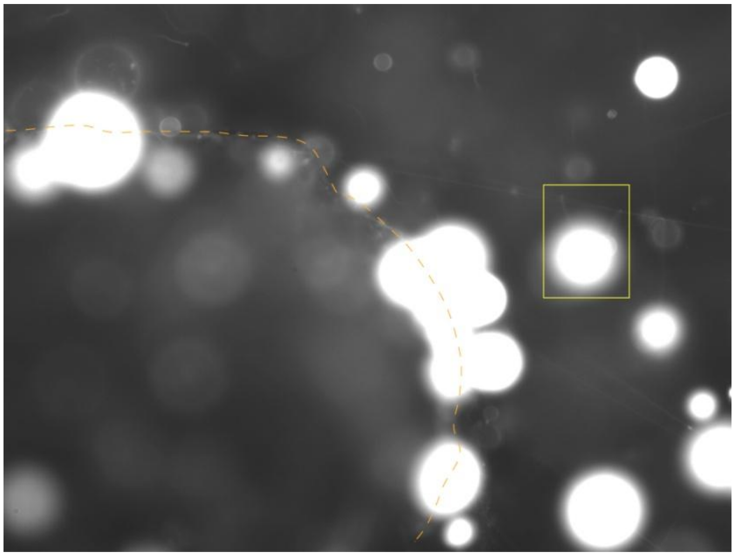

Full image corresponding to Fig. 6F:

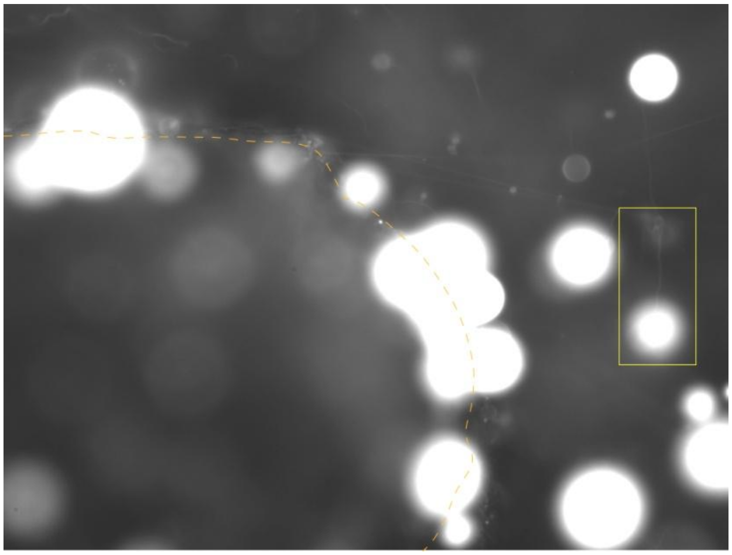

### 3. Analyses of all particles used in the experiments

In this section the SEM images of reference particles, raw EDX spectra and analysis data for all particles are provided:

**Micrometeorite 1:** Corresponds to the micrometeorite shown in Fig. 2 panels B-E-H

- Exposure to archaeal lipids: Fig. 5A - m1
- Exposure to E.coli lipids: Fig. 5B - m1
- Exposure to plant lipids mixture: Fig. 5C - m1

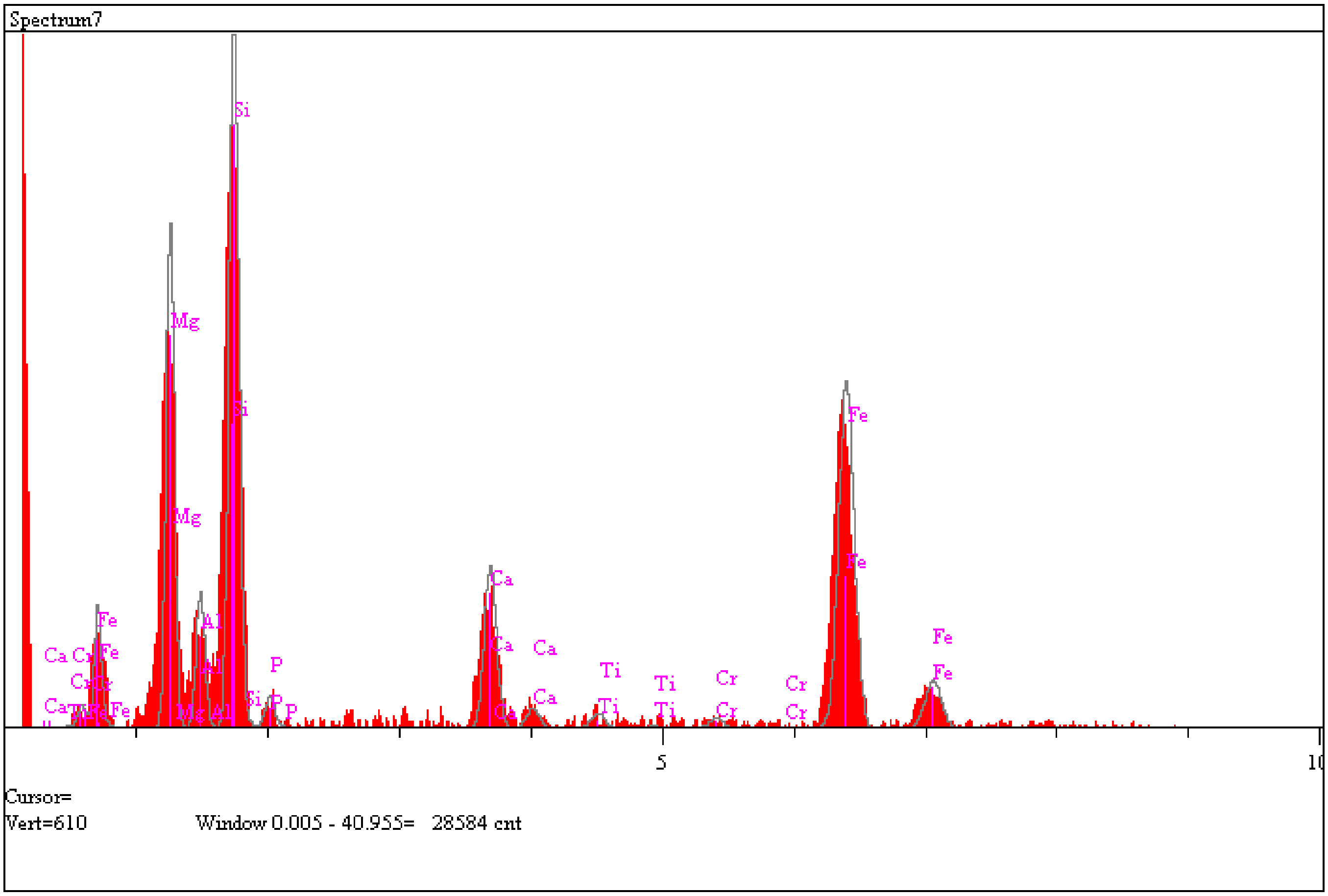

| Elt. | Line | Intensity (c/s) | Error 2-sig | Atomic % | K-Ratio |
|---|---|---|---|---|---|
| N | Ka | 0.00 | 0.000 | 0.000 | 0.0000 |
| Mg | Ka | 57.90 | 1.965 | 11.256 | 0.0421 |
| Al | Ka | 16.28 | 1.042 | 2.793 | 0.0129 |
| Si | Ka | 99.00 | 2.569 | 15.808 | 0.0876 |

| | | | | | | |
|---|---|---|---|---|---|---|
| P | Ka | 4.32 | 0.537 | 0.697 | 0.0044 | |
| Ca | Ka | 27.25 | 1.348 | 5.424 | 0.0550 | |
| Ti | Ka | 2.66 | 0.421 | 0.673 | 0.0079 | |
| Cr | Ka | 2.00 | 0.365 | 0.690 | 0.0099 | |
| Fe | Ka | 74.05 | 2.222 | 62.659 | 0.7802 | |
| | | | | 100.000 | | Total |

**Micrometeorite 2:** Corresponds to the micrometeorite shown in Fig. 2 panels A-D-G

- Exposure to archaeal lipids: Fig. 5A – m2
- Exposure to E.coli lipids: Fig. 5B – m2
- Exposure to plant lipids mixture: Fig. 5C – m2
- Fluorescence micrograph in Fig. 3I and Fig. 3G corresponds to this particle.

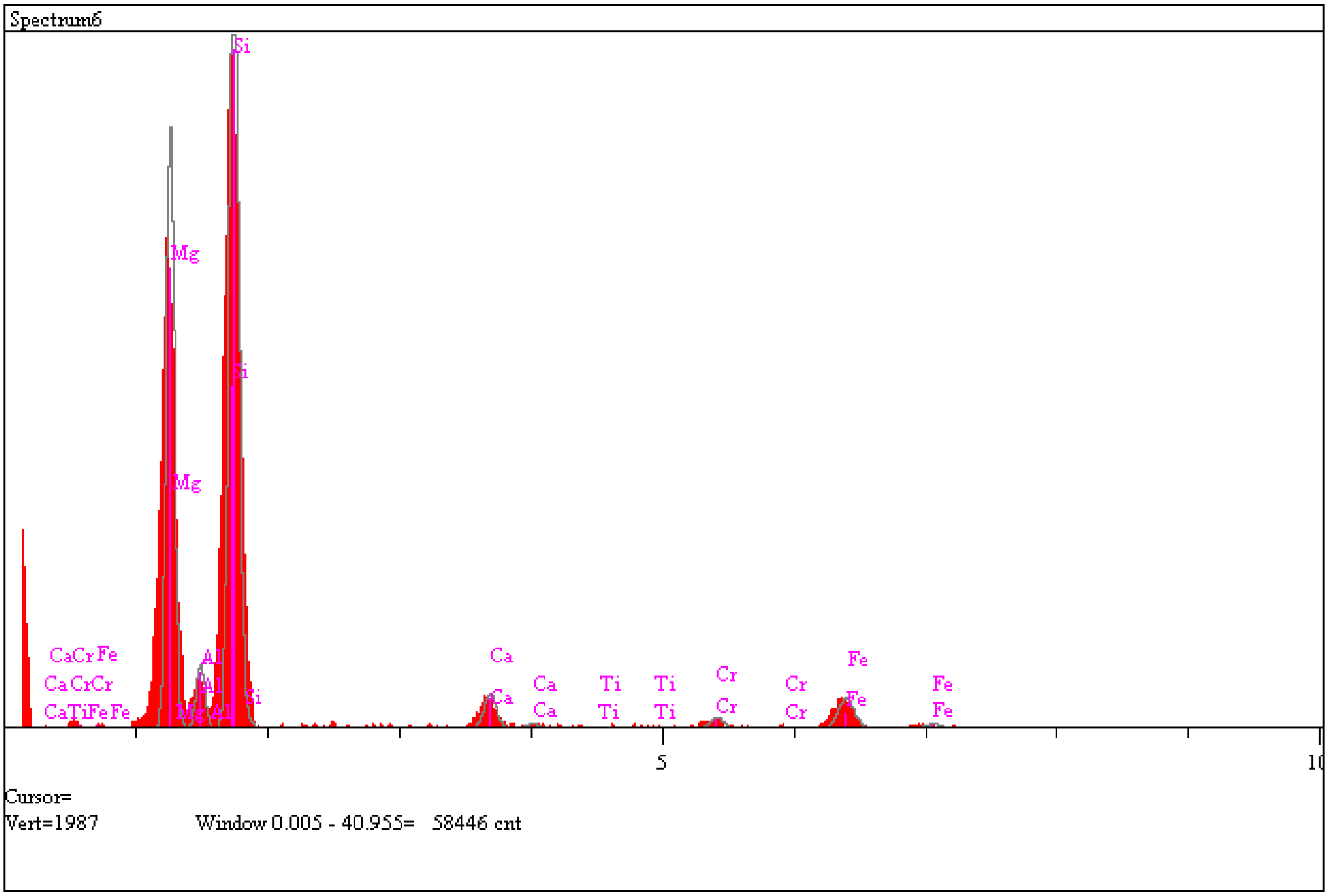

| Elt. | Line | Intensity<br>(c/s) | Error<br>2-sig | Atomic<br>% | K-Ratio |
|---|---|---|---|---|---|
| Mg | Ka | 224.03 | 3.864 | 28.483 | 0.1992 |
| Al | Ka | 24.54 | 1.279 | 3.203 | 0.0238 |
| Si | Ka | 360.64 | 4.903 | 45.666 | 0.3907 |
| Ca | Ka | 18.72 | 1.117 | 3.308 | 0.0462 |
| Ti | Ka | 0.69 | 0.215 | 0.162 | 0.0025 |
| Cr | Ka | 6.57 | 0.662 | 2.308 | 0.0399 |
| Fe | Ka | 21.18 | 1.188 | 15.539 | 0.2731 |
| Ni | Ka | 0.57 | 0.194 | 1.330 | 0.0245 |

| | | | | | | |
|---|---|---|---|---|---|---|
| | | | | 100.000 | | Total |

**Micrometeorite 3:** Corresponds to the micrometeorite shown in Fig. 2 panels C-F-I

- Exposure to archaeal lipids: Fig. 5A – m3
- Exposure to E.coli lipids: Fig. 5B – m3
- Exposure to plant lipids mixture: Fig. 5C – m3
- Fluorescence micrograph in Fig. 3H corresponds to this particle.

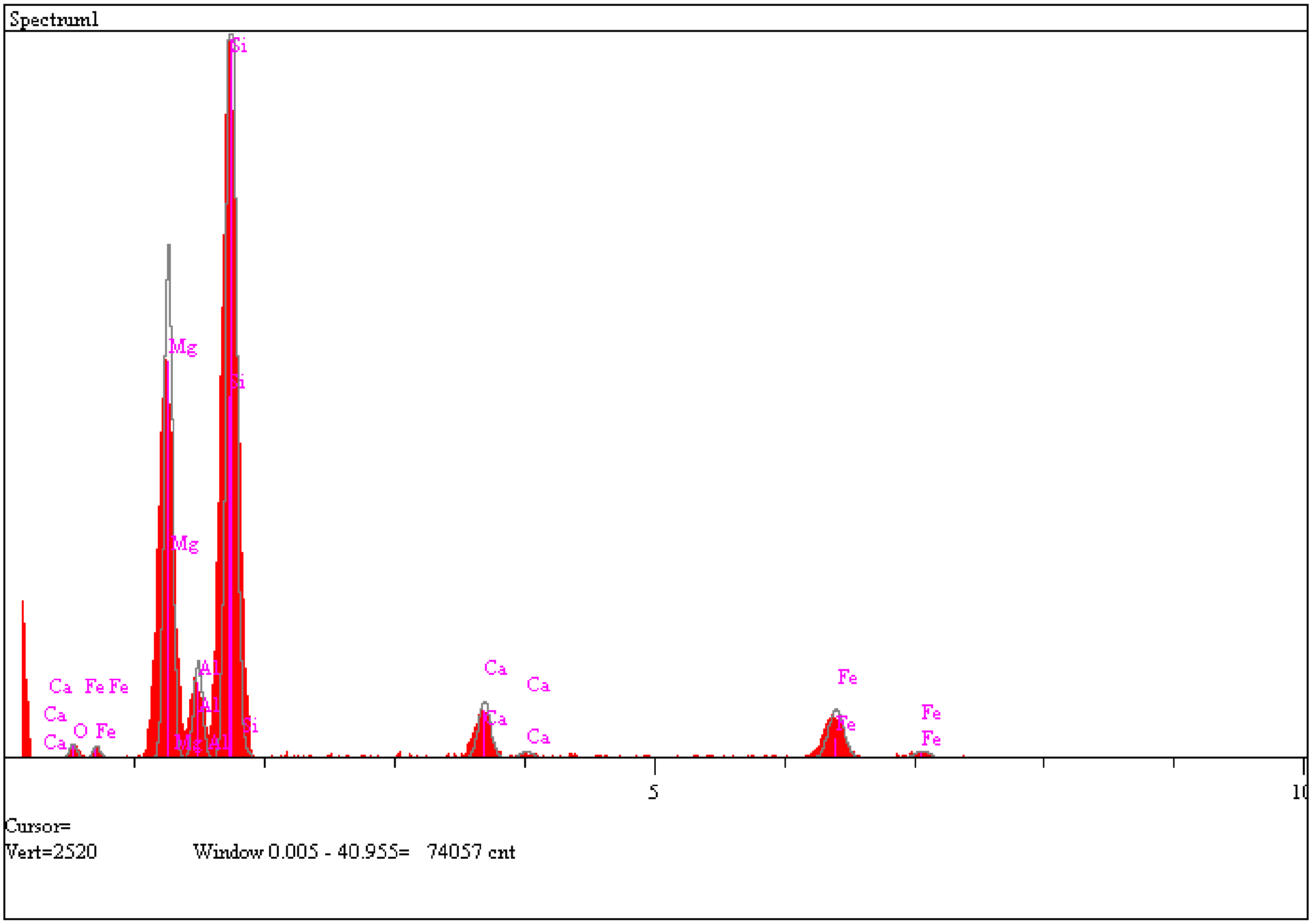

| Elt. | Line | Intensity | Error | Atomic | K-Ratio |
|---|---|---|---|---|---|

| | | (c/s) | 2-sig | % | | |
|---|---|---|---|---|---|---|
| N | Ka | 0.00 | 0.000 | 0.000 | 0.0000 | |
| O | Ka | 5.32 | 0.596 | 1.565 | 0.0049 | |
| Mg | Ka | 232.28 | 3.935 | 22.709 | 0.1486 | |
| Al | Ka | 45.65 | 1.744 | 4.408 | 0.0319 | |
| Si | Ka | 466.27 | 5.575 | 43.874 | 0.3634 | |
| Ca | Ka | 38.12 | 1.594 | 4.973 | 0.0677 | |
| Fe | Ka | 41.36 | 1.660 | 22.471 | 0.3836 | |
| | | | | 100.000 | | Total |

**Model Micrometeorite 1:**

- Exposure to archaeal lipids: Fig. 5A – p1
- Exposure to E.coli lipids: Fig. 5B – p1
- Exposure to plant lipids mixture: Fig. 5C – p1
- Fluorescence micrograph in Fig. 3E corresponds to this particle.

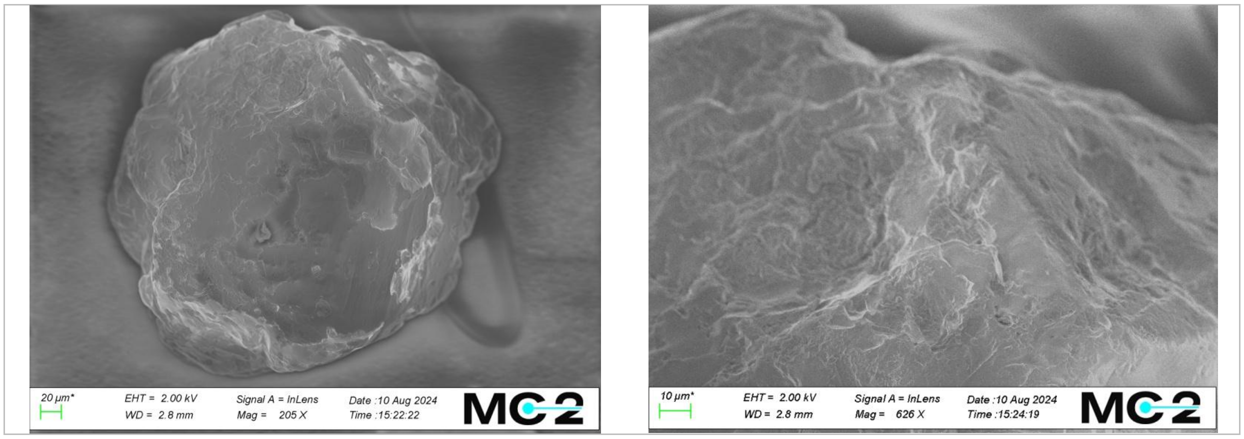

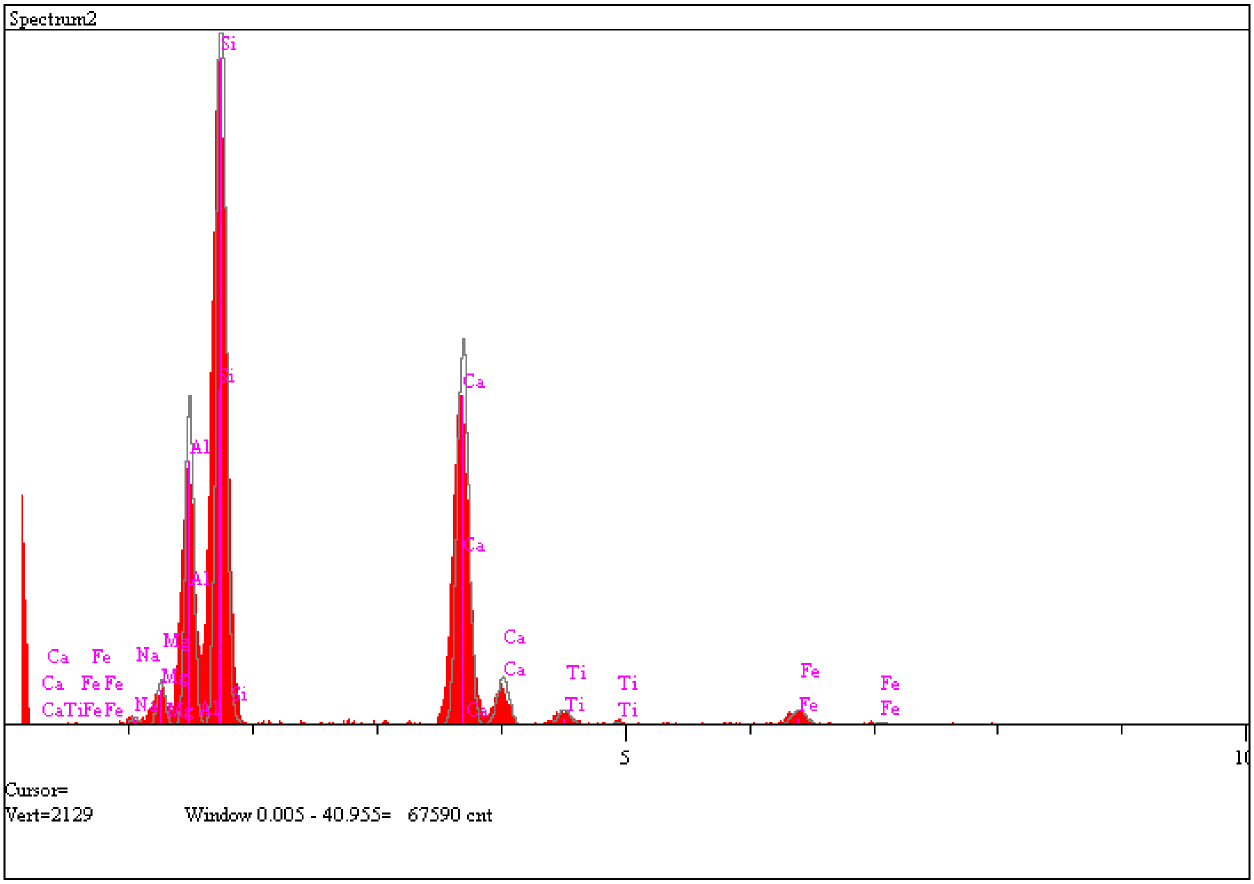

| Elt. | Line | Intensity (c/s) | Error 2-sig | Atomic % | K-Ratio | |
|---|---|---|---|---|---|---|
| Na | Ka | 3.35 | 0.472 | 0.448 | 0.0022 | |
| Mg | Ka | 17.89 | 1.092 | 1.922 | 0.0121 | |
| Al | Ka | 137.26 | 3.025 | 13.628 | 0.1012 | |
| Si | Ka | 381.74 | 5.044 | 39.625 | 0.3137 | |
| Ca | Ka | 227.95 | 3.898 | 34.640 | 0.4268 | |
| Ti | Ka | 10.20 | 0.825 | 2.151 | 0.0281 | |
| Fe | Ka | 11.85 | 0.889 | 7.586 | 0.1159 | |
| | | | | 100.000 | | Total |

**Model Micrometeorite 2:**

- Exposure to archaeal lipids: Fig. 5A – p2
- Exposure to E.coli lipids: Fig. 5B – p2
- Exposure to plant lipids mixture: Fig. 5C – p2
- Fluorescence micrograph in Fig. 3D corresponds to this particle.

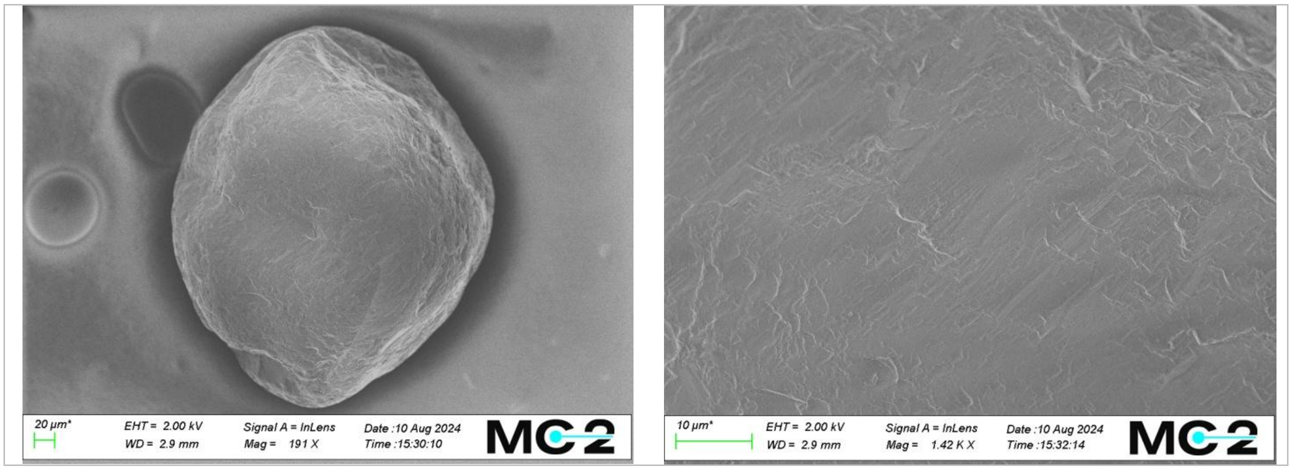

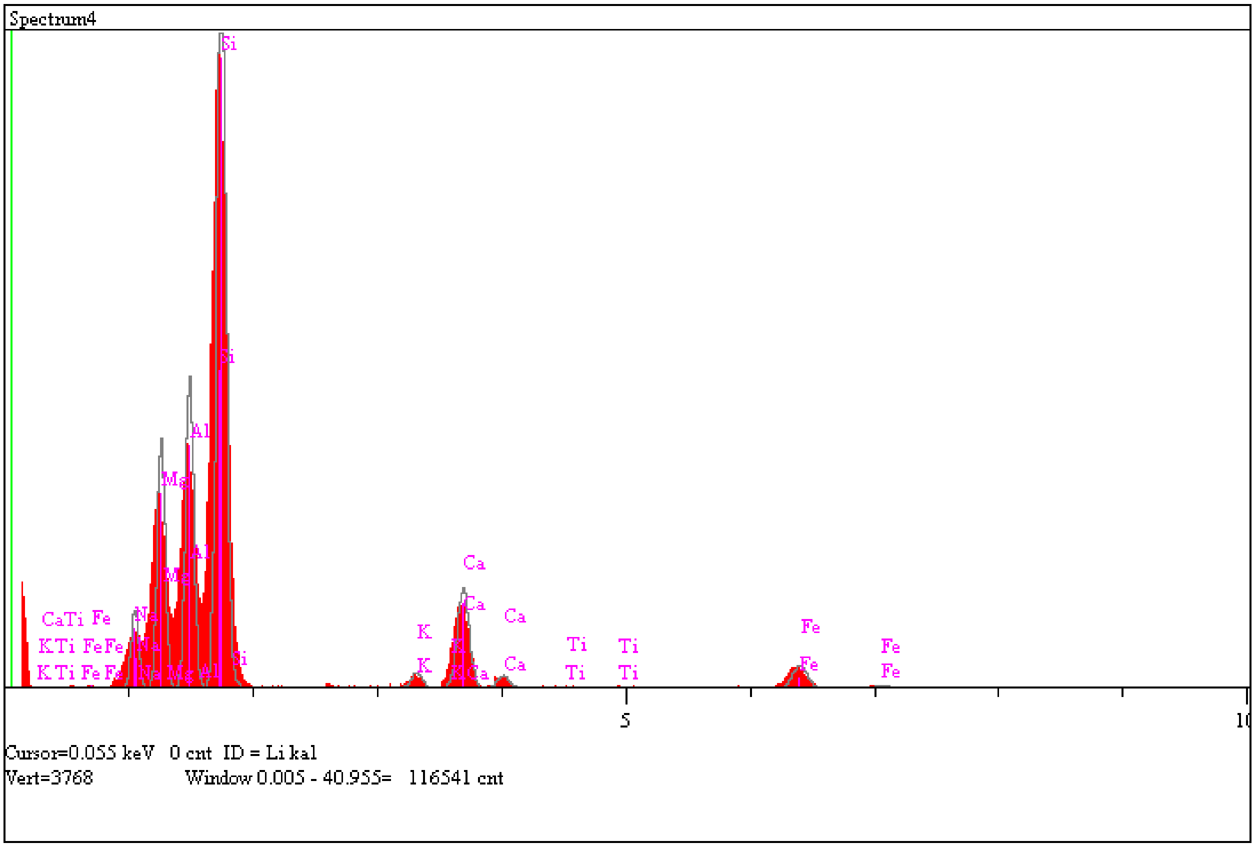

| Elt. | Line | Intensity (c/s) | Error 2-sig | Atomic % | K-Ratio | |
|---|---|---|---|---|---|---|
| Na | Ka | 54.48 | 1.906 | 4.231 | 0.0248 | |
| Mg | Ka | 186.45 | 3.525 | 12.033 | 0.0870 | |
| Al | Ka | 242.70 | 4.022 | 15.208 | 0.1238 | |
| Si | Ka | 681.32 | 6.739 | 44.572 | 0.3874 | |
| K | Ka | 16.25 | 1.041 | 1.346 | 0.0179 | |
| Ca | Ka | 110.47 | 2.714 | 10.267 | 0.1431 | |
| Ti | Ka | 2.09 | 0.374 | 0.260 | 0.0040 | |
| Fe | Ka | 31.32 | 1.445 | 12.083 | 0.2120 | |
| | | | | 100.000 | | Total |

**Model Micrometeorite 3:**

- Exposure to archaeal lipids: Fig. 5A – p3
- Exposure to E.coli lipids: Fig. 5B – p3
- Exposure to plant lipids mixture: Fig. 5C – p3
- Fluorescence micrograph in Fig. 3F corresponds to this particle.

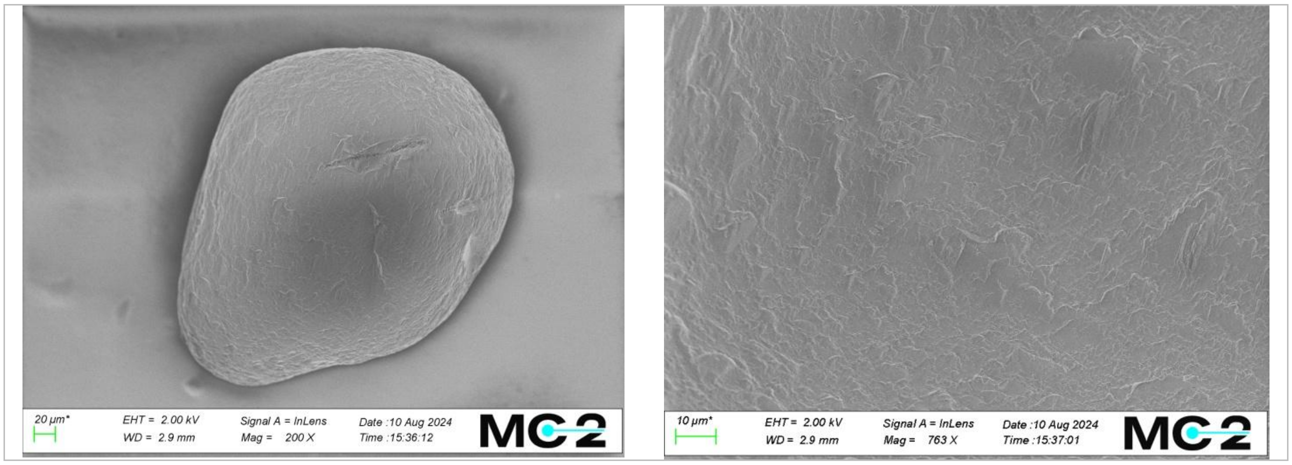

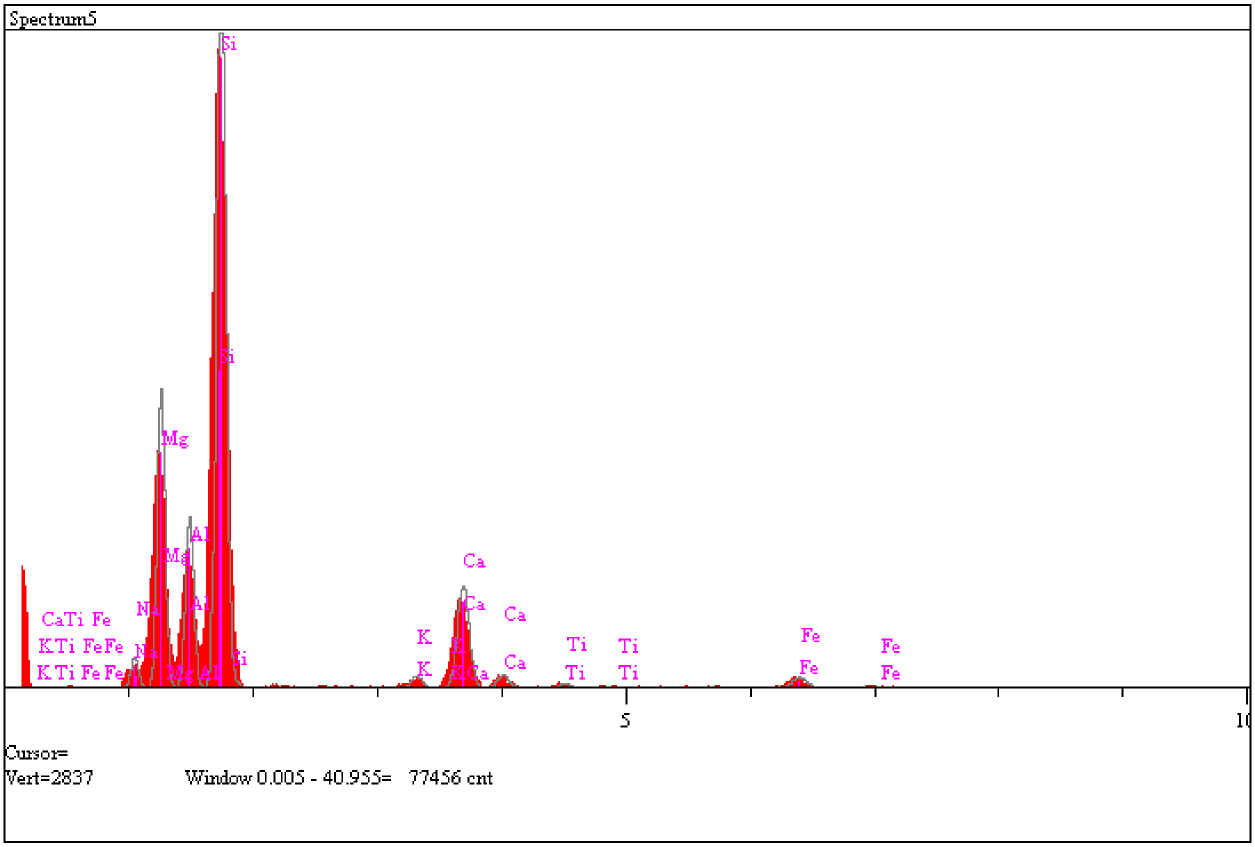

| Elt. | Line | Intensity (c/s) | Error 2-sig | Atomic % | K-Ratio | |
|---|---|---|---|---|---|---|
| Na | Ka | 16.32 | 1.043 | 1.871 | 0.0117 | |
| Mg | Ka | 168.88 | 3.355 | 16.159 | 0.1241 | |
| Al | Ka | 100.70 | 2.591 | 9.685 | 0.0809 | |
| Si | Ka | 511.69 | 5.840 | 50.775 | 0.4580 | |
| K | Ka | 9.34 | 0.789 | 1.214 | 0.0162 | |
| Ca | Ka | 85.28 | 2.384 | 12.478 | 0.1739 | |
| Ti | Ka | 4.19 | 0.528 | 0.826 | 0.0126 | |
| Fe | Ka | 11.52 | 0.876 | 6.993 | 0.1227 | |
| | | | | 100.000 | | Total |

**Natural particle 1 (with no plasma treatment)**

- Exposure to archaeal lipids, Fig. 5A - s1

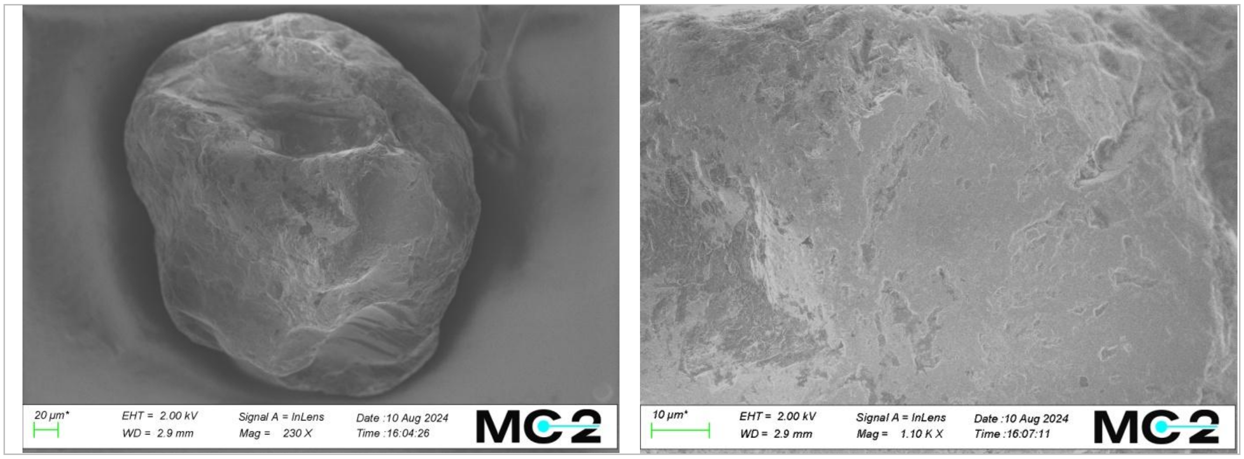

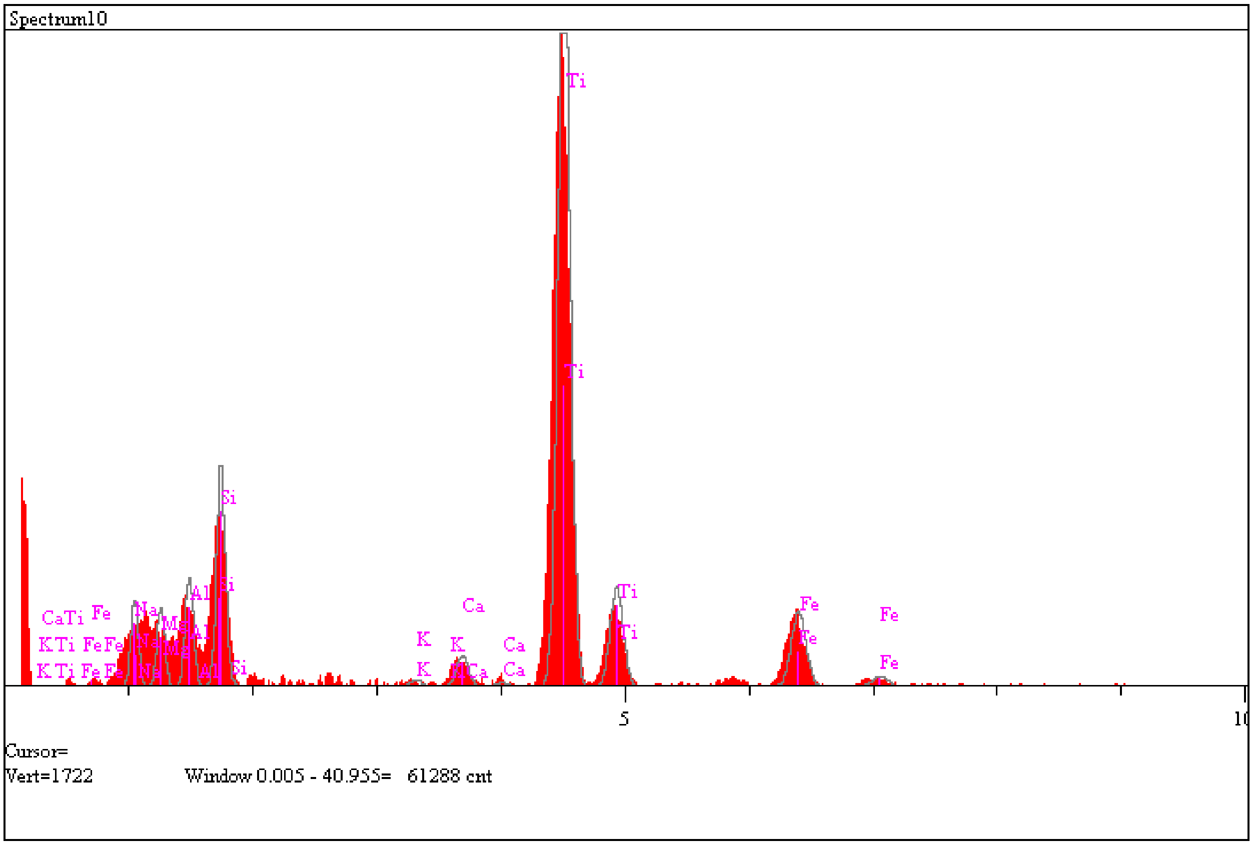

| Elt. | Line | Intensity (c/s) | Error 2-sig | Atomic % | K-Ratio | |
|---|---|---|---|---|---|---|
| Na | Ka | 27.64 | 1.357 | 3.530 | 0.0104 | |
| Mg | Ka | 27.05 | 1.343 | 2.658 | 0.0105 | |
| Al | Ka | 39.15 | 1.615 | 3.380 | 0.0166 | |
| Si | Ka | 83.98 | 2.366 | 6.906 | 0.0396 | |
| K | Ka | 3.36 | 0.473 | 0.306 | 0.0031 | |
| Ca | Ka | 15.95 | 1.031 | 1.537 | 0.0171 | |
| Ti | Ka | 400.53 | 5.167 | 59.078 | 0.6332 | |
| Fe | Ka | 48.03 | 1.789 | 22.606 | 0.2696 | |
| | | | | 100.000 | | Total |

**Natural particle 2 (with no plasma treatment)**

- Exposure to archaeal lipids, Fig. 5A – s2
- Fluorescence micrograph in Fig. 3A corresponds to this particle.

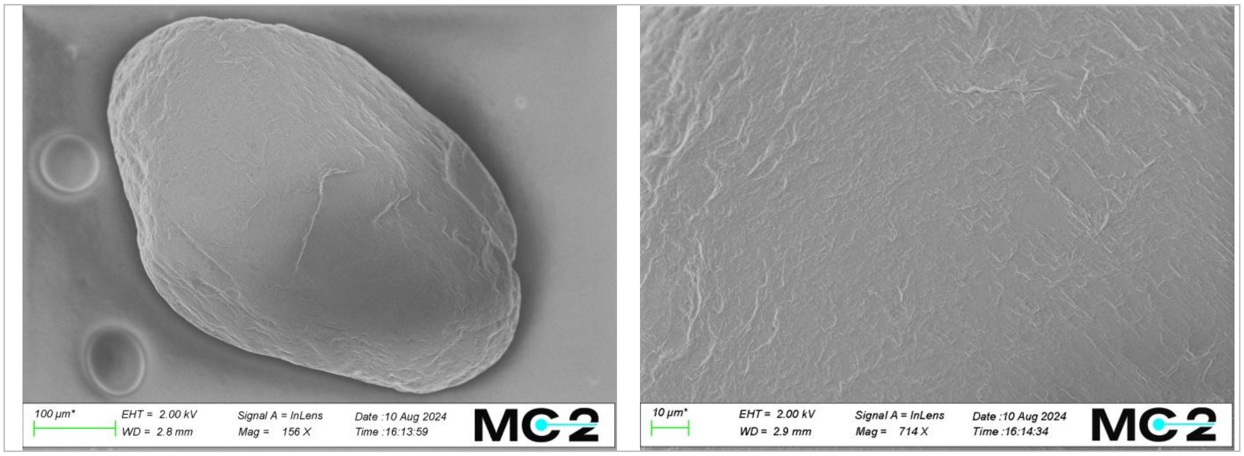

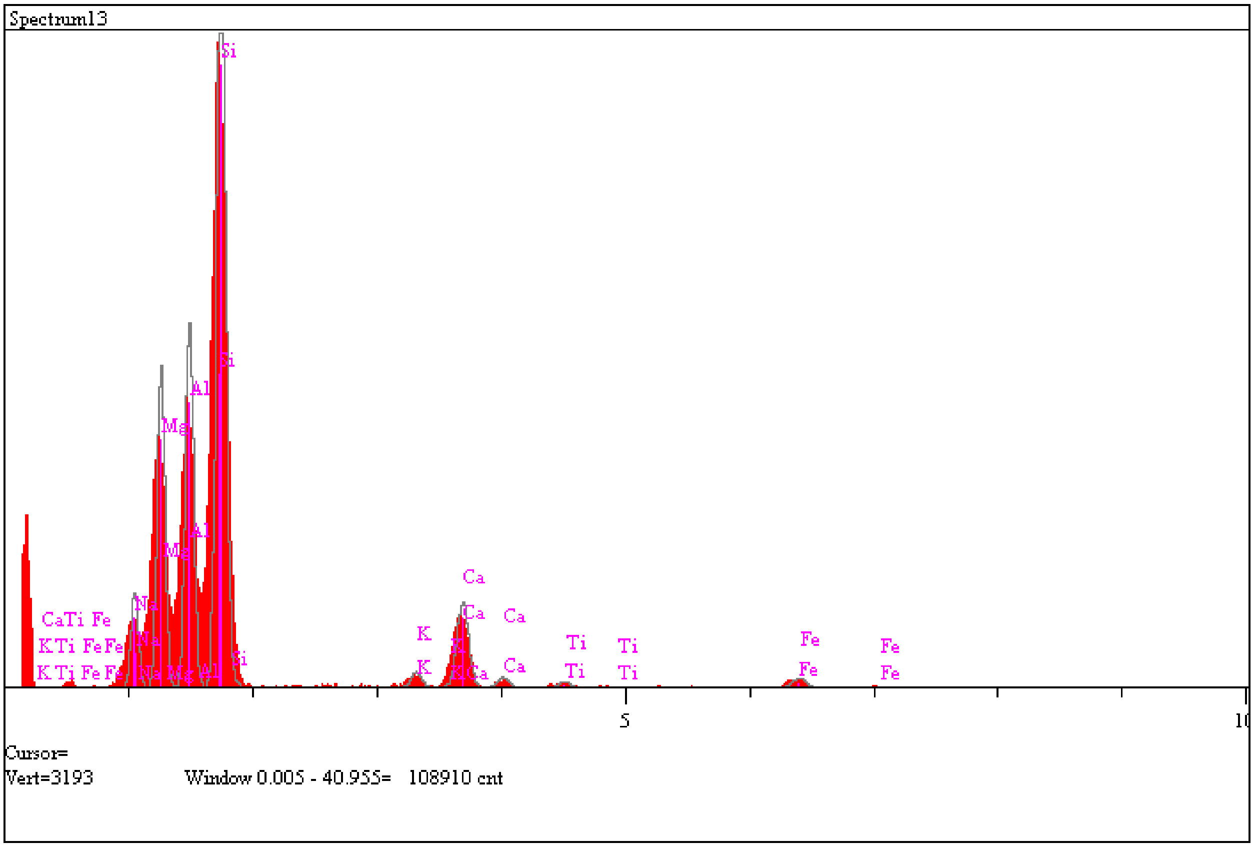

| Elt. | Line | Intensity (c/s) | Error 2-sig | Atomic % | K-Ratio | |
|---|---|---|---|---|---|---|
| Na | Ka | 56.65 | 1.943 | 4.793 | 0.0327 | |
| Mg | Ka | 204.25 | 3.690 | 14.846 | 0.1213 | |
| Al | Ka | 241.03 | 4.009 | 17.712 | 0.1564 | |
| Si | Ka | 579.99 | 6.218 | 45.625 | 0.4196 | |
| K | Ka | 14.22 | 0.973 | 1.432 | 0.0199 | |
| Ca | Ka | 80.40 | 2.315 | 9.080 | 0.1325 | |
| Ti | Ka | 6.32 | 0.649 | 0.959 | 0.0153 | |
| Fe | Ka | 11.87 | 0.889 | 5.554 | 0.1022 | |
| | | | | 100.000 | | Total |

**Natural particle 3 (with no plasma treatment)**

- Exposure to archaeal lipids, Fig. 5A – s3
- Fluorescence micrograph in Fig. 3C corresponds to this particle.

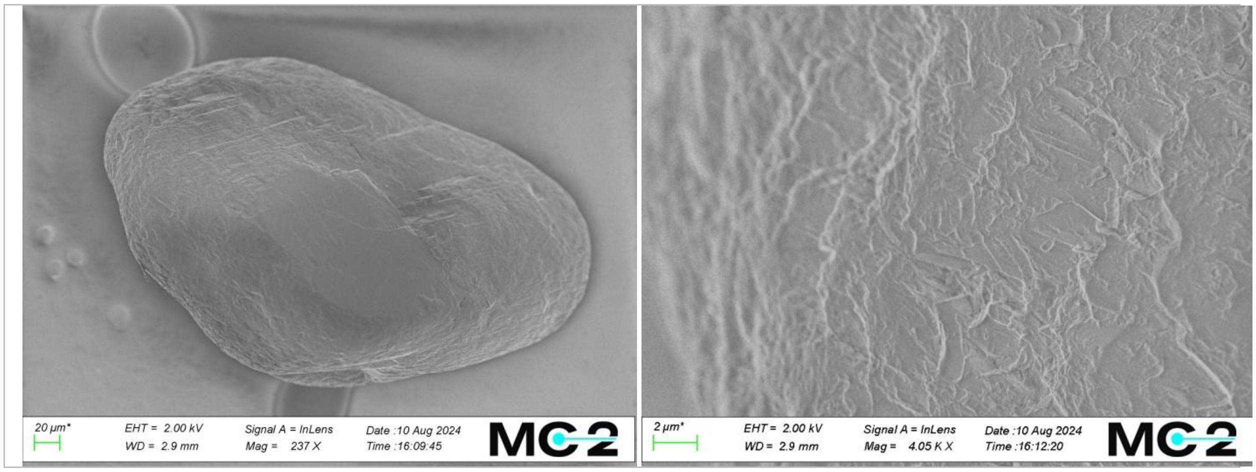

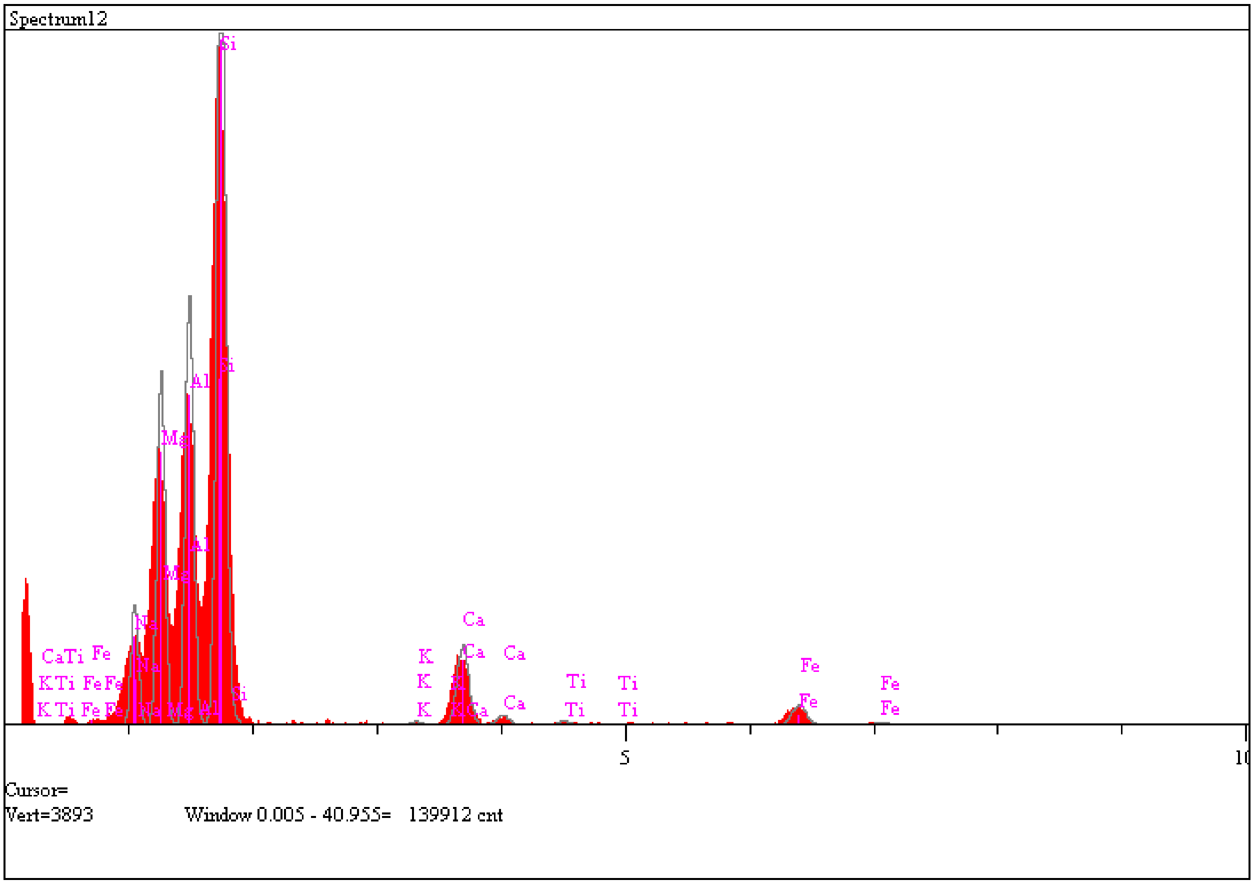

| Elt. | Line | Intensity (c/s) | Error 2-sig | Atomic % | K-Ratio | |
|---|---|---|---|---|---|---|
| Na | Ka | 82.70 | 2.348 | 5.558 | 0.0361 | |
| Mg | Ka | 259.90 | 4.162 | 14.850 | 0.1165 | |
| Al | Ka | 327.34 | 4.671 | 18.650 | 0.1603 | |
| Si | Ka | 713.03 | 6.894 | 43.163 | 0.3893 | |
| K | Ka | 3.75 | 0.500 | 0.286 | 0.0040 | |
| Ca | Ka | 86.38 | 2.399 | 7.327 | 0.1074 | |
| Ti | Ka | 4.65 | 0.556 | 0.527 | 0.0085 | |
| Fe | Ka | 27.37 | 1.351 | 9.640 | 0.1779 | |
| | | | | 100.000 | | Total |

**Natural particle 4 (with no plasma treatment)**

- Exposure to E.coli lipids, Fig. 5B – s4

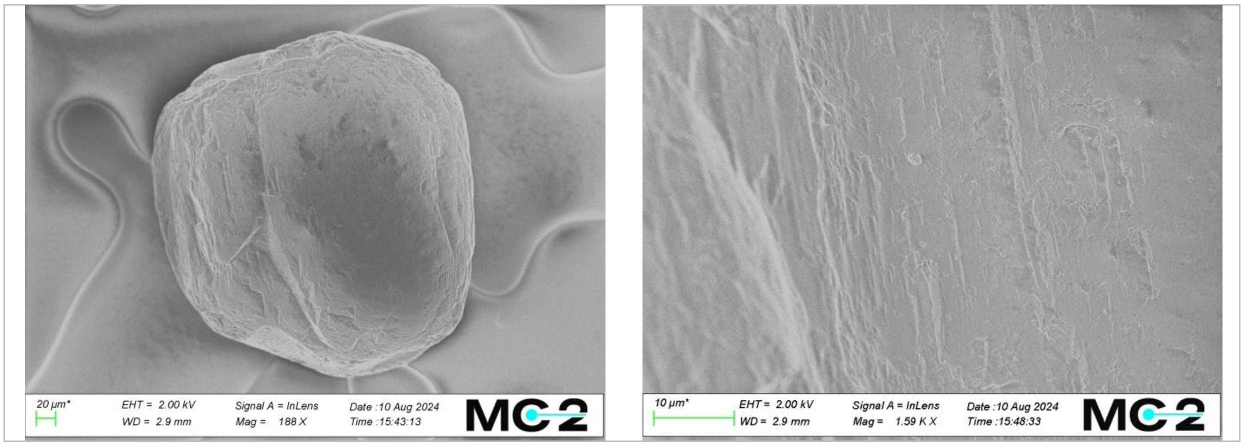

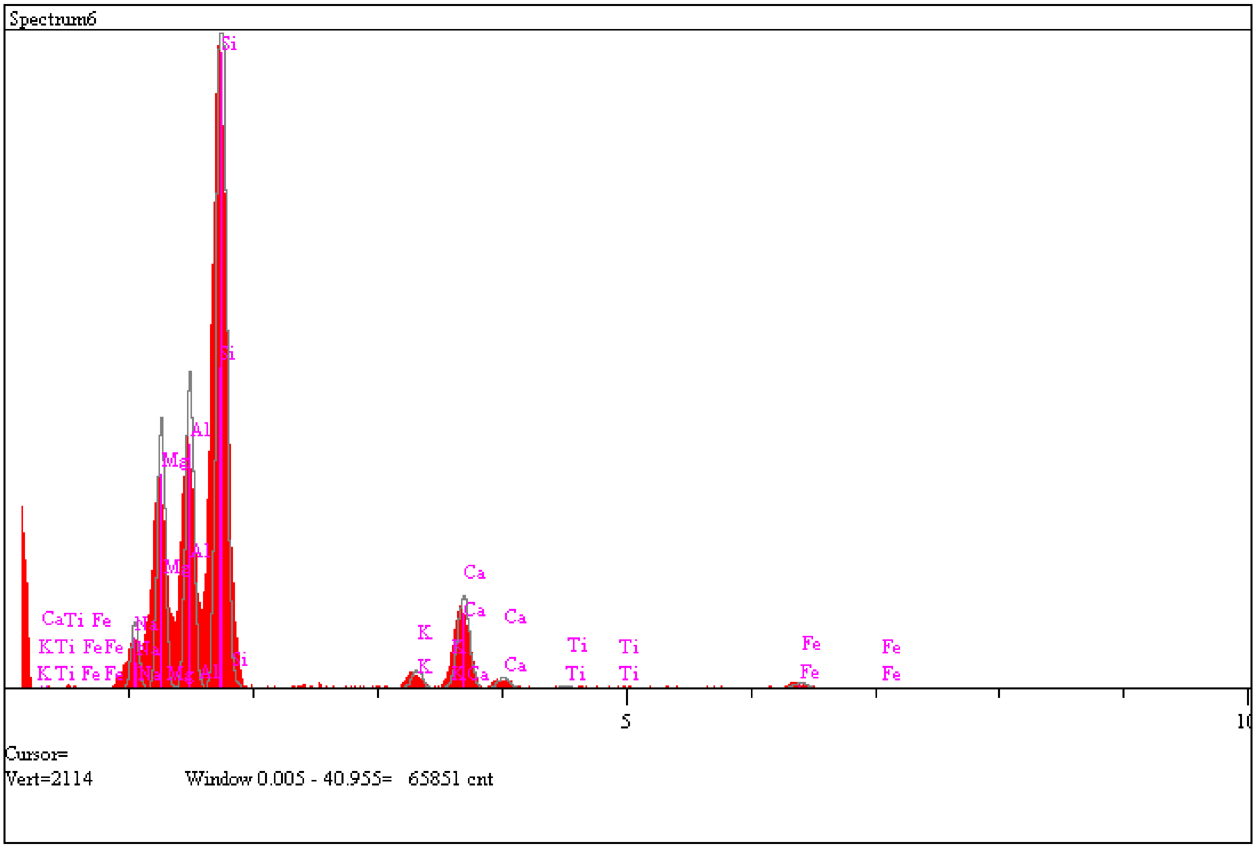

| Elt. | Line | Intensity (c/s) | Error 2-sig | Atomic % | K-Ratio | |
|---|---|---|---|---|---|---|
| Na | Ka | 26.69 | 1.334 | 3.667 | 0.0254 | |
| Mg | Ka | 114.05 | 2.757 | 13.462 | 0.1116 | |
| Al | Ka | 138.74 | 3.041 | 16.562 | 0.1484 | |
| Si | Ka | 385.09 | 5.066 | 49.476 | 0.4591 | |
| K | Ka | 10.88 | 0.852 | 1.826 | 0.0251 | |
| Ca | Ka | 57.68 | 1.961 | 10.885 | 0.1567 | |
| Ti | Ka | 1.82 | 0.349 | 0.464 | 0.0073 | |
| Fe | Ka | 4.69 | 0.559 | 3.658 | 0.0665 | |
| | | | | 100.000 | | Total |

**Natural particle 5 (with no plasma treatment)**

- Exposure to E.coli lipids, Fig. 5B – s5

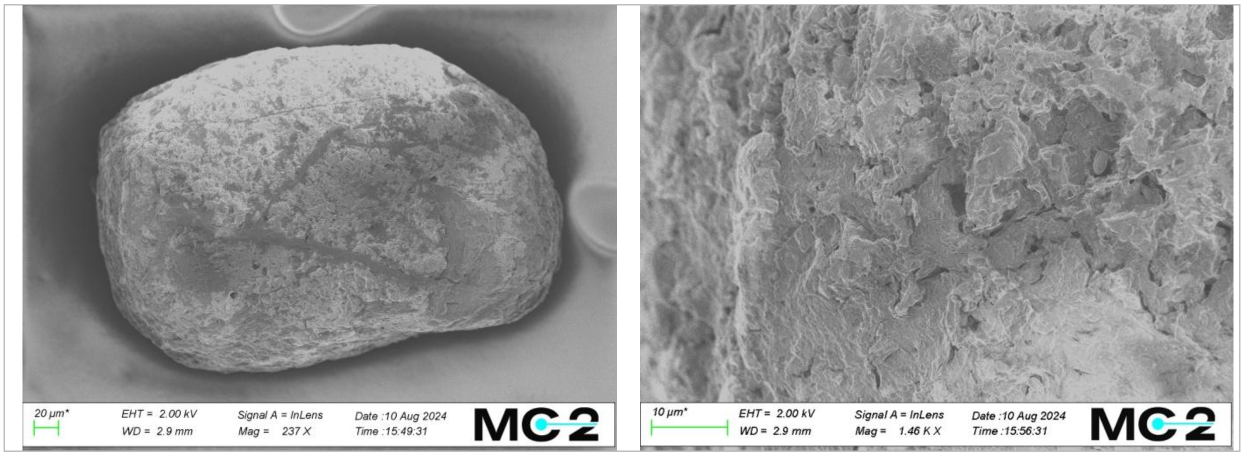

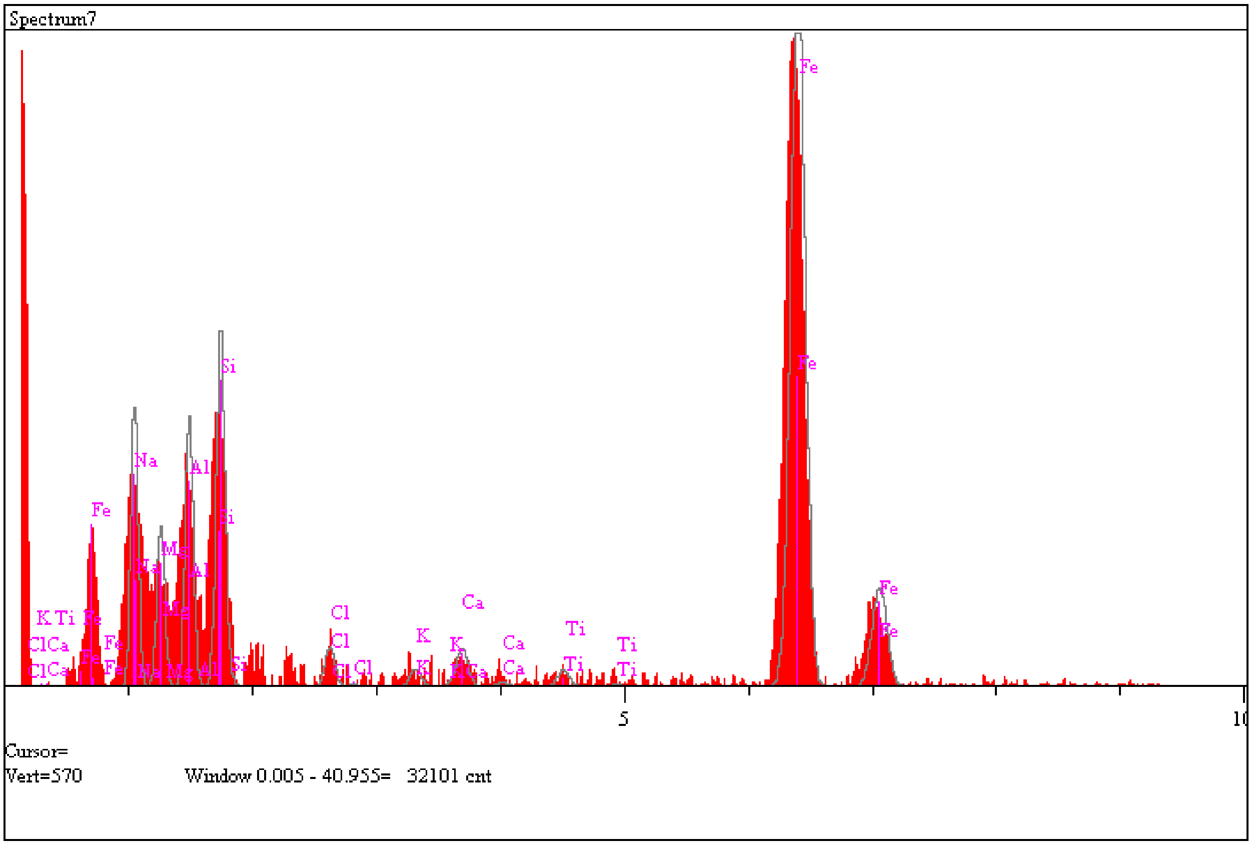

| Elt. | Line | Intensity (c/s) | Error 2-sig | Atomic % | K-Ratio | |
|---|---|---|---|---|---|---|
| Na | Ka | 30.10 | 1.416 | 5.441 | 0.0124 | |
| Mg | Ka | 18.24 | 1.103 | 2.456 | 0.0077 | |
| Al | Ka | 31.87 | 1.457 | 3.650 | 0.0147 | |
| Si | Ka | 44.68 | 1.726 | 4.717 | 0.0230 | |
| Cl | Ka | 5.72 | 0.617 | 0.586 | 0.0043 | |
| K | Ka | 2.86 | 0.437 | 0.331 | 0.0029 | |
| Ca | Ka | 6.27 | 0.647 | 0.788 | 0.0074 | |
| Ti | Ka | 2.92 | 0.441 | 0.442 | 0.0050 | |
| Fe | Ka | 150.33 | 3.165 | 81.589 | 0.9226 | |
| | | | | 100.000 | | Total |

**Natural particle 6 (with no plasma treatment)**

- Exposure to E.coli lipids, Fig. 5B – s6
- Fluorescence micrograph in Fig. 3B corresponds to this particle.
-

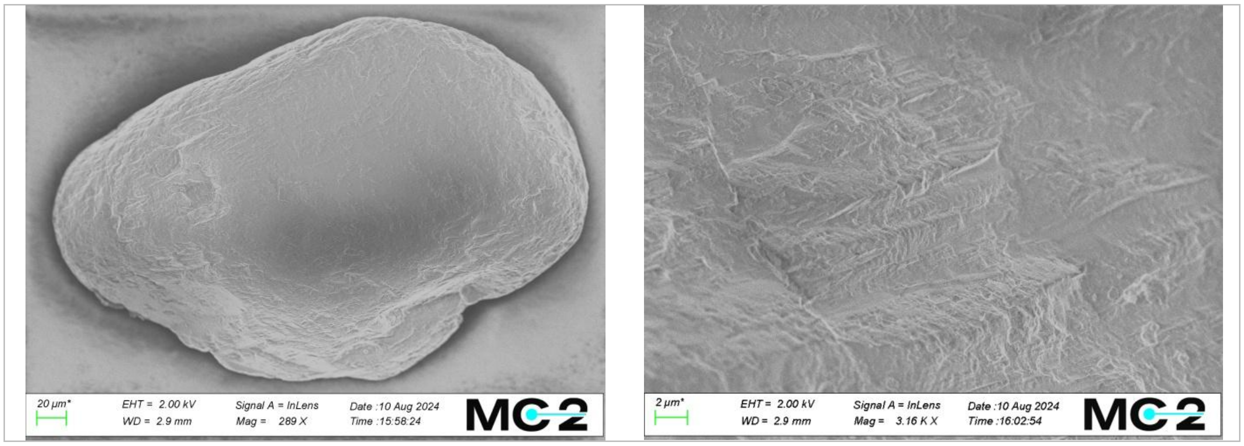

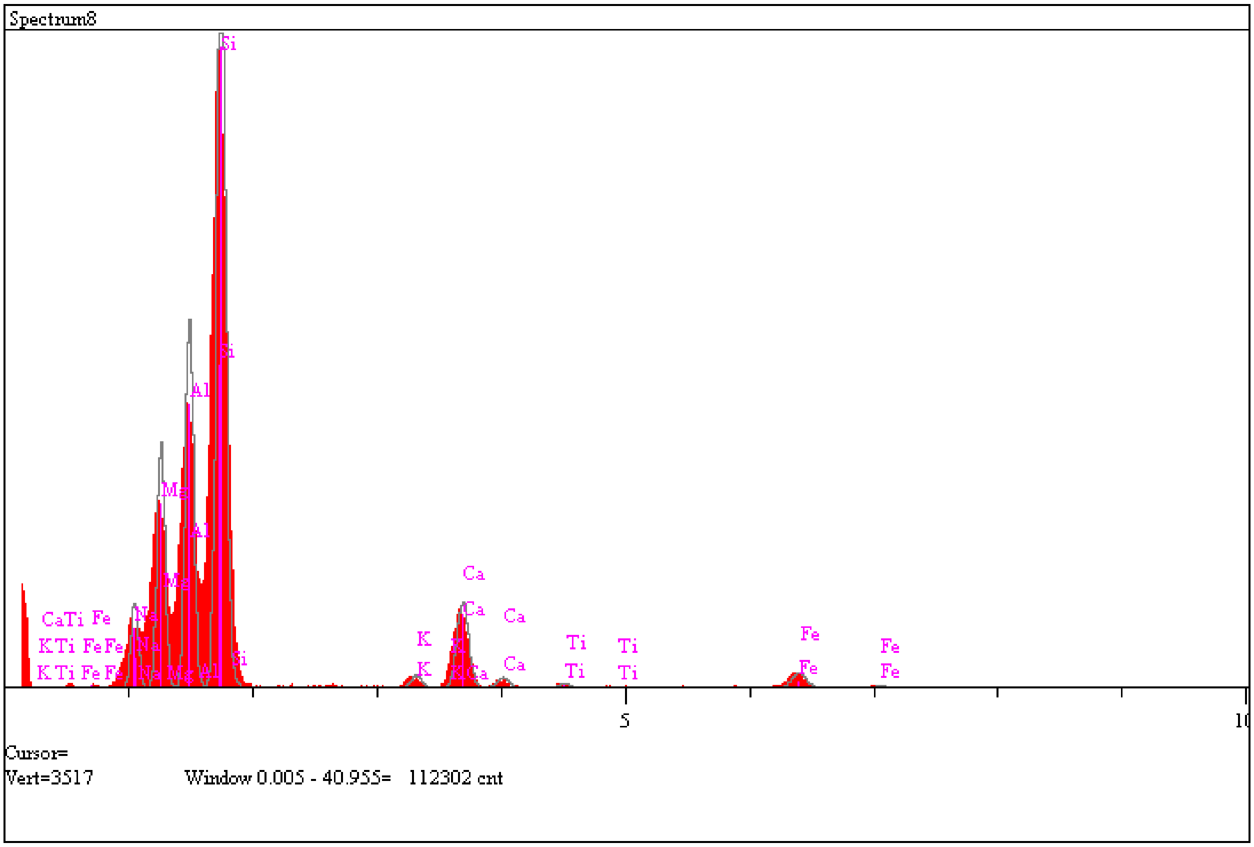

| Elt. | Line | Intensity (c/s) | Error 2-sig | Atomic % | K-Ratio | |
|---|---|---|---|---|---|---|
| Na | Ka | 56.35 | 1.938 | 4.562 | 0.0291 | |
| Mg | Ka | 171.84 | 3.384 | 11.768 | 0.0912 | |
| Al | Ka | 268.31 | 4.229 | 18.094 | 0.1557 | |
| Si | Ka | 639.58 | 6.529 | 46.097 | 0.4137 | |
| K | Ka | 12.57 | 0.915 | 1.156 | 0.0157 | |
| Ca | Ka | 89.12 | 2.437 | 9.178 | 0.1313 | |
| Ti | Ka | 5.25 | 0.592 | 0.725 | 0.0114 | |
| Fe | Ka | 19.72 | 1.146 | 8.421 | 0.1518 | |
| | | | | 100.000 | | Total |

**Natural particle 7 (with no plasma treatment)**

- Exposure to reference lipids mixture, Fig. 5C – s7

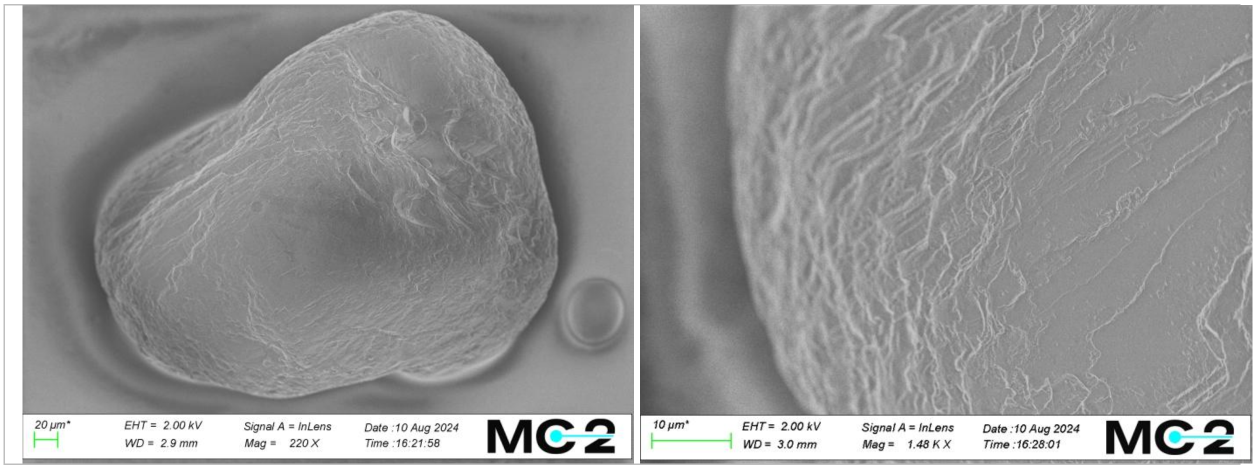

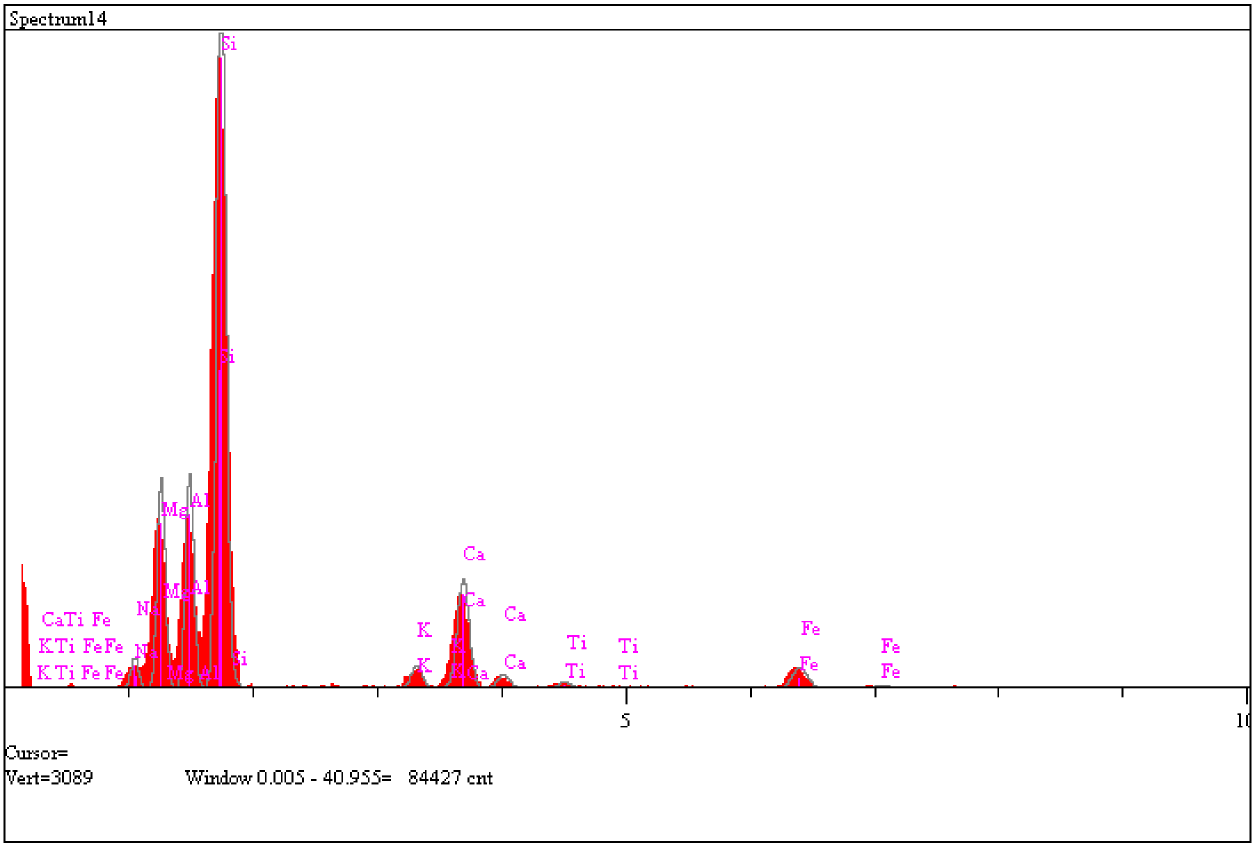

| Elt. | Line | Intensity (c/s) | Error 2-sig | Atomic % | K-Ratio | |
|---|---|---|---|---|---|---|
| Na | Ka | 17.59 | 1.083 | 1.863 | 0.0103 | |
| Mg | Ka | 129.19 | 2.935 | 11.169 | 0.0774 | |
| Al | Ka | 137.35 | 3.026 | 11.428 | 0.0899 | |
| Si | Ka | 556.19 | 6.089 | 47.696 | 0.4061 | |
| K | Ka | 19.74 | 1.147 | 2.174 | 0.0279 | |
| Ca | Ka | 98.52 | 2.563 | 12.228 | 0.1639 | |
| Ti | Ka | 6.08 | 0.637 | 1.013 | 0.0149 | |
| Fe | Ka | 24.12 | 1.268 | 12.430 | 0.2096 | |
| | | | | 100.000 | | Total |

**Natural particle 8 (with no plasma treatment)**

- Exposure to reference lipids mixture, Fig. 5C – s8

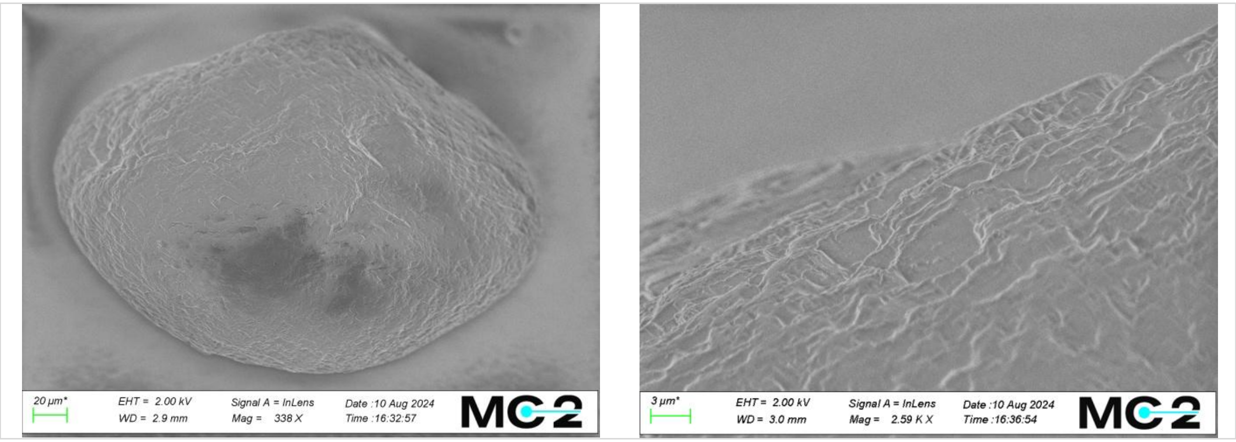

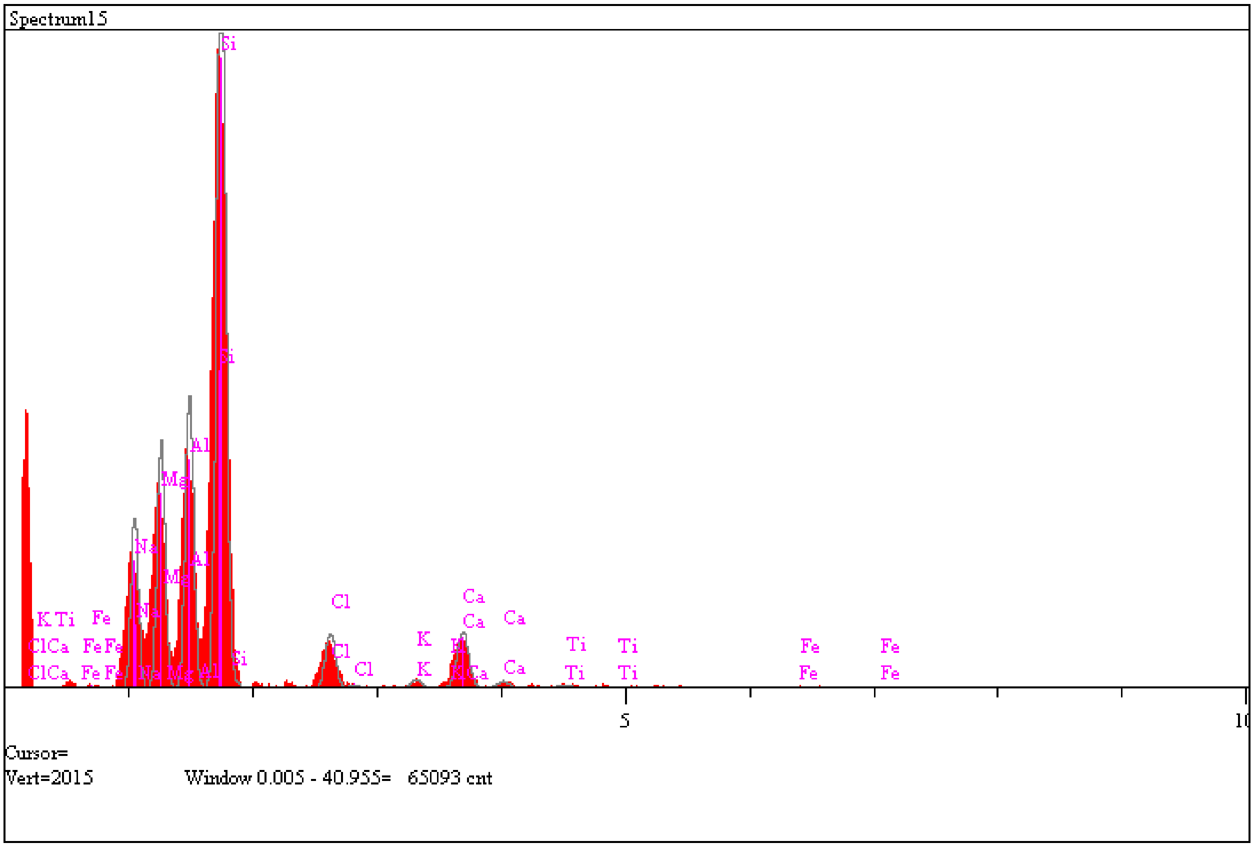

| Elt. | Line | Intensity (c/s) | Error 2-sig | Atomic % | K-Ratio |
|---|---|---|---|---|---|
| Na | Ka | 64.38 | 2.072 | 8.837 | 0.0685 |
| Mg | Ka | 99.00 | 2.569 | 12.344 | 0.1082 |
| Al | Ka | 121.87 | 2.850 | 15.410 | 0.1455 |
| Si | Ka | 364.01 | 4.926 | 49.729 | 0.4847 |
| Cl | Ka | 28.19 | 1.371 | 4.546 | 0.0542 |
| K | Ka | 4.83 | 0.567 | 0.887 | 0.0124 |
| Ca | Ka | 33.34 | 1.491 | 6.823 | 0.1011 |
| Ti | Ka | 2.14 | 0.378 | 0.587 | 0.0095 |
| Fe | Ka | 0.99 | 0.257 | 0.836 | 0.0157 |

| | | | | | | |
|---|---|---|---|---|---|---|
| | | | | 100.000 | | Total |

**Natural particle 9 (with no plasma treatment)**

- Exposure to reference lipids mixture, Fig. 5C – s9

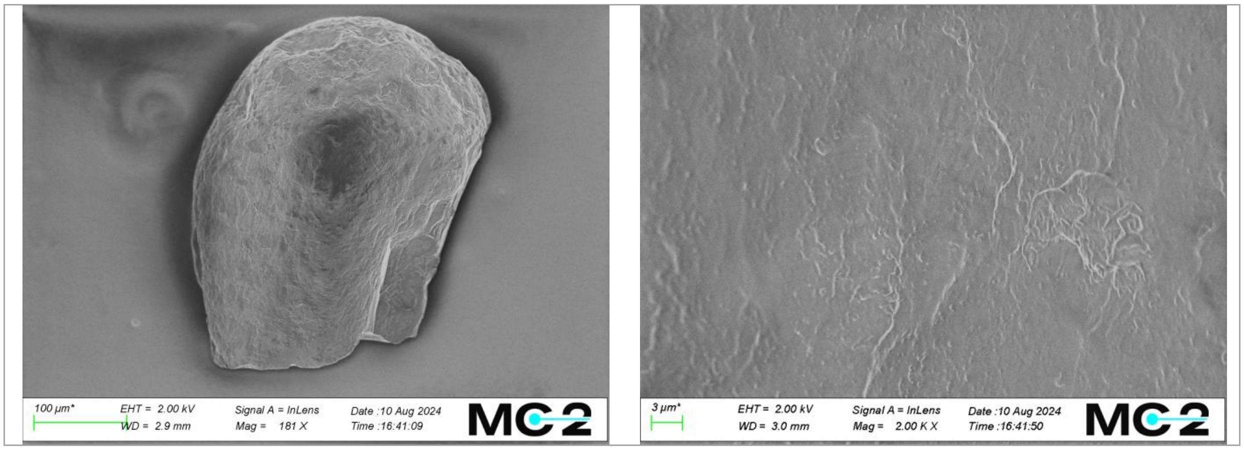

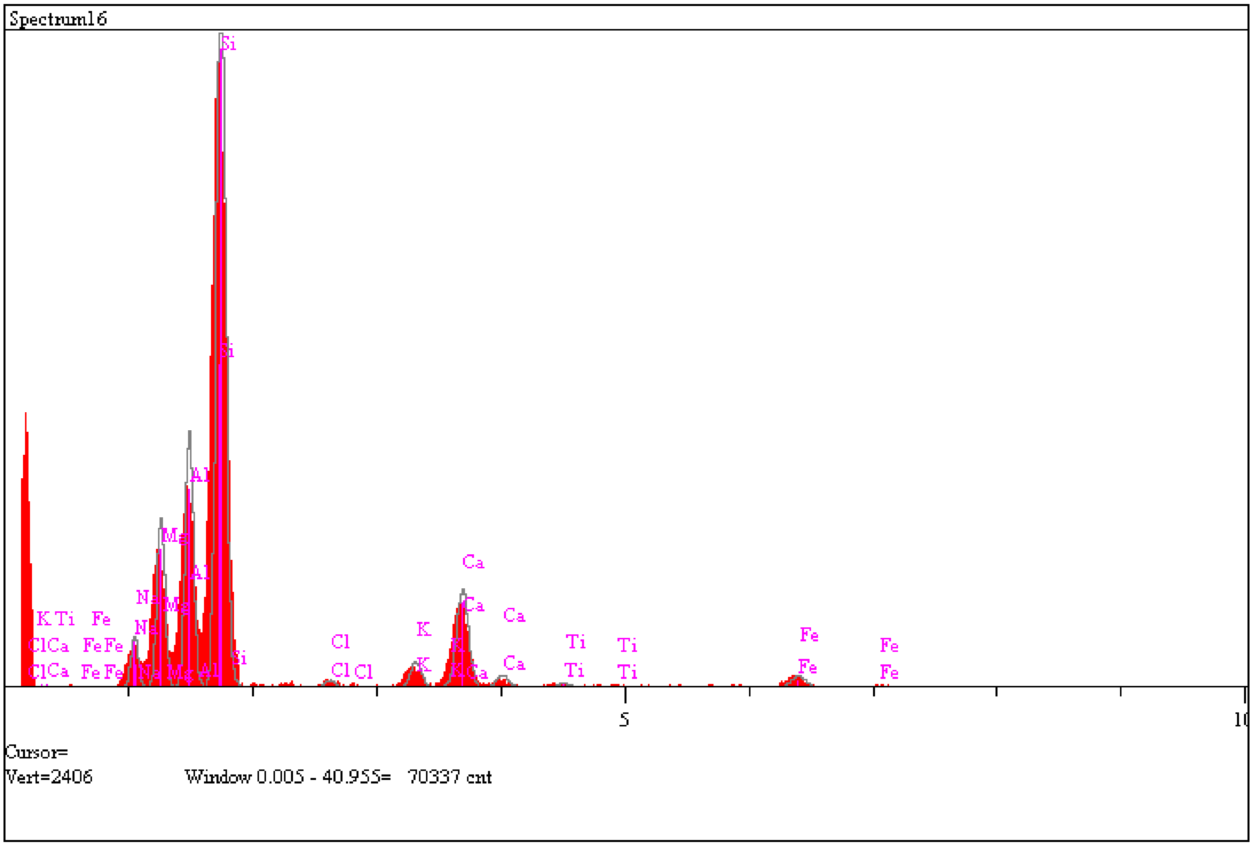

| Elt. | Line | Intensity (c/s) | Error 2-sig | Atomic % | K-Ratio |
|---|---|---|---|---|---|
| Na | Ka | 22.89 | 1.235 | 3.113 | 0.0190 |
| Mg | Ka | 81.32 | 2.328 | 9.250 | 0.0694 |
| Al | Ka | 127.43 | 2.914 | 14.064 | 0.1188 |
| Si | Ka | 430.03 | 5.354 | 50.229 | 0.4470 |
| Cl | Ka | 4.41 | 0.542 | 0.598 | 0.0066 |
| K | Ka | 16.94 | 1.063 | 2.591 | 0.0341 |
| Ca | Ka | 68.85 | 2.142 | 11.865 | 0.1630 |
| Ti | Ka | 3.30 | 0.469 | 0.765 | 0.0115 |
| Fe | Ka | 10.55 | 0.839 | 7.525 | 0.1306 |

| | | | | | | |
|---|---|---|---|---|---|---|
| | | | | 100.000 | | Total |

### 4. All images used in Fig. 5

The images are seperately attached as supporting information.
