## Supplementary figures and images for "Protocell formation on micrometeorites"

### 1_1_58.986.tif

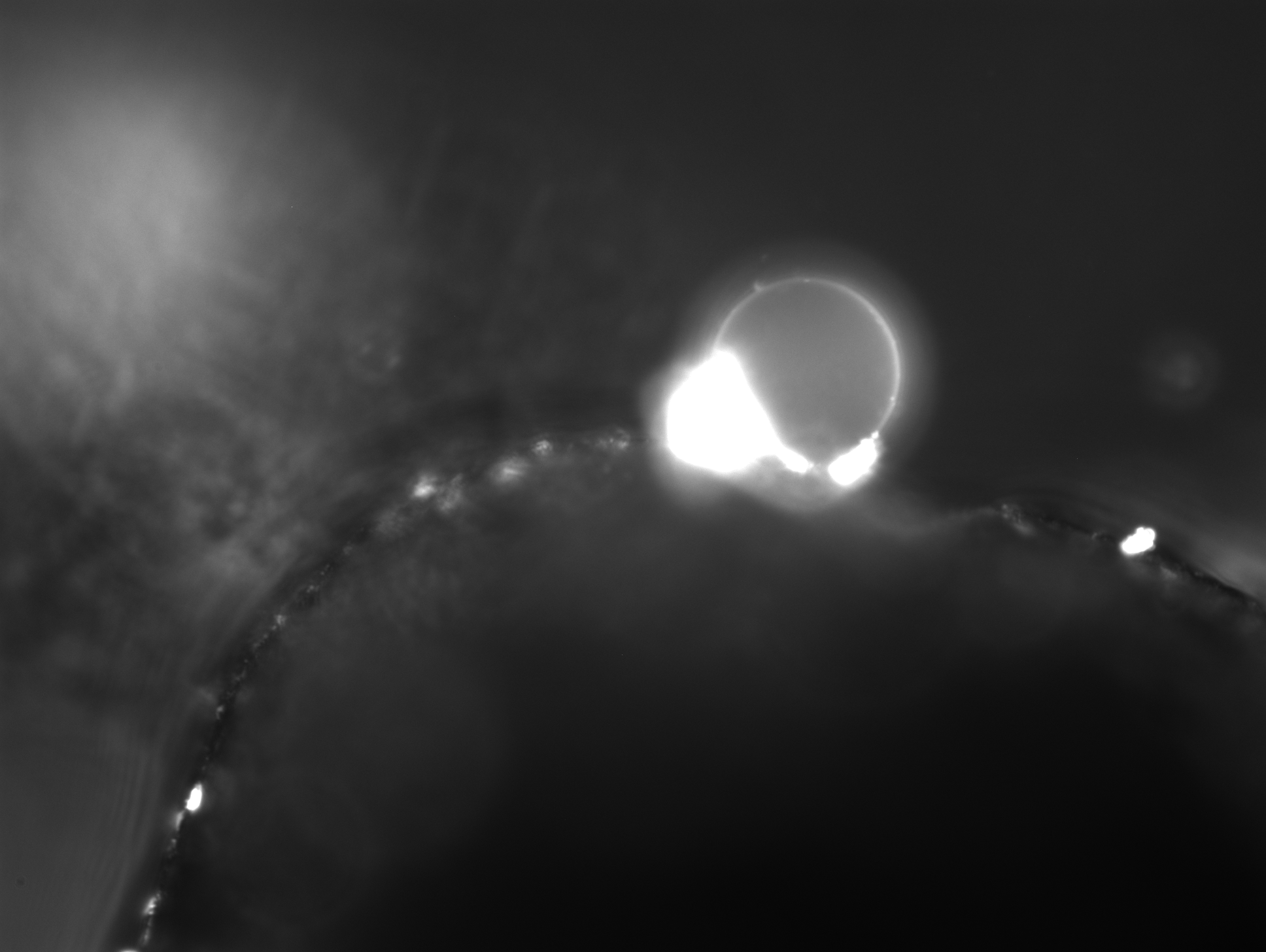

### 1_11_07.635.tif

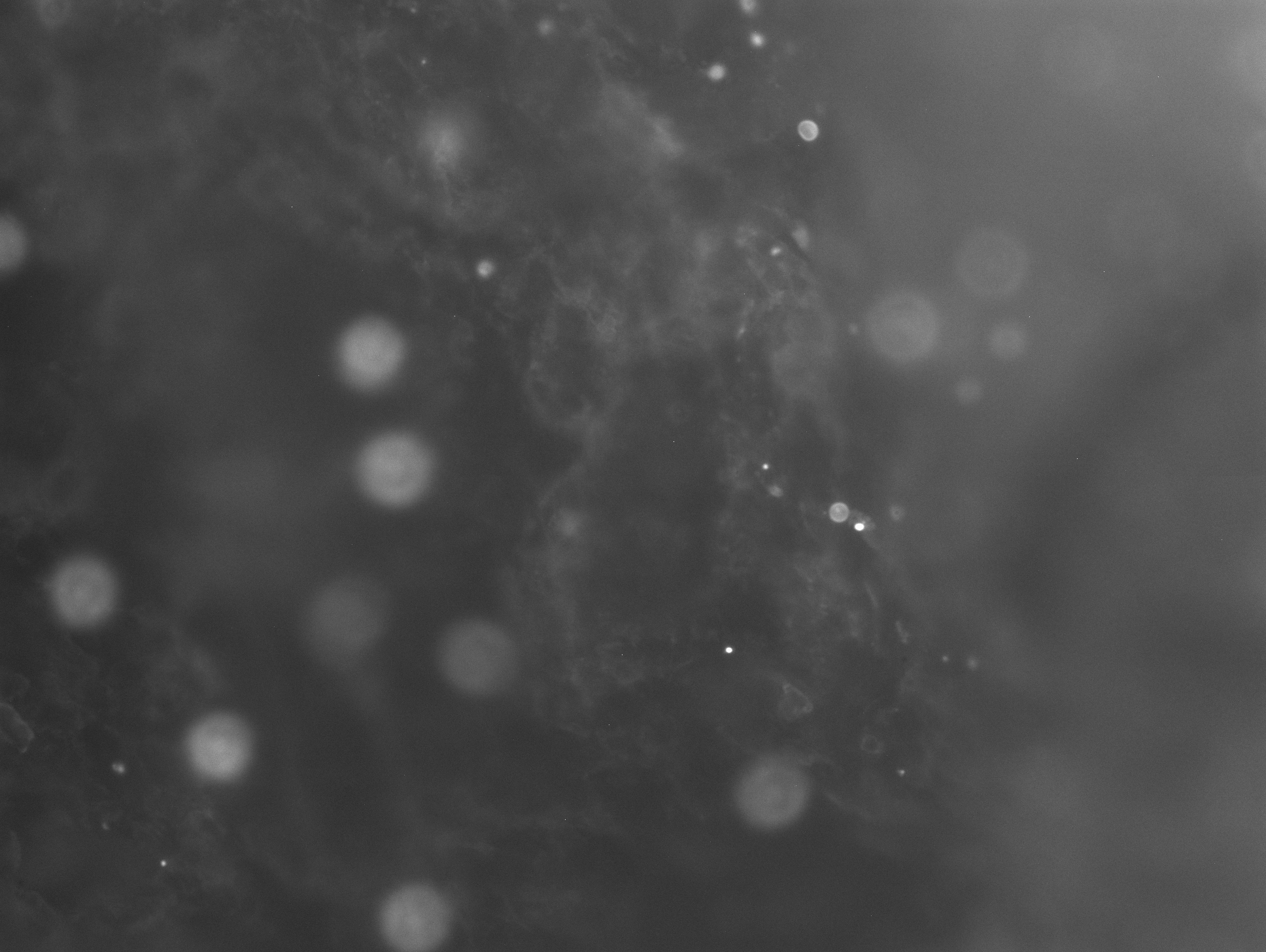

### 1_16_39.580.tif

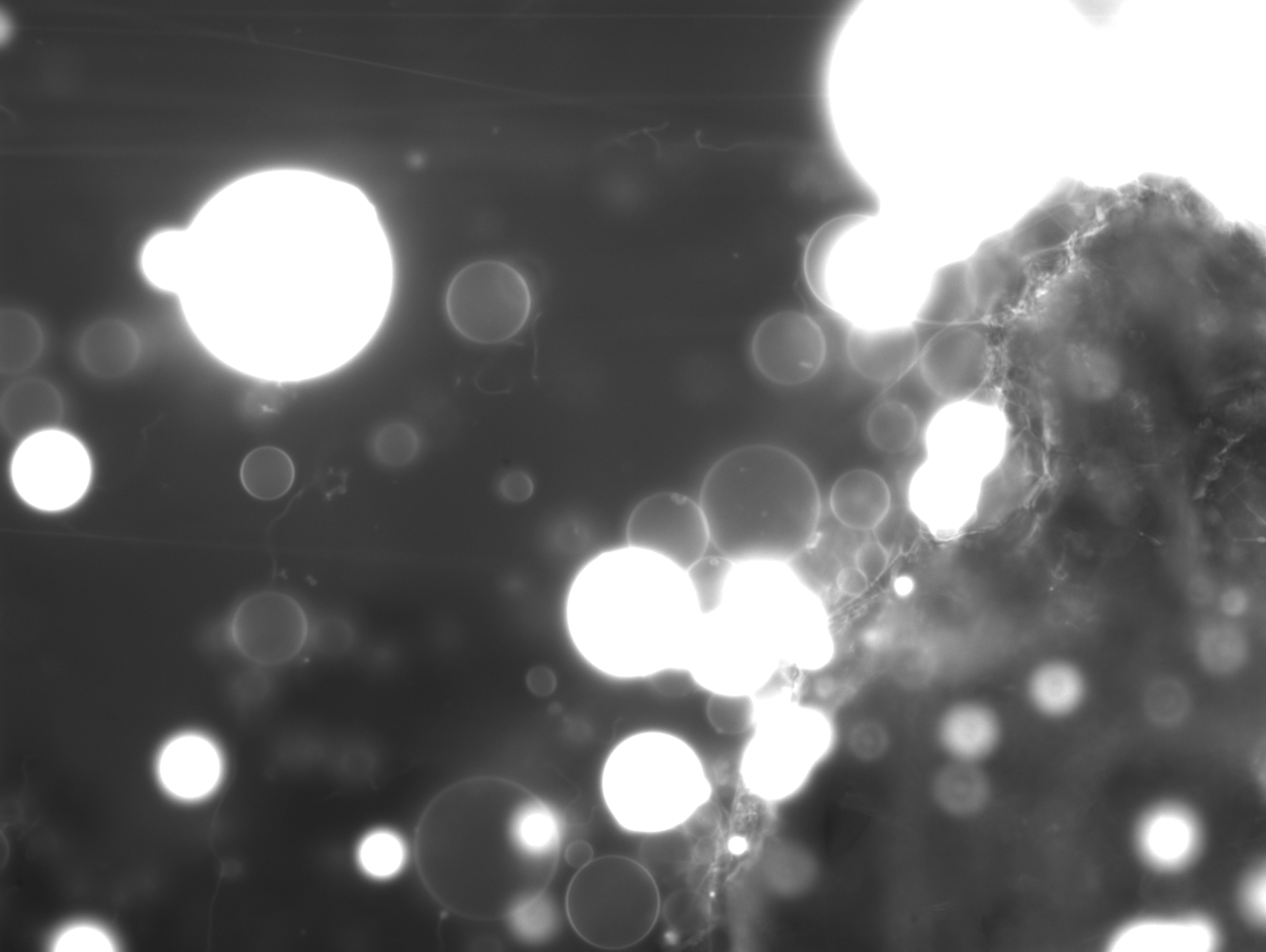

### 1_20_42.053.tif

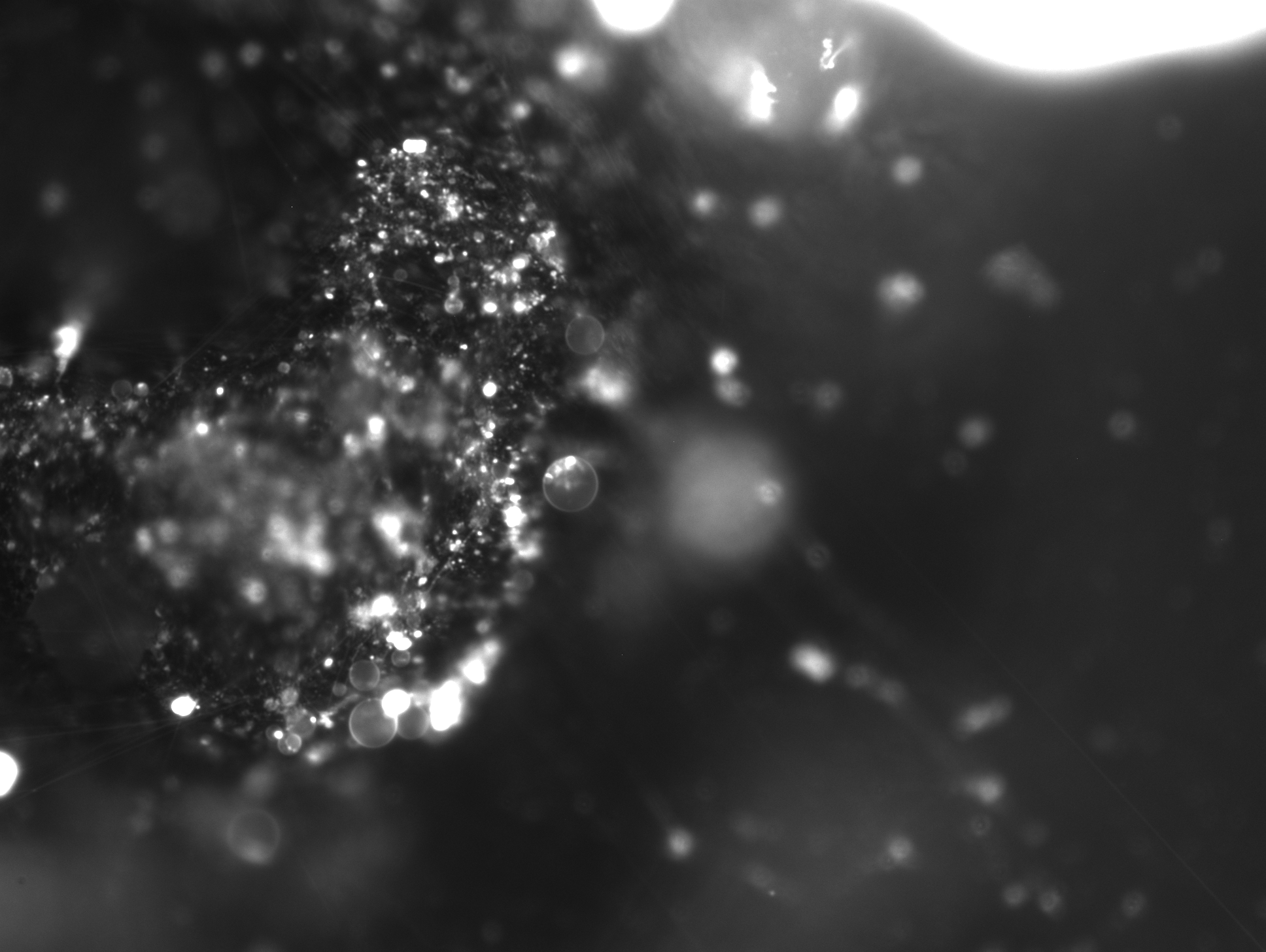

### 1_23_44.971.tif

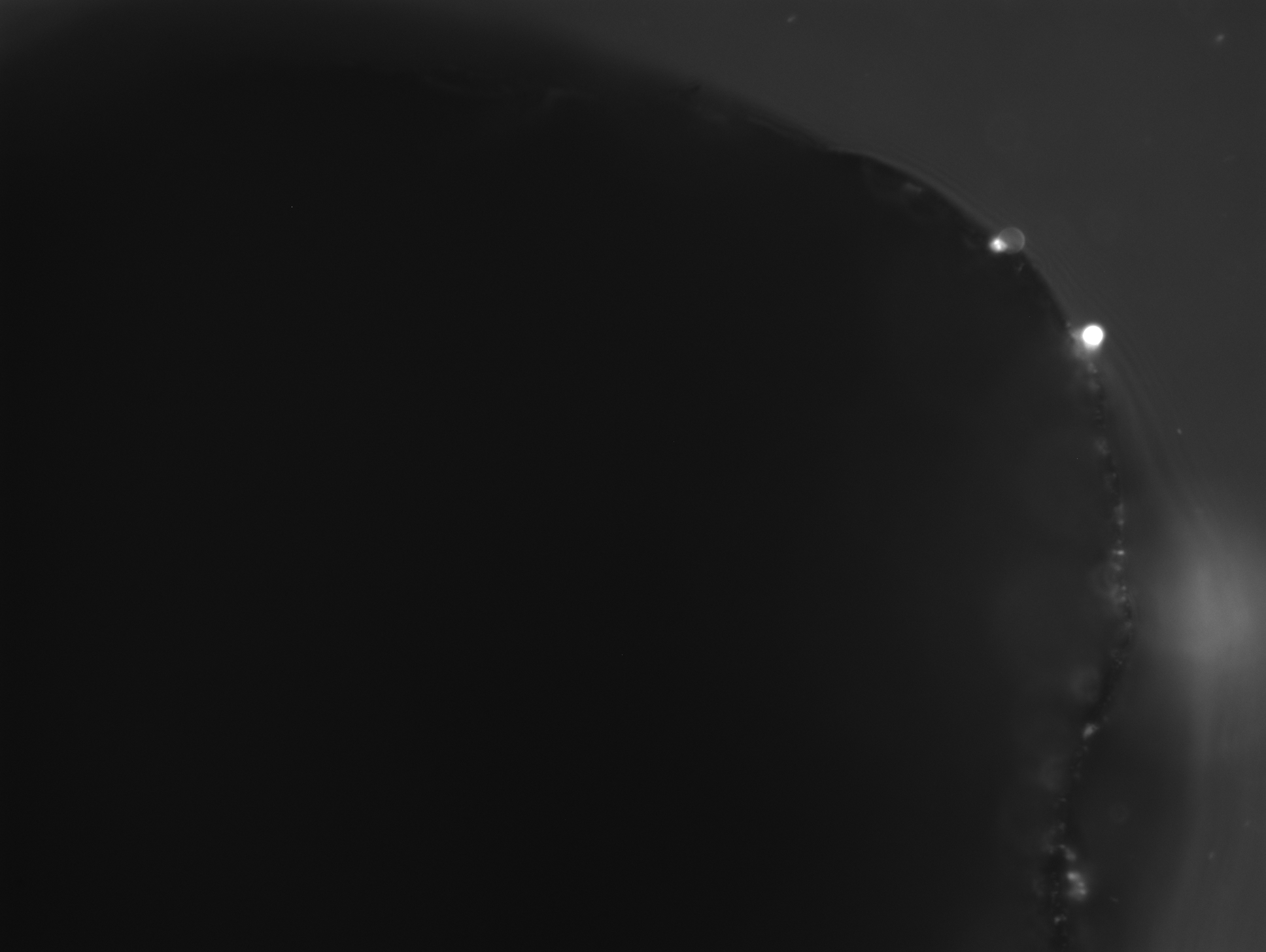

### 1_28_40.081.tif

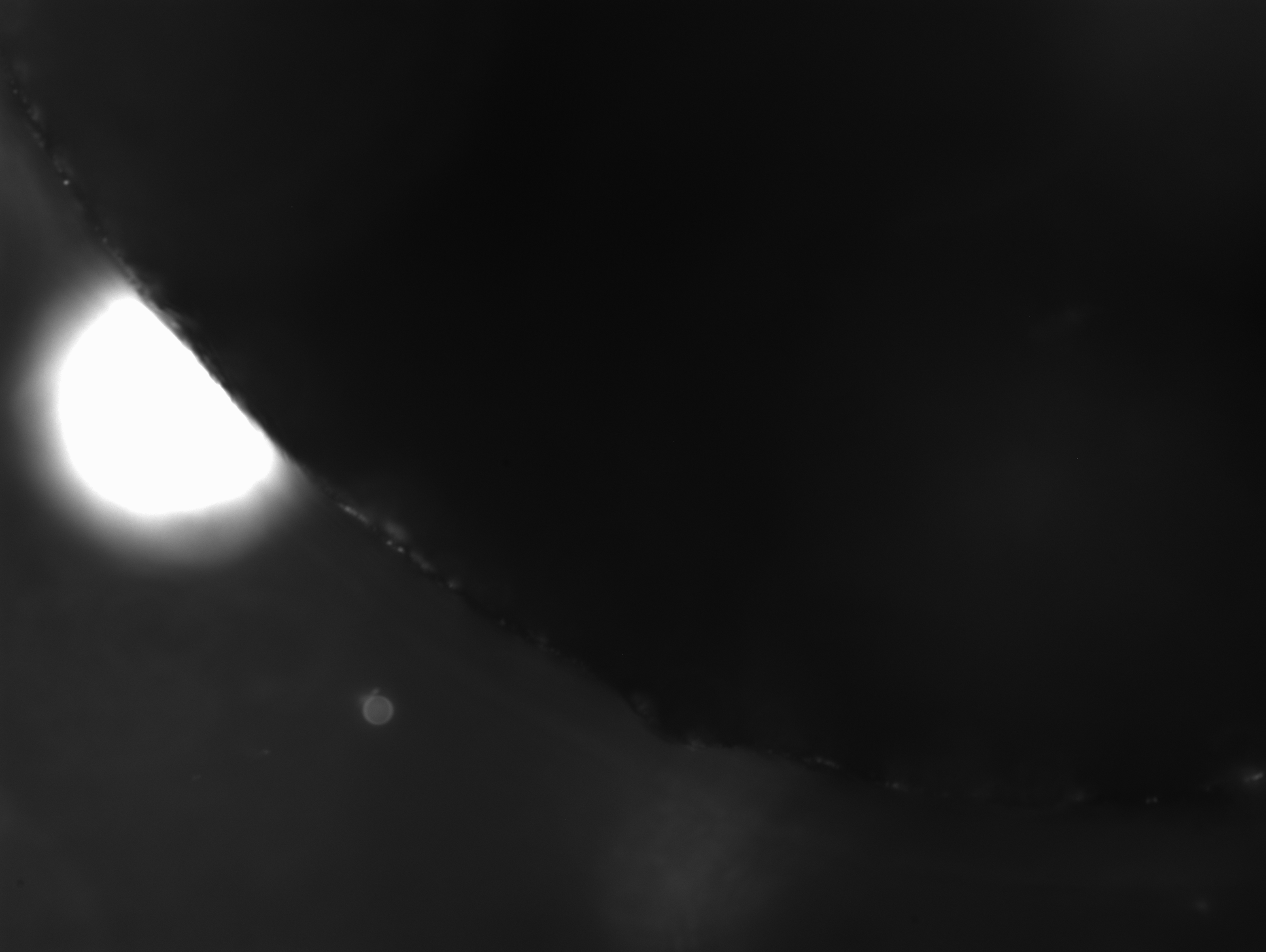

### 1_31_44.811.tif

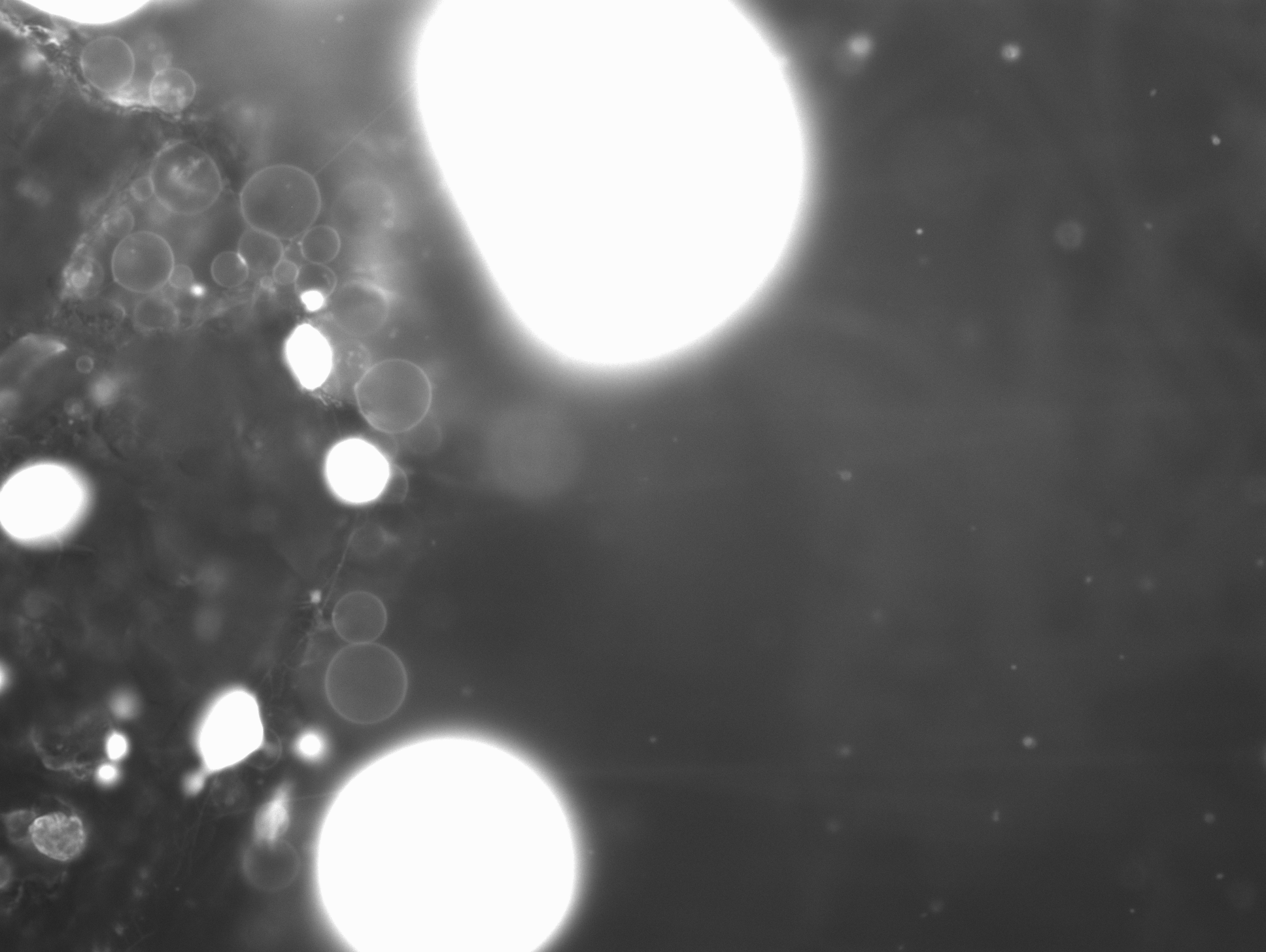

### 1_37_52.531.tif

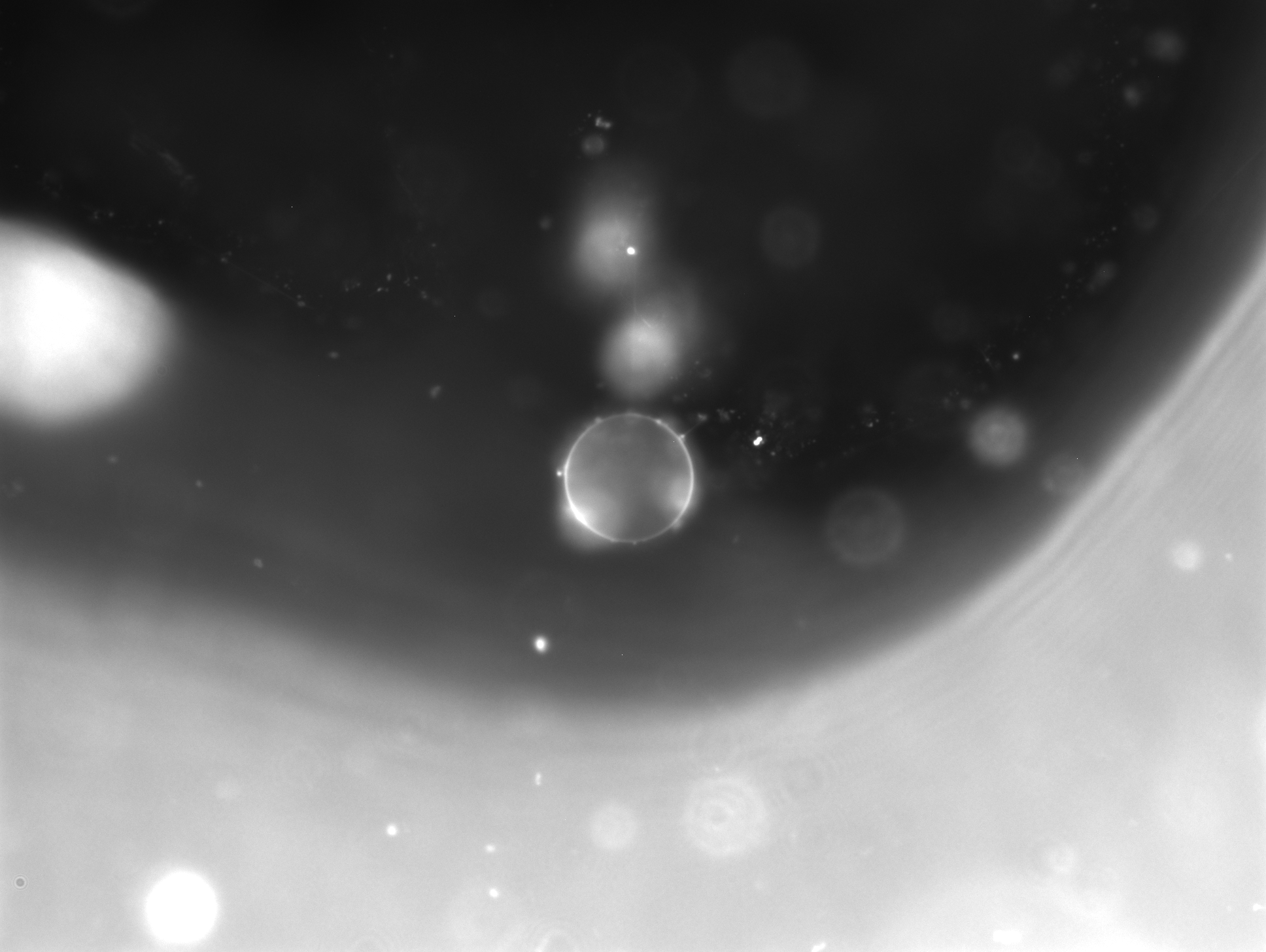

### 1_40.002.tif

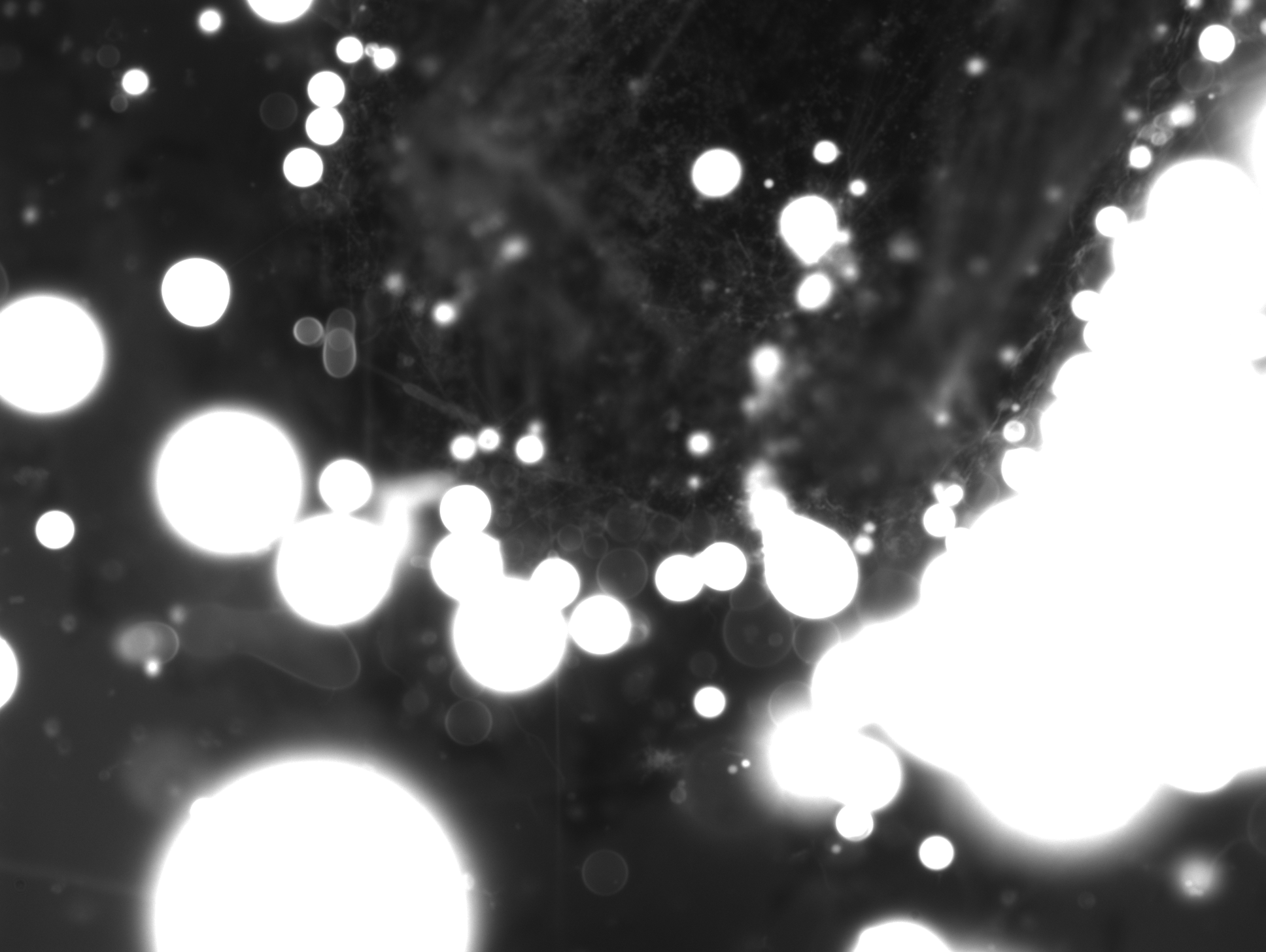

### 1_40.581-1.tif

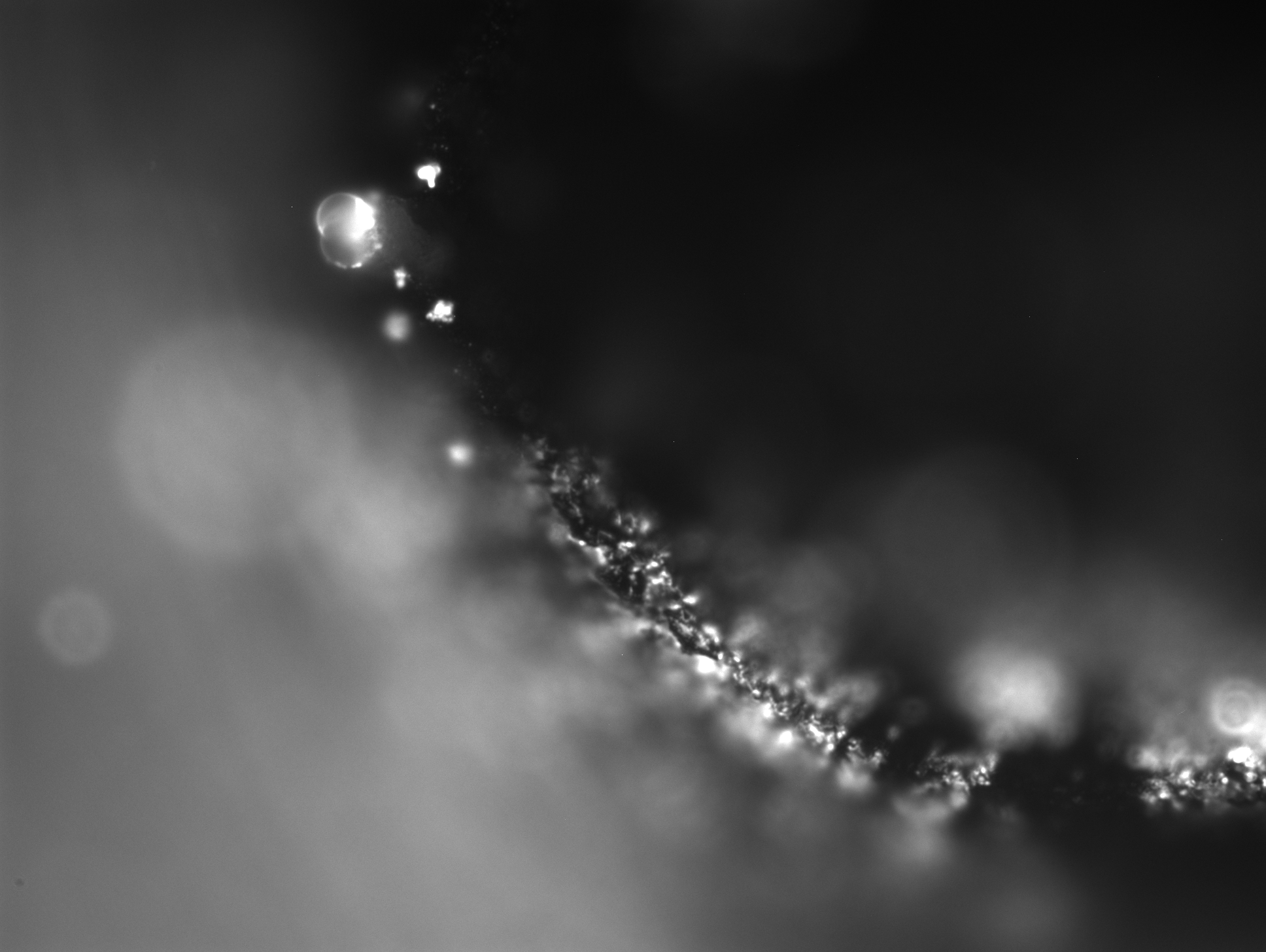

### 1_42_36.540.tif

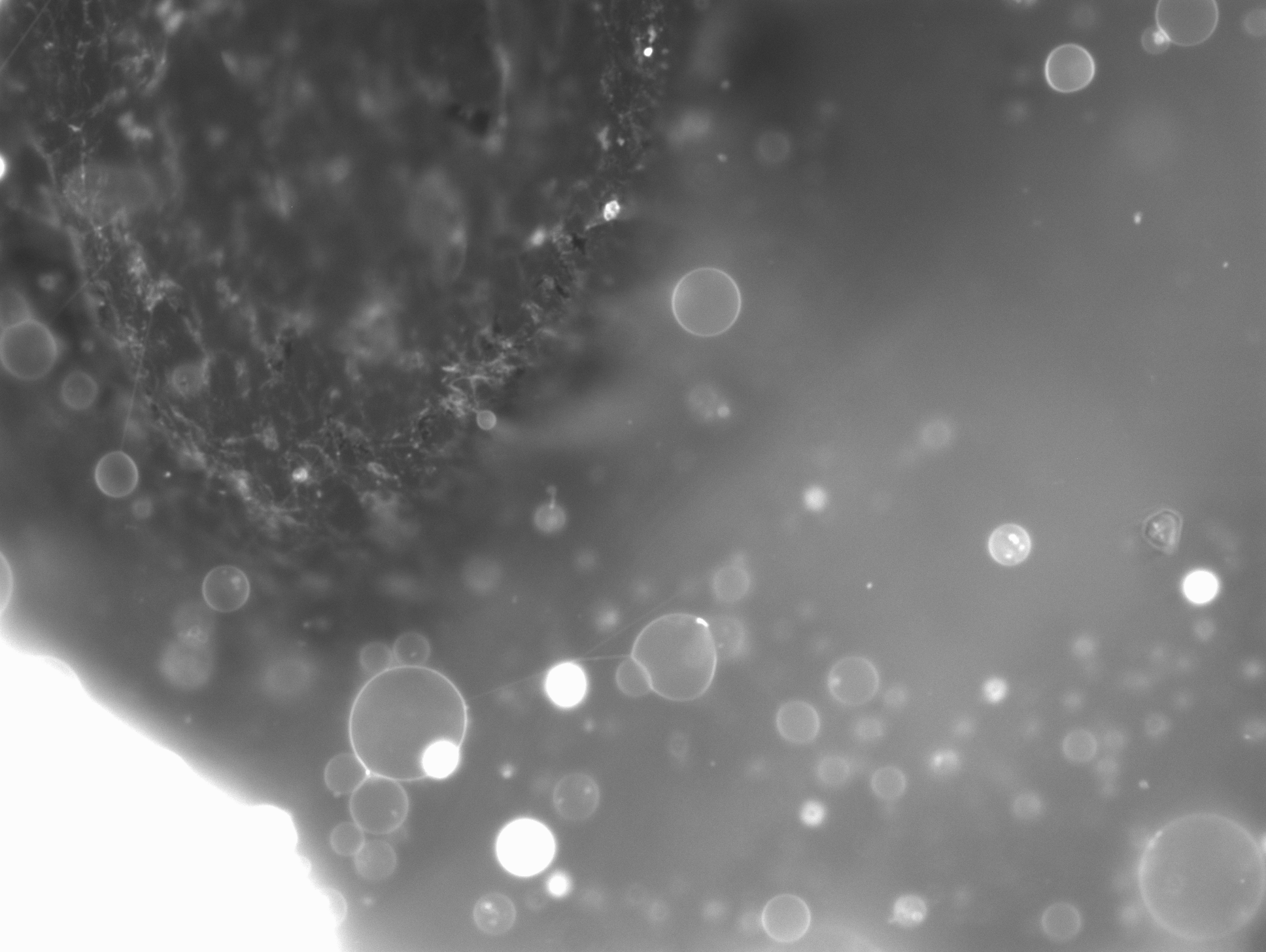

### 1_44.502-1.tif

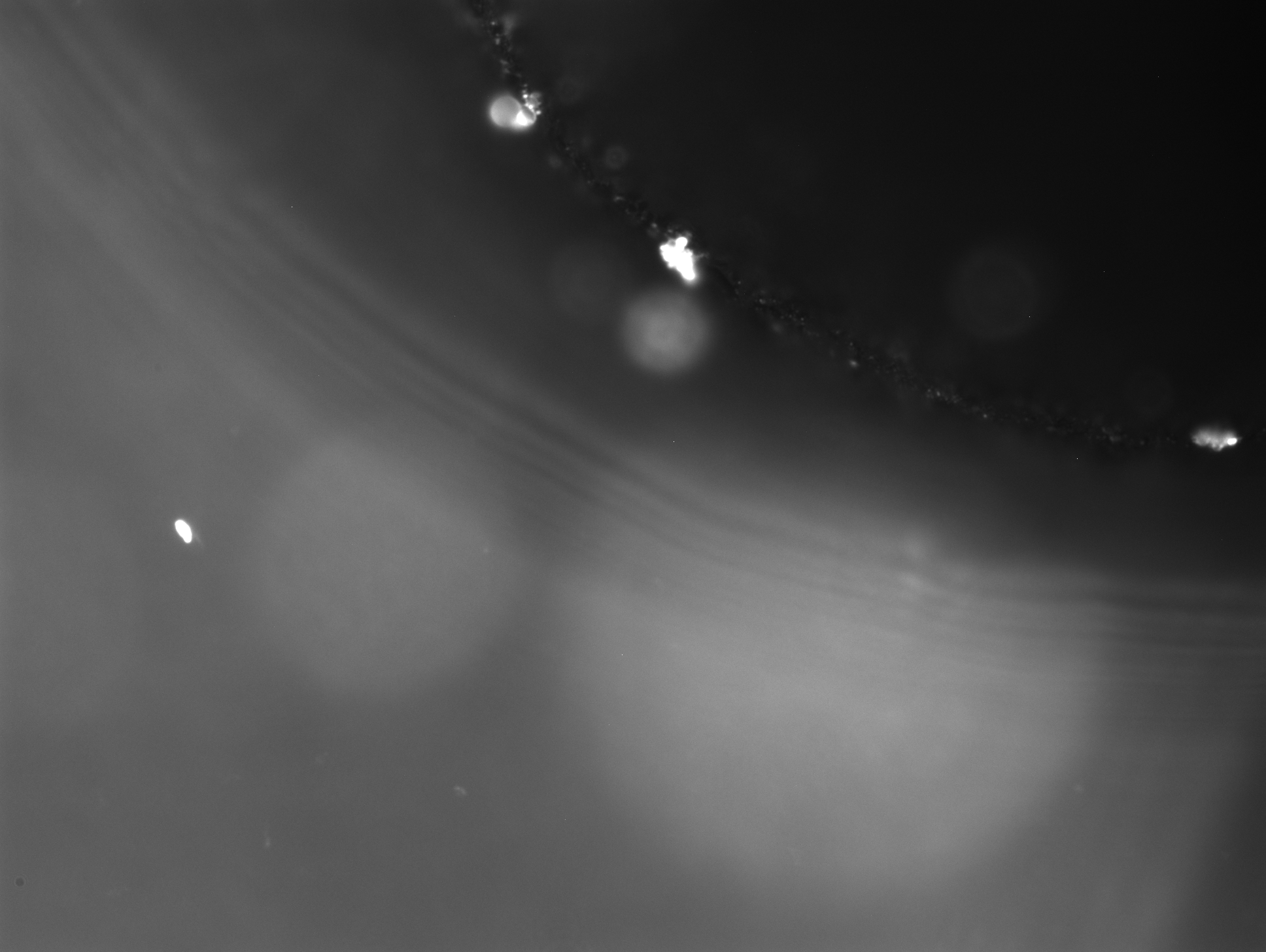

### 1_52_54.811.tif

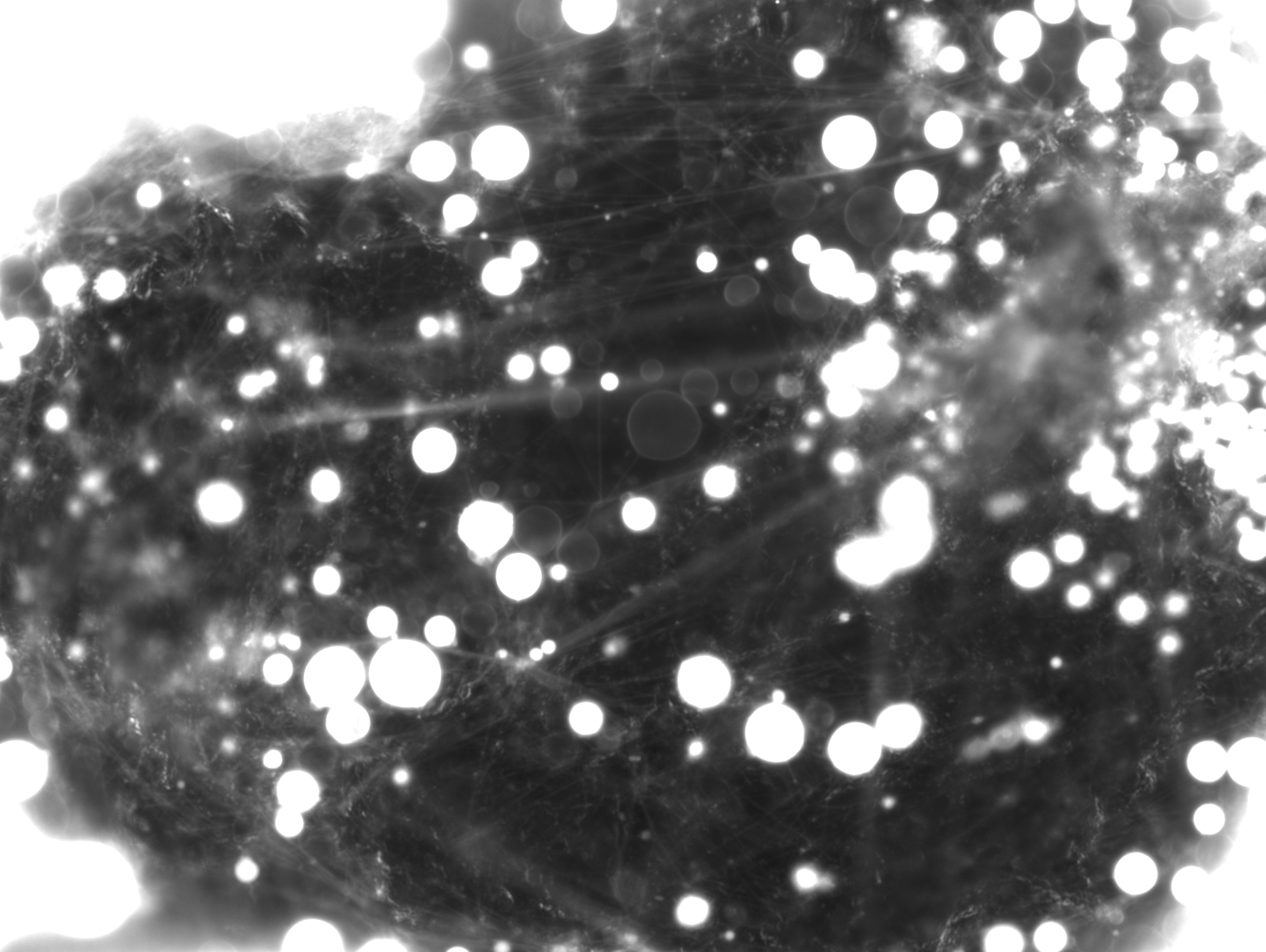

### 1_53_42.138.tif

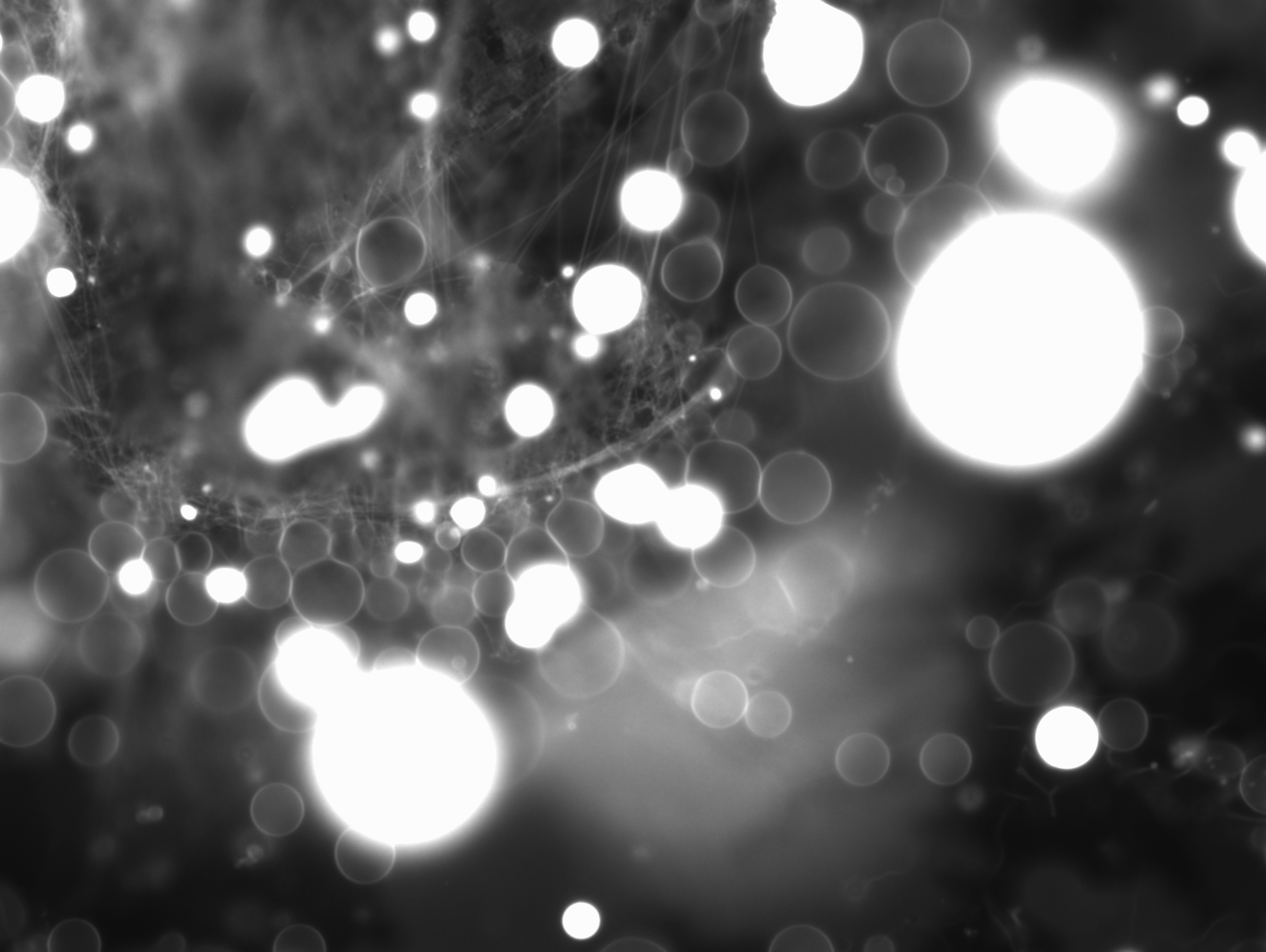

### 1_57_21.553.tif

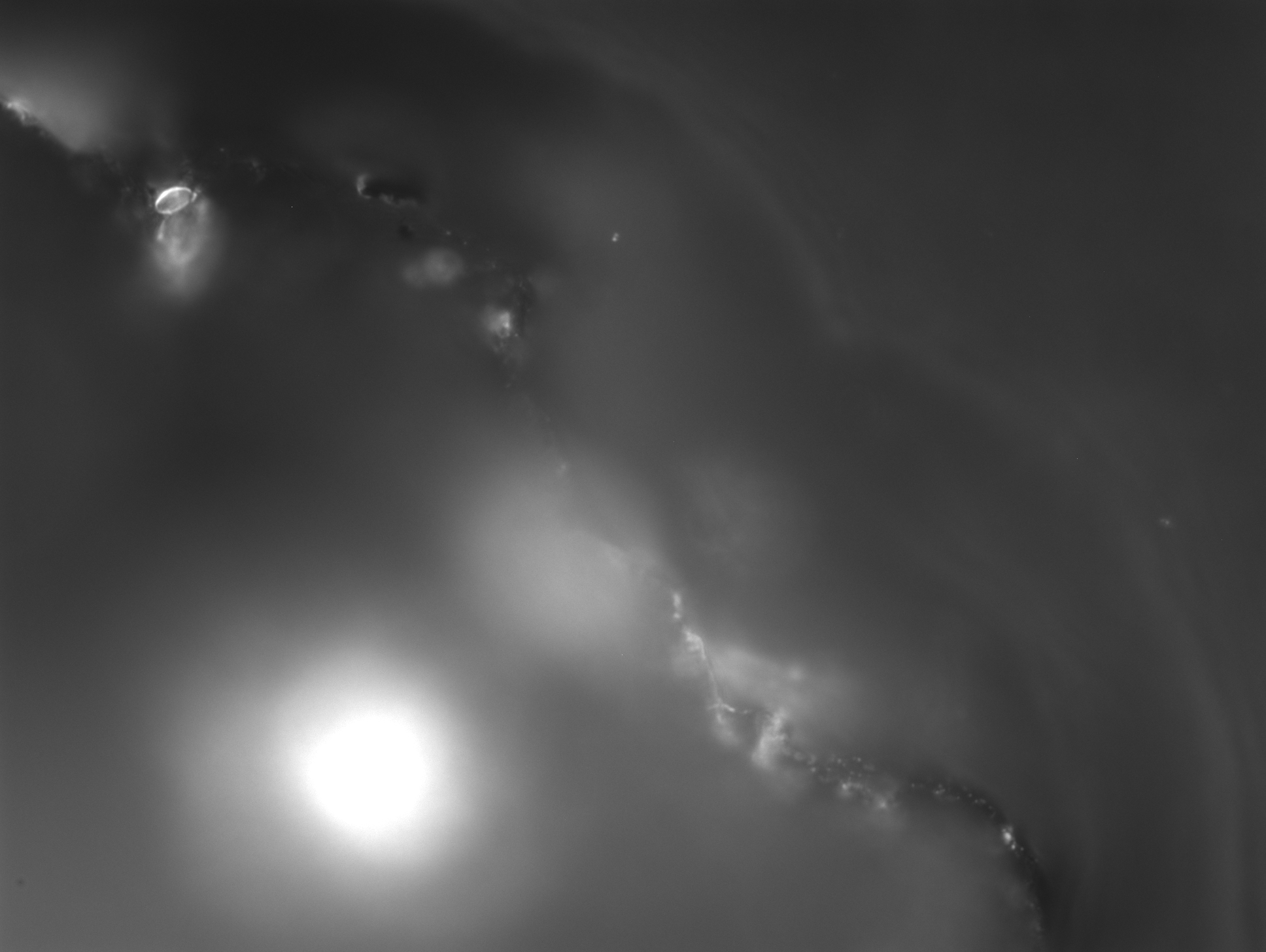

### 1_58_56.395.tif

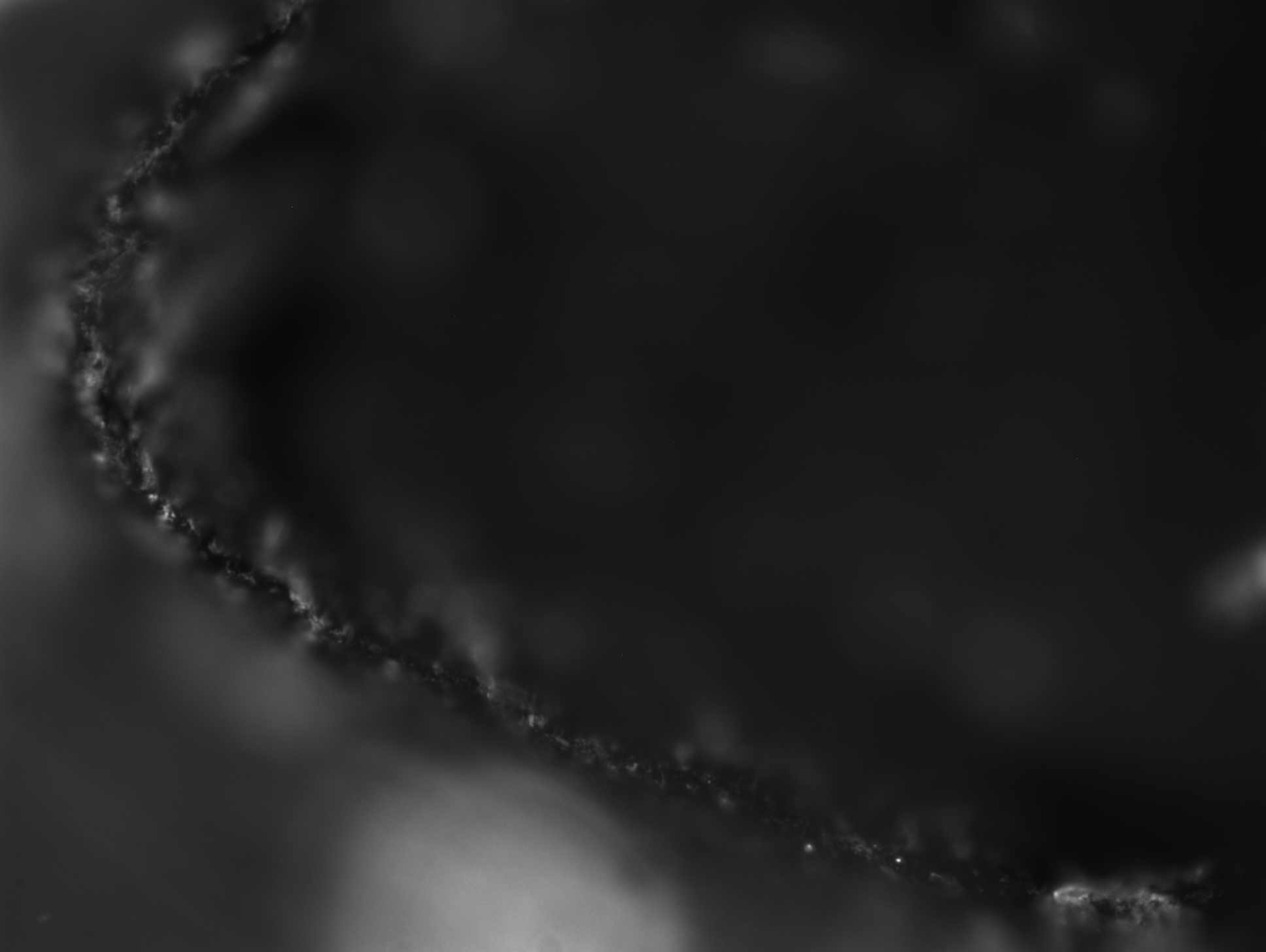

### 1_74_58.164.tif

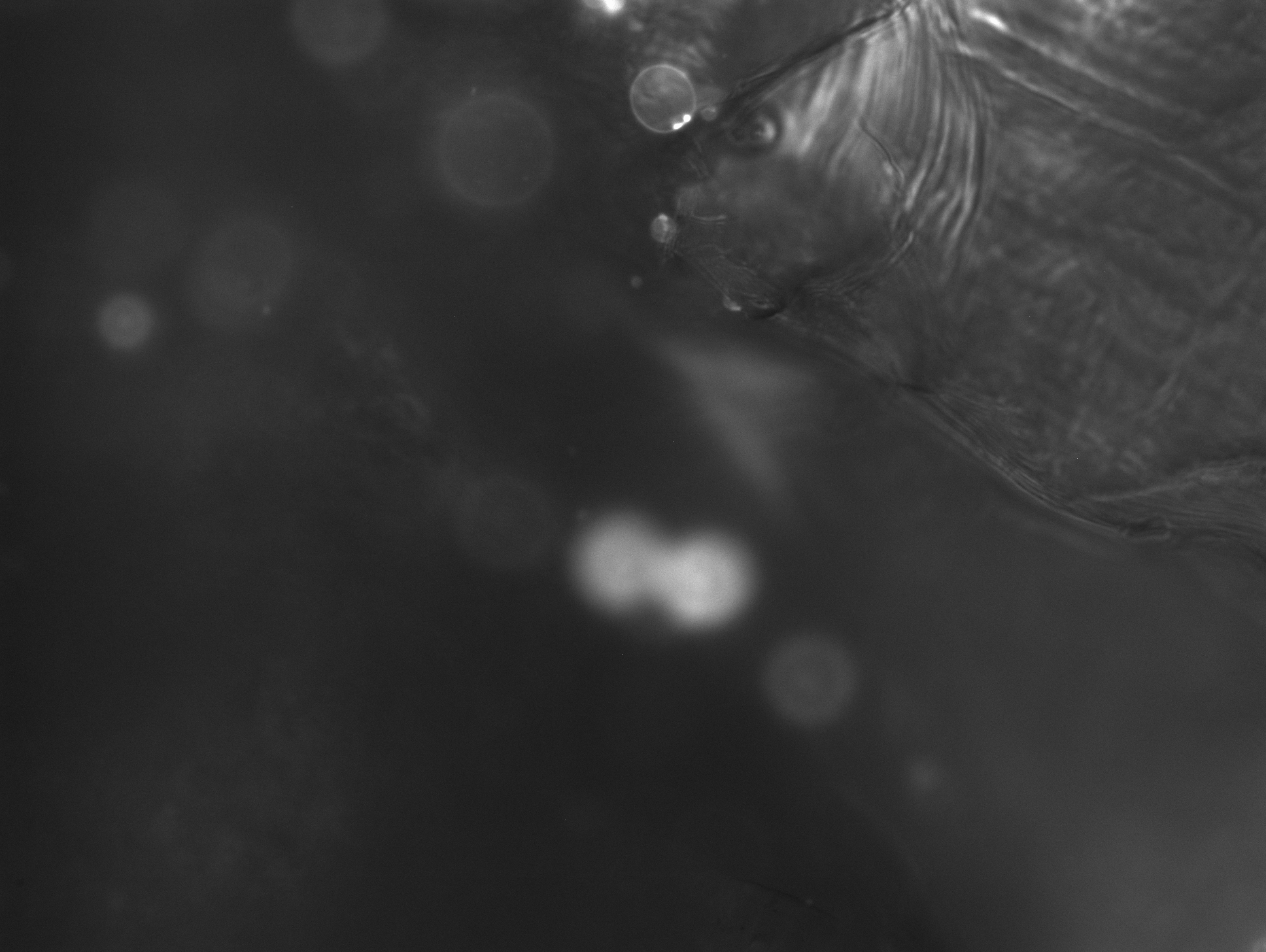

### 1_75_12.251.tif

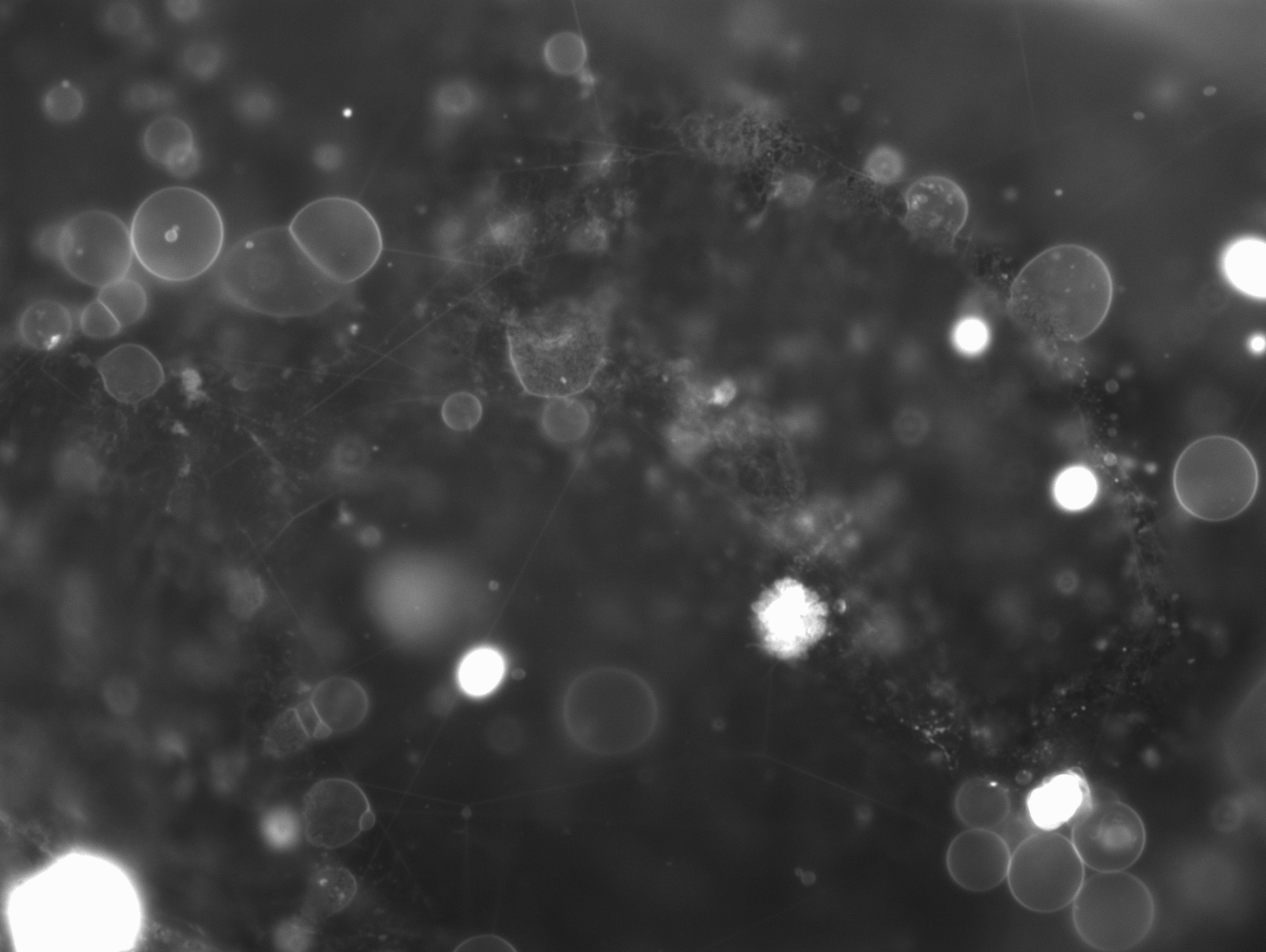

### 1_77_17.910.tif

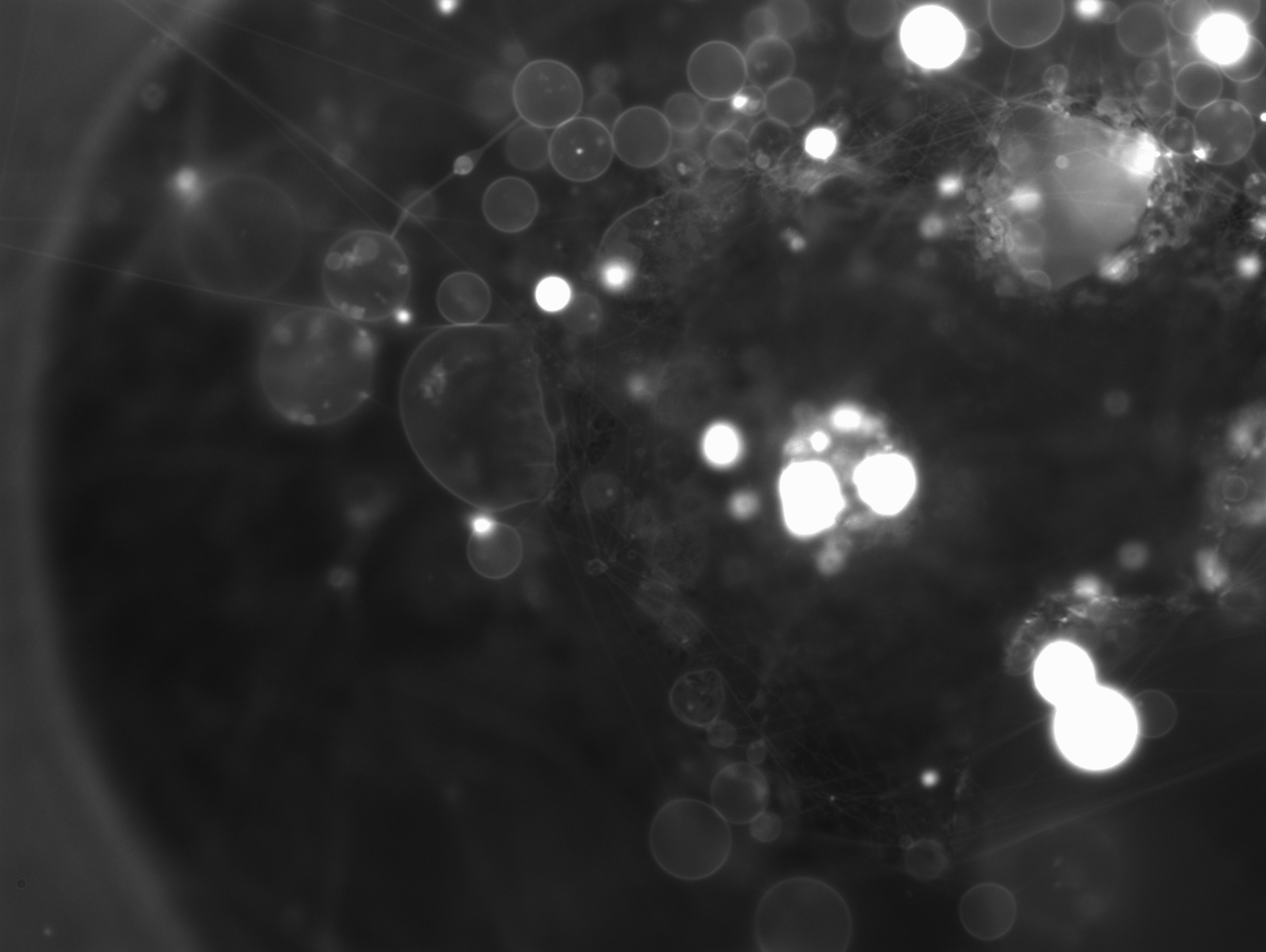

### 1_88_16.009.tif

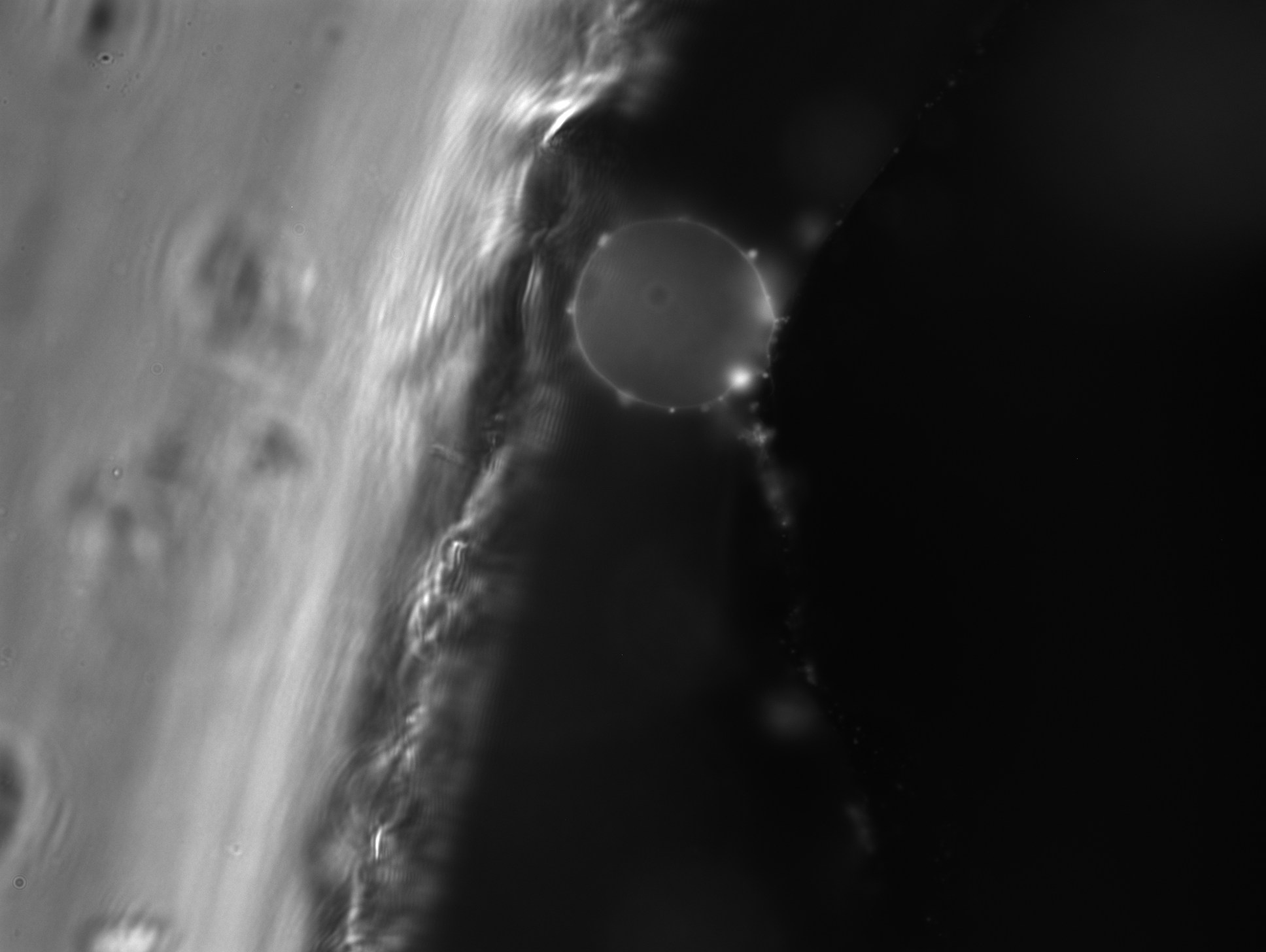

### 1_110_40.299.tif

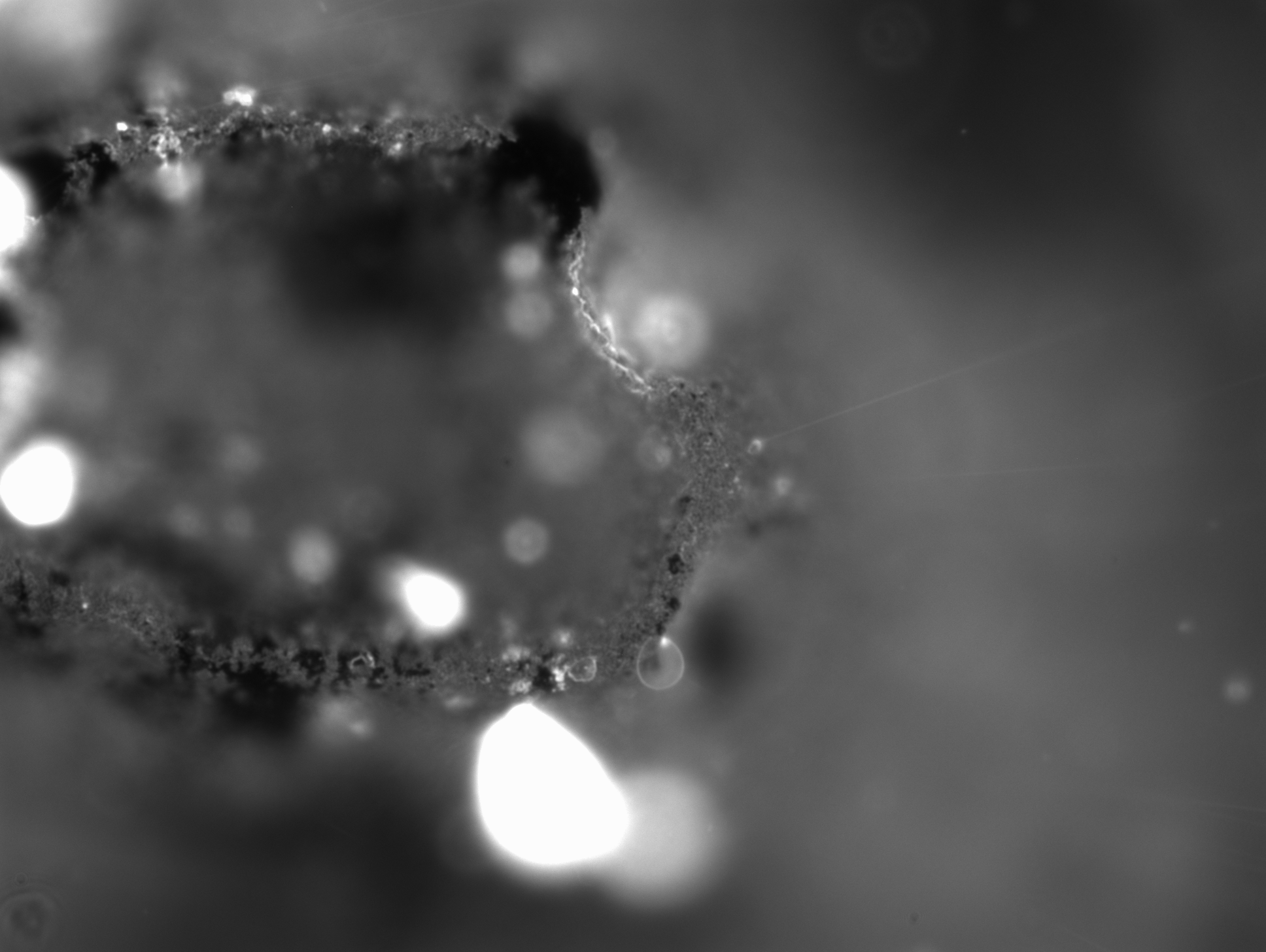

### 1_120_23.472.tif

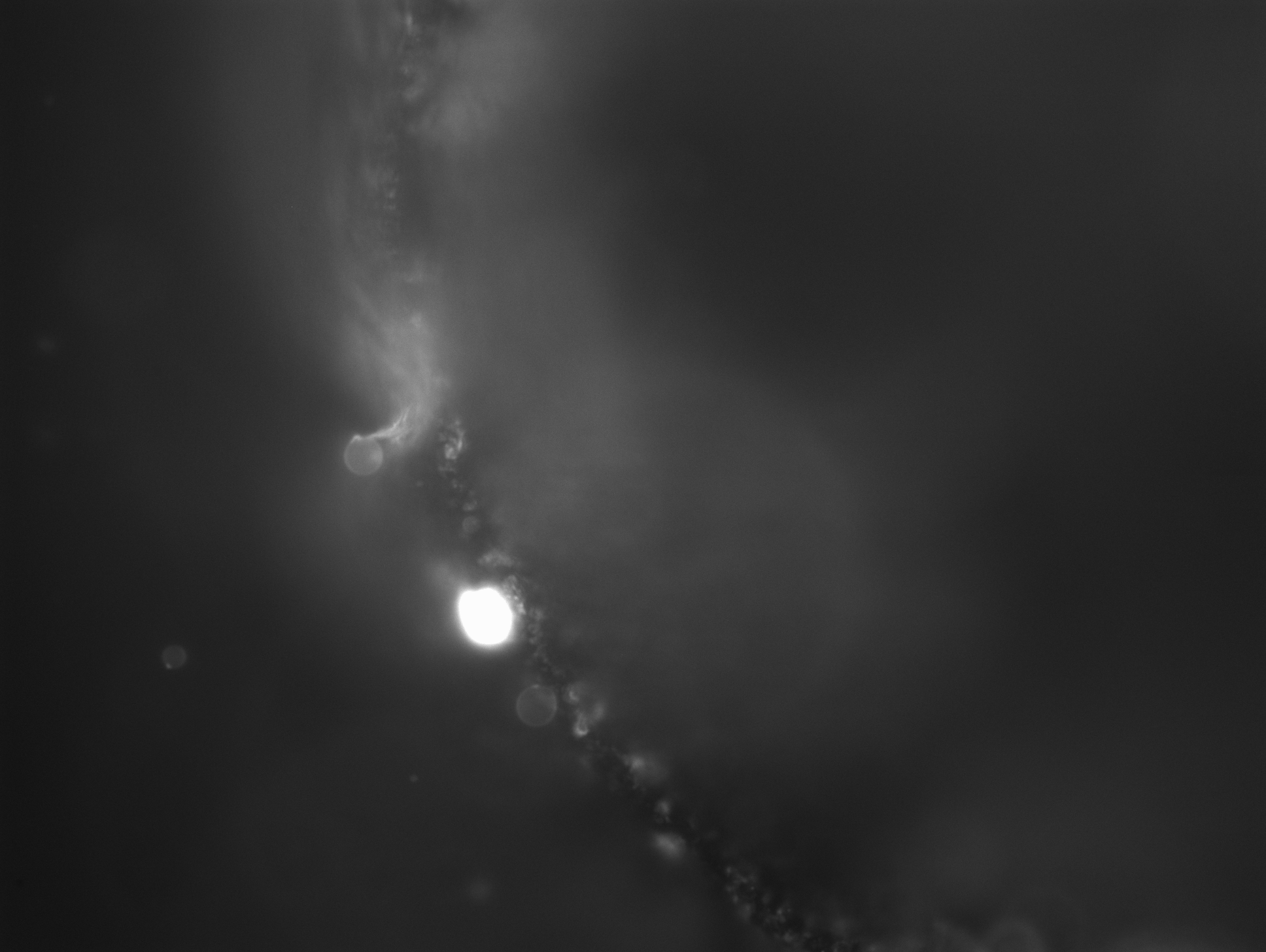

### 1_141_50.833.tif

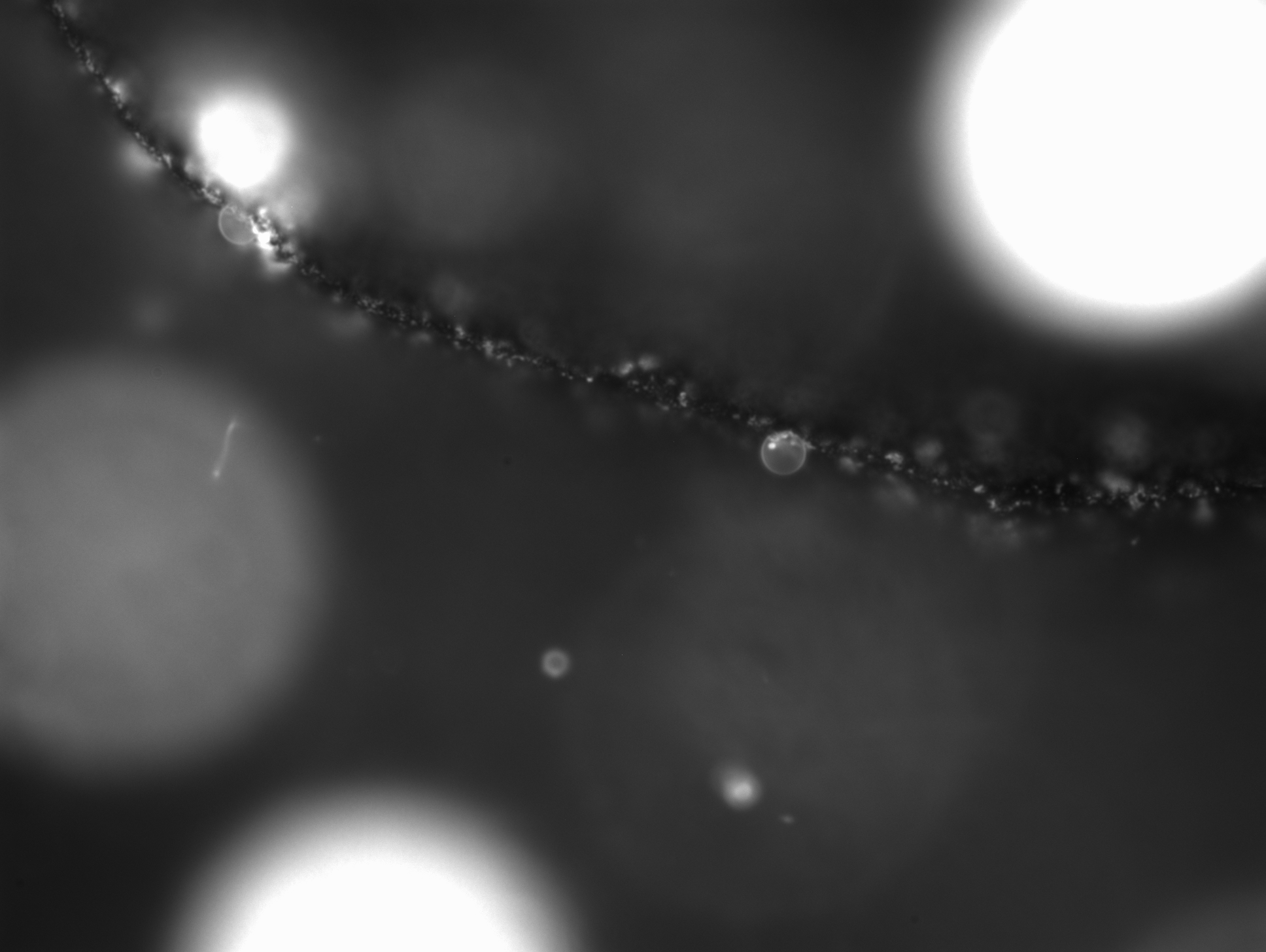

### 1_154_46.961.tif

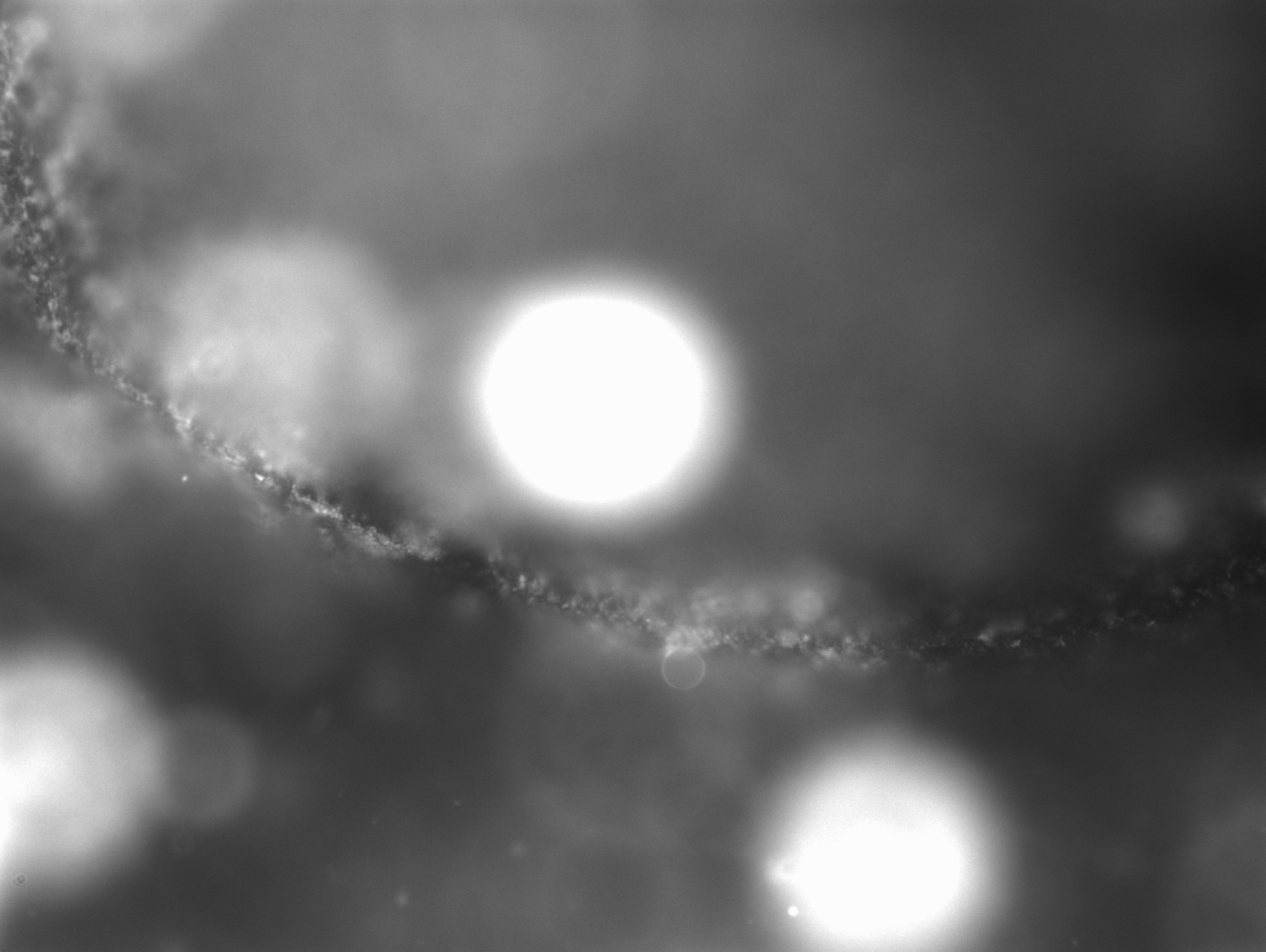

### 1_203_30.714.tif

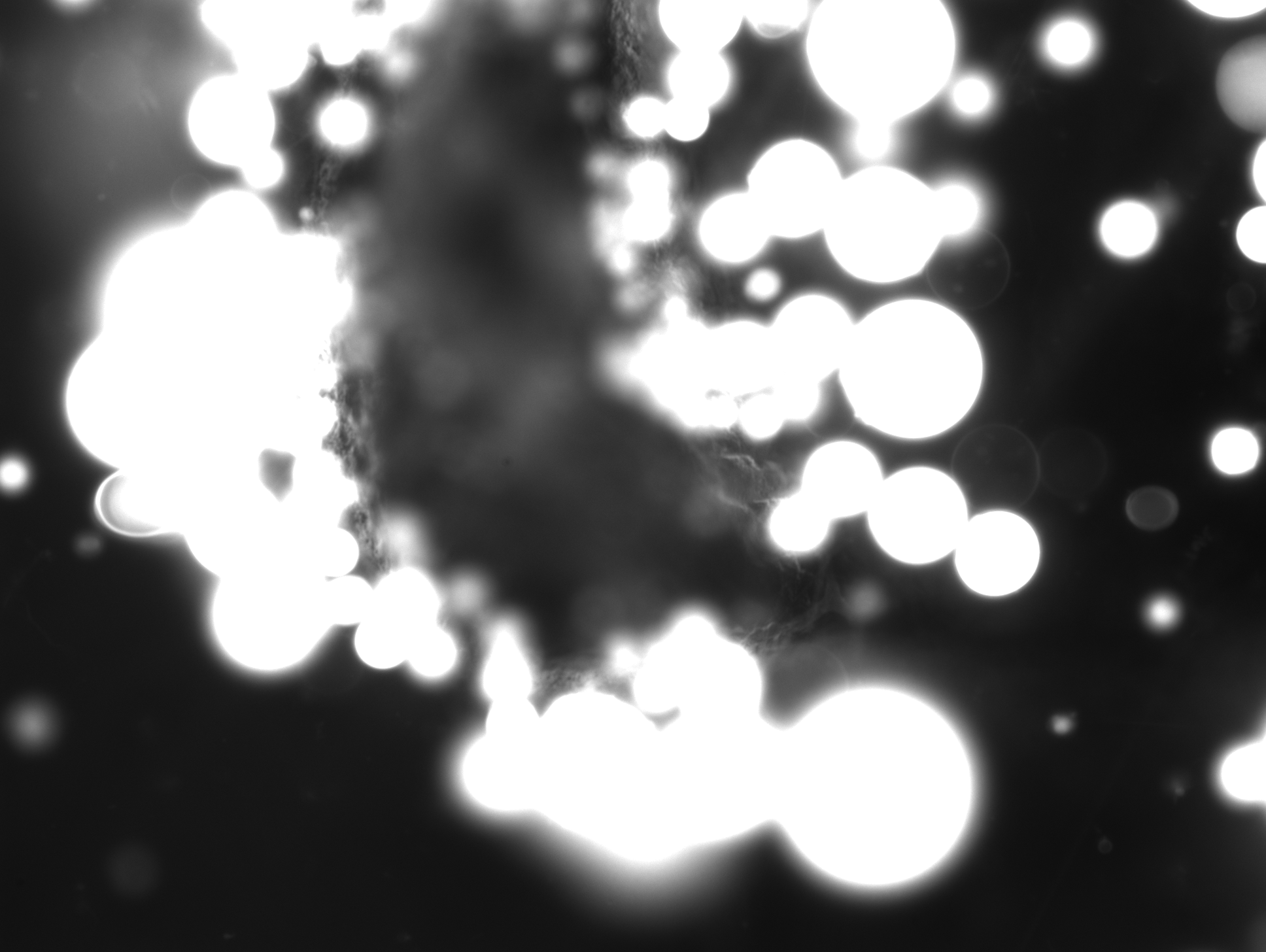

### 1_257_15.419.tif

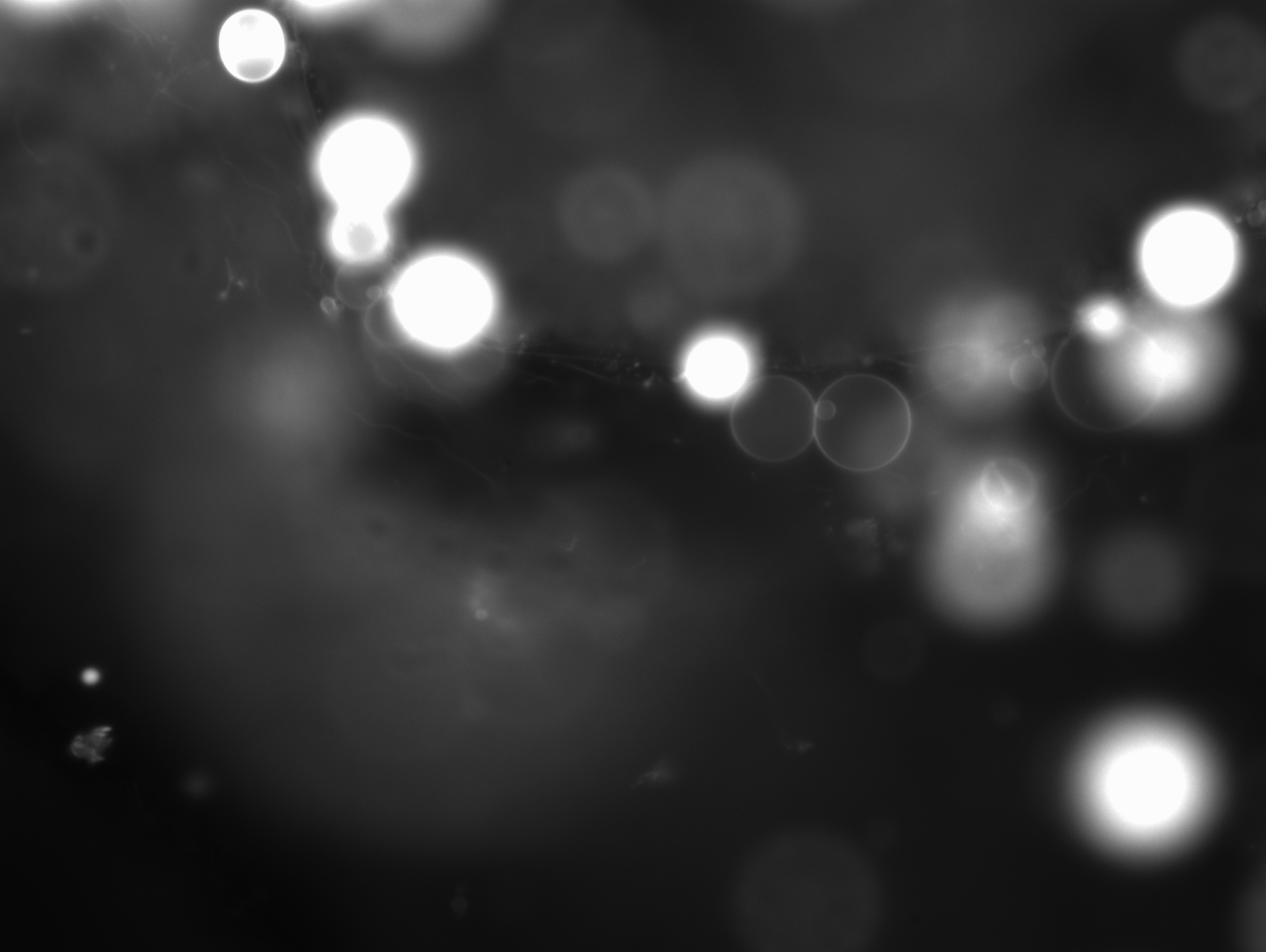

### 1_319_22.891.tif

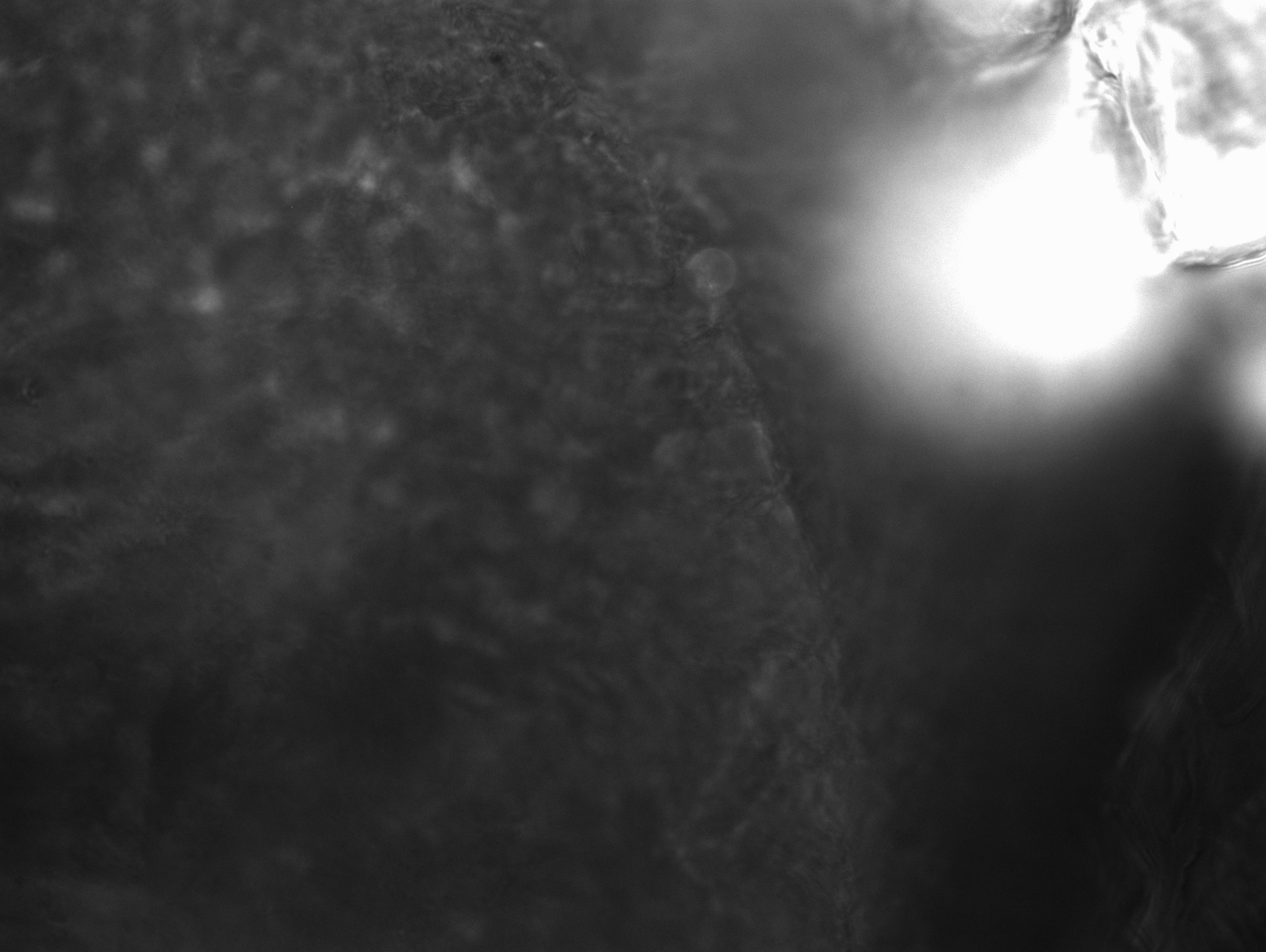

### 2_7_46.292.tif

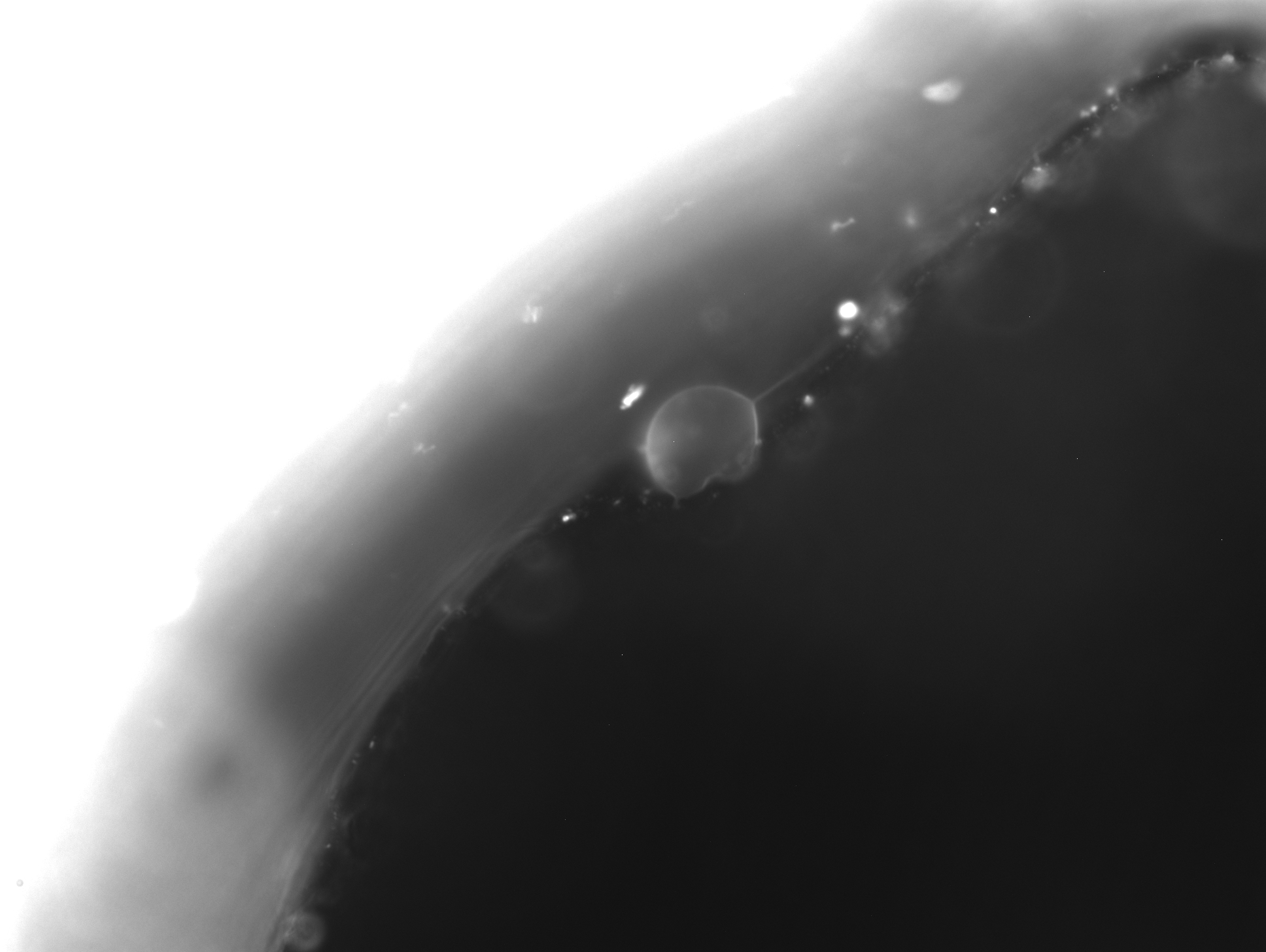

### 2_17.188.tif

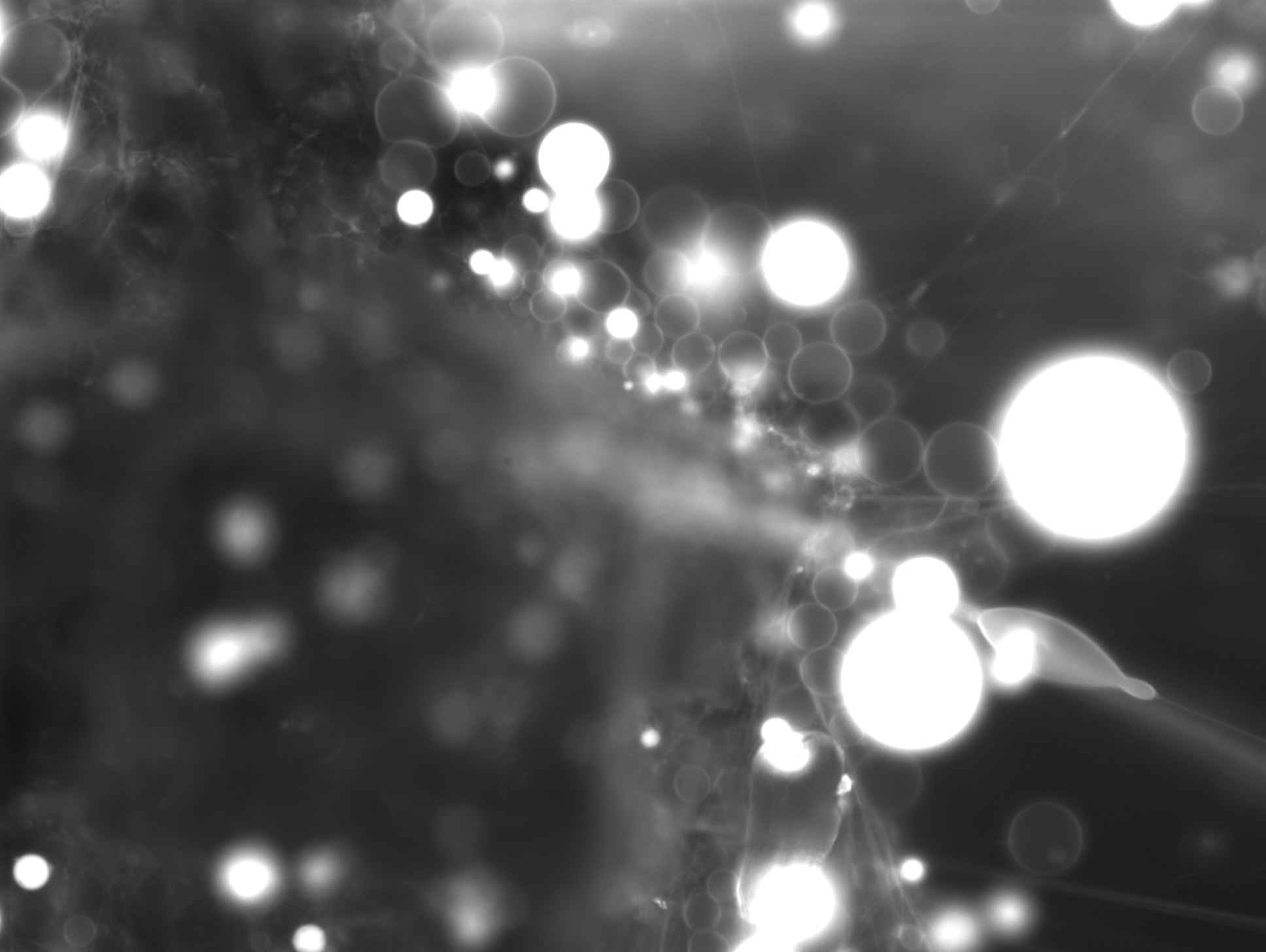

### 2_29.742-1.tif

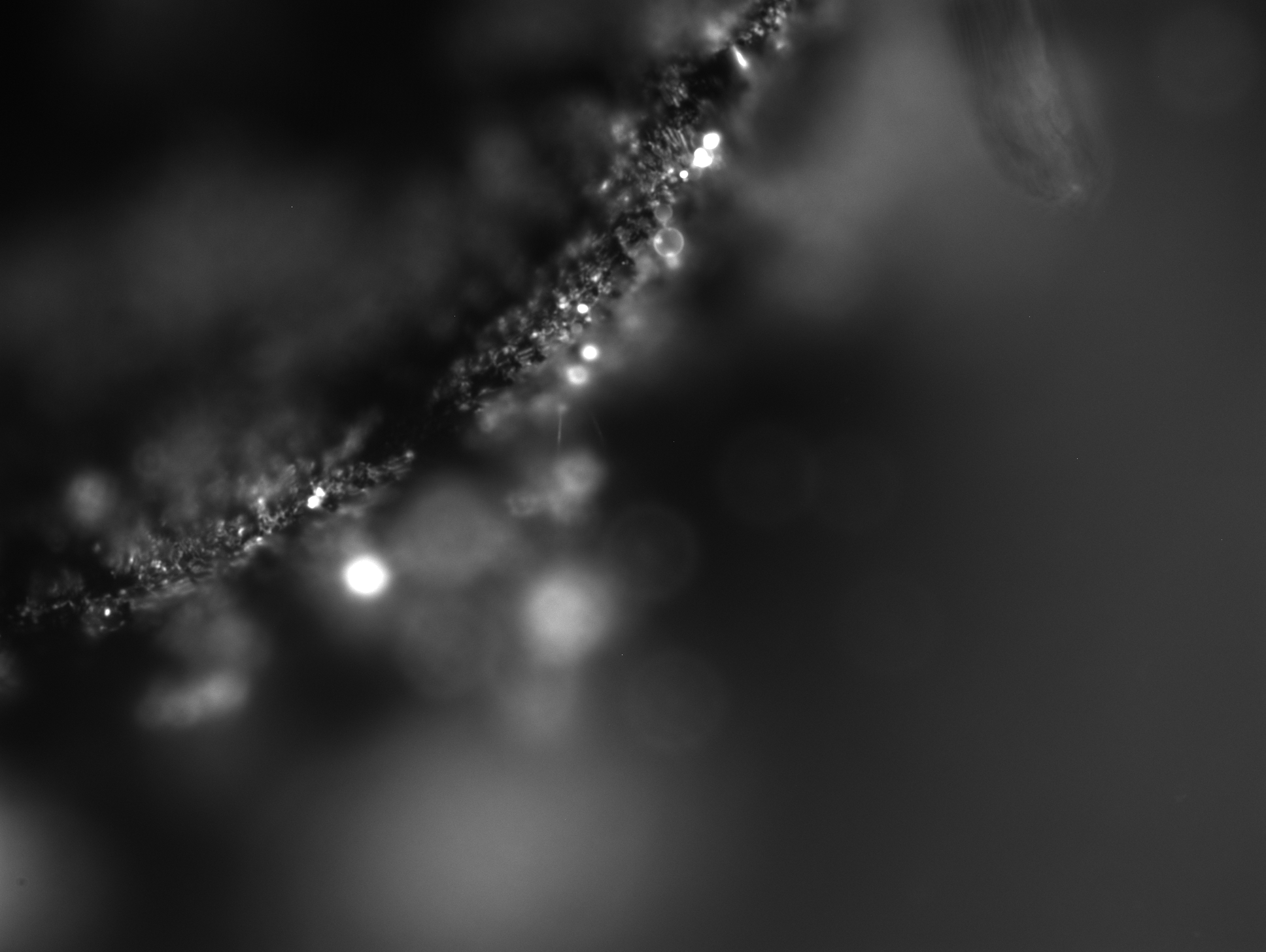

### 2_31_26.363.tif

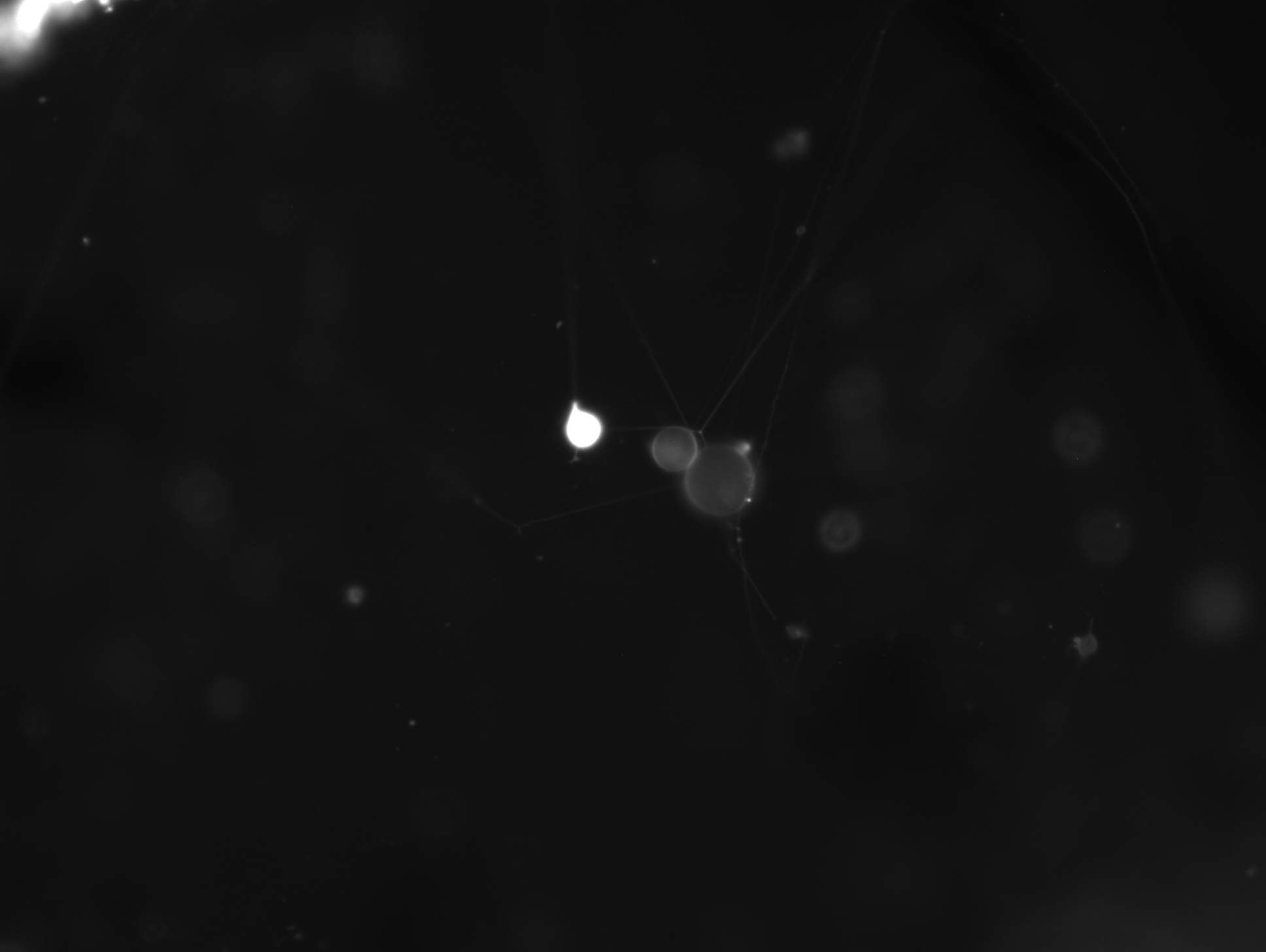

### 2_36_27.123.tif

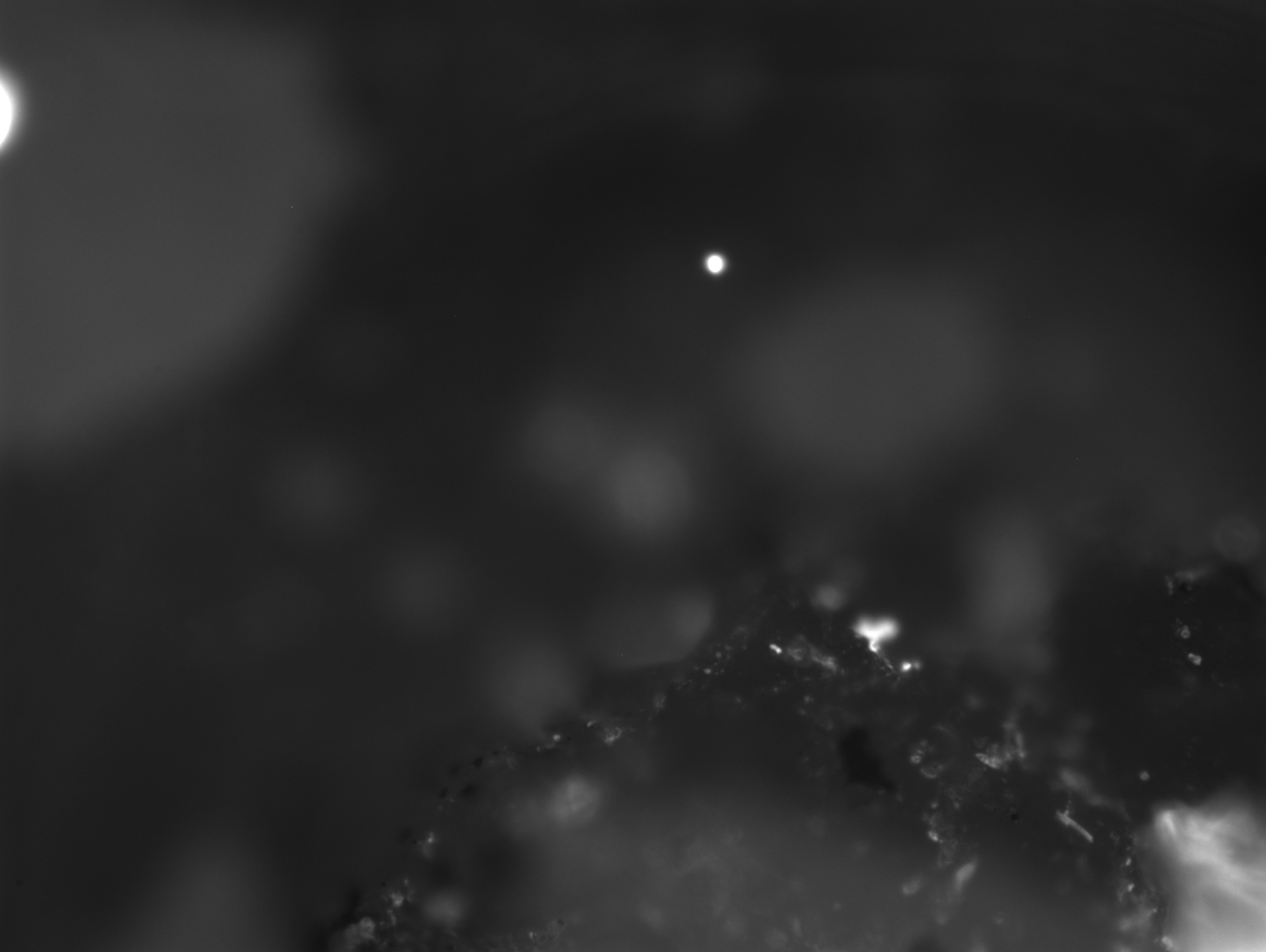

### 2_43_09.268.tif

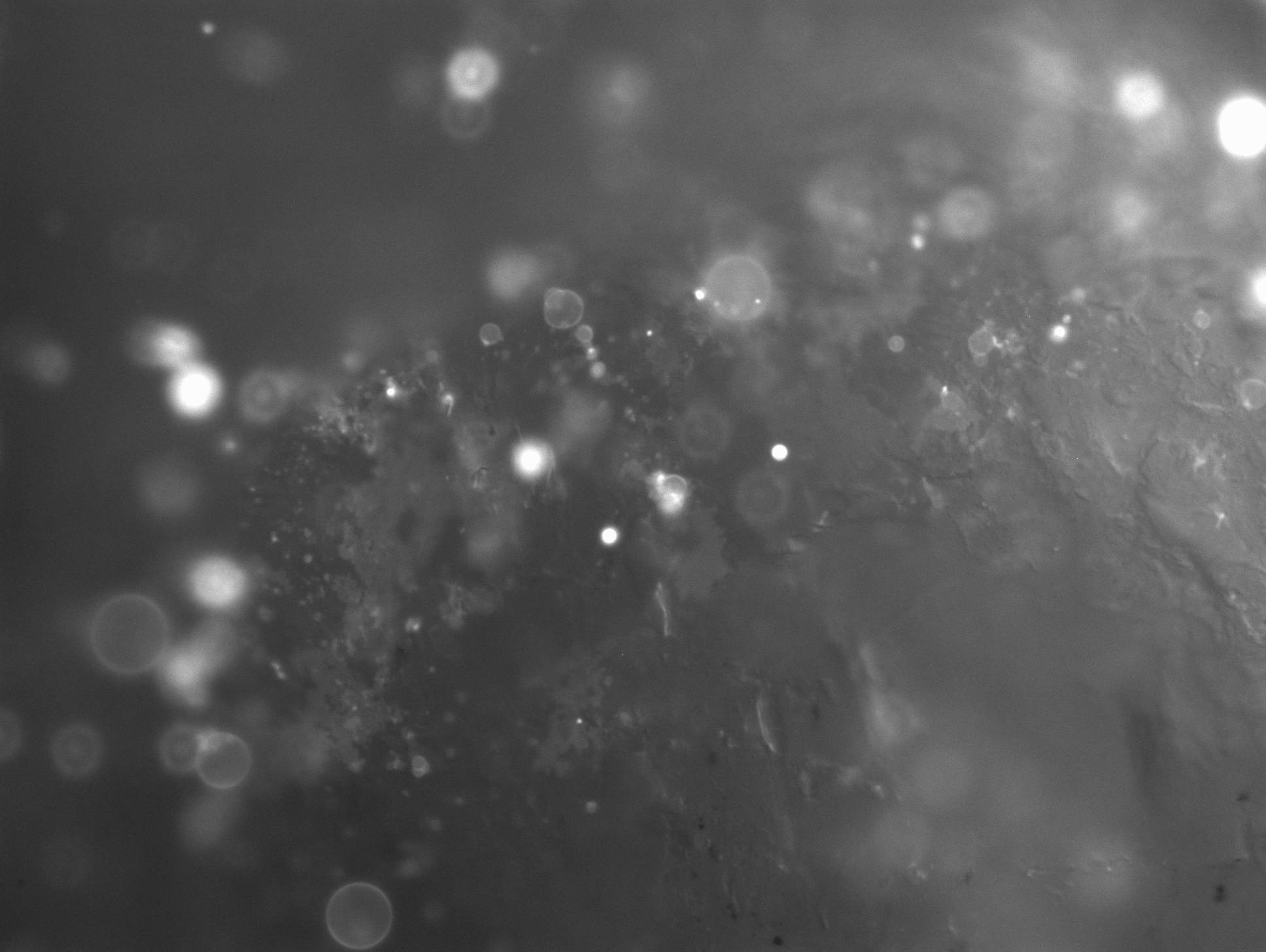

### 2_50_42.881.tif

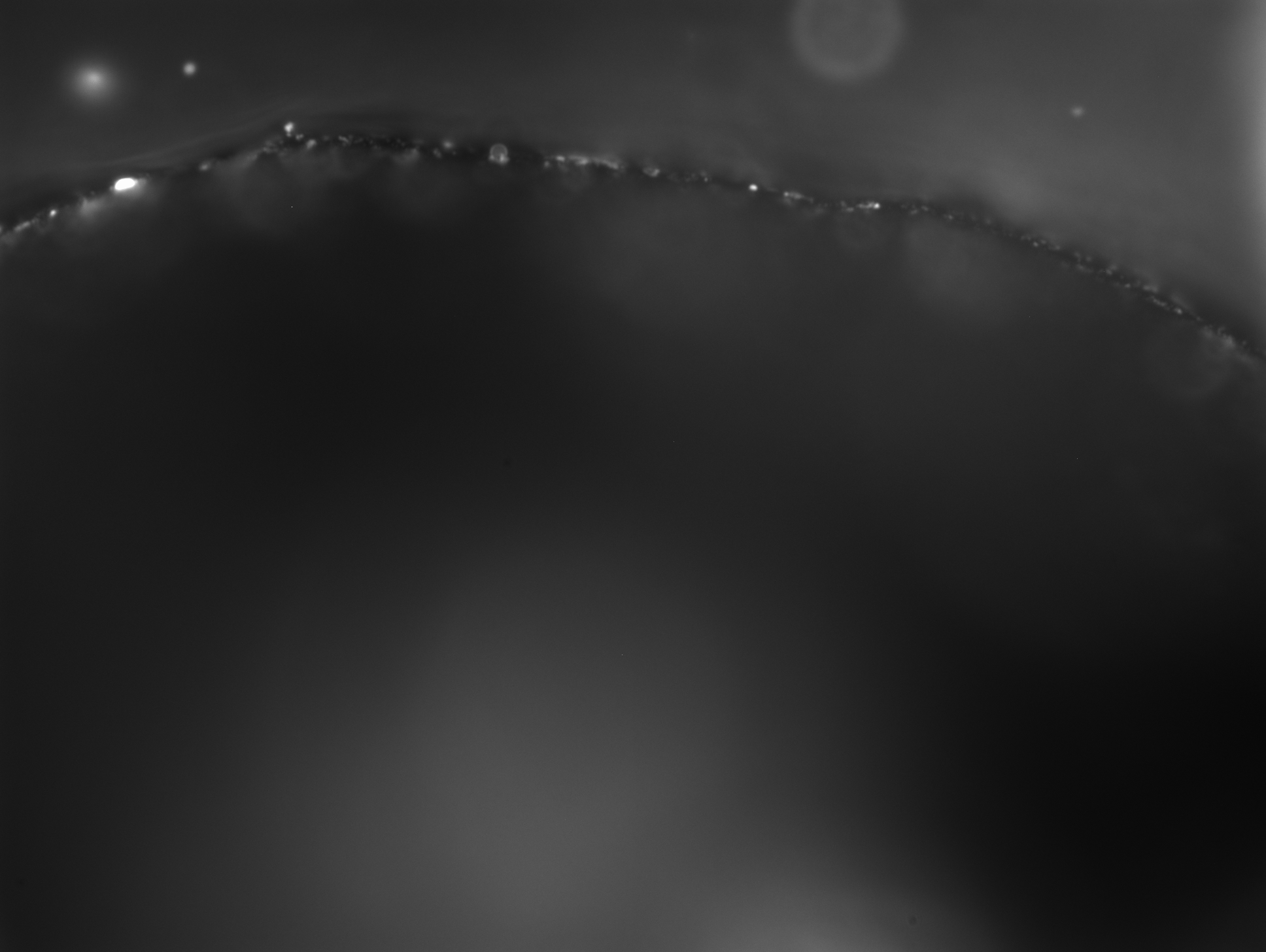

### 2_52.230-1.tif

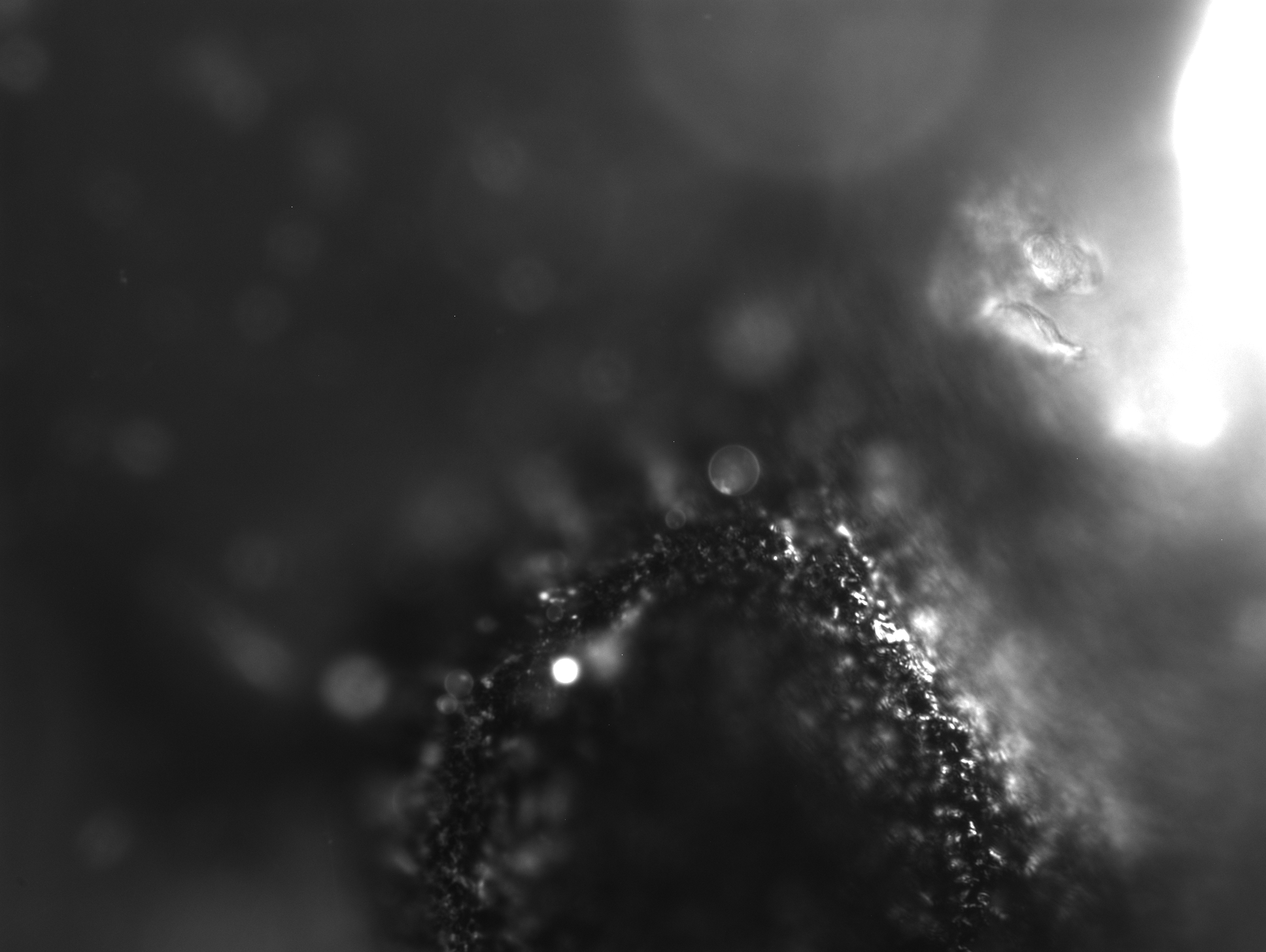

### 2_66_06.370.tif

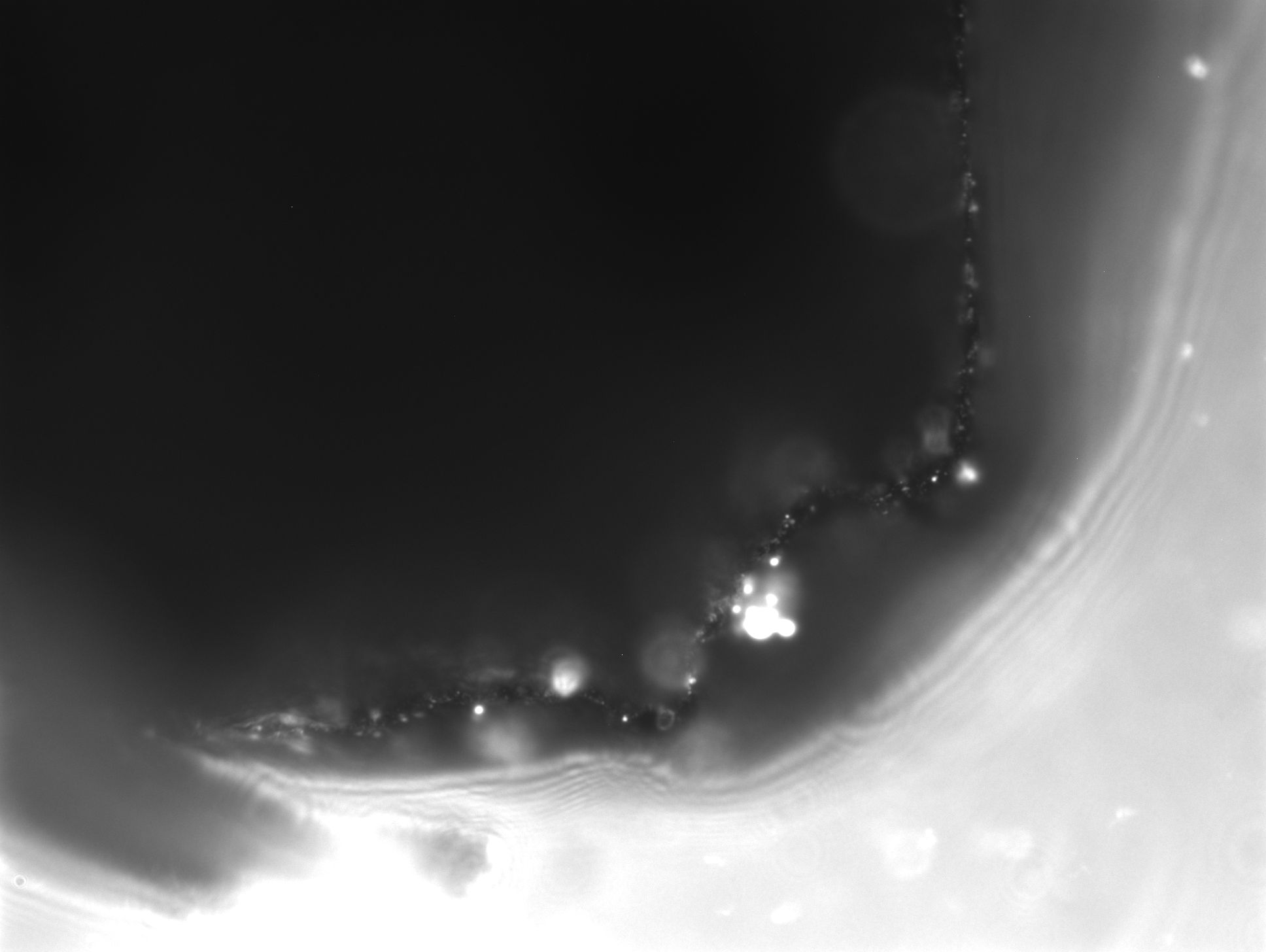

### 2_72_06.402.tif

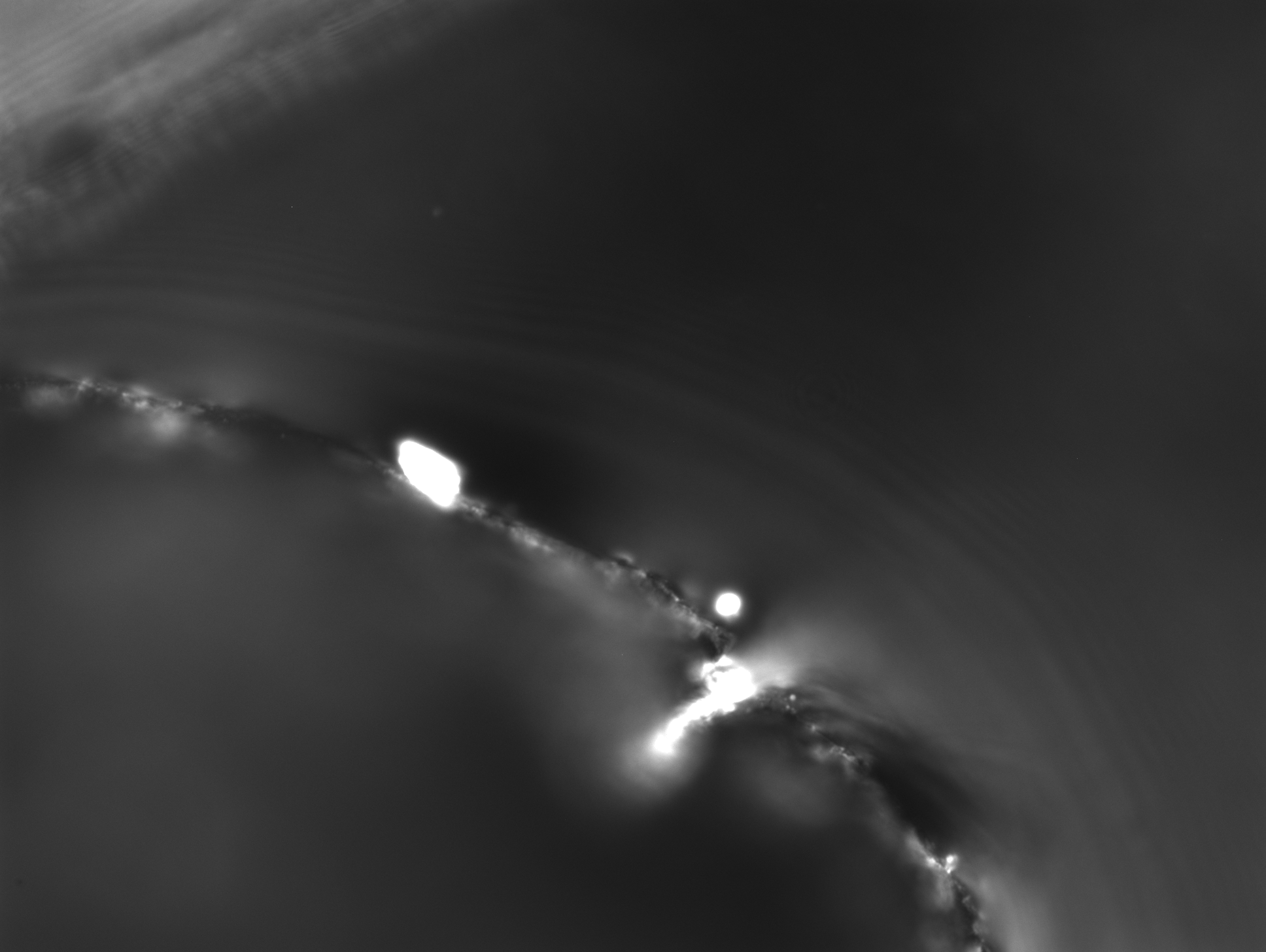

### 2_74_54.700.tif

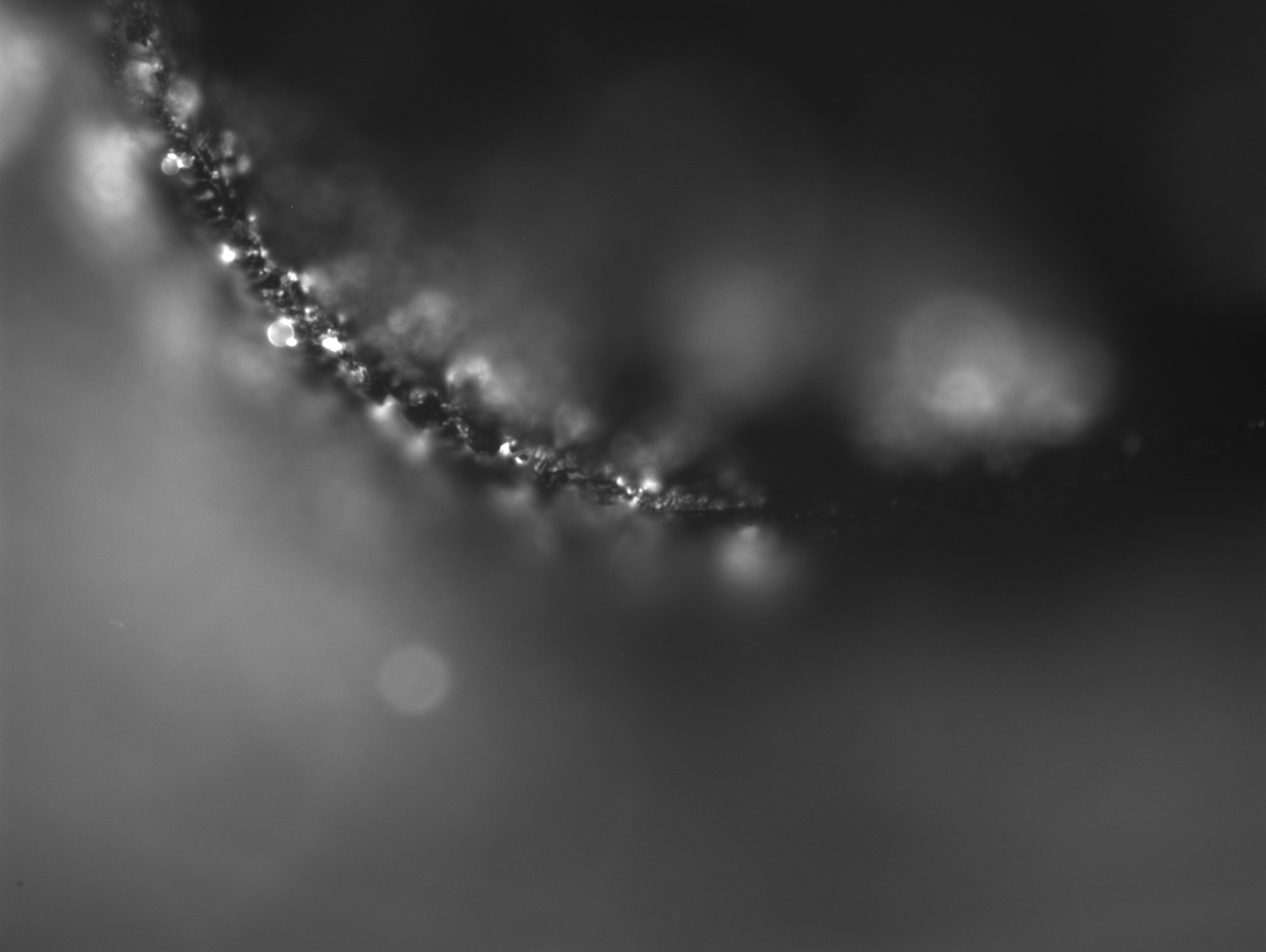

### 2_76_49.715.tif

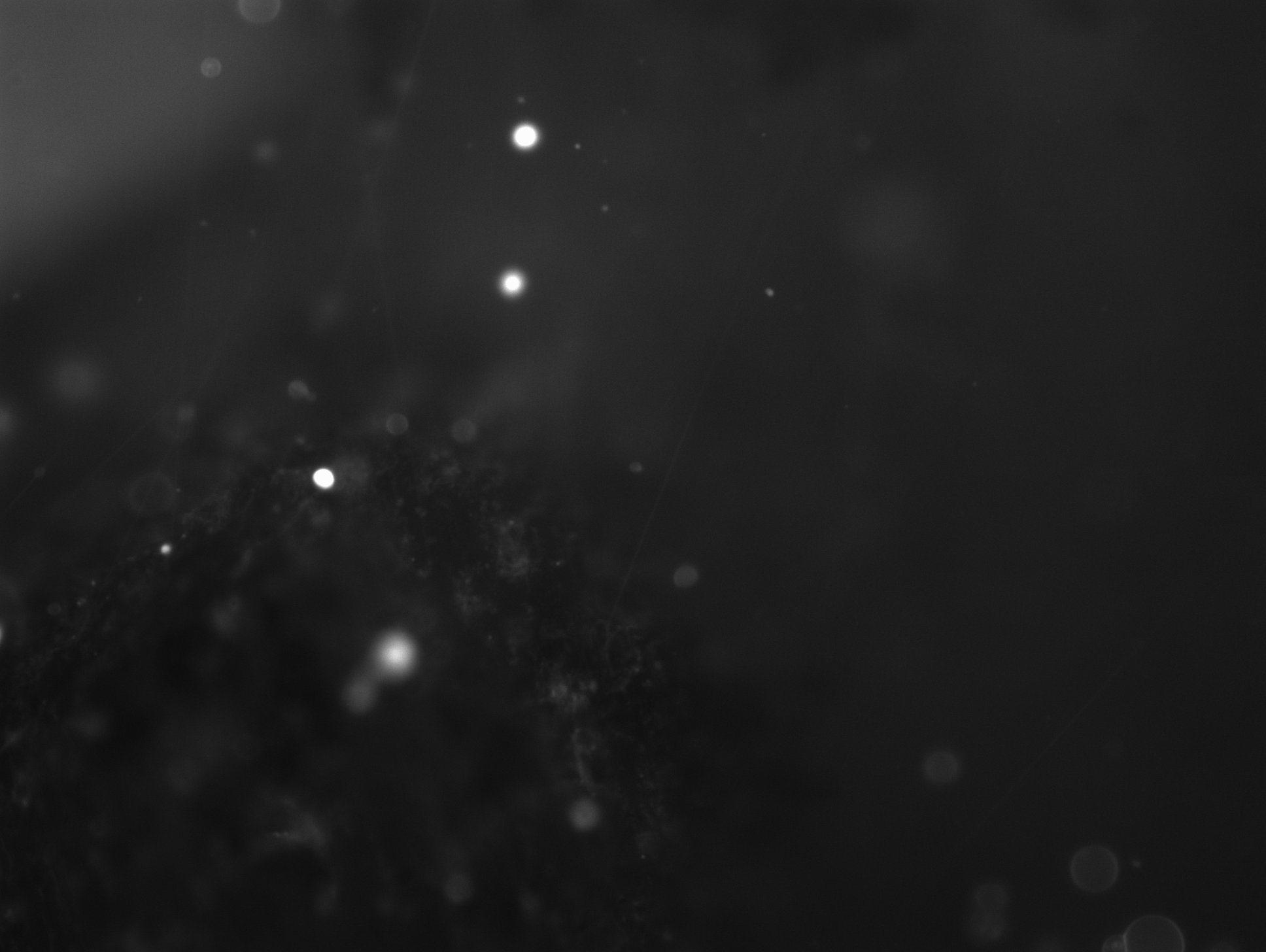

### 2_81_48.363.tif

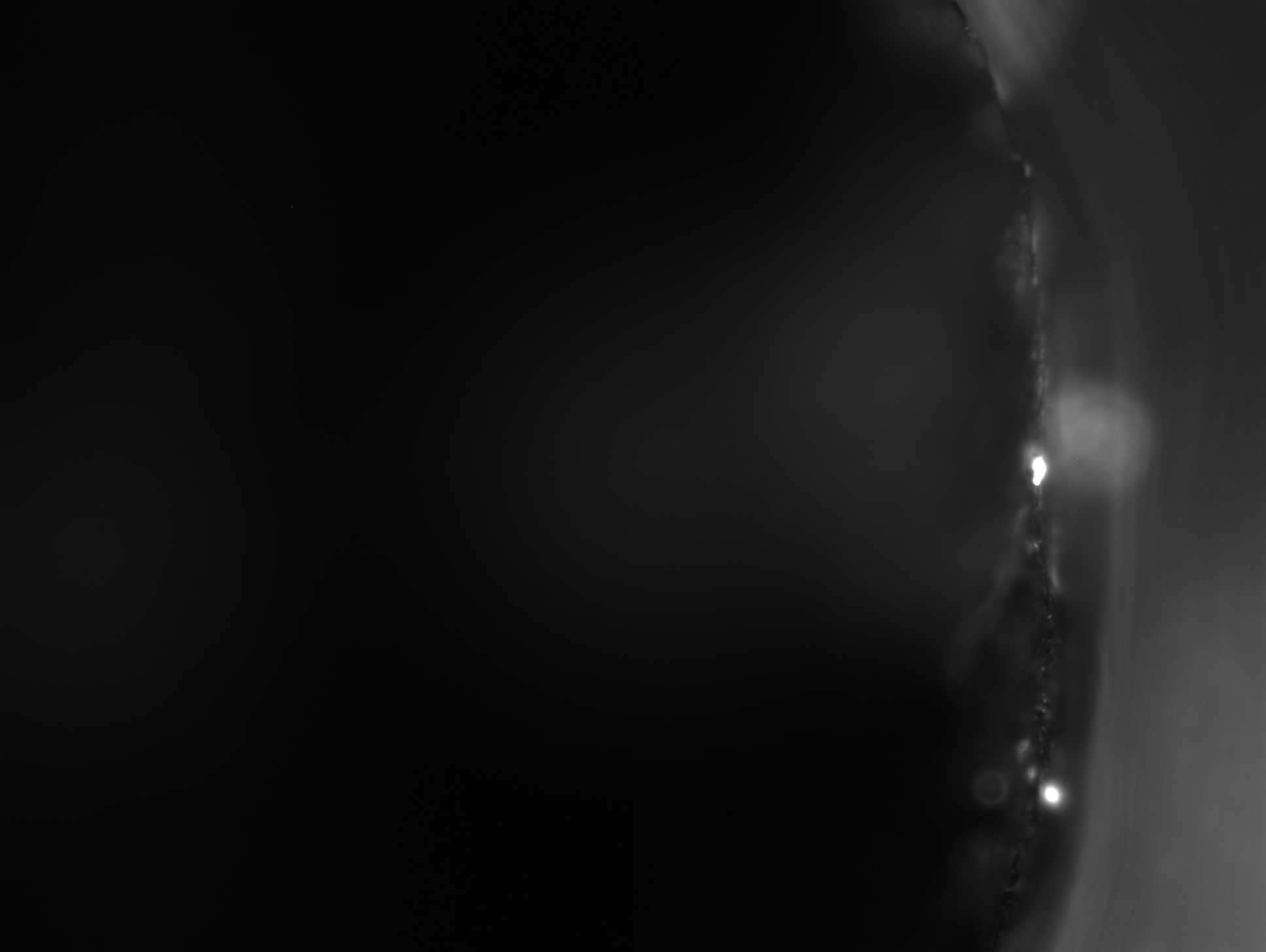

### 2_94_17.769.tif

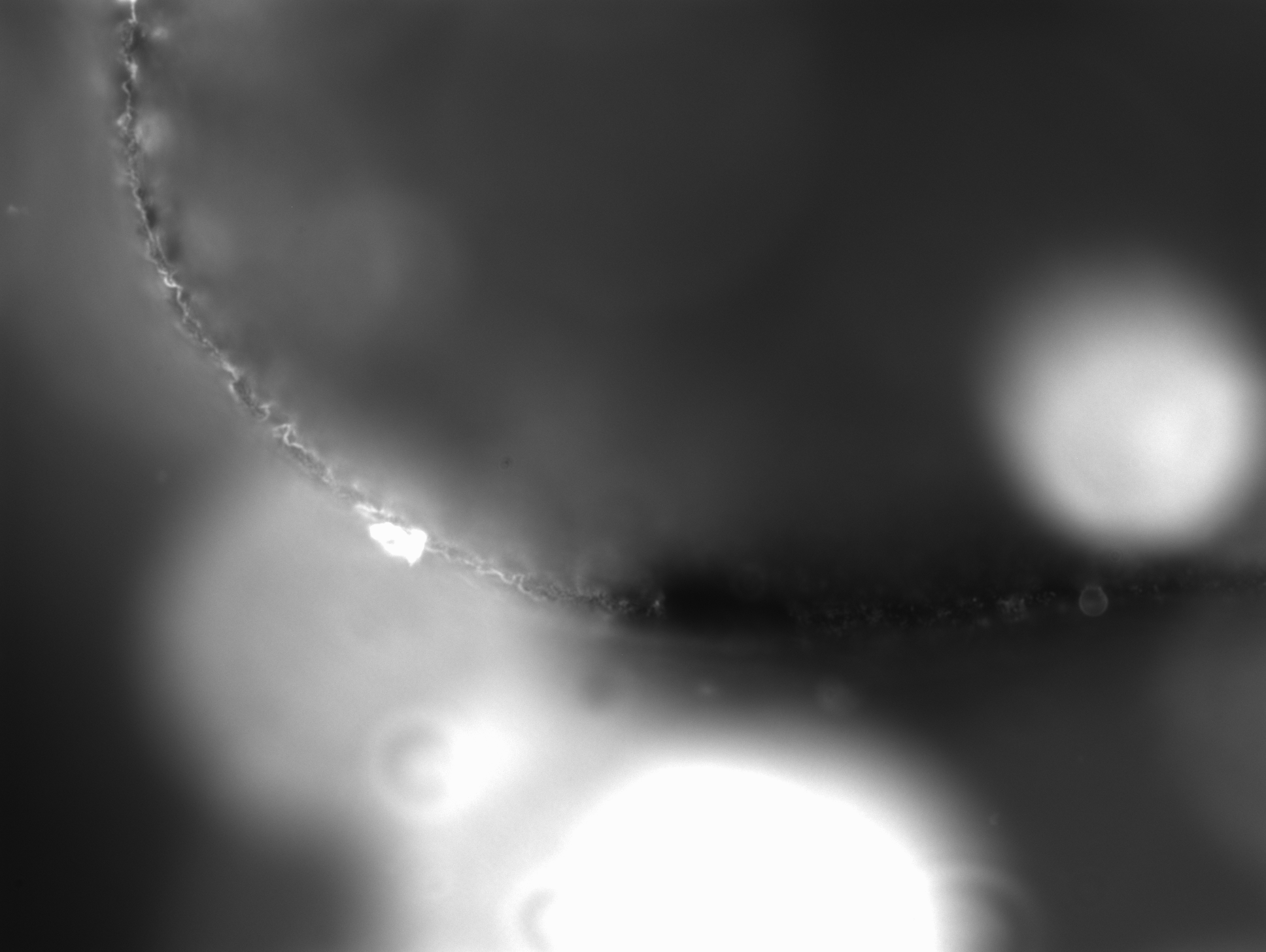

### 2_100_00.009.tif

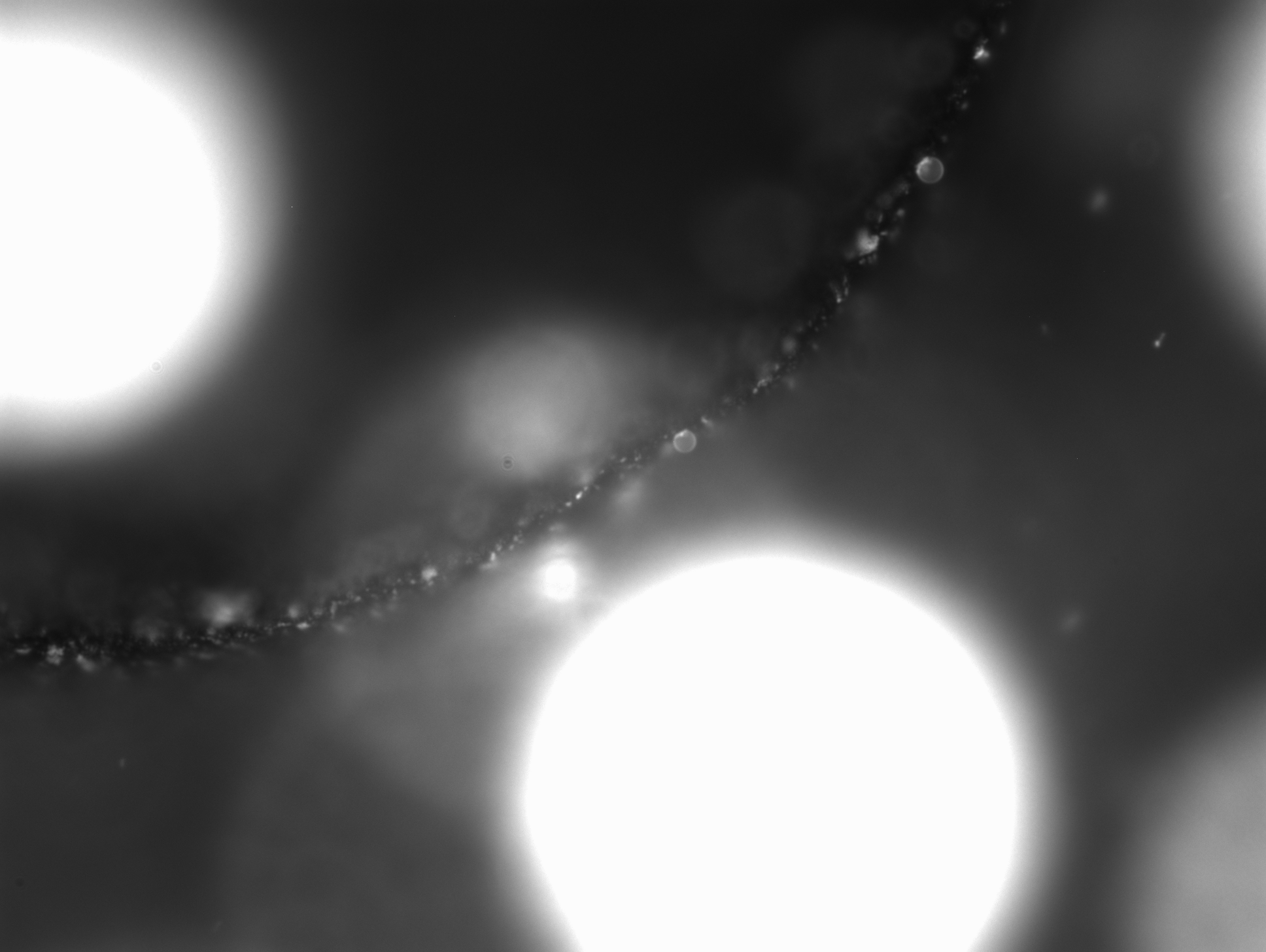

### 2_111_15.419.tif

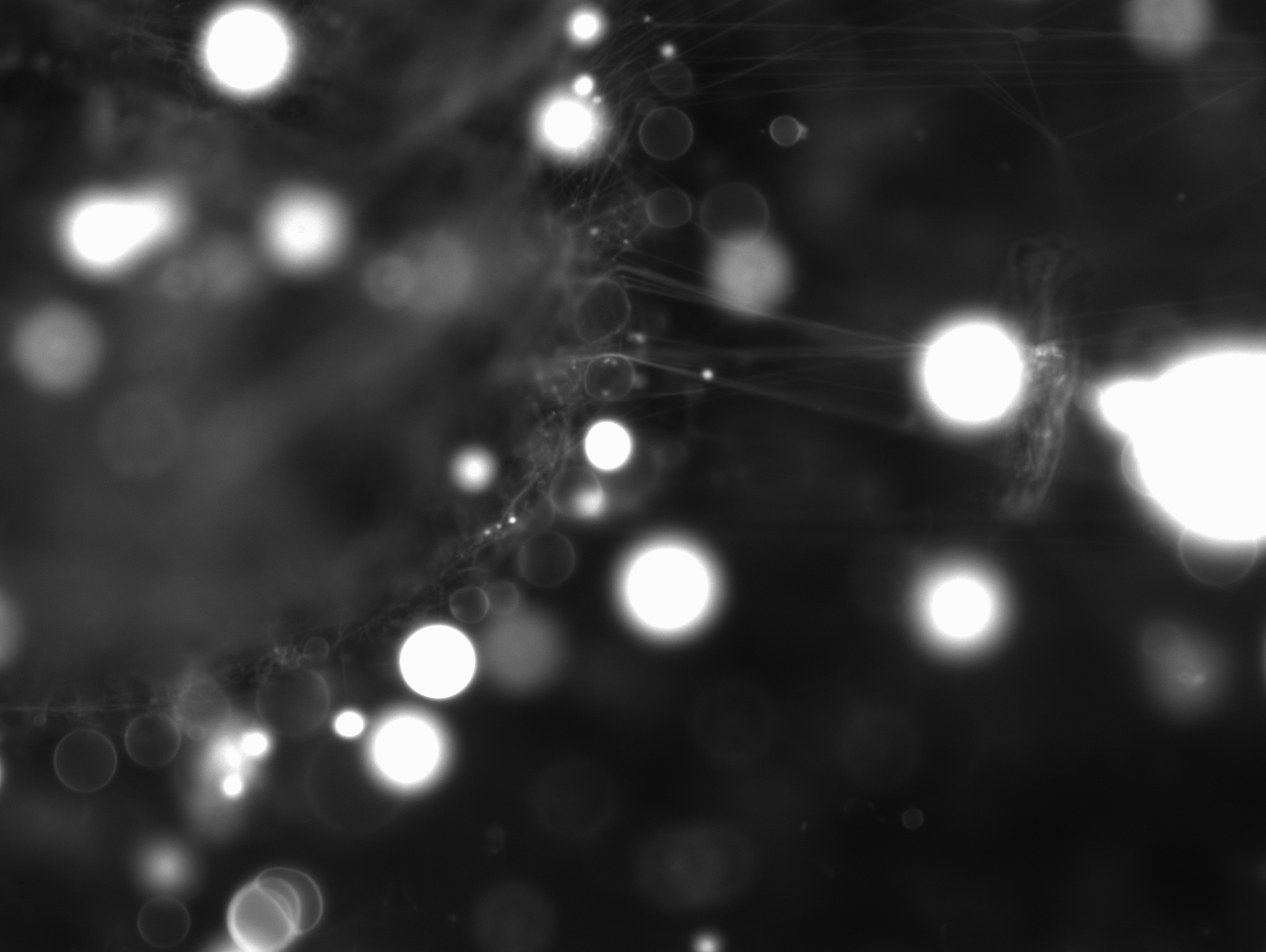

### 2_117_25.868.tif

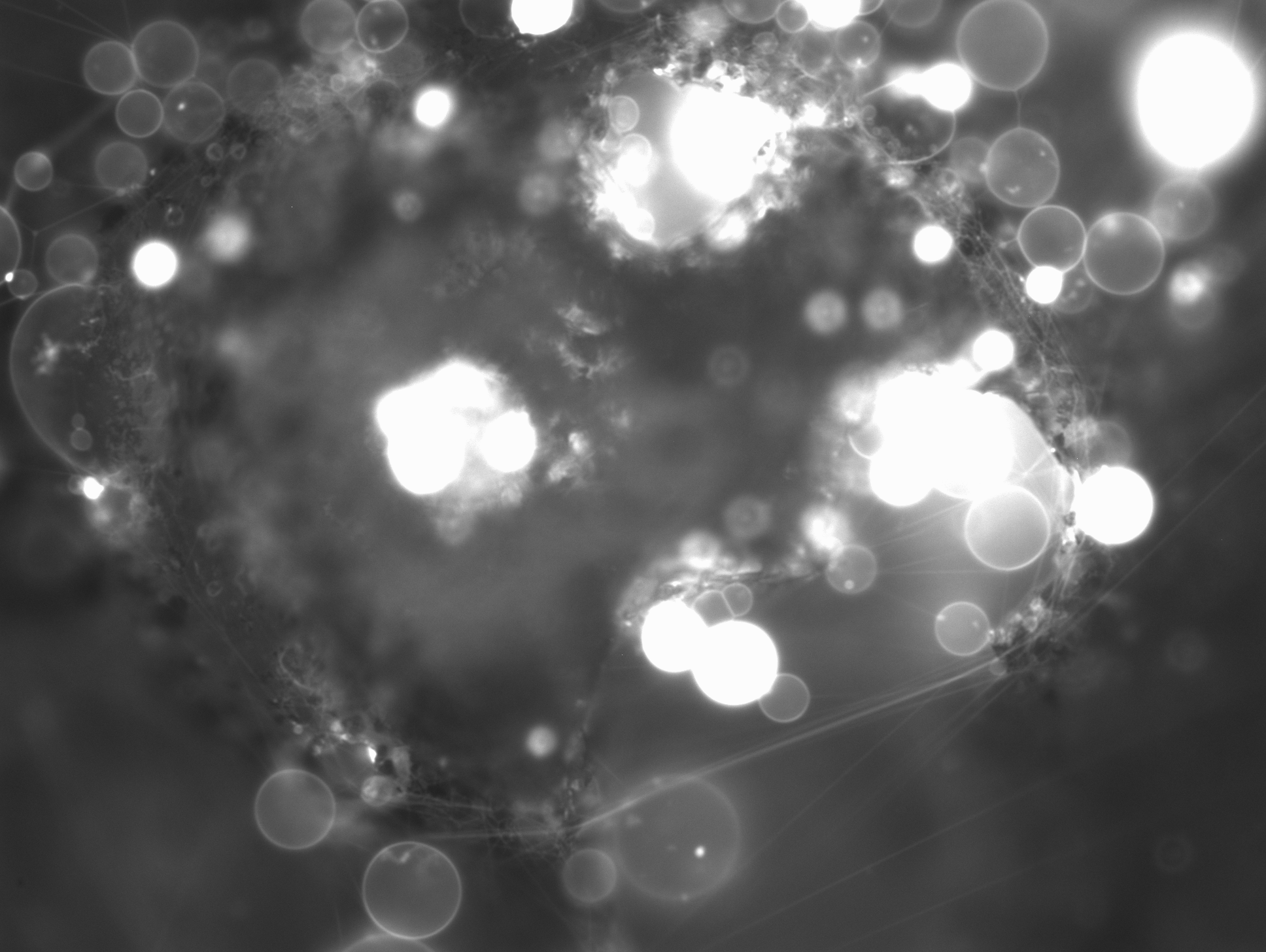

### 2_120_42.138.tif

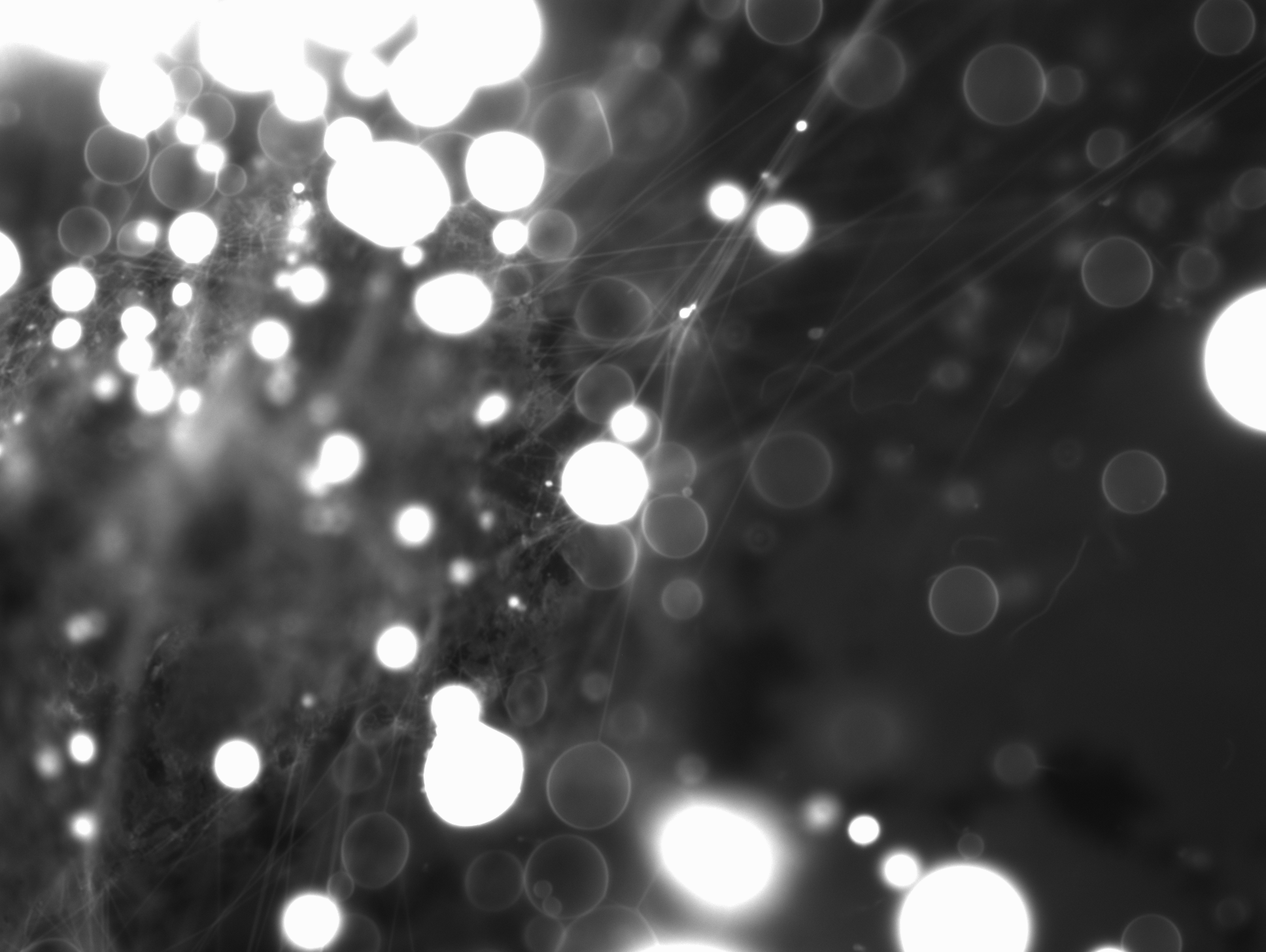

### 2_134_48.976.tif

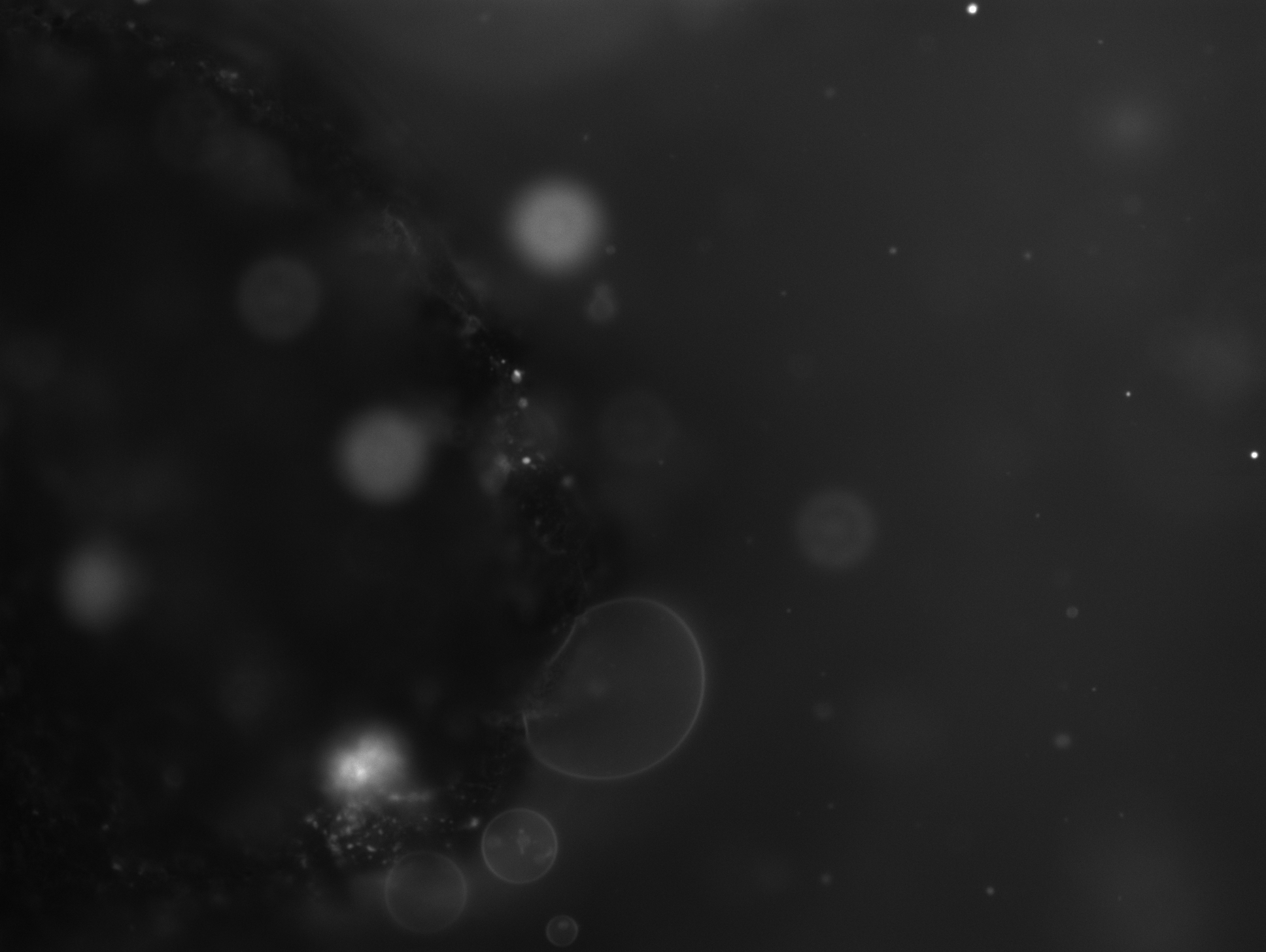

### 2_169_09.552.tif

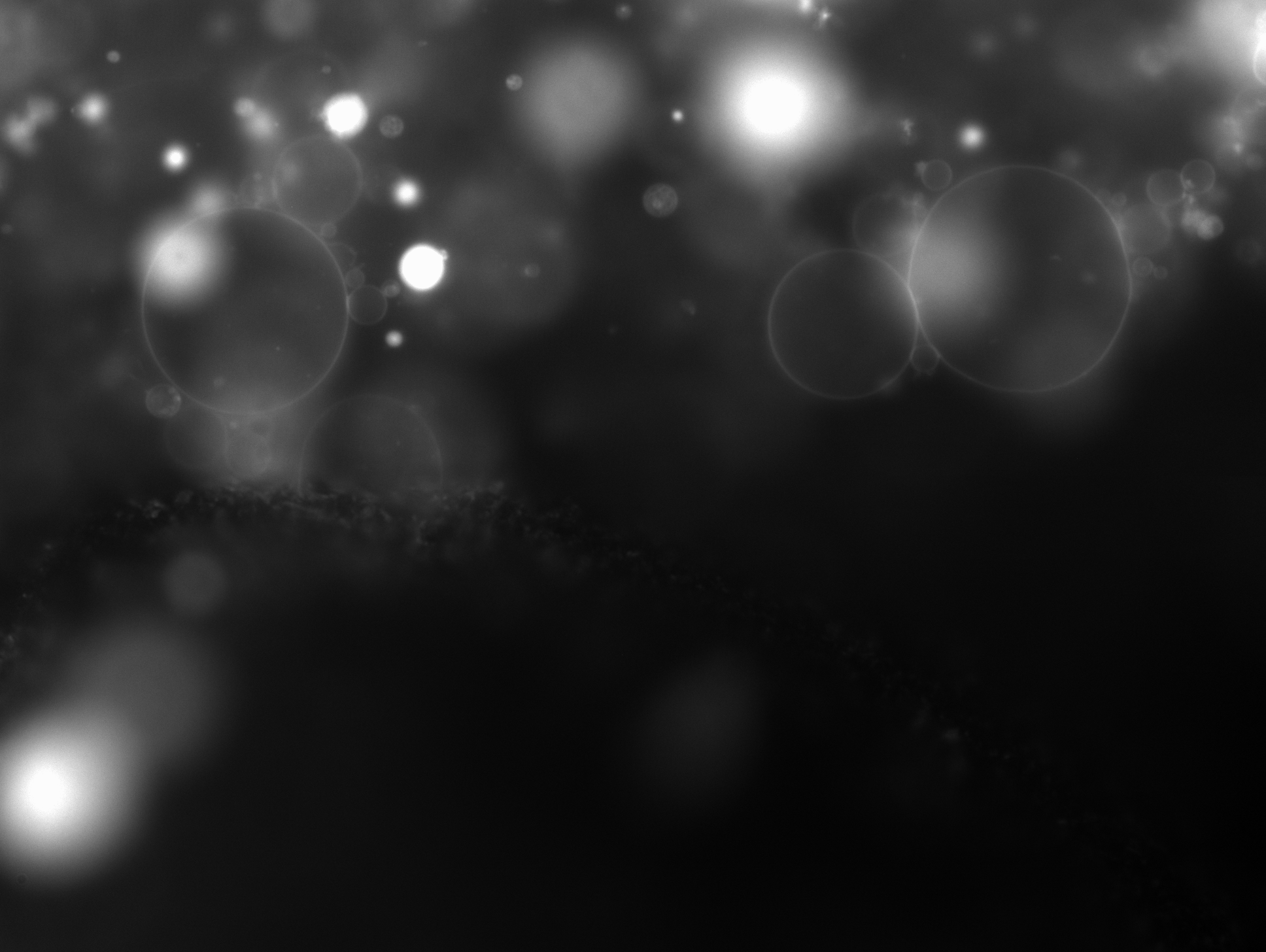

### 2_195_03.795.tif

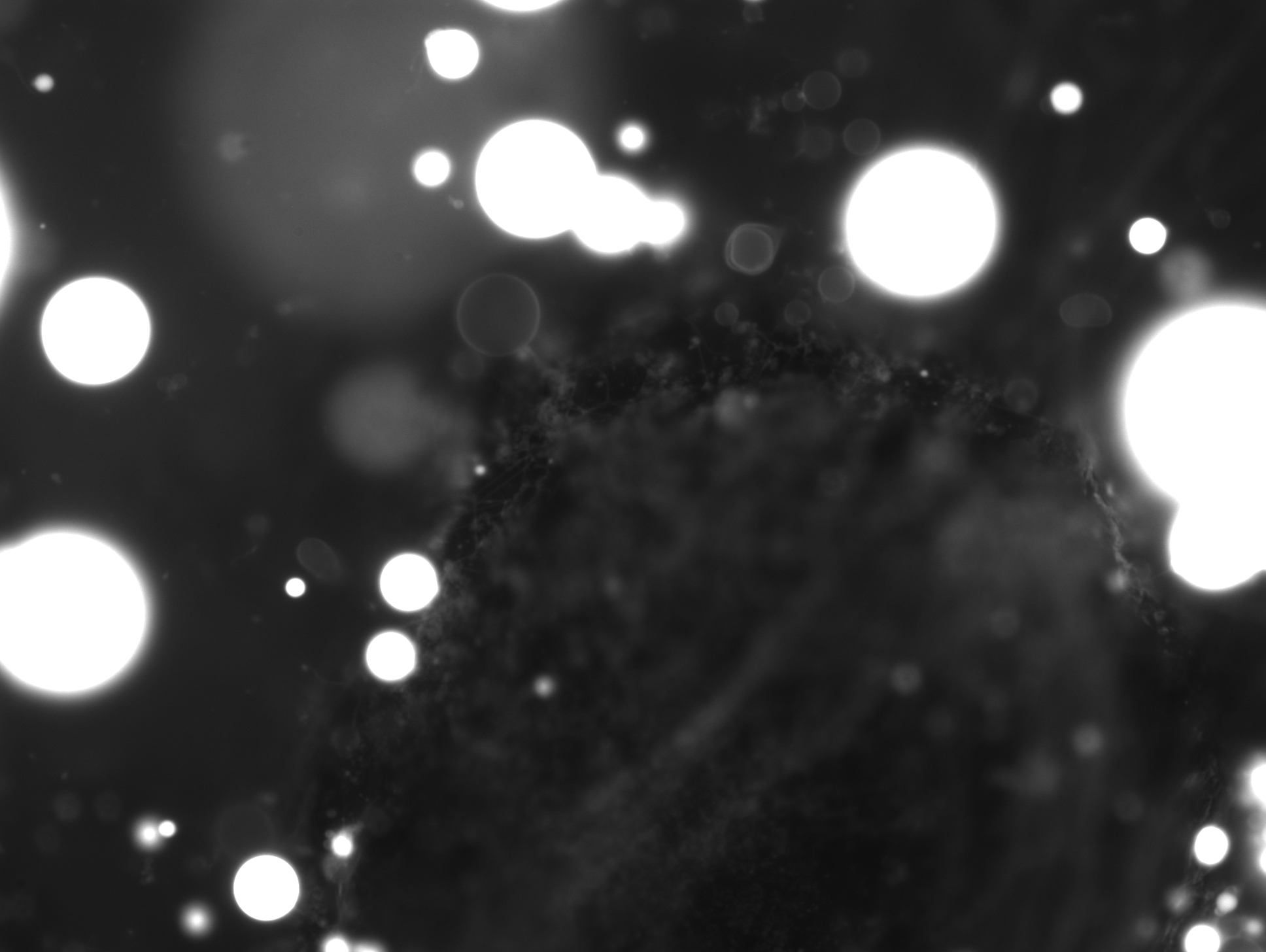

### 2_228_02.132.tif

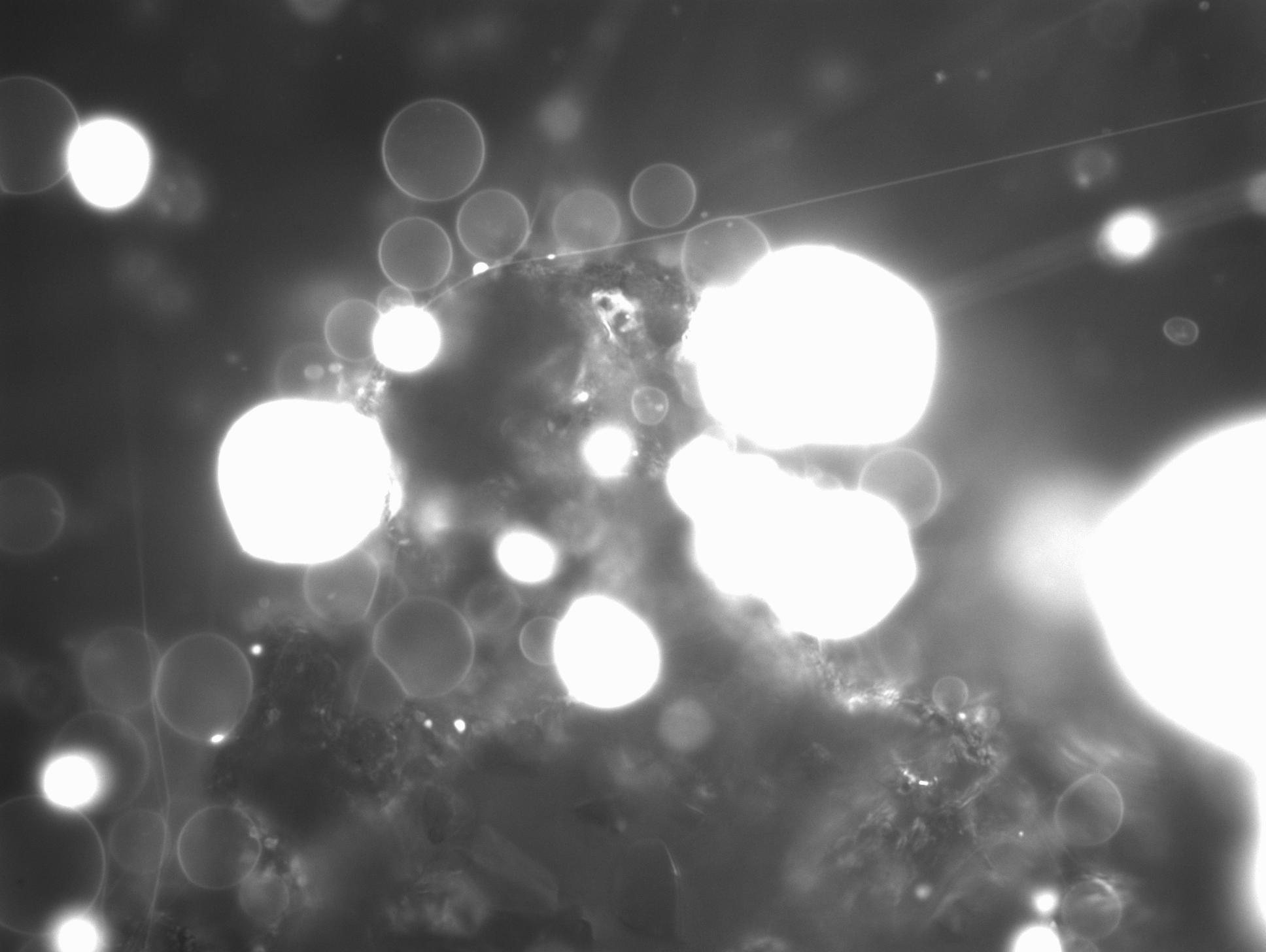

### 2_249_18.467.tif

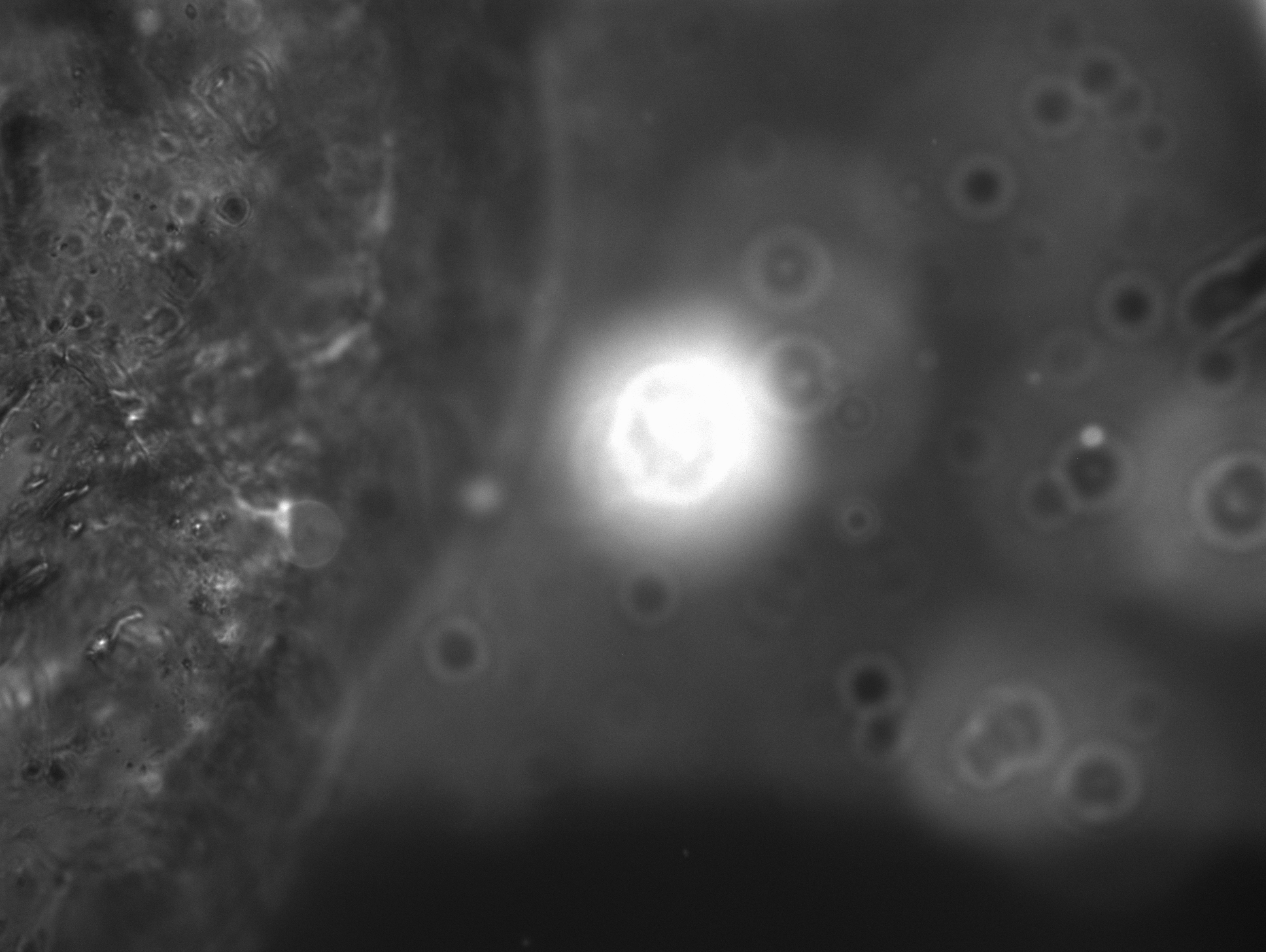

### 2_382_38.836.tif

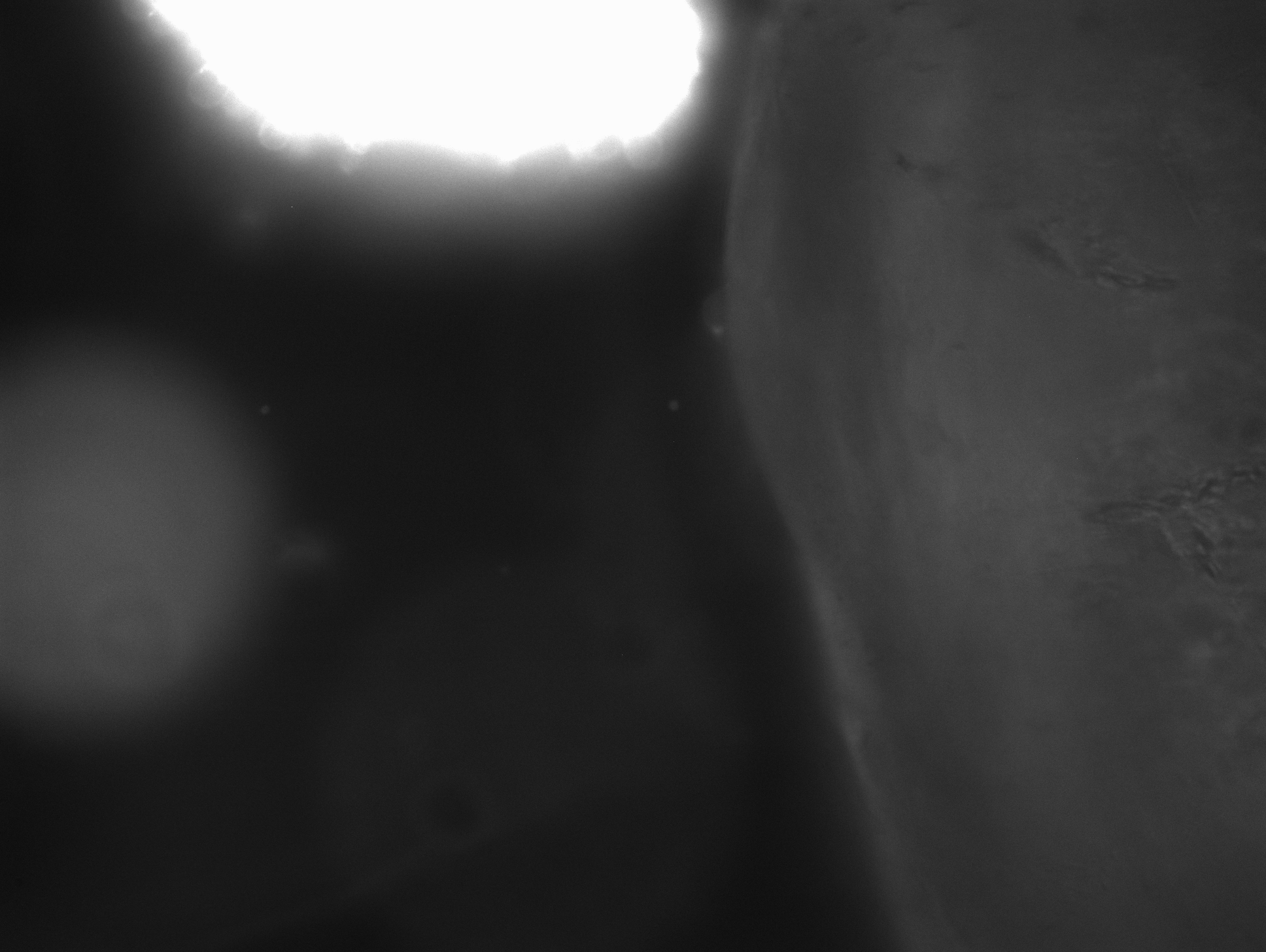

### 2_416_54.811.tif

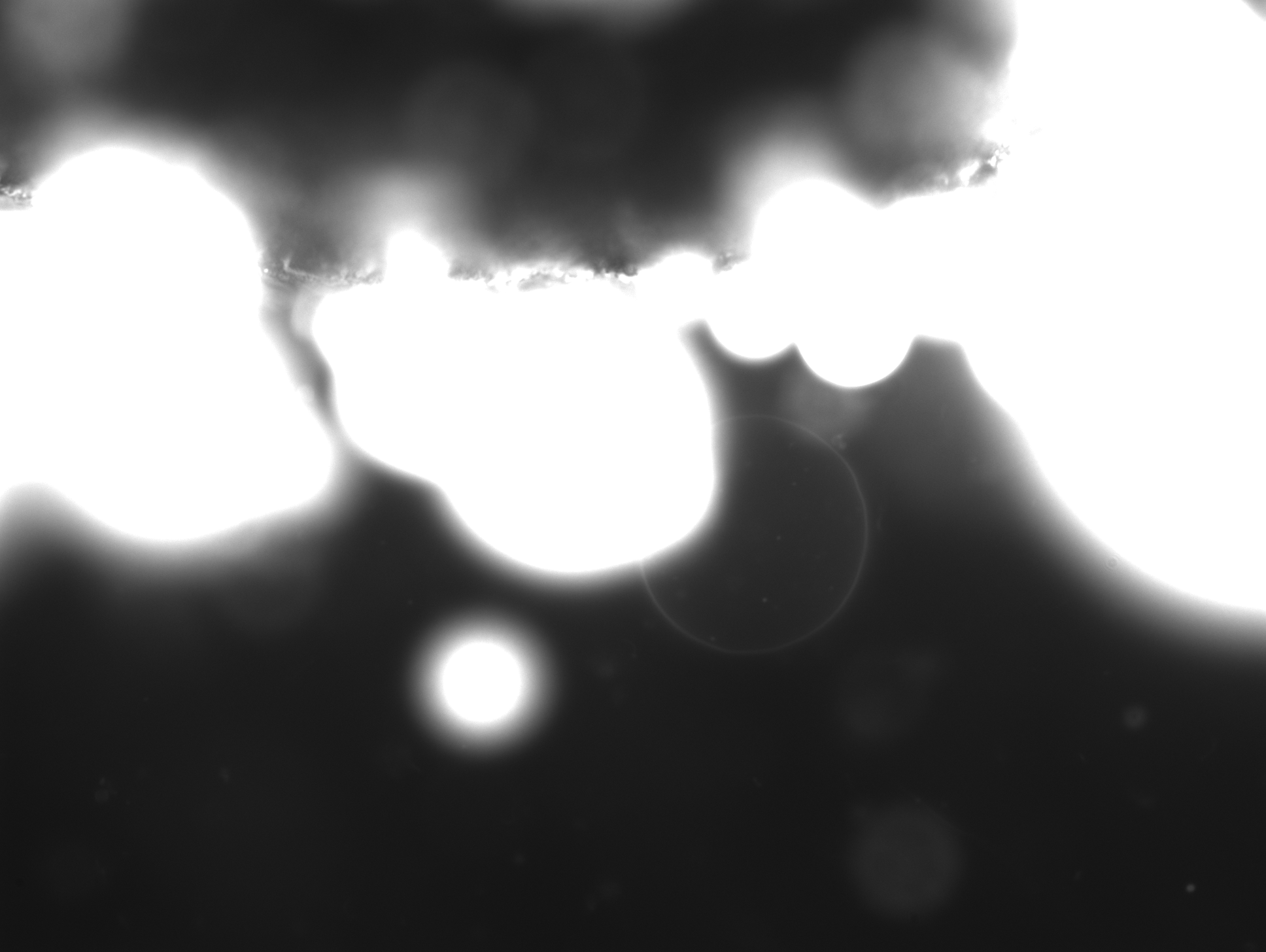

### 2_470_51.140.tif

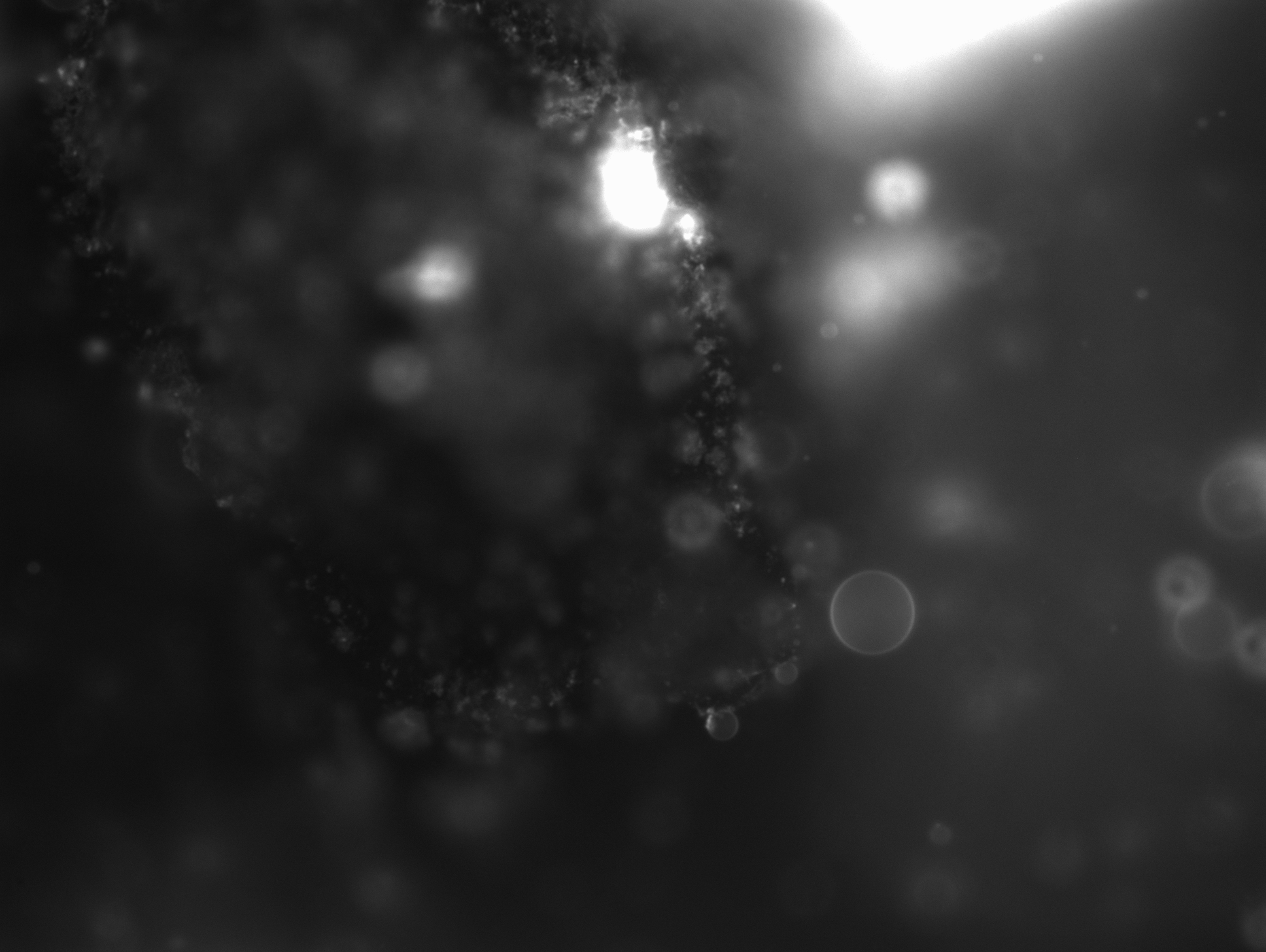

### 2_692_30.714.tif

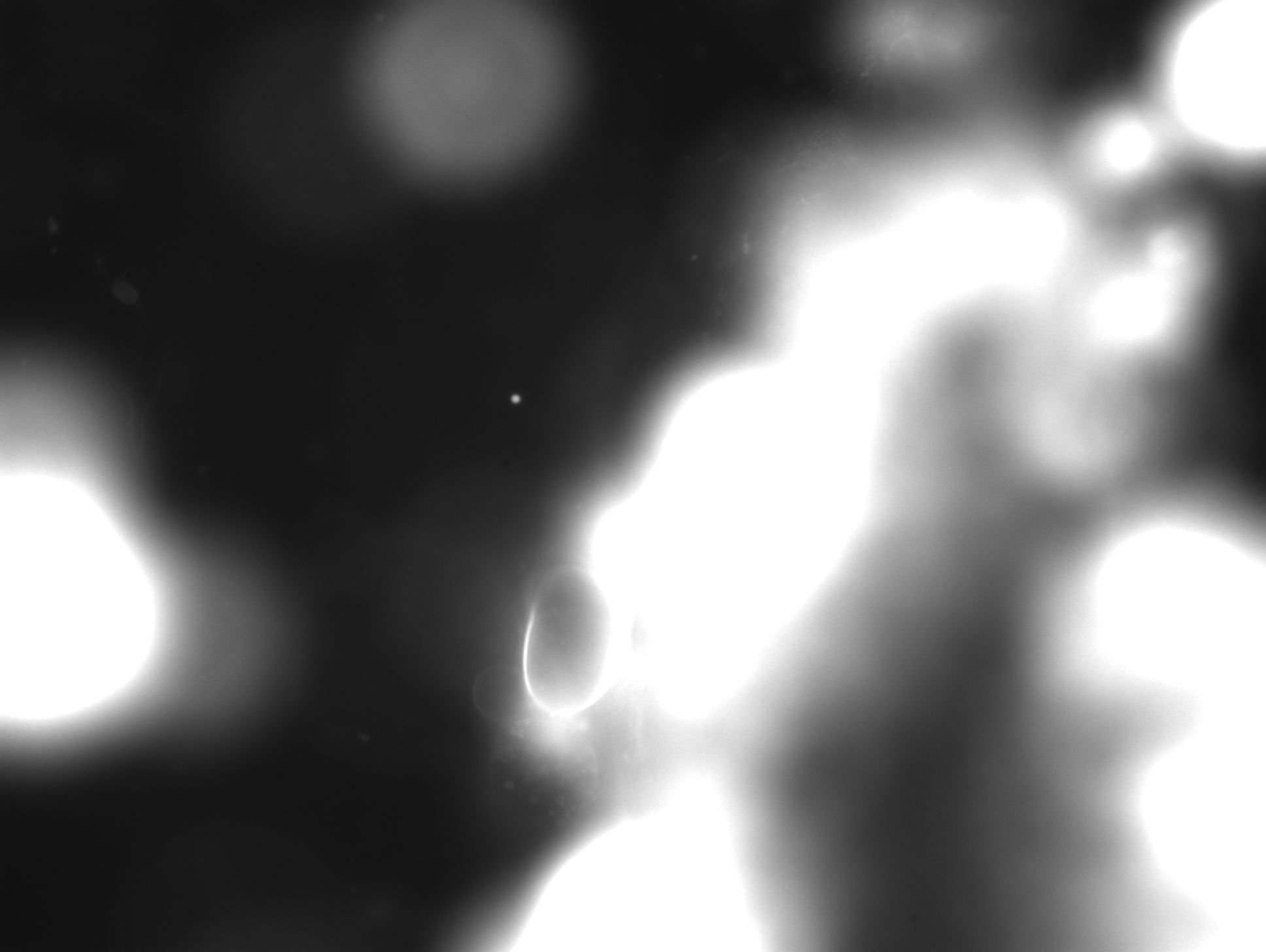

### 3_12.094.tif

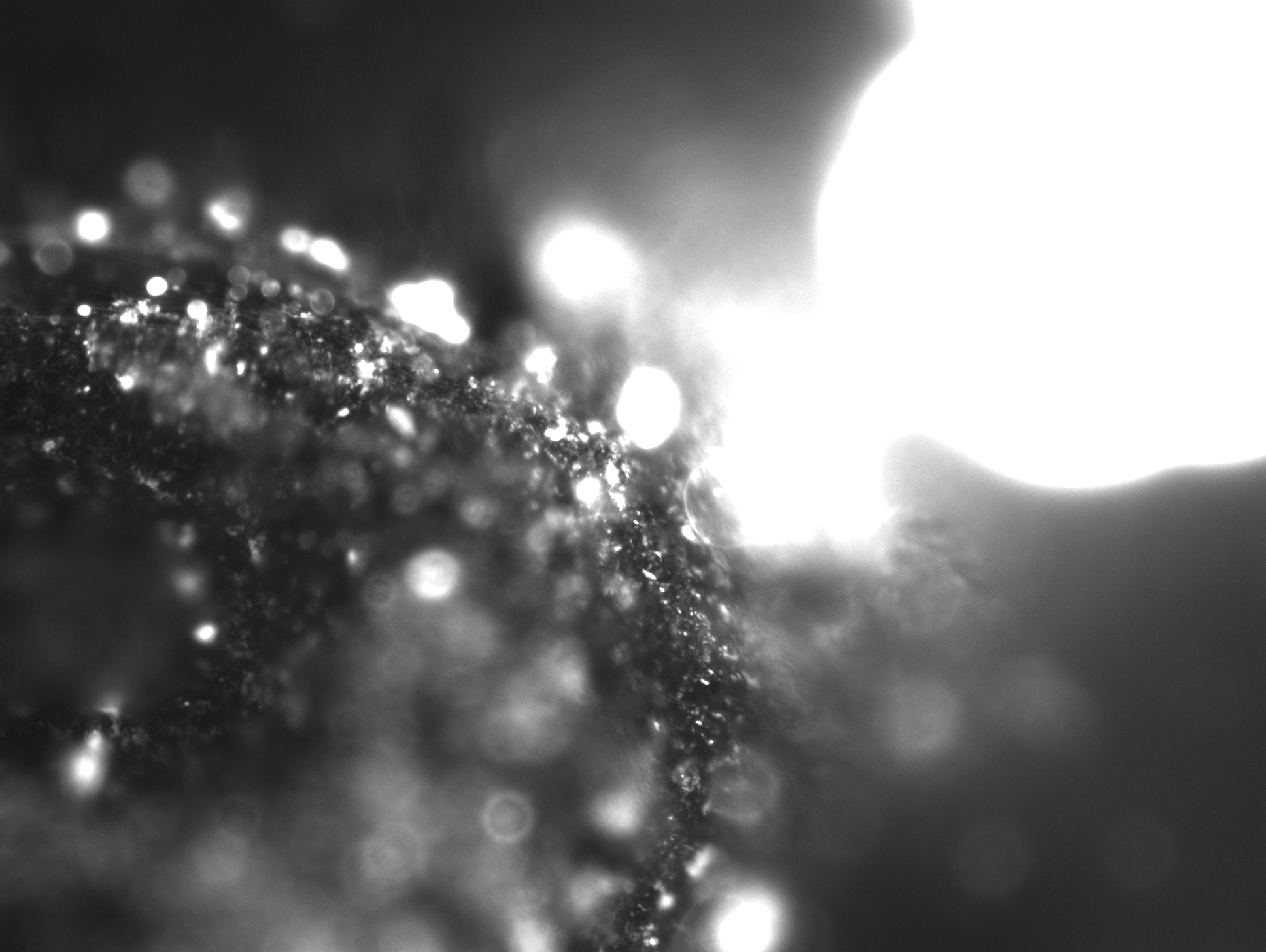

### 3_22.838-1.tif

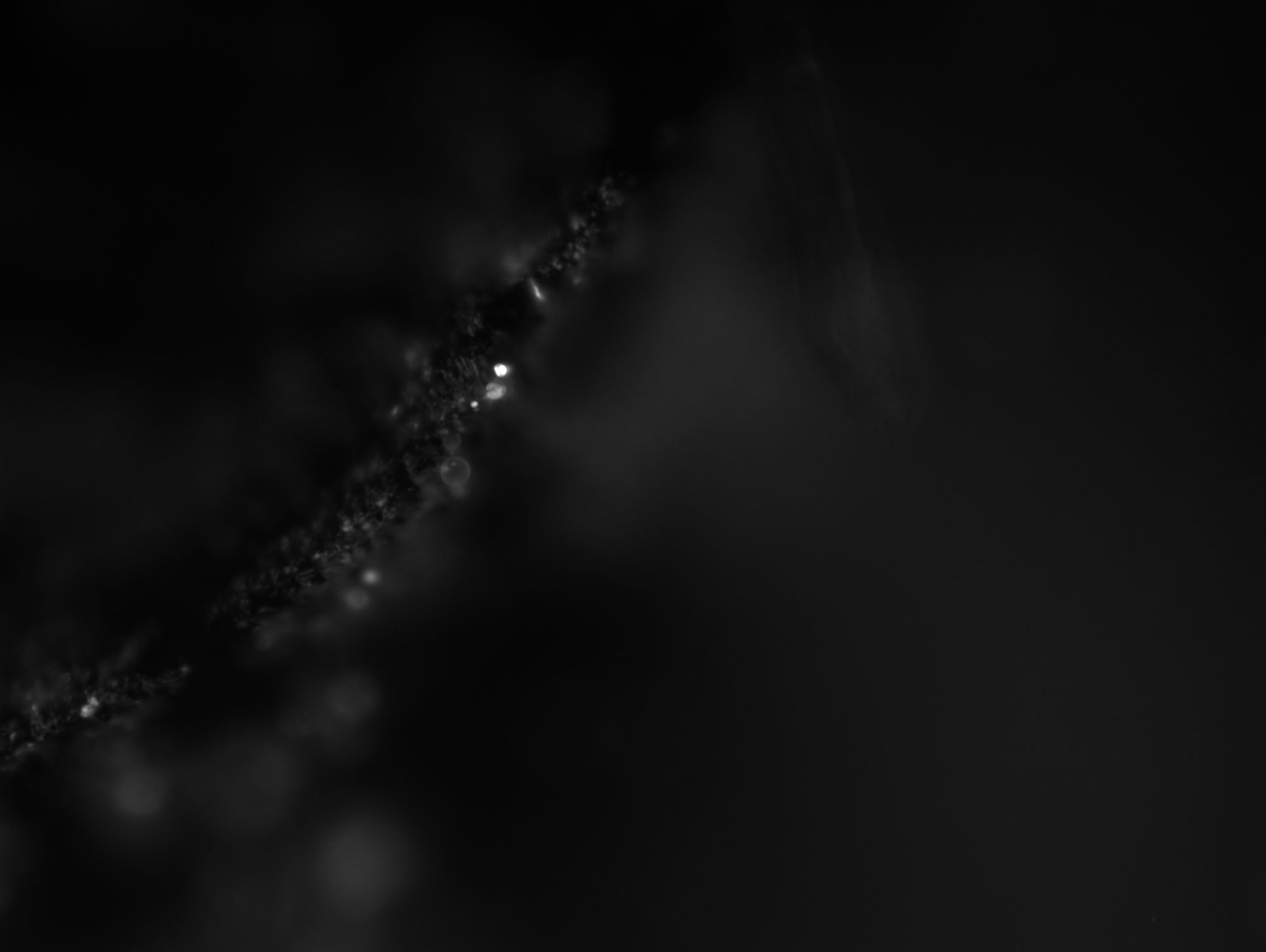

### 3_23_09.316.tif

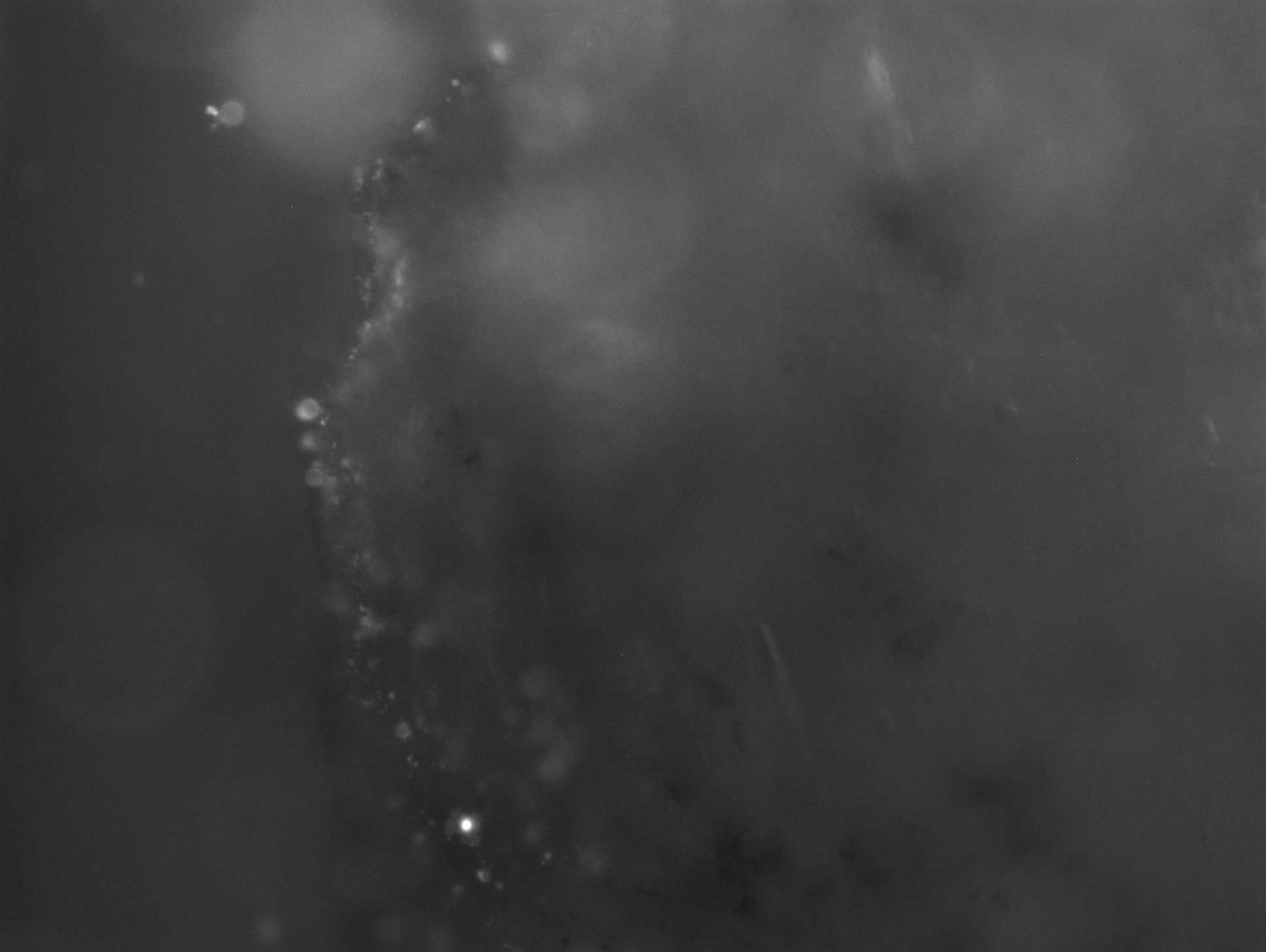

### 3_23_21.050.tif

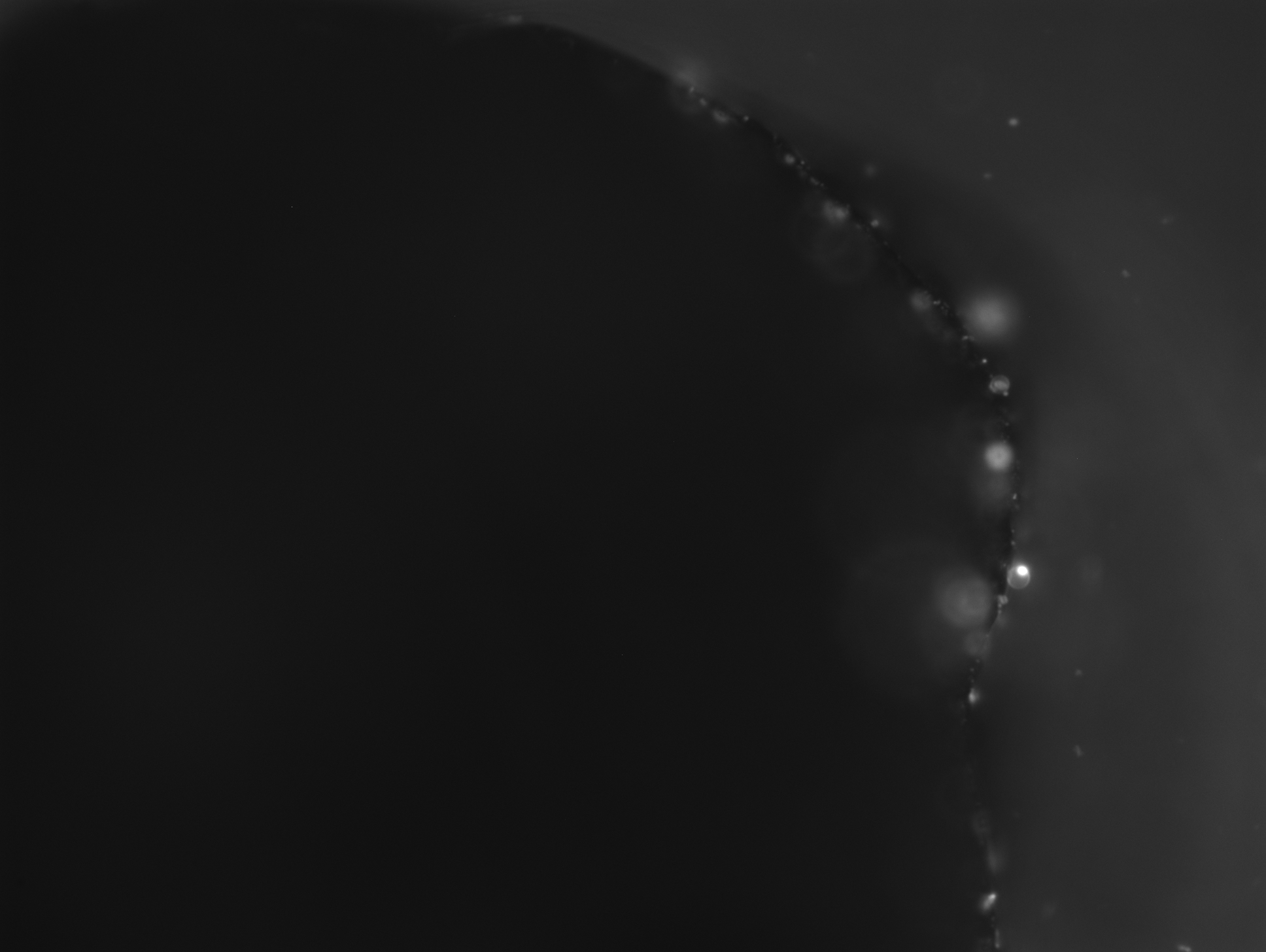

### 3_25_58.710.tif

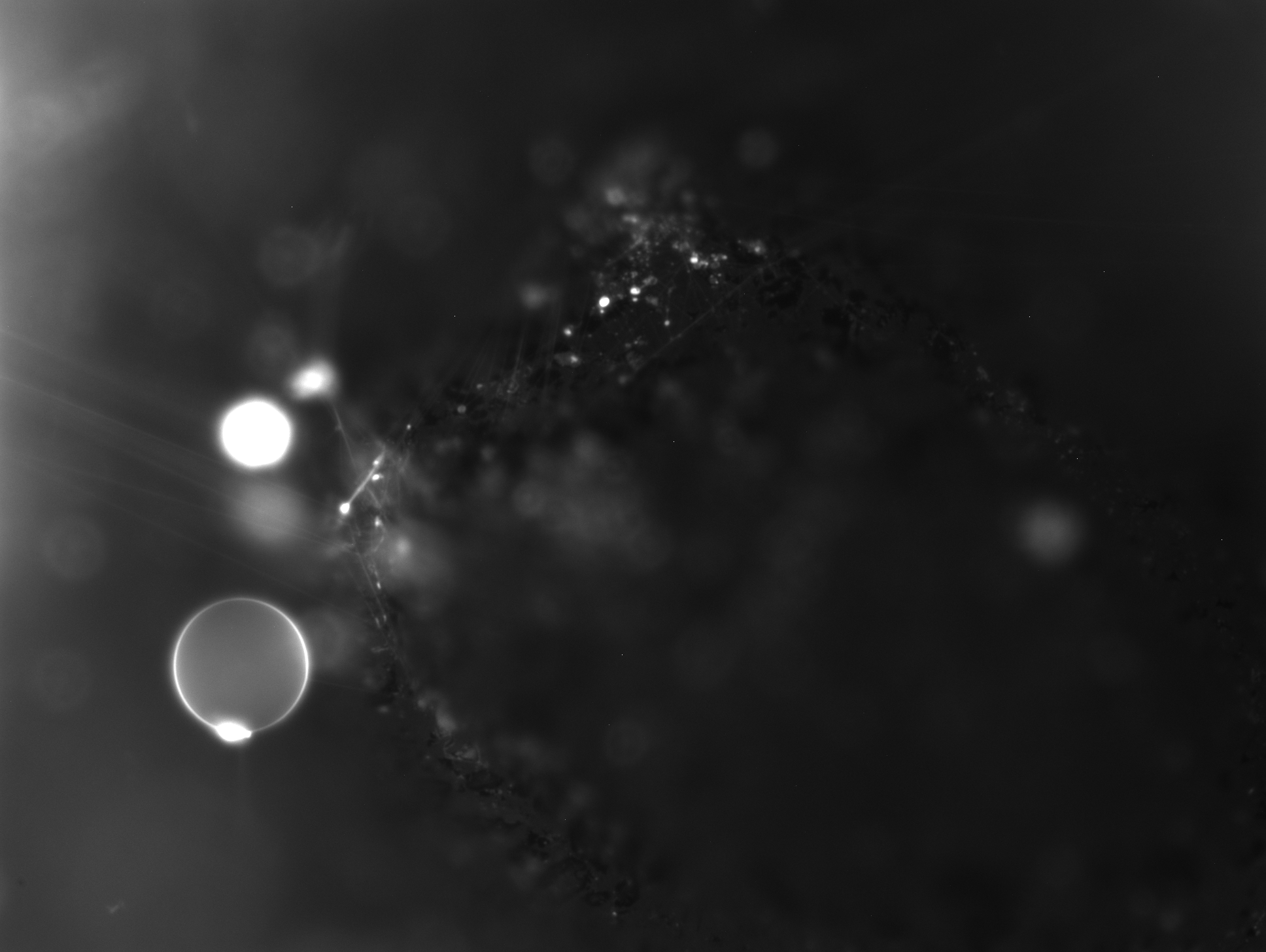

### 3_36_46.675.tif

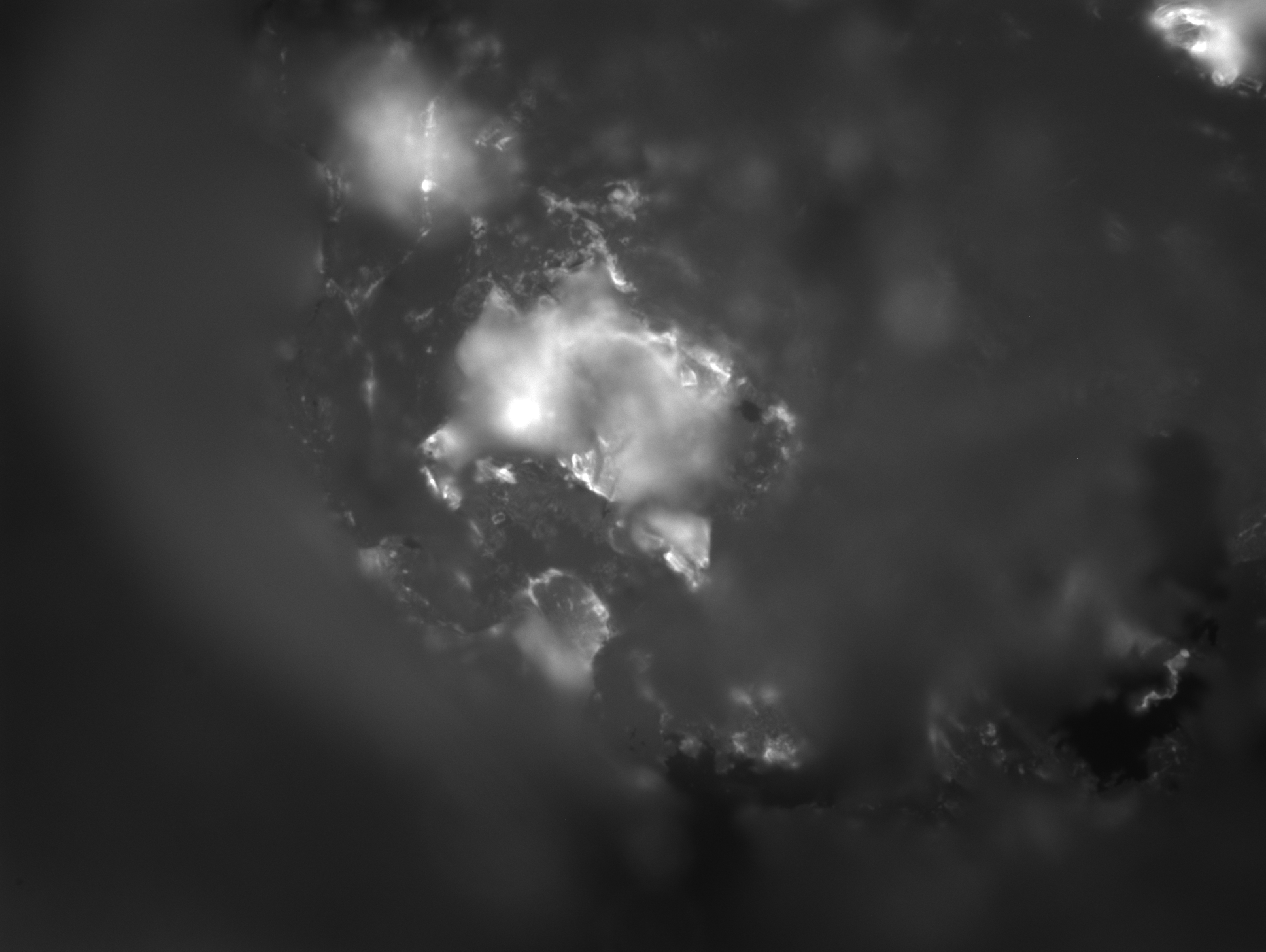

### 3_39_17.433.tif

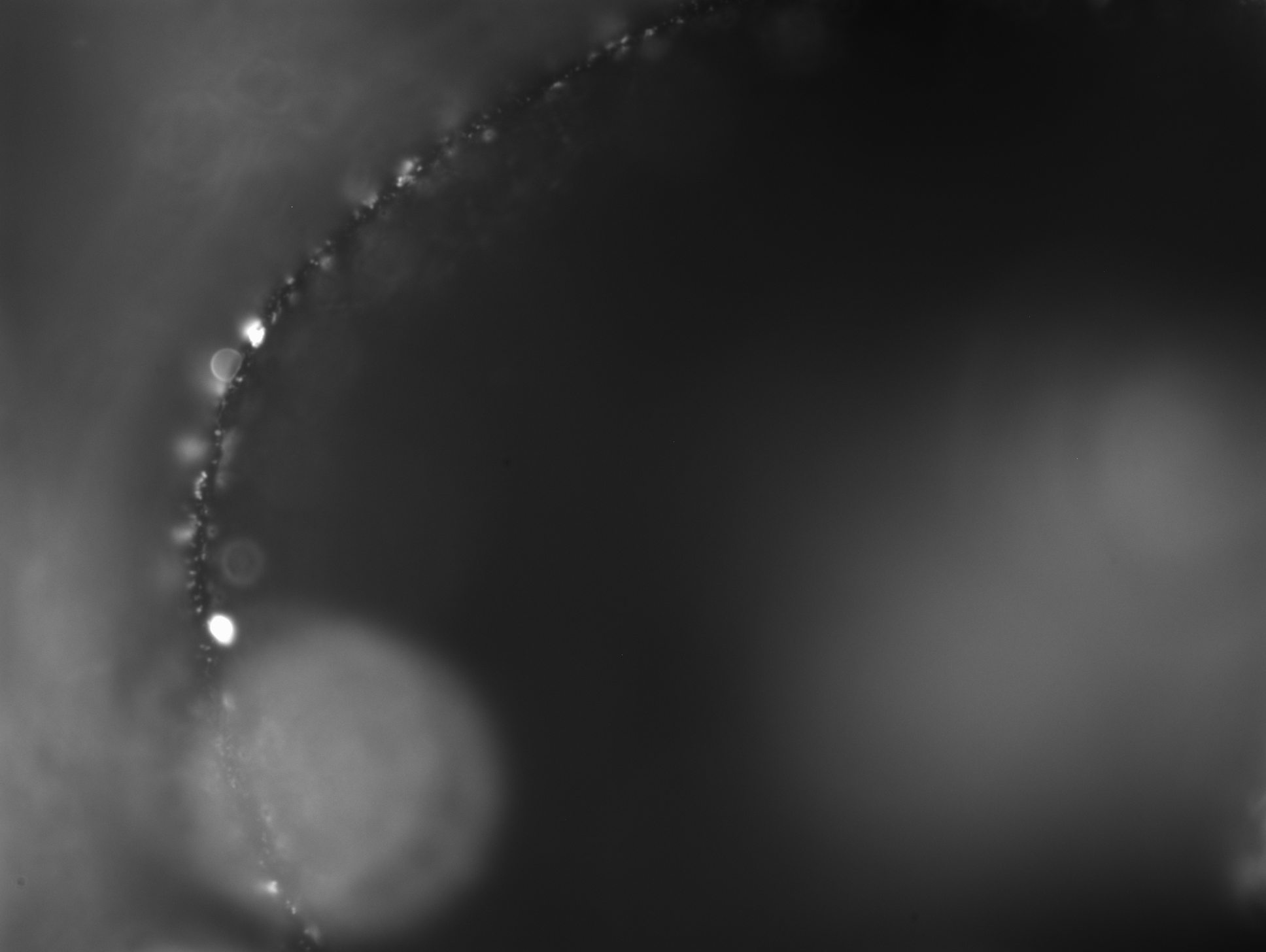

### 3_45_22.969.tif

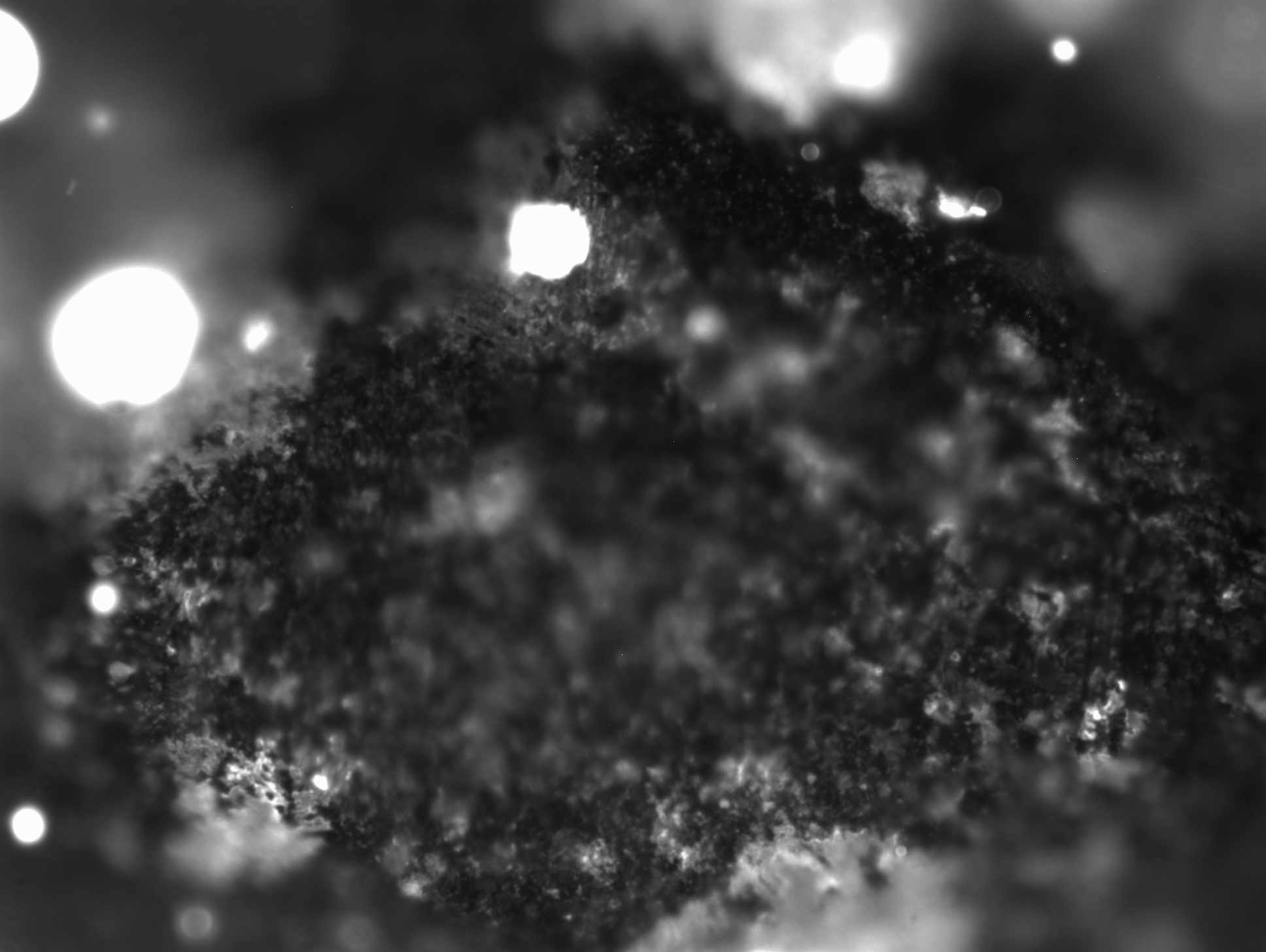

### 3_65_18.857.tif

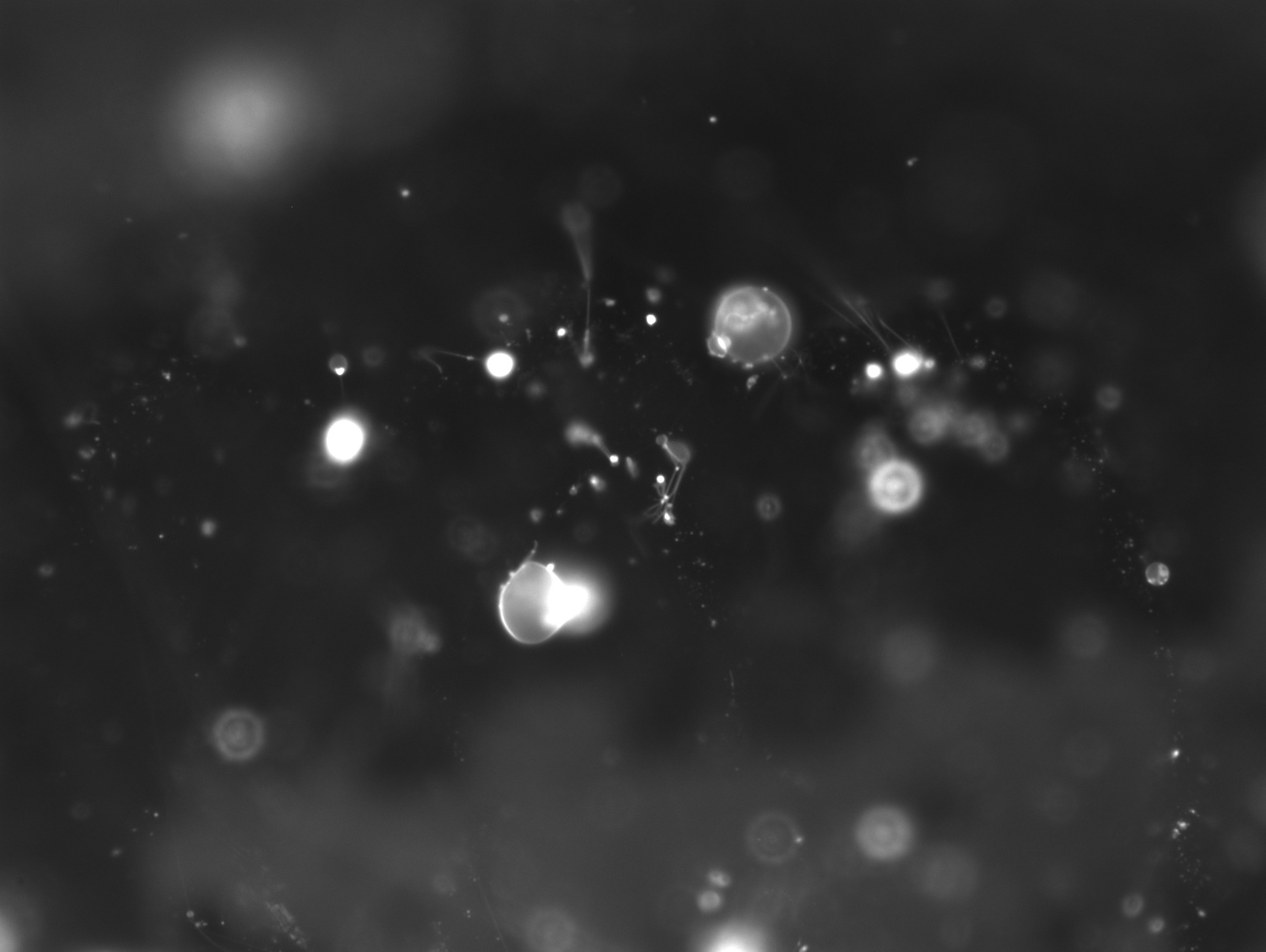

### 3_69_21.530.tif

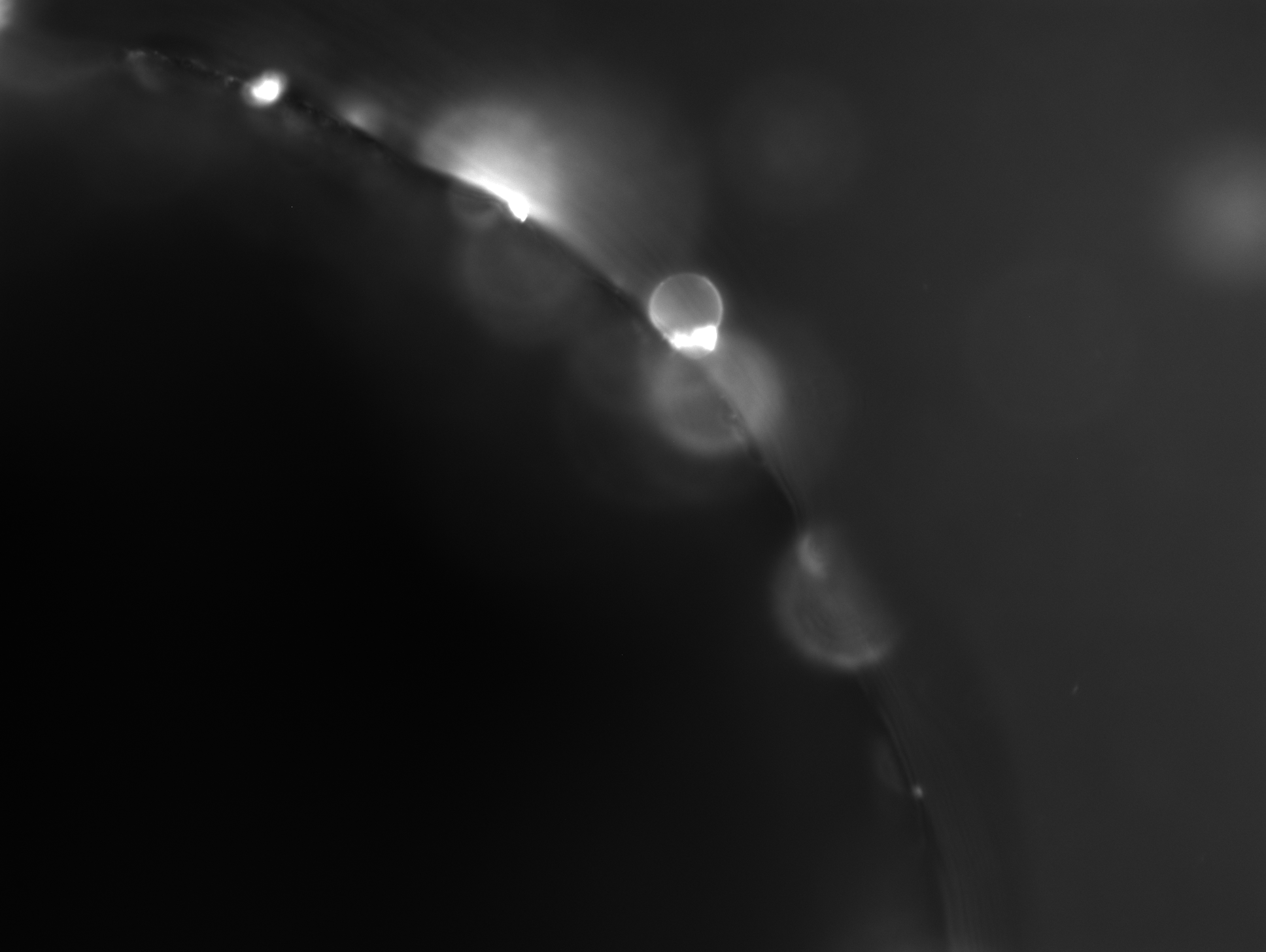

### 3_91_31.401.tif

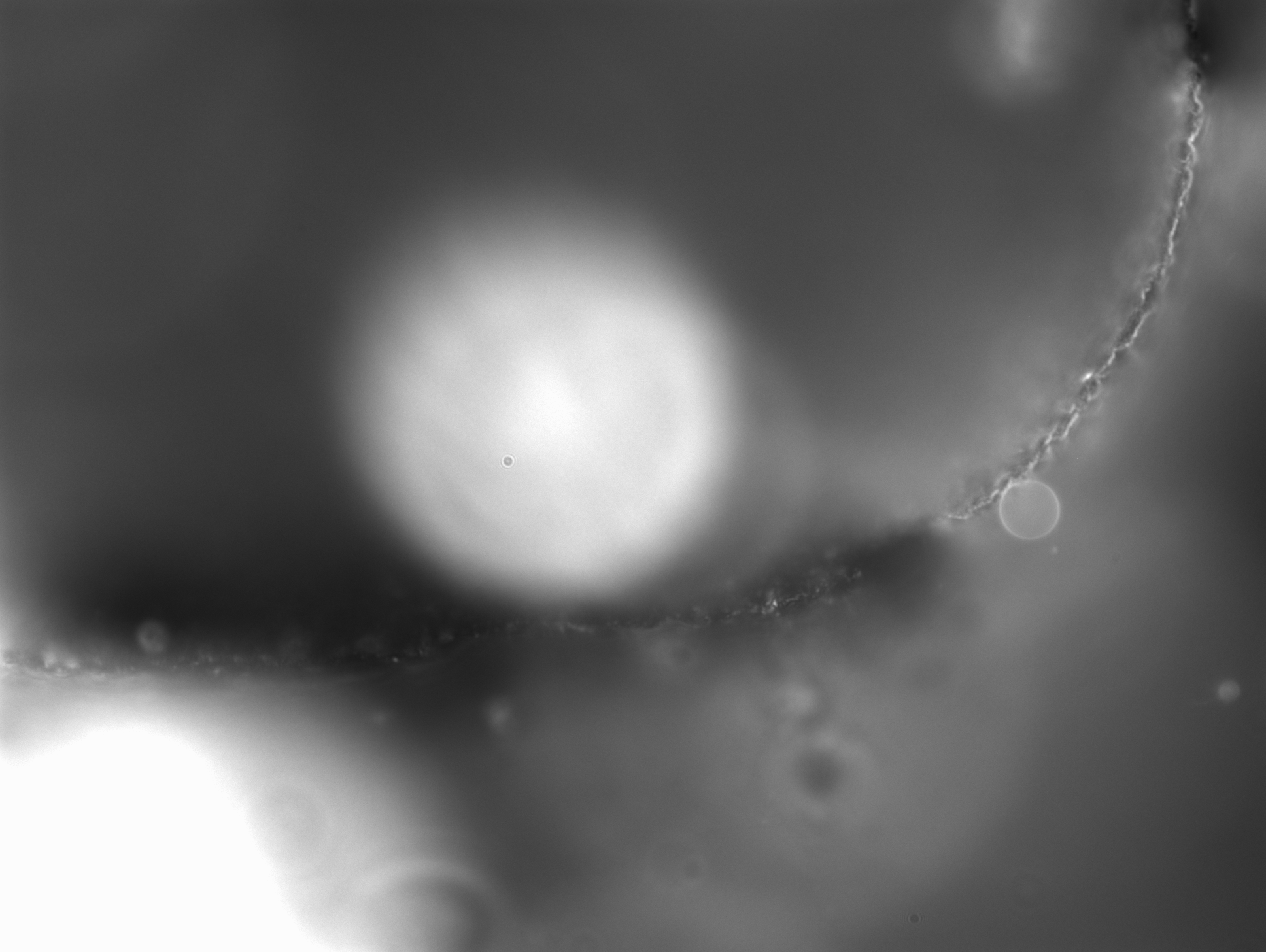

### 3_94_21.620.tif

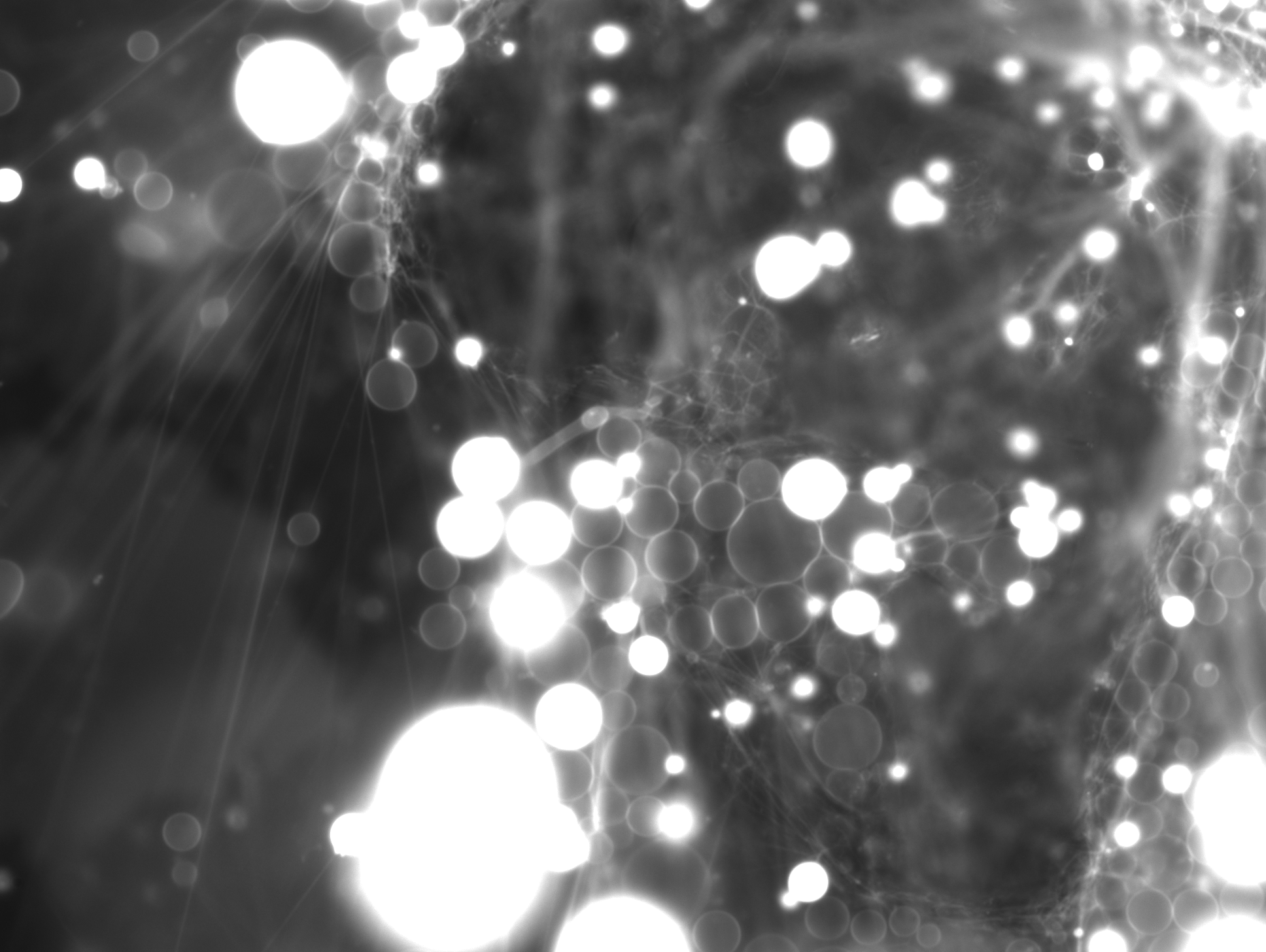

### 3_103_42.138.tif

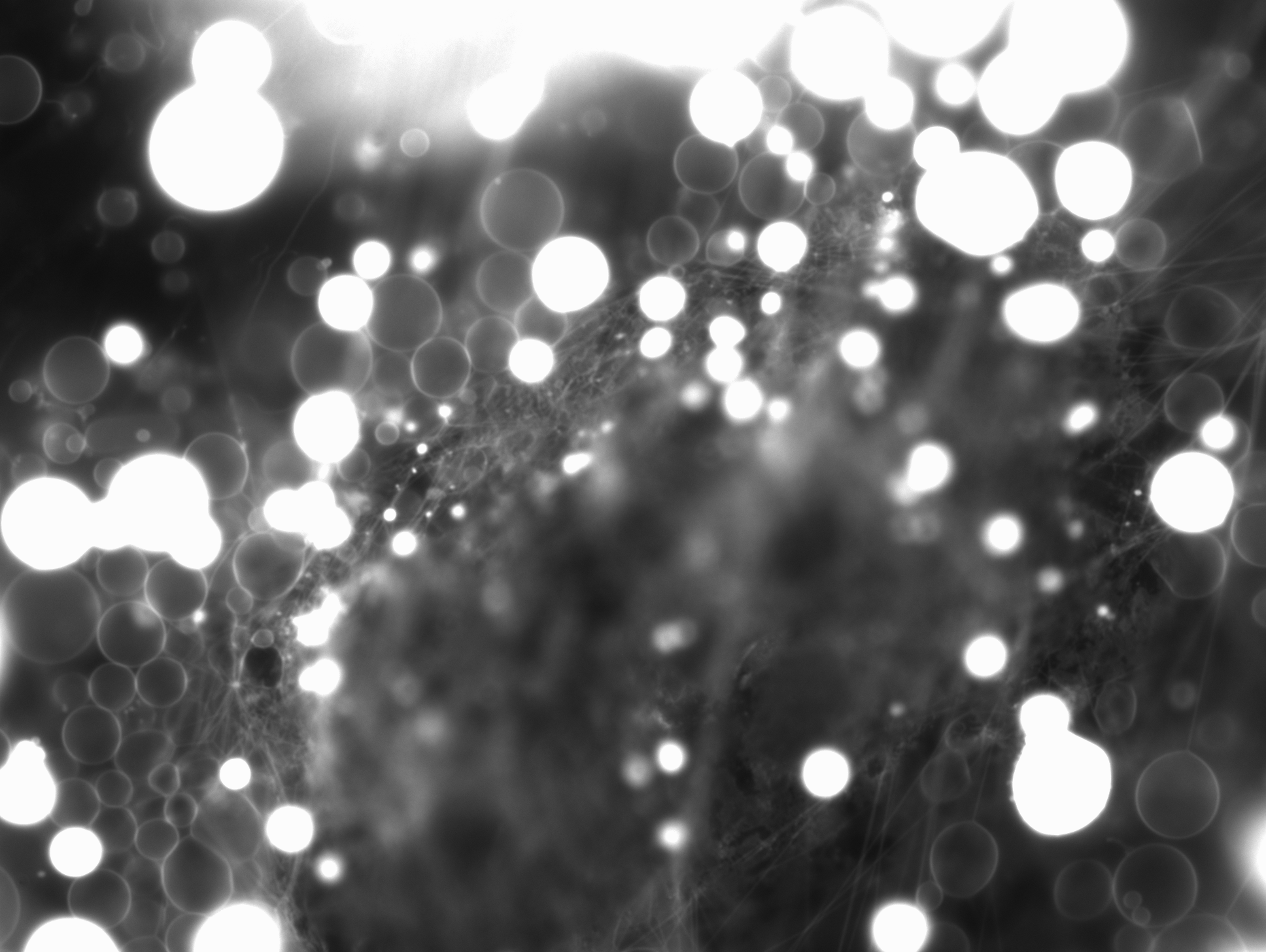

### 3_107_32.506.tif

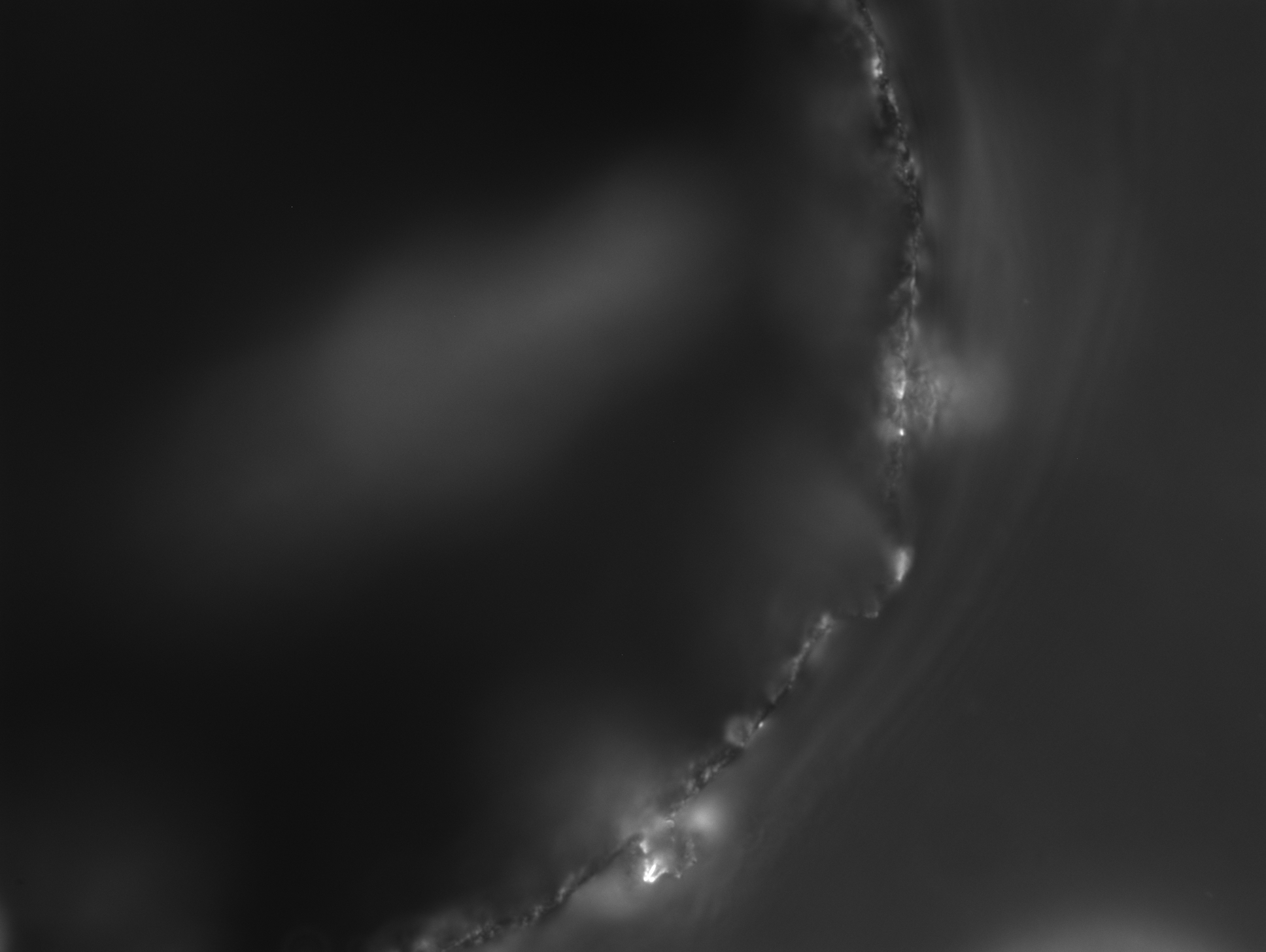

### 3_109_05.979.tif

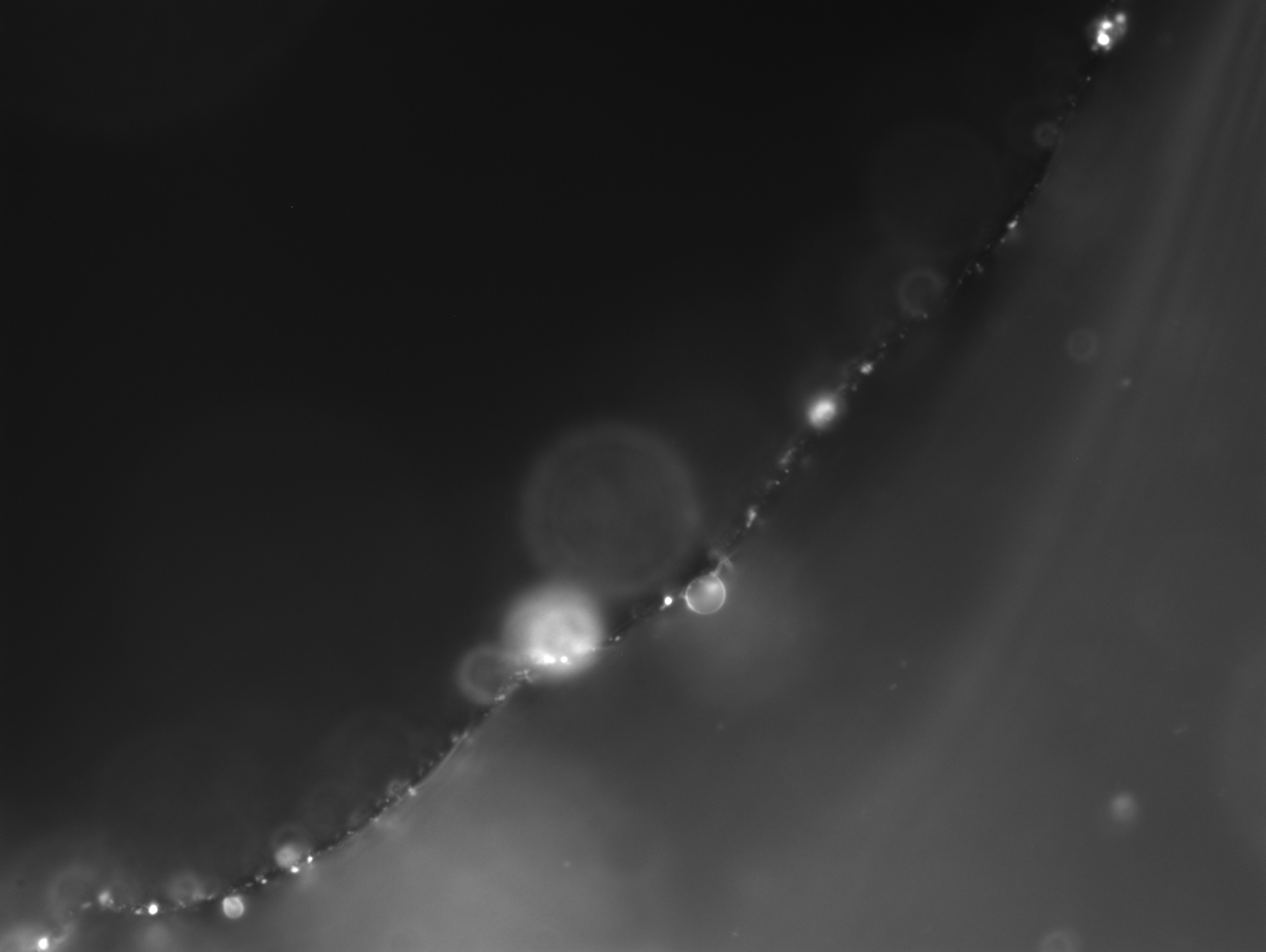

### 3_128_23.044.tif

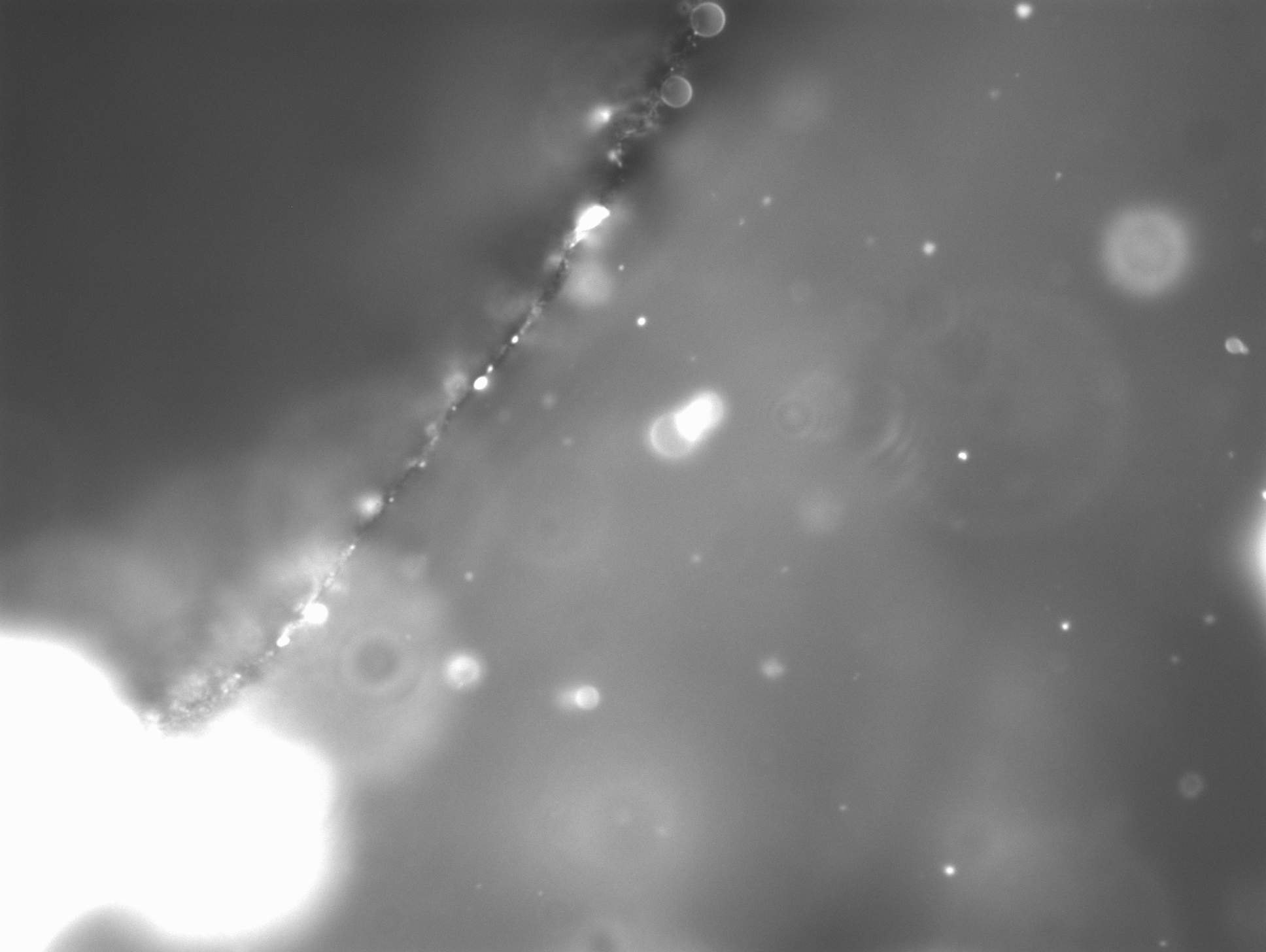

### 3_146_27.308.tif

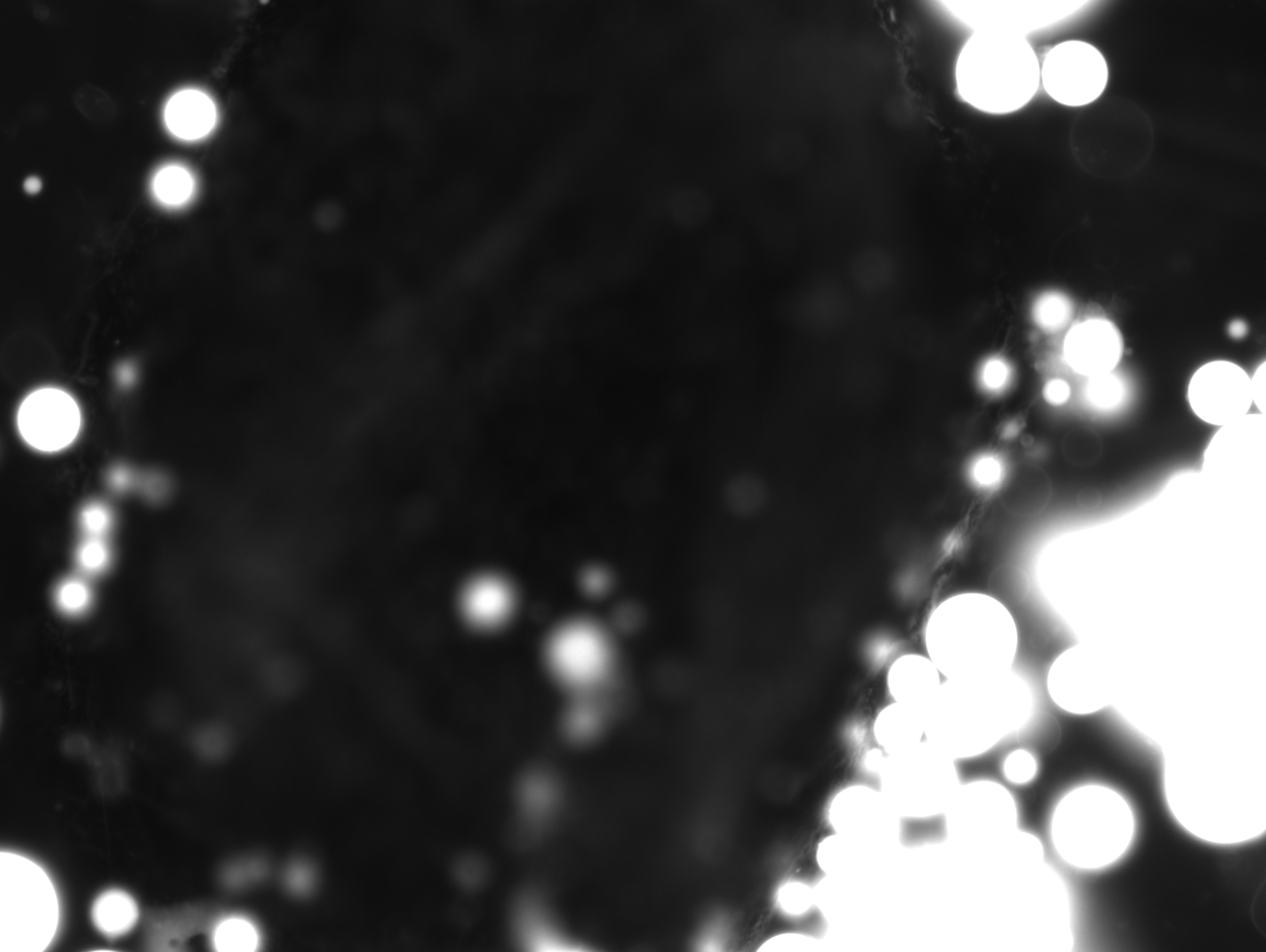

### 3_188_53.018.tif

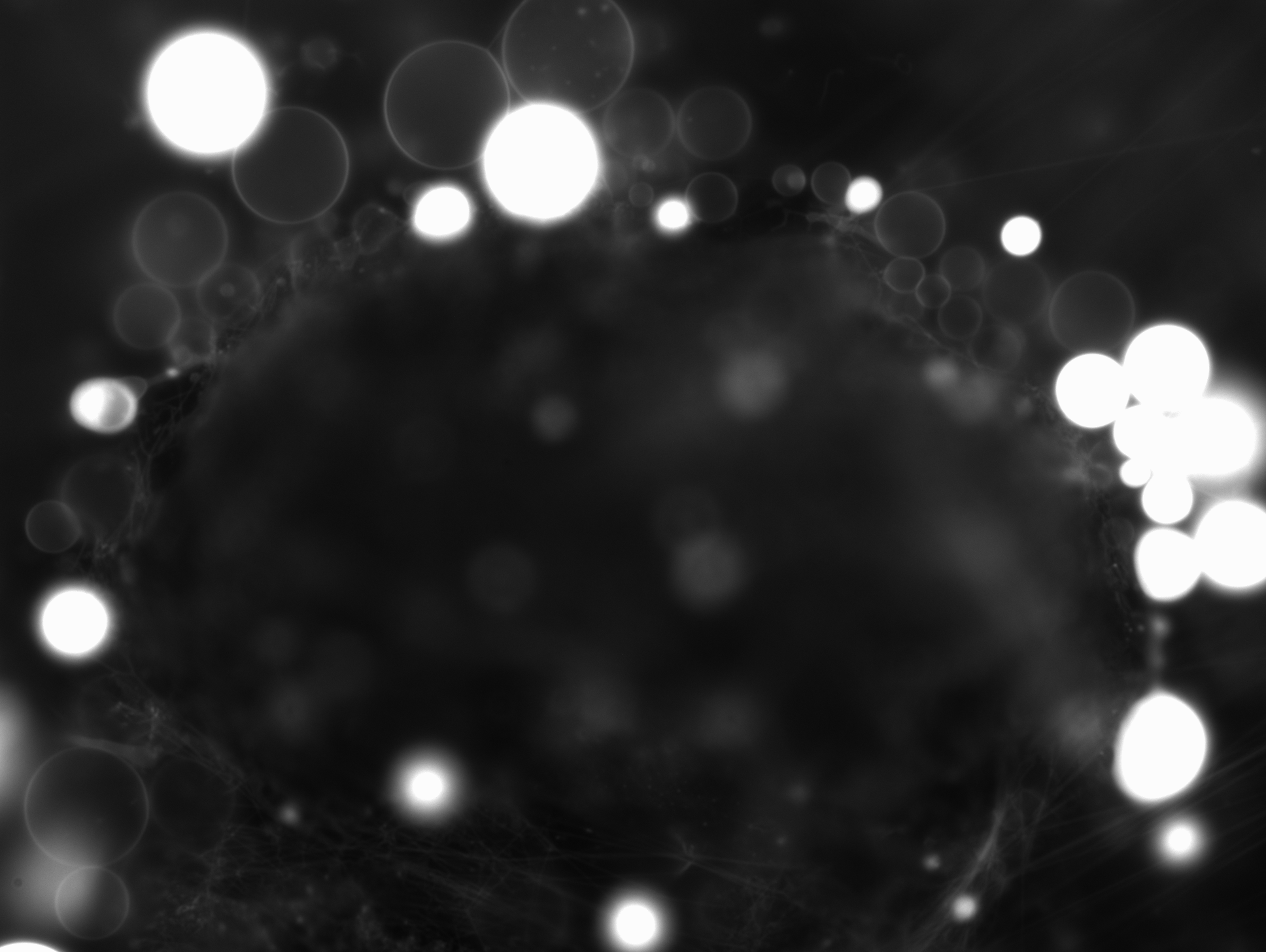

### 3_210_59.802.tif

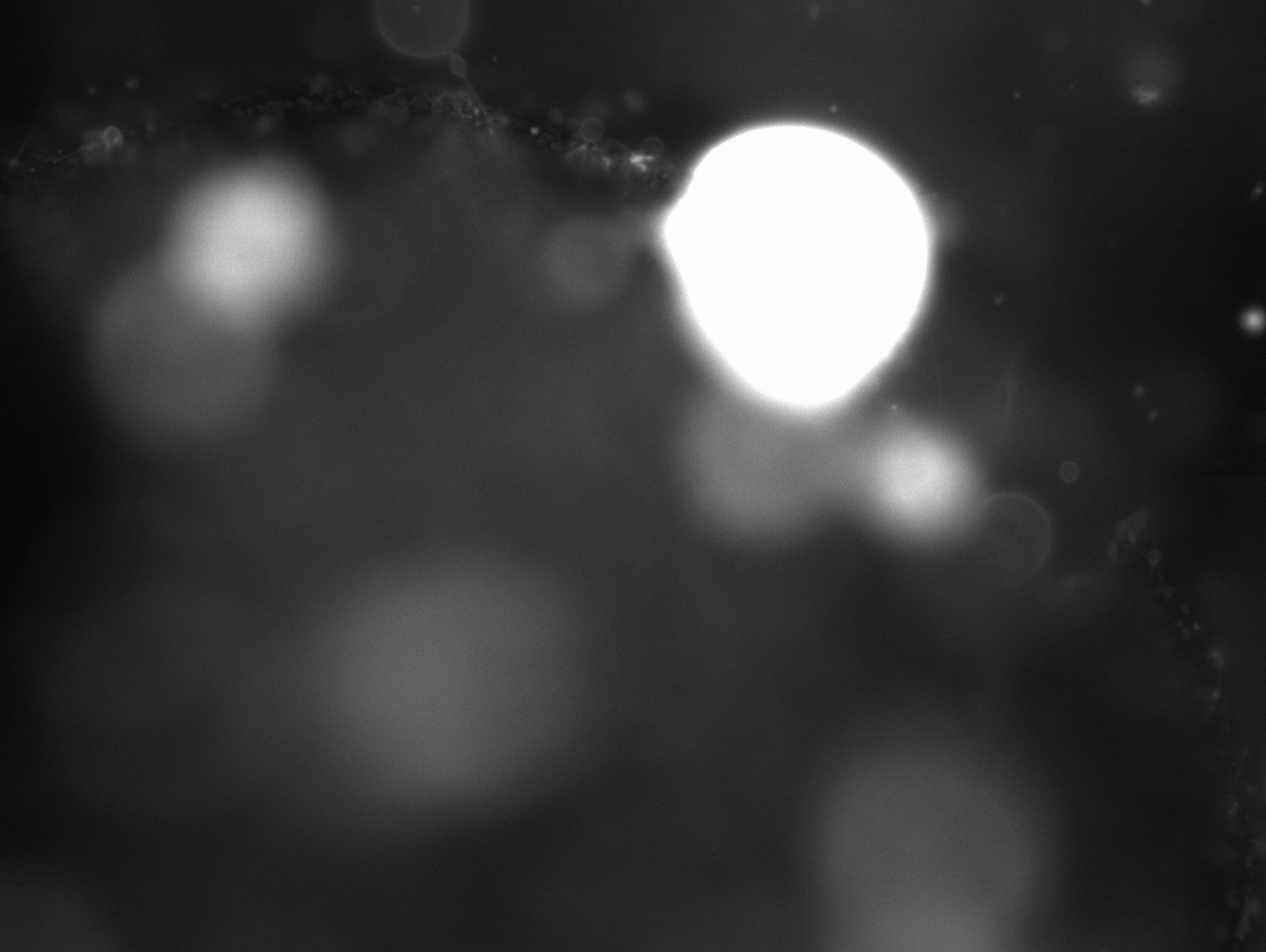

### 3_214_41.042.tif

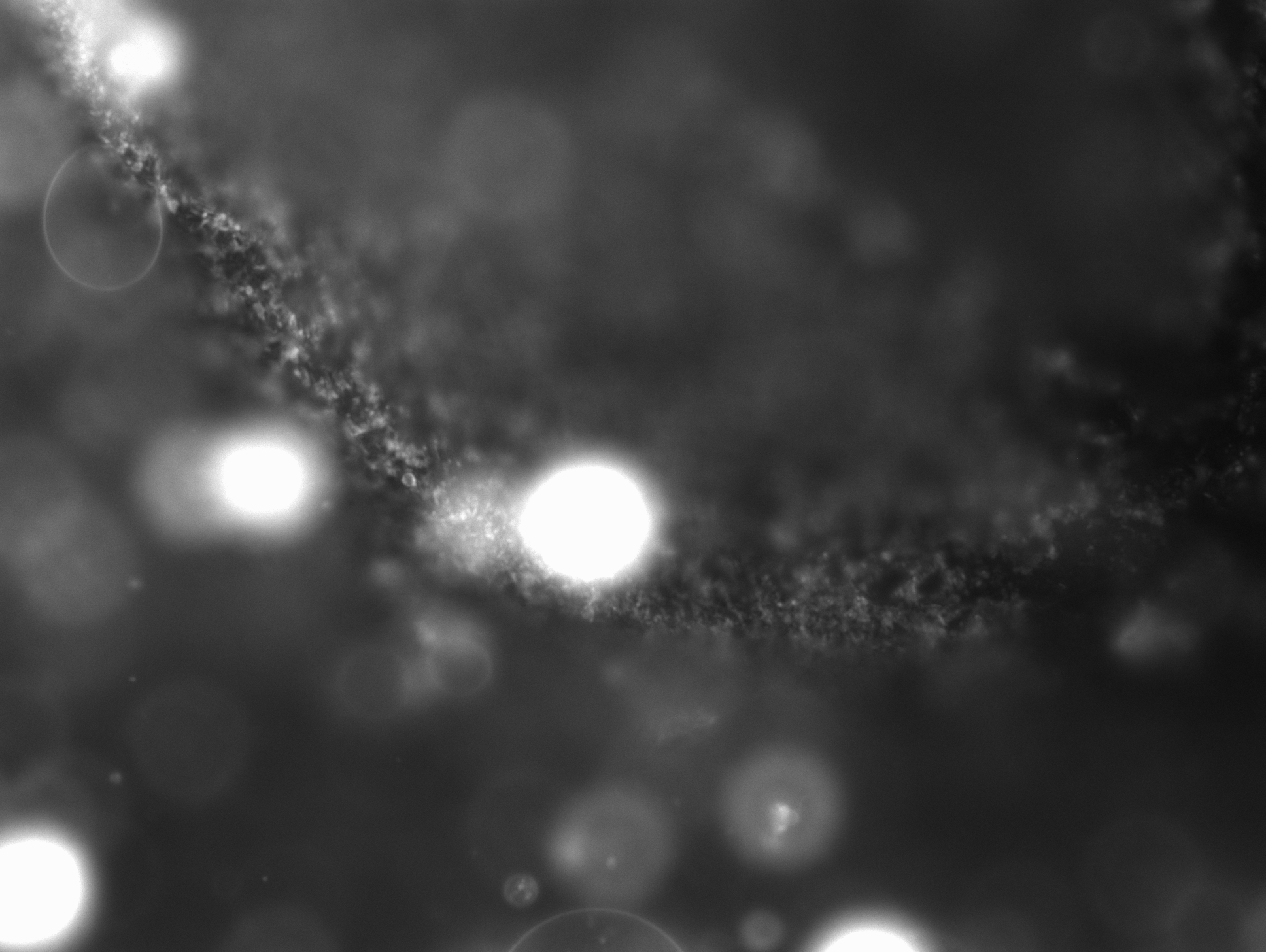

### 3_280_40.603.tif

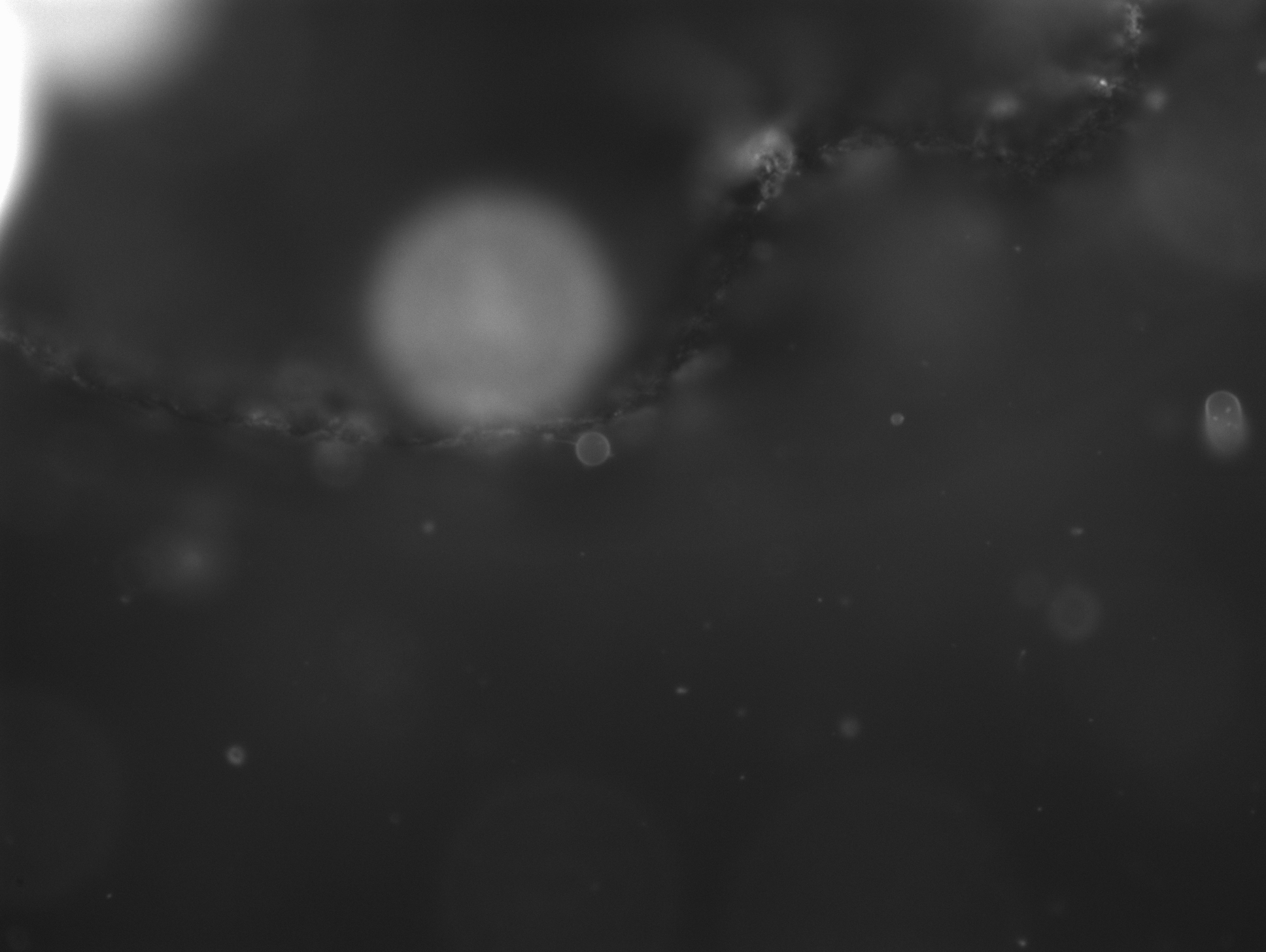

### 3_329_49.020.tif

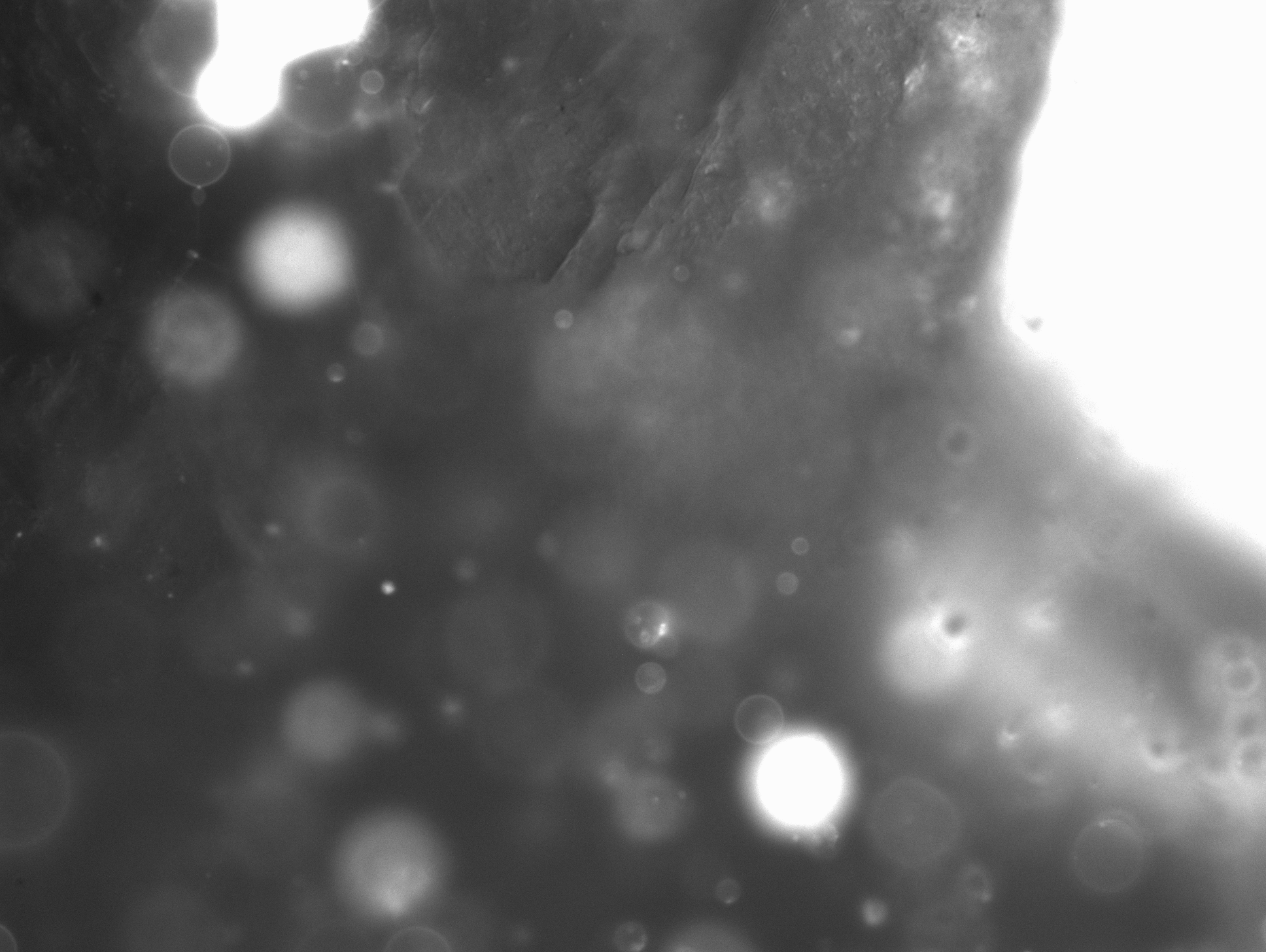

### 3_431_09.601.tif

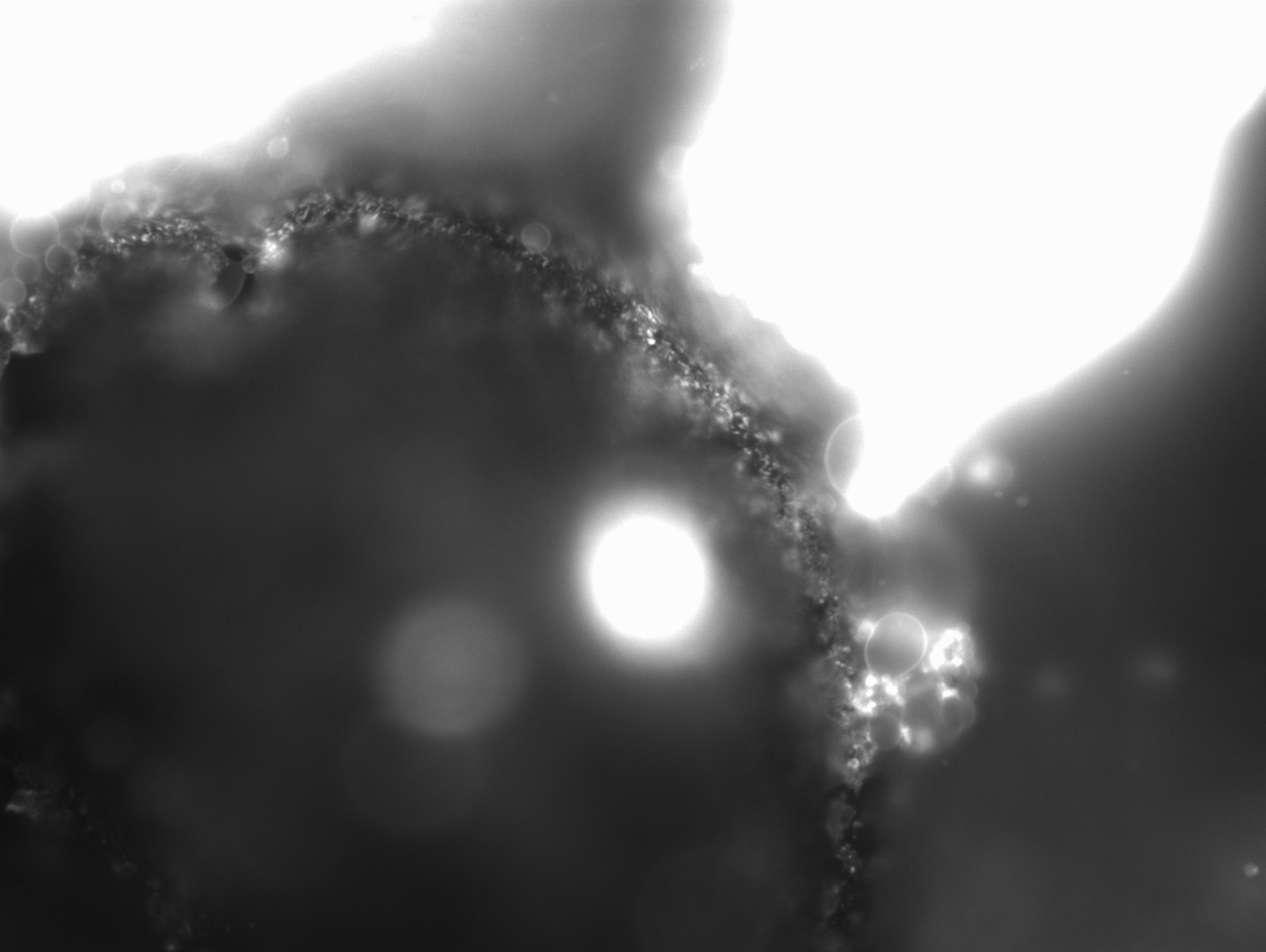

### 3_448_50.339.tif

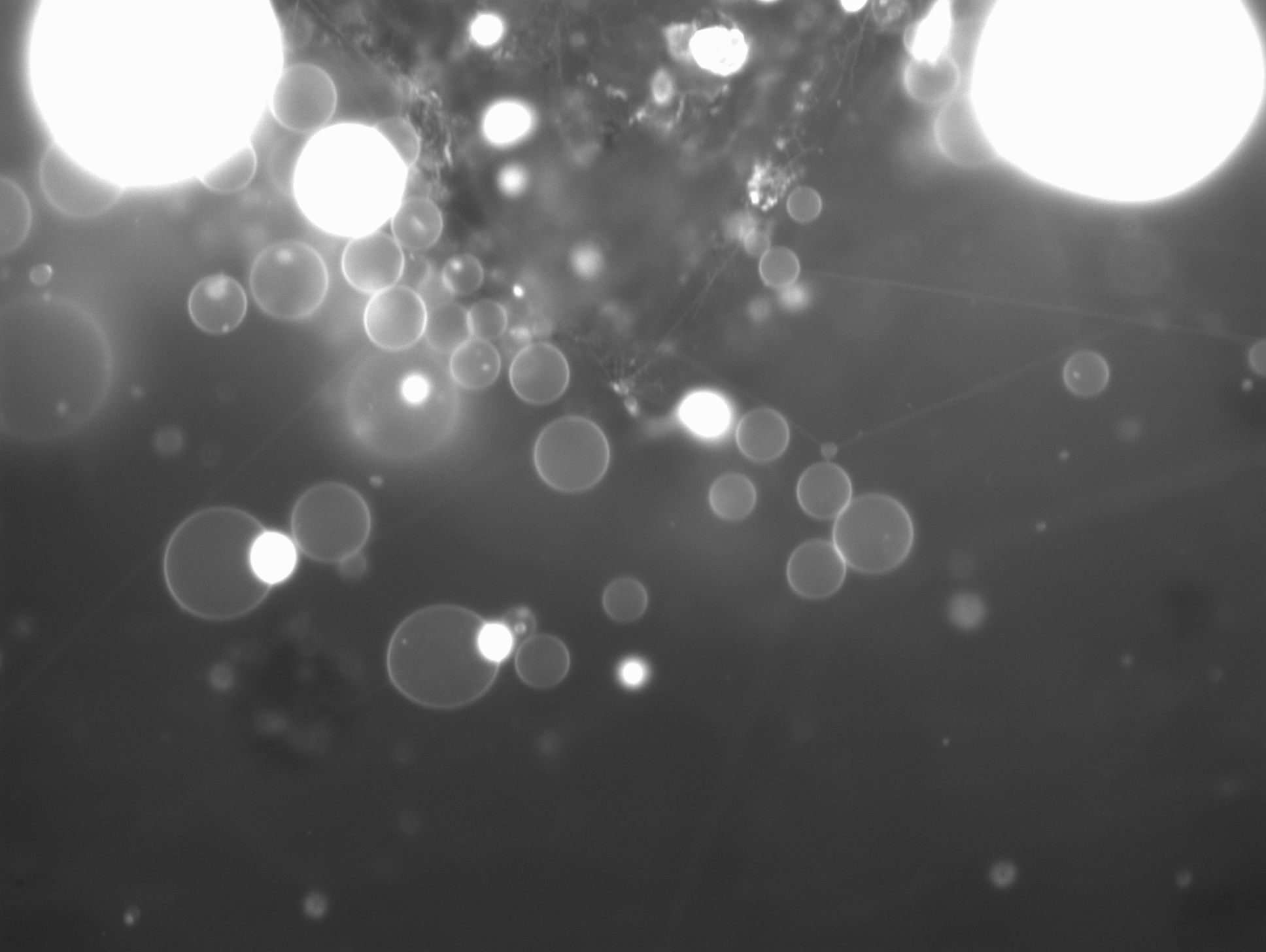

### 3_701_30.458.tif

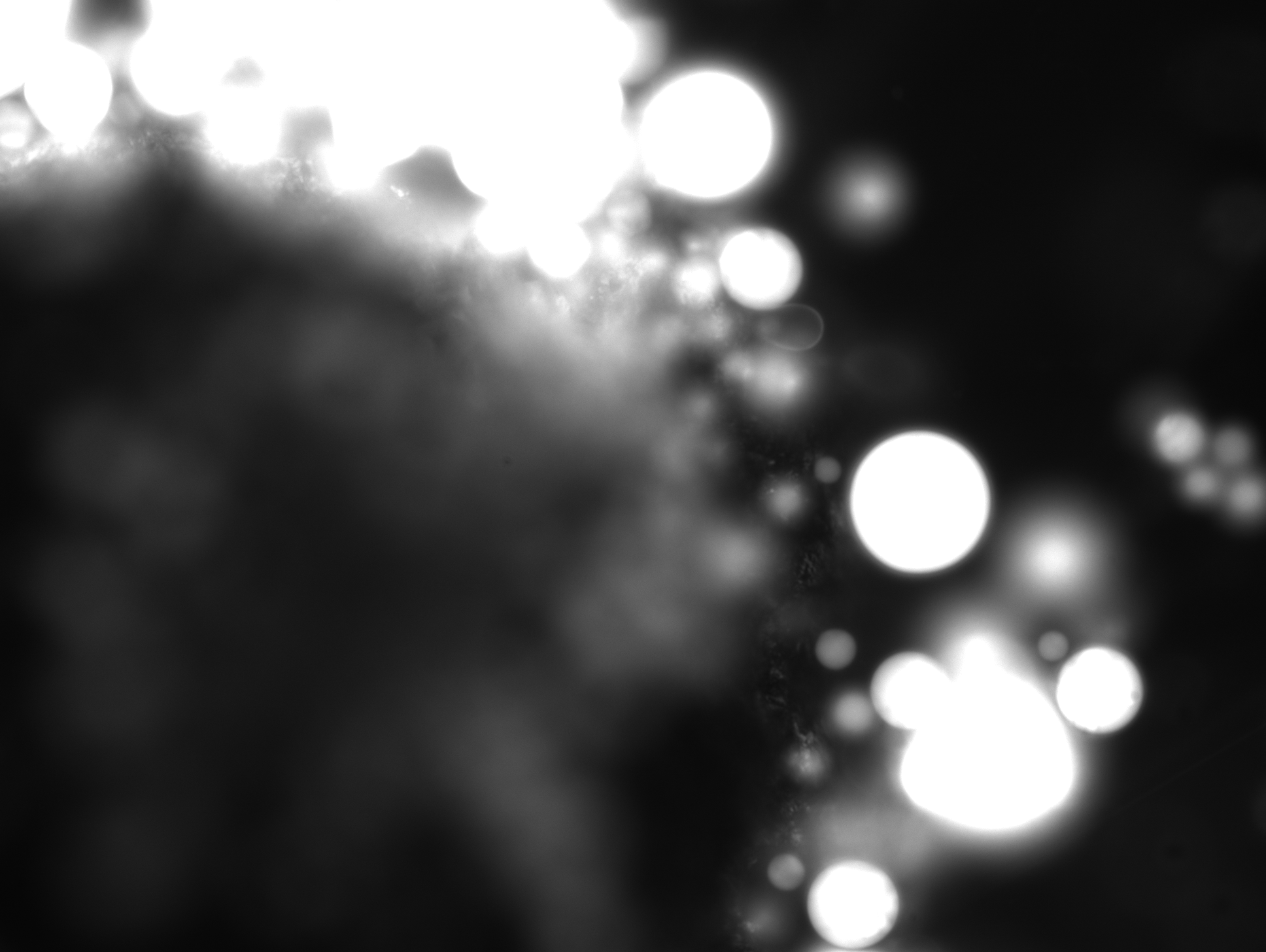

### 3_718_22.891.tif

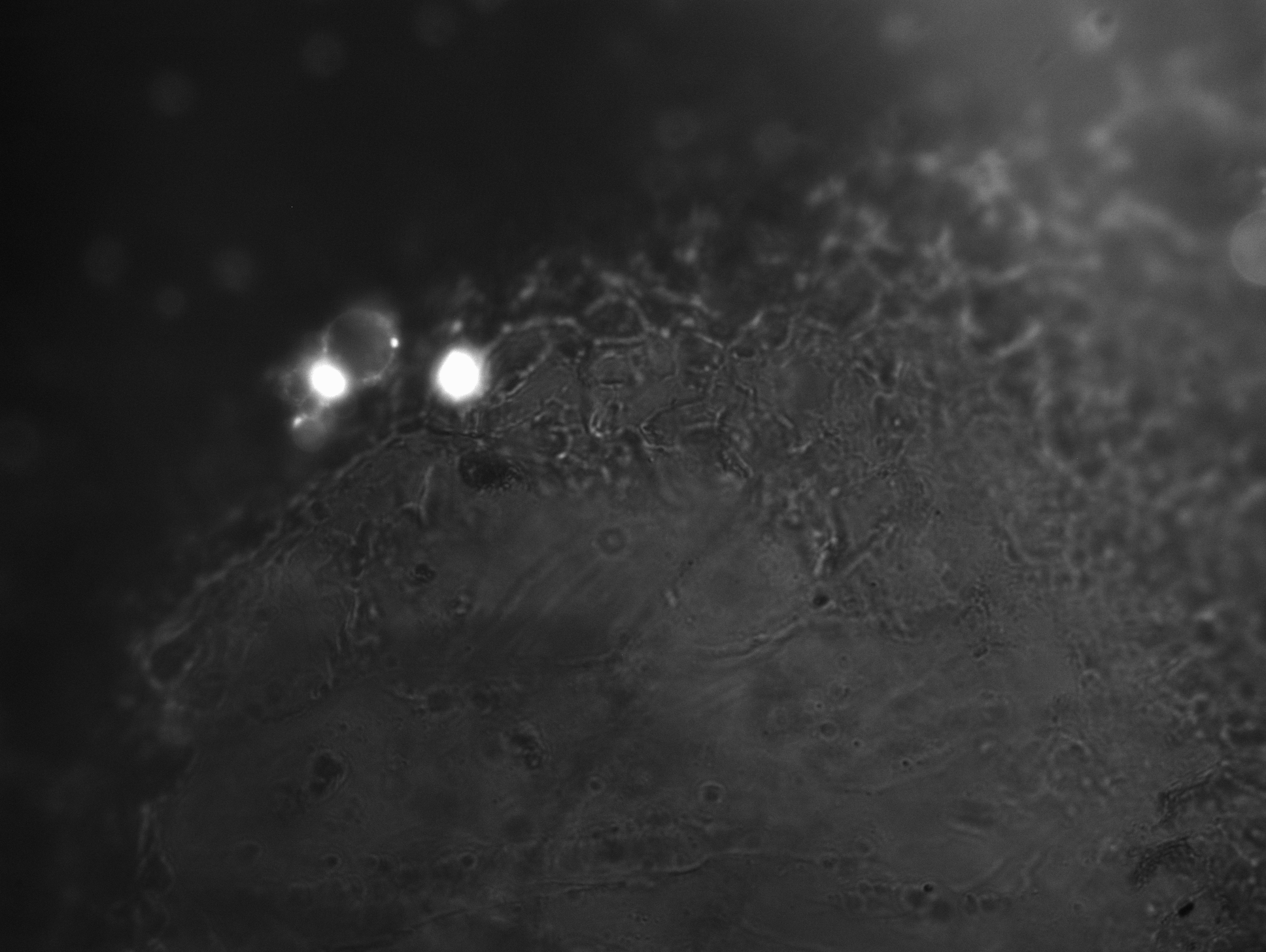

### 3_975_30.714.tif

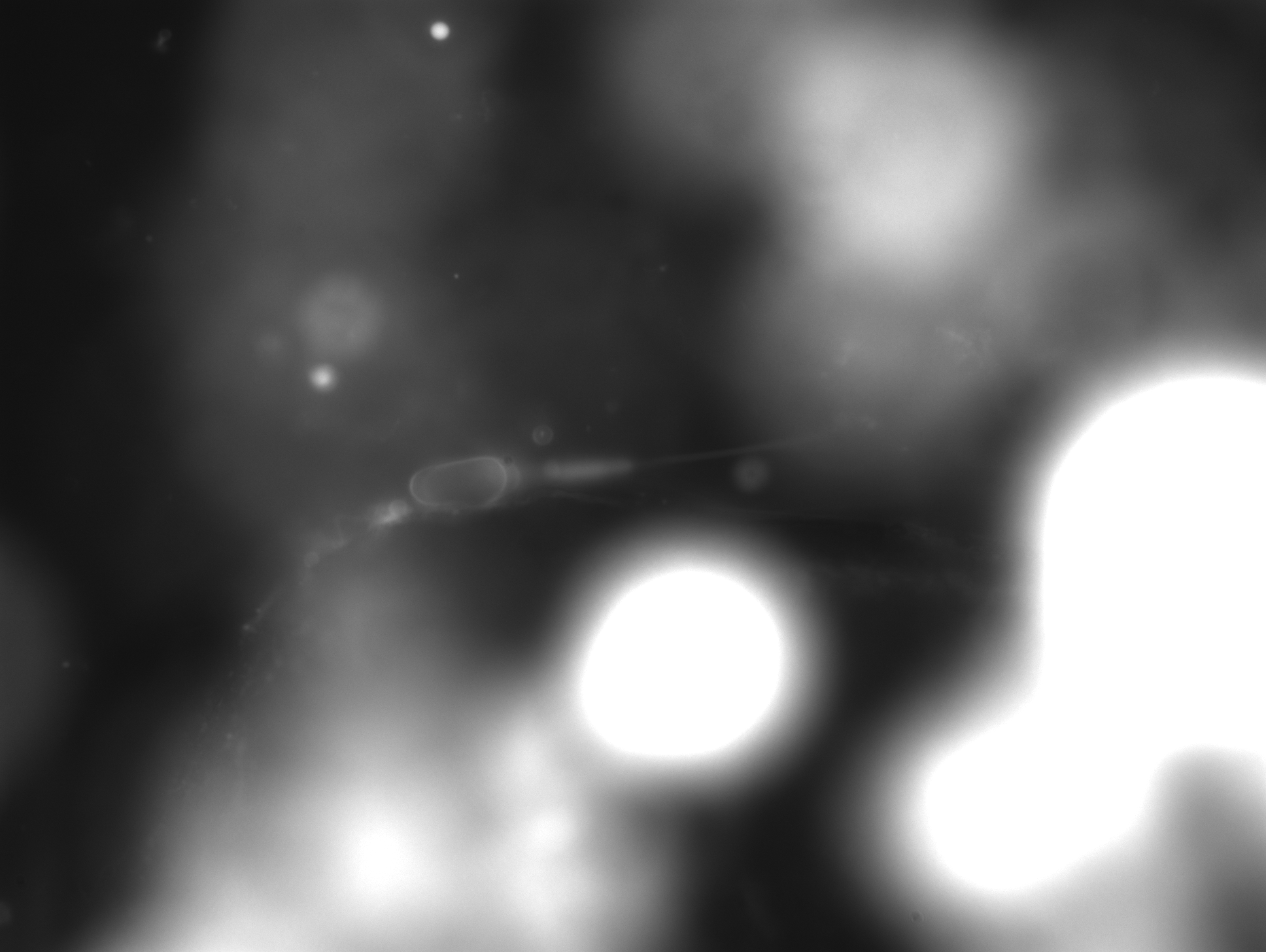
